# Evaluating genomic offset predictions in a forest tree with high population genetic structure

**DOI:** 10.1101/2024.05.17.594631

**Authors:** Juliette Archambeau, Marta Benito-Garzón, Marina de-Miguel, Alexandre Changenet, Francesca Bagnoli, Frédéric Barraquand, Maurizio Marchi, Giovanni G. Vendramin, Stephen Cavers, Annika Perry, Santiago C. González-Martínez

## Abstract

Predicting how tree populations will respond to climate change is an urgent societal concern. An increasingly popular way to make such predictions is the genomic offset (GO) approach, which aims to use genomic and climate data to identify populations that may experience climate maladaptation in the near future. More precisely, GO tries to represent the change in allele frequencies required to maintain the current gene-climate relationships under climate change. However, the GO approach has major limitations and, despite promising validation of its predictions using height data from common gardens, it still lacks broad empirical testing. In the present study, we evaluated the consistency and empirical validity of GO predictions in maritime pine (*Pinus pinaster* Ait.), a tree species from southwestern Europe and North Africa with a marked population genetic structure. First, gene-climate relationships were estimated using 9,817 SNPs genotyped in 454 trees from 34 populations; and candidate SNPs potentially involved in climate adaptation were identified. Second, GO was predicted using four methods, namely Gradient Forest (GF), Redundancy Analysis (RDA), latent factor mixed model (LFMM) and Generalised Dissimilarity Modeling (GDM), two sets of SNPs (candidate and control SNPs) and five climate general circulation models (GCMs) to account for uncertainty in future climate predictions. Last, the empirical validity of GO predictions was evaluated within a Bayesian framework by estimating the associations between GO predictions and two independent data sources: mortality data from National Forest Inventories (NFI), and mortality and height data from five common gardens in contrasting environments. We found high variability in GO predictions across methods, SNP sets and GCMs. Regarding validation, GO predictions with GDM and GF (and to a lesser extent RDA) based on the candidate SNPs showed the strongest and most consistent associations with mortality rates in common gardens and NFI plots. We found almost no association between GO predictions and tree height in common gardens, most likely due to the overwhelming effect of population genetic structure on tree height in this species. Our study demonstrates the imperative to validate GO predictions with a range of independent data sources before they can be used as informative and reliable metrics in conservation or management strategies.

## 1 Introduction

Mounting evidence shows that anthropogenic climate change is already affecting biodiversity (Parmesan and Yohe 2003), exposing populations and species to unprecedented risks of decline and extinction (Urban 2015, Wiens 2016). By shifting fitness optima, climate change increases the mismatch between the current phenotypes of well-adapted populations and their environment, thereby decreasing the average fitness of the populations (i.e., scenario of the moving target in Brady et al. 2019a). The consequent maladaptation can be counterbalanced by sufficient and appropriate adaptive variation, which can enable rapid evolution towards the new phenotypic optimum (Brady et al. 2019b). However, if the extent and rate of climate change are too severe, the demographic costs of maladaptation might be too high relative to the capacity and speed of adaptation, and populations will be pushed toward extinction (Bürger and Lynch 1995, Gomulkiewicz and Holt 1995, Chevin et al. 2010, Uecker et al. 2014).

To identify populations at risk of short-term maladaptation under climate change, Fitzpatrick and Keller (2015) developed the concept of genomic offset (or genetic offset; GO). They modeled the non-linear turnover in allele frequencies along climatic gradients with two approaches from community-level modeling, namely Gradient Forest (GF; Ellis et al. 2012) and Generalised Dissimilarity Modeling (GDM; Ferrier et al. 2007). They then calculated GO as the distance between the current genomic composition of the populations and the genomic composition required to maintain the estimated gene-climate relationships (Fitzpatrick and Keller 2015). Since then, GO has been estimated with a variety of other methods including Redundancy Analysis (RDA; Capblancq and Forester 2021), the risk of non adaptedness (RONA; Rellstab et al. 2016) and latent factor mixed models (LFMM; Gain and François 2021). As they are straightforward to implement (genomic and climatic data from several populations are sufficient) and seemingly easy to interpret, these methods have already been applied to a wide range of species, and GO predictions are now often recommended to guide management and conservation strategies (e.g., Rhoné et al. 2020, Lachmuth et al. 2023, Yuan et al. 2023).

In this context, it is crucial to emphasize that GO is not informative about the capacity of populations to adapt, i.e., to evolve towards the new fitness optima of future climates, and therefore captures only part of the vulnerability of populations to climate change (IPCC 2007, Foden et al. 2019). Moreover, the GO concept relies on strong assumptions that undermine the robustness of its predictions (Rellstab et al. 2021, Ahrens et al. 2023). One key assumption is that populations are currently optimally adapted to their local environment, which is often violated. For instance, populations of temperate and boreal tree species at the northern edges of their distribution would benefit from a temperature increase, at least in the short term (Rehfeldt et al. 2002, 2003, Savolainen et al. 2007, Pedlar and McKenney 2017, Rehfeldt et al. 2018, Fréjaville et al. 2020). Another key assumption is that populations from different geographic locations but occupying similar environments have the same adaptive alleles; but the same phenotypes can be produced by multiple different combinations of genetic variants (i.e., ’genotypic redundancy’; Láruson et al. 2020) and how populations evolve towards the fitness optimum depends on the shape of the adaptive landscape (Lotterhos 2023). GO predictions are thus impacted by trait genomic architecture and the environmental landscape, as demonstrated in a simulation study based on the GF method (Láruson et al. 2022). The study also showed that the predicted rate of allele frequency change did not reflect the steepness of adaptive clines, when simulated as linear, although it did capture the extent to which the environmental gradient explained the turnover in allele frequencies (Láruson et al. 2022).

Neutral evolutionary and demographic processes also impact GO predictions and how to account for them remains unclear. In particular, greater genetic drift in small populations results in greater allele turnover (e.g., in ecologically-marginal populations; Theraroz et al. 2023), and thus higher GO predictions without any selection pressure being involved (Láruson et al. 2020). Then, to avoid the confounding effects of the population genetic structure, it was initially recommended to predict GO based on a set of genetic markers previously identified as potentially involved in climate adaptation (Fitzpatrick and Keller 2015, Fitzpatrick et al. 2018), e.g., using gene-environment association (GEA) analyses correcting for population genetic structure. However, when patterns of demographic and adaptive history are confounded (e.g., species that migrated along climatic gradients after the last glaciation), correcting for population genetic structure is likely to hamper the detection of climate adaptation signals, and adopting a less strict approach to SNP selection might be more appropriate (Capblancq et al. 2023). Moreover, some studies have suggested that GO can be predicted equally well by random loci, thus questioning the relevance of prior selection of a set of candidate loci (Láruson et al. 2022, Lind et al. 2023). Finally, the uncertainty in GO predictions also stems from the inherent uncertainty in future climate forecasts, which comes from unknowns in future greenhouse gas emission scenarios (which depends on socioeconomic factors, technological advancements, and policy decisions), internal climate variability (which hinders the estimation of long-term climate trends), uncertainty in the magnitude and timing of external forcings (e.g., volcanic eruptions, changes in solar radiation), as well as uncertainty in the parameterization, process representation, simplifications and approximations of the general circulation models (GCMs) used to simulate the Earth’s climate system (Collins et al. 2013).

Despite these limitations, the GO concept has been repeatedly validated with empirical and simulation studies showing that fitness reductions in the novel environment of common garden experiments were well explained by GO predictions, and often better than by the commonly used climatic transfer distances (CTDs; e.g., Capblancq et al. 2018, Lind et al. 2018, Fitzpatrick et al. 2021, Láruson et al. 2022). Bay et al. (2018) also showed that higher GO predictions were associated with decreasing population sizes over the past half century in a North American migratory bird, though further investigation is necessary to determine whether this association results from higher genetic drift in decreasing populations or from population declines caused by climate maladaptation (Fitzpatrick et al. 2018, Láruson et al. 2022). The concept of GO therefore offers great promise as an informative metric for identifying populations at risk of maladaptation, but its predictions must be evaluated across a wider range of scenarios, species and validation approaches.

To test the method, we evaluated the consistency and empirical validity of GO predictions in maritime pine (*Pinus pinaster* Ait.), a tree species with fragmented populations in southwestern Europe and North Africa and a strong neutral population genetic structure (Alberto et al. 2013, Jaramillo-Correa et al. 2015). Using climatic and genomic data (9,817 SNPs) from 34 populations (454 trees clonally replicated in five common gardens), we studied the variability of GO predictions from four methods (namely GF, RDA, GDM and LFMM), two sets of SNPs (SNPs previously identified as candidate loci potentially involved in climate adaptation and neutral loci), and five climate general circulation models (GCMs). We then estimated the association between GO predictions and mortality rates in natural stands sampled in the French and Spanish National Forest Inventories (NFI), as well as height and mortality data from five clonal common gardens planted in contrasted environments (the CLONAPIN network; Archambeau et al. 2022, de-Miguel et al. 2022). Finally, we discussed the risk of maladaptation in maritime pine based on GO predictions from the two best-validated methods, and more generally the role and limits of GO predictions in conservation and management strategies, and how they can be used effectively.

## 2 Materials & Methods

### 2.1 Focal species

Maritime pine (*Pinus pinaster* Ait., Pinaceae) is a wind-pollinated, outcrossing and long-lived tree species with large economic and ecological importance in southwestern Europe and North Africa. The species is widely exploited for its wood, and has a key role stabilizing coastal and fossil dunes and, as a foundation species, supporting biodiversity (Viñas et al. 2016). Natural populations of maritime pine are scattered over a large range of climatic conditions: the dry climate along the northern coasts of the Mediterranean Basin (from southern Spain to western Italy), the mountainous climates of southeastern Spain and Morocco (Rif and Atlas mountains), the wetter climate of the Atlantic region (from northwestern Spain and Portugal to the southwestern part of France), and the continental climate of central Spain (Fig. 1). Maritime pine populations are also found on a wide range of substrates (from sandy and acidic soils to more calcareous ones) and in fire-prone regions, showing intraspecific variability in fire-related traits such as early flowering and serotiny (Tapias et al. 2004, Budde et al. 2014). Maritime pine is therefore a particularly interesting species for studying local adaptation and several studies have already provided evidence of genetic differentiation for adaptive traits (e.g., González-Martínez et al. 2002, de-Miguel et al. 2022).

**Figure 1:**
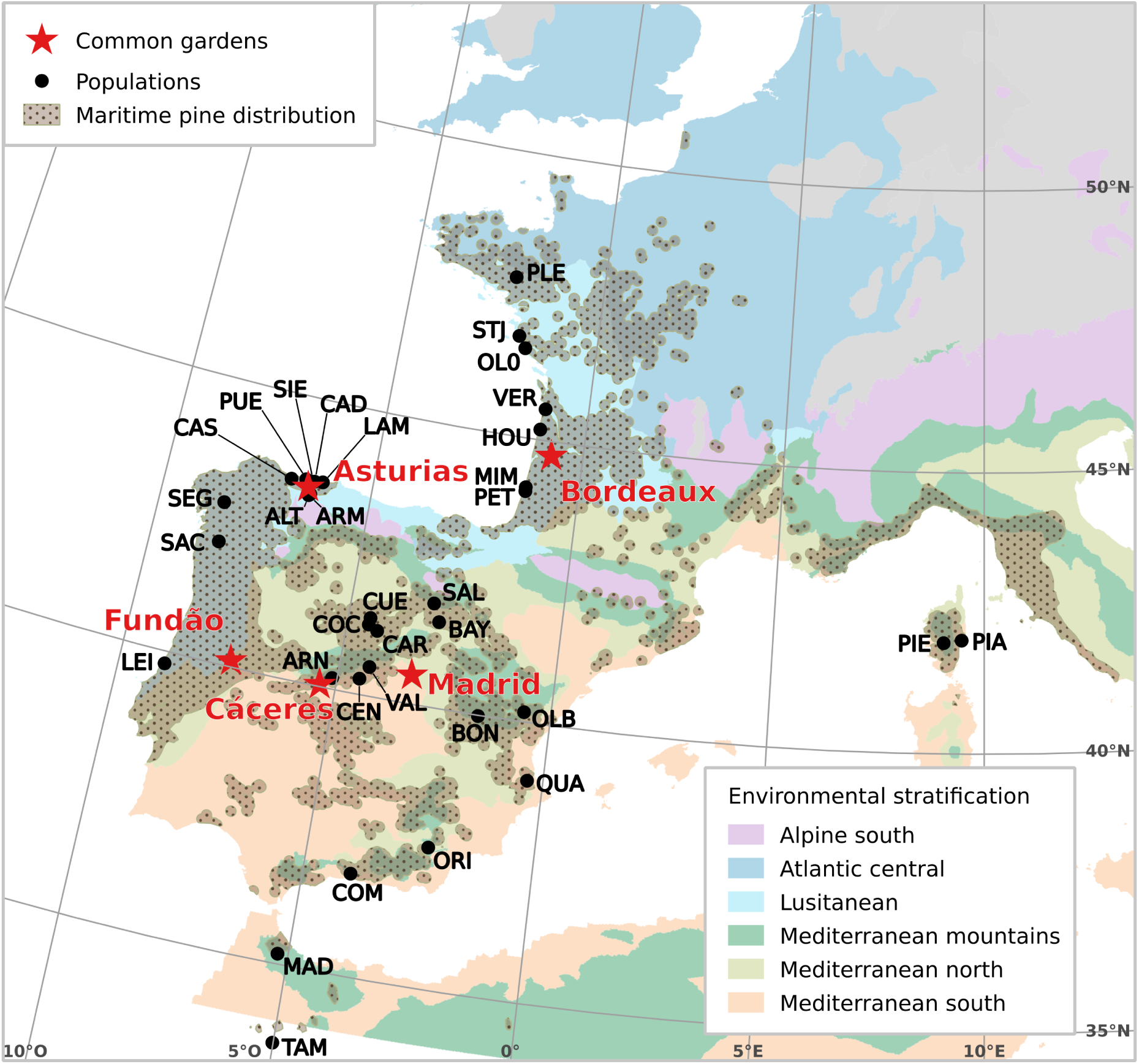
Location of the 34 genotyped maritime pine populations (filled dots) and the five common gardens (red stars) used in the validation steps and in which the same 34 populations were planted (CLONAPIN network). The environment stratification from Metzger (2018) shows the variety of environments inhabited. The maritime pine distribution depicted in the figure combines the widely-used EUFORGEN map (http://www.euforgen.org/) and 10-km radius areas around the French and Spanish National Forest Inventory (NFI) plots with maritime pine occurrence. NFI plots were surveyed between 2000 and 2014 for the French inventory and between 1986 and 2008 for the Spanish inventory.

Maritime pine has a strong population genetic structure (Alberto et al. 2013), with eight gene pools and two isolated one-population genetic clusters described in Theraroz et al. (2023). The sampled populations originated from six of the main gene pools previously identified in Jaramillo-Correa et al. (2015), each probably originating from an unique glacial refugia (Bucci et al. 2007, Santos-del-Blanco et al. 2012).

### 2.2 Clonal common garden network (CLONAPIN)

For GO estimation and validation in common gardens, we used genomic and phenotypic data from a network of five clonal common gardens (CLONAPIN; Fig. 1), in which the clones (i.e., genotypes) were obtained from open-pollinated seeds collected in natural populations and vegetatively propagated by cuttings (eight clonal replicates per genotype in each common garden; see details in de-Miguel et al. 2022). Two trial sites, Asturias (Spain) and Bordeaux (France), benefit from the favorable climates of the Atlantic European region (i.e., mild winters, high annual rainfall and relatively wet summers; Table S8) and have low proportions of dead trees: ∼ 2.8% in Asturias and ∼ 3.6% in Bordeaux, when trees were 37 and 85 months old, respectively (Tables S9 & S10). Two trial sites, Cáceres and Madrid (both in central Spain), experienced a strong drought during the summer following planting (Table S8), worsened by the presence of clay soils and leading to the death of 92% and 72% of the trees, respectively (Tables S9 & S10). The last trial site, Fundão (Portugal), is located on the border between the Atlantic and Mediterranean climates, and thus also experiences relatively dry summers (Table S8), leading to the mortality of ∼ 36.3% of the trees (recorded when the trees were 27 months old; Tables S9 & S10). One ramet of each clone in the Asturias common garden were genotyped with the Illumina Infinium assay described in Plomion et al. (2016) and with the new ThermoFisher 4TREE multispecies Axiom assay (Guilbaud et al. 2020). In our study, we used 454 genotypes from 34 populations (13 genotypes per population on average; Table S1), and with less than 18% missing data. We filtered out SNPs with minor allele frequencies below 1% or with more than 20% missing data, which resulted in 9,817 high-quality polymorphic SNPs, of which 2,855 were genotyped by both assays to ensure sample identity and estimate genotyping errors.

### 2.3 Climatic data

Climatic data at the location of the populations, common gardens and NFI plots were extracted from the Climate Downscaling Tool (Marchi et al. 2024). The climatic variables used to predict and validate GO were the mean annual temperature (*bio1*), the isothermality (*bio3*, ratio of the mean diurnal temperature range to the annual temperature range), the temperature seasonality (*bio4*), the annual precipitation (*bio12*), the precipitation seasonality (*bio15*) and the summer heat moisture (*SHM*) index (Table S3). They were selected based on their biological relevance for maritime pine, their contribution to the genetic variance using a RDA-based stepwise selection (Capblancq and Forester 2021) and the magnitude of their exposure to climate change (see details in section 3 of the Supplementary Information). For future climates, we used the climatic forecasts for the period 2041-2060 under the shared socio-economic pathway (SSP) 3.7-0 and from five GCMs, namely GFDL-ESM4, IPSL-CM6A-LR, MPI-ESM1-2-HR, MRI-ESM2-0 and UKESM1-0-LL.

### 2.4 Partitioning of the genetic variance

Following Capblancq and Forester (2021), we estimated the proportion of genetic variation that can be uniquely attributed to climate (the five climatic variables selected for GO predictions), neutral population structure (first three axes of a PCA based on genomic data) and geography (distance-based Moran’s eigenvector maps or geographical coordinates) using a combination of RDAs and partial RDAs (pRDAs) implemented in the *vegan* R package (Oksanen et al. 2022; see section 4 of the Supplementary Information).

### 2.5 Identification of SNPs potentially involved in climate adaptation

We identified SNPs covarying with climatic gradients using five GEA approaches: RDA, pRDA, LFMM, BayPass and GF. These approaches were chosen based on their different but complementary features (e.g., univariate vs multivariate, account or not for population genetic structure). Prior to GEA analyses, missing allelic values were imputed using the most common allele within the main gene pool of the genotype of concern (although we acknowledge that some genotypes had high admixture rates). GEAs were estimated based on the average climate over the period 1901-1950 to capture the climatic conditions under which the populations have may evolved.

The RDA-based GEA approach is a multivariate constrained ordination method that models the linear associations among the climatic variables and genomic variation (Capblancq et al. 2018, Capblancq and Forester 2021). In the pRDA approach, the first three axes of a principal component analysis (PCA) based on genomic data not filtered for minor allele frequencies (Figure S1) were added as conditioning variables to the RDA model to account for the population genetic structure. Both in the RDA and pRDA, we identified outlier SNPs by first estimating the Mahalanobis distances between each locus and the center of the RDA space along the significant axes (Capblancq et al. 2018), and then selecting SNPs with extreme Mahalanobis distances based on a False Discovery Rate (FDR) of 5%.

The LFMM-based GEA relies on the new version of the LFMM algorithm available in the *LEA* R package, which estimates locus-specific effect sizes and latent factors by using a least-square method and incorporating multiple climatic variables (Gain and François 2021). The population genetic structure is accounted for in this method by using ancestry coefficients. As with RDA, outlier SNPs were identified based on a FDR threshold of 5%.

The GEA implemented with the standard covariate model from the BayPass software is an univariate approach in which the neutral genetic structure is accounted for using a population covariance matrix based on the population allele frequencies (Gautier 2015). SNPs were considered outliers if the median Bayes Factor (calculated over five independent runs) of their association with at least one climatic variable was higher than 10 deciban units, which corresponds to, at minimum, strong evidence on the Jeffrey’s scale (Jeffreys 1961).

The GF-based GEA consists in fitting GF models individually to each SNP and calculating empirical 𝑝-values based on their R^2^ rank (Lotterhos and Whitlock 2014), as described in Fitzpatrick et al. (2021) and Capblancq et al. (2023). We ran three independent runs and selected the 0.5% SNPs with the lowest empirical 𝑝-values for each run (i.e., 49 SNPs). We then considered as outliers only those SNPs identified by the three runs.

To obtain the set of candidate SNPs used for GO predictions, we selected outlier SNPs identified by at least two GEA methods. When some outlier SNPs were located on the same scaffold/contig (a scaffold/contig measuring approximately between 400 and 10,000 bp), we kept the SNP with the lower *p*-value in the RDA. Among the loci that were not identified as outliers by any of the GEA methods, we randomly sampled the same number of SNPs as in the candidate SNP set to obtain a set of control SNPs (i.e., loci expected not to be involved in climate adaptation).

### 2.6 Genomic offset (GO) predictions

We evaluated GO predictions from four methods: GF, RDA, LFMM and GDM. These methods rely on common steps: (i) estimating gene-climate relationships under a reference period (1901-1950 in our study), (ii) projecting the genomic composition across the landscape under past and future climates, and (iii) calculating the difference between the predicted current and future genomic composition for each pixel of the landscape, i.e., the genetic change required to maintain the estimated gene-climate relationships under future climates (Capblancq et al. 2020). RDA and LFMM approaches are based on the extrapolation of linear gene-climate relationships (Capblancq and Forester 2021, Gain and François 2021), while GF and GDM are nonparametric approaches that model nonlinear allele turnover functions along climatic gradients (Fitzpatrick and Keller 2015). For each method, we predicted GO at the location of the populations under future climates for the sets of candidate and control SNPs, as well as all SNPs for LFMM, as this method is designed to work without pre-selecting candidate SNPs. We also assessed the relationship between GO predictions and Euclidean climate distances (i.e., the differences in climate between the reference period and the future period for each population) to identify the GO predictions that deviate the most from this widely-used climate-only index.

### 2.7 Validation of genomic offset (GO) predictions

We used two approaches to validate GO predictions involving either natural populations or common gardens in contrasted environments.

First, we evaluated which methods provided the strongest associations between GO predictions and mortality rates in natural populations. We used mortality data from the French and Spanish NFIs harmonised in Changenet et al. (2021) and covering the period 2000-2014 for the French inventory and 1986-2008 for the Spanish inventory. Note that we did not attempt to validate GO predictions with tree height data from NFIs because the French NFI does not provide tree age. Tree diameter at breast height (1.30 m, DBH) ranged from 10 to 263 cm, hence including both saplings and adult trees. The 34 sampled populations cover most climatic space of the NFI plots, except for locations with high *SHM* (Figure 4a), or low *bio15*, *bio3* or *bio1* (Figures S59 and S60). GO predictions at the locations of the NFI plots capture the mismatch between the projected current genomic composition of the populations and the genomic composition predicted to be optimal under the climatic conditions of the inventory period (e.g., Figure 4b for GO predictions with GF and the candidate SNPs). We modeled the probability of mortality 𝑝_𝑖_ of maritime pine in the plot 𝑖 during the census interval Δ_𝑖_ with a complementary log-log link as follows:

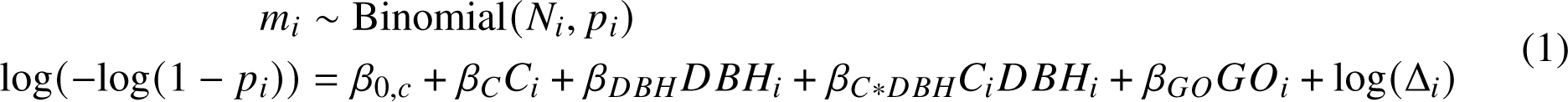

𝑁_𝑖_ is the total number of maritime pines in the plot 𝑖. 𝑚_𝑖_ is the number of maritime pines that died during the census interval Δ_𝑖_ in the plot 𝑖. 𝐶_𝑖_ is the basal area of all tree species confounded in the plot 𝑖 (to account for the competition among trees), with 𝛽_𝐶_ being its associated regression coefficient. 𝐷𝐵𝐻_𝑖_ is the mean DBH of all maritime pines (including adults, dead trees and saplings) in the plot 𝑖 (to account for age-related mortality), with 𝛽_𝐷𝐵𝐻_ being its associated regression coefficient and 𝛽_𝐶∗𝐷𝐵𝐻_ being the regression coefficient capturing the interaction between 𝐶_𝑖_ and 𝐷𝐵𝐻_𝑖_. 𝐺𝑂_𝑖_is the GO predicted in the plot 𝑖, with 𝛽_𝐺𝑂_ being its associated regression coefficient. 𝛽_0,𝑐_ are country-specific intercepts that account for the methodological differences between the French and Spanish NFIs. N (0, 1) prior distributions were used for parameter estimation.

Second, we identified which methods provided the strongest associations between GO predictions and mortality and height data in the five common gardens from the CLONAPIN network (Figure 1). GO predictions at the locations of the five common gardens capture the mismatch between the current genomic composition of the populations and the genomic composition predicted to be optimal under the climates of the common gardens between the planting and the measurement dates. We also calculated the climatic transfer distances, CTDs (i.e., absolute climatic difference between the location of the populations and the common gardens), for the six climatic variables used and compared them with GO predictions. We modeled the probability of mortality 𝑝 _𝑝_ in the population 𝑝 independently in the five common gardens as follows:

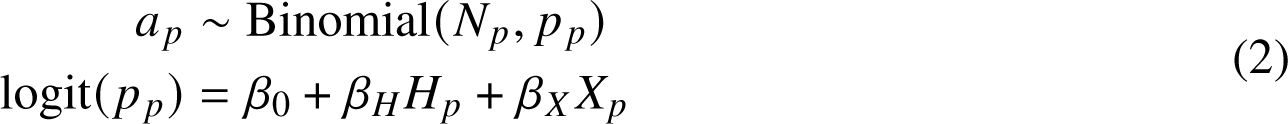

𝑎 _𝑝_ is the count of individuals that died in the population 𝑝. 𝑁_𝑝_ is the total number of individuals from the population 𝑝 that were initially planted in the common garden. 𝑋_𝑝_is the GO or CTD of the population 𝑝, with 𝛽_𝑋_ being its associated coefficient. 𝐻_𝑝_ is a proxy of the initial tree height at the planting date (i.e., population intercept estimated across all common gardens in model 1 of Archambeau et al. 2022), with 𝛽_𝐻_being its associated regression coefficient. Initial height at planting was included in the models to account for the observed higher survival of taller trees in the common gardens. N (0, 5) prior distributions were used for 𝛽_0_, 𝛽_𝐻_ and 𝛽_𝑋_ .

We modeled tree height independently in the five common gardens as follows:

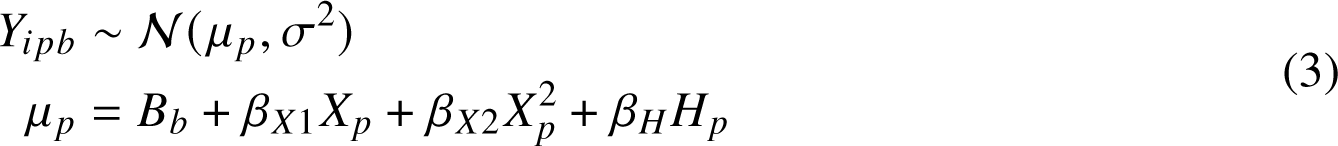

𝑌_𝑖𝑝𝑏_is the height of the individual 𝑖 in the population 𝑝 and the block 𝑏. 𝐵_𝑏_are the block intercepts. 𝜎^2^ is the residual variance. 𝛽_𝑋1_ and 𝛽_𝑋2_ are the linear and quadratic regression coefficients of 𝑋_𝑝_, respectively, and 𝛽_𝐻_ is the linear regression coefficient of 𝐻_𝑝_ (see equation 2). The quadratic form was used to allow the slope of the association between height and GO to vary (such as in Fitzpatrick et al. 2021). N (0, 1) prior distributions were used for 𝐵_𝑏_, 𝛽_𝑋1_, 𝛽_𝑋2_ and 𝛽_𝐻_ and Exp(1) for 𝜎.

The statistical models were implemented in a Bayesian framework using the Stan probabilistic programming language (Carpenter et al. 2017), based on the no-U-turn sampler algorithm. All analyses were undertaken using R Statistical Software v4.2.2 (R Core Team 2022) and scripts are available at https://github.com/JulietteArchambeau/GOPredEvalPinpin.

## 3 Results

Based on RDA/pRDA approaches, geography, climate and the population neutral genetic structure explained 80.3% of the variation in population-allele frequencies (Figure S5 and Table S6). Single variables explained 29.8% of the variance (7.9% by climate, 8.3% by geography and 13.6% by the neutral genetic structure) while the rest (50.5%) could not be uniquely attributed to a given variable (i.e., confounded effects).

The GEA analyses identified 240 outlier SNPs with the RDA, 178 with the pRDA, 46 with BayPass, 40 with GF and 120 with LFMM (Figure 2). 85 SNPs were identified by at least two GEA methods and after retaining a single outlier SNP from those located on the same scaffold/contig, we ended up with a final set of 69 candidate SNPs.

**Figure 2:**
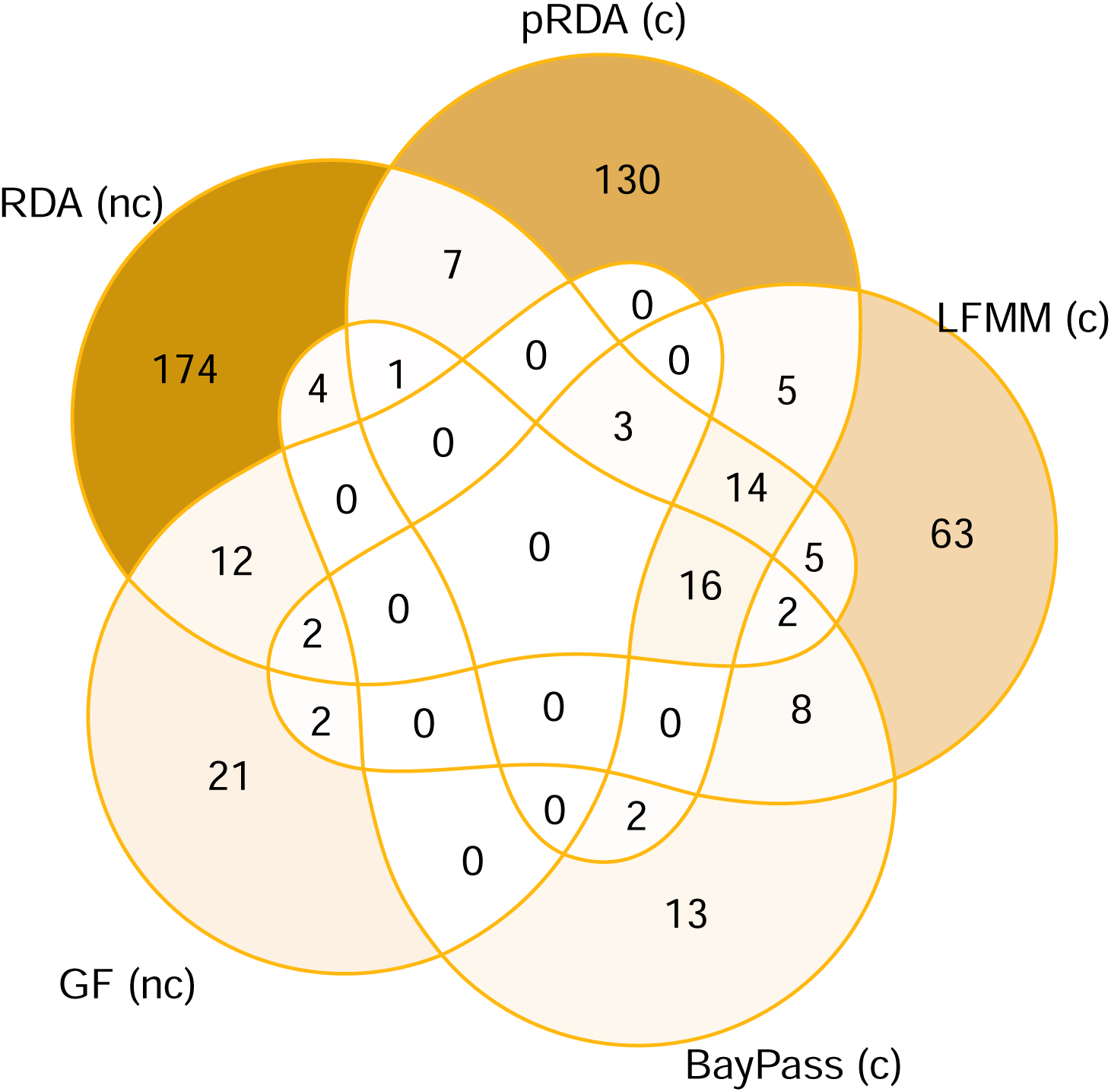
Venn diagram showing the number of outlier SNPs in common among the five gene-environment association (GEA) approaches, namely Redundancy analysis (RDA), partial RDA (pRDA), Gradient Forest (GF), BayPass and latent factor mixed models (LFMM). Approaches annotated with ’(c)’ incorporate corrections for population structure, while approaches annotated with ’(nc)’ do not.

### 3.1 Variability in genomic offset (GO) predictions

We observed substantial variation among GO predictions from the different methods, SNP sets and GCMs (Figure 3). GO predictions from GDM and GF and the candidate SNPs were negatively or poorly correlated with the other GO predictions (Figure 3a) and the Euclidean climatic distances (Figures S33 and S38). By contrast, GO predictions from GDM and GF and the control SNPs showed positive correlations with predictions from LFMM and RDA (Figure 3a) and with the Euclidean climatic distances (Figures S32 and S37). GO predictions from LFMM and RDA were highly correlated (Figure 3a) and were also correlated with the Euclidean climatic distances (Figures S30-S31 and S34-36). As a result, populations with the highest predicted GO (i.e., populations that are predicted to be at risk of maladaptation) were not the same depending on which method and SNP set was used (Figure 3b). As an example, the Iberian Atlantic populations of Armayán

**Figure 3:**
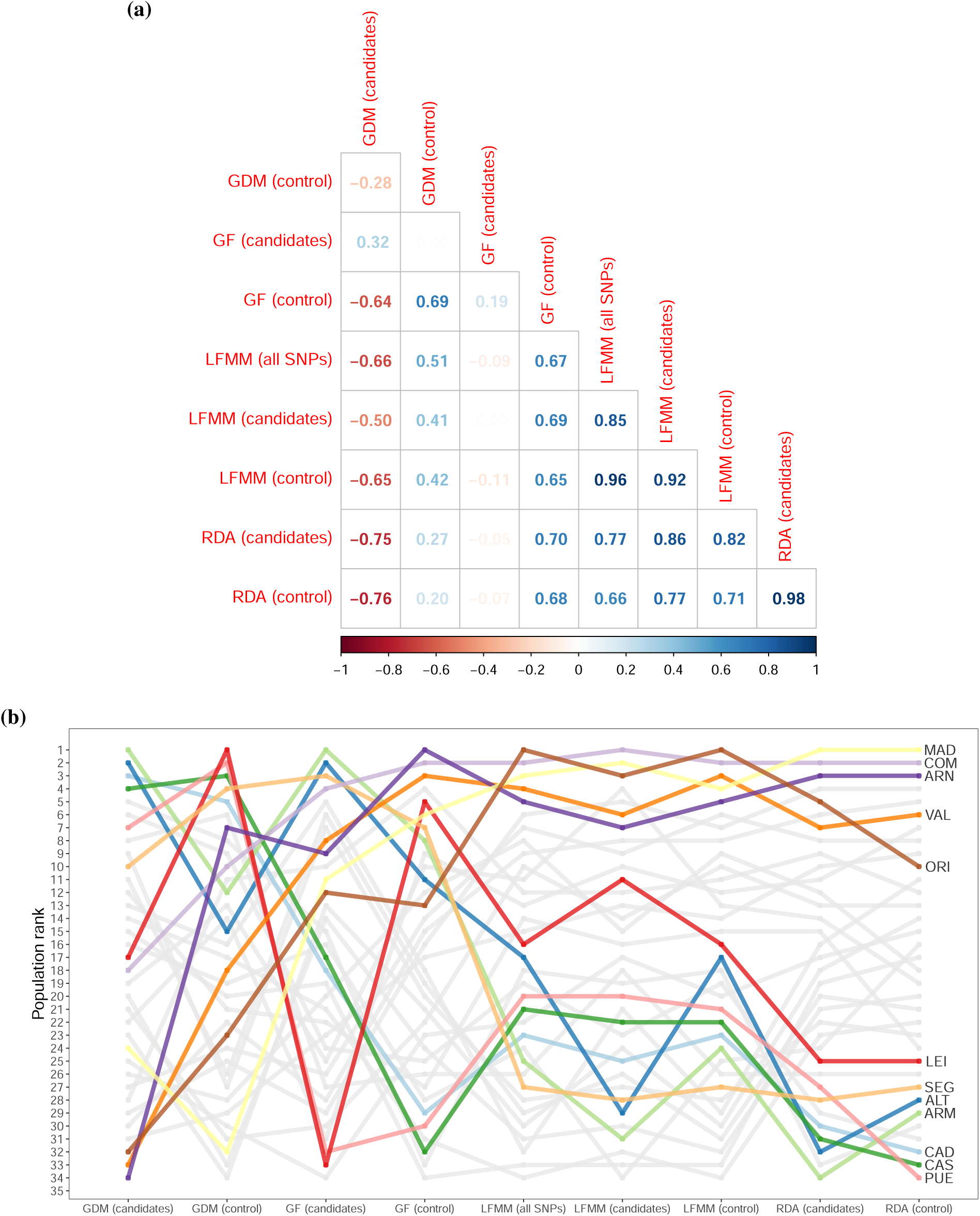
Variability in GO predictions across methods and SNP sets. GO predictions correspond to the averaged prediction across the five GCMs. Similar figures for GO predictions for each GCM can be found in section 7.6 of the Supplementary Information. (a) Pairwise correlations between GO predictions for each combination of method (GF, RDA, GDM and LFMM) and SNP set (candidate and control SNPs, and also all SNPs for LFMM only). (b) Population ranks for GO predictions for each combination of method and SNP set. Populations with higher GO have a lower value rank. Colored populations are those having GO values in the top three highest values in at least one column (i.e., combination of SNP set and method).

(ARM), Alto de la Llama (ALT) and Cadavedo (CAD) had the highest predicted GO based on GDM and candidate SNPs but were among the populations with the lowest predicted GO based on RDA or LFMM and candidate SNPs (Figure 3b). Conversely, Oria (ORI) population from southeastern Spain, and Valdemaqueda (VAL) and Arenas de San Pedro (ARN) populations from central Spain, had the lowest GO predictions based on GDM and candidate SNPs but were among the populations with the highest GO based on RDA or LFMM and candidate SNPs (Figure 3b). Moreover, for a given combination of method and SNP set, populations with the highest predicted GO were not the same across GCMs, though some combinations (e.g., GF or LFMM and candidate SNPs) showed more consistency than others (see section 7.6.2.2 of the Supplementary Information).

### 3.2 Validation of genomic offset (GO) predictions

GO predictions based on candidate SNPs were positively associated with mortality rates in the NFI plots for GDM, GF and RDA (95% credible intervals not overlapping zero; Figure 4c), and these associations were highly unlikely to be obtained by chance (Figure S73). More specifically, a one-standard deviation increase in GO predictions resulted in an estimated mean increase in tree mortality probability of 23.1% in Spain and 24.5% in France with GDM and the candidate SNPs, and 19.3% in Spain and 19.2% in France with GF and the candidate SNPs (Table S7). Figure 4 illustrates how higher GO predictions based on GF and the candidate SNPs (e.g. in Galicia, south-eastern Spain or French Brittany; Figure 4b) translate in higher predictions of annual probability of tree mortality in the NFI plots (when the other covariates of the validation models are fixed to the mean and the country fixed to Spain; Figure 4d). By contrast, none of GO predictions with LFMM were positively associated with mortality rates, and GO predictions based on control SNPs were positively associated with mortality rates only for GF (Figure 4c).

**Figure 4:**
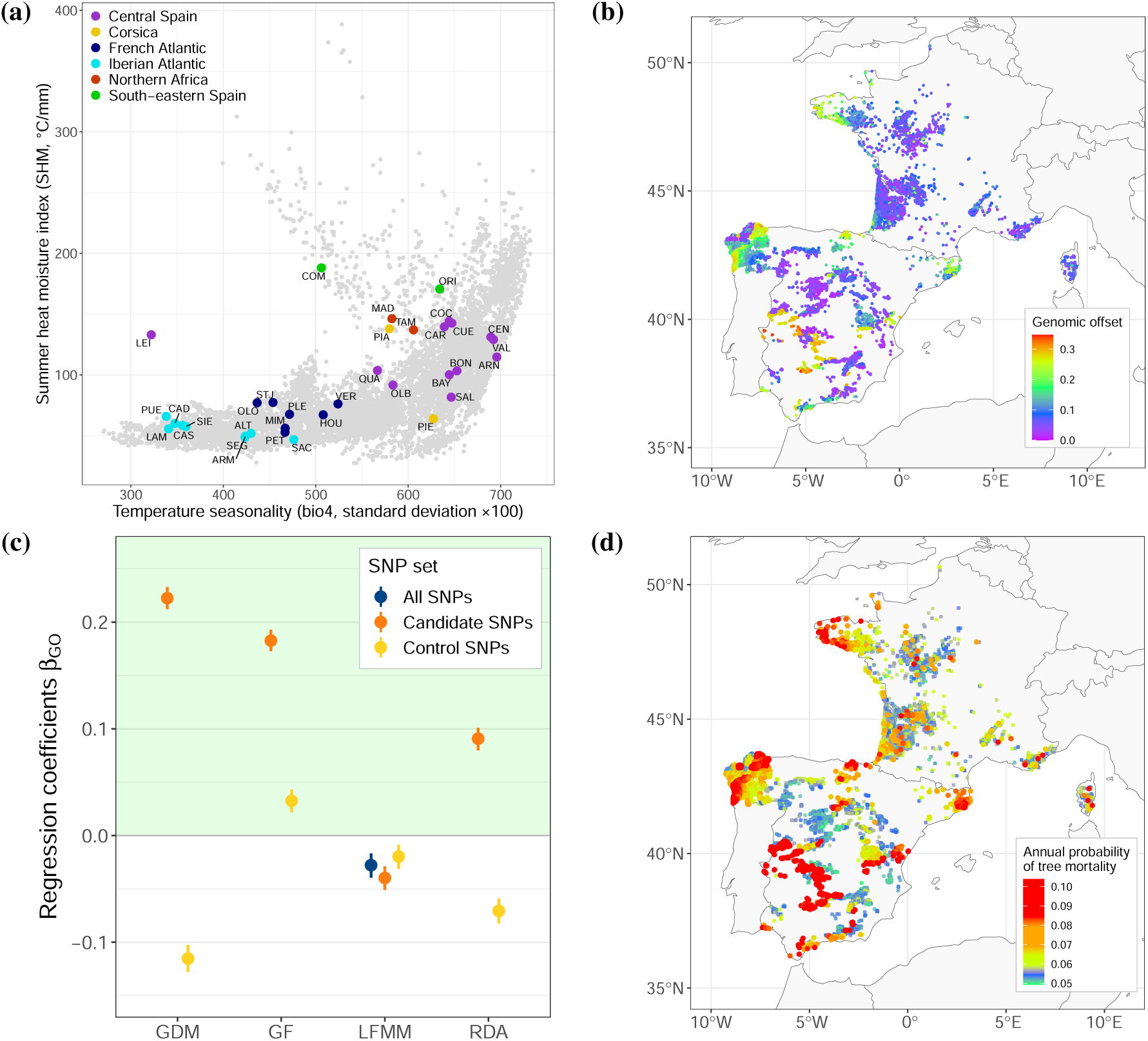
Validation of GO predictions using mortality data from the French and Spanish National Forest Inventory (NFI) plots. (a) Summer heat moisture (SHM) index and temperature seasonality (the two main predictors of the GF models) at the location of the NFI plots (in gray) and the 34 sampled populations (coloured according to their main gene pool). Climatic values correspond to average climates over the reference period 1901-1950. (b) Estimated mean and 95% credible intervals of the regression coefficients 𝛽_𝐺𝑂_ standing for the association between mortality rates and GO predictions in the NFI plots. Coefficients in the green area have the expected sign, indicating that plots with higher GO predictions also have higher mortality rates. See Figures S69, S70 and S71 for the estimated regression coefficients 𝛽_𝐶_, 𝛽_𝐷𝐵𝐻_ and 𝛽_𝐶∗𝐷𝐵𝐻_. (c) GO predictions with GF and the candidate SNPs at the locations of the NFI plots. These predictions capture the mismatch between the projected current genomic composition of the populations and the genomic composition predicted to be optimal under the climatic conditions of the NFI period. (d) Predicted annual probability of mortality in each NFI plot with covariates 𝐶 (basal area of all tree species) and 𝐷𝐵𝐻 (mean DBH of all maritime pines) fixed to the mean and country fixed for Spain.

In the three common gardens with dry and hot summers (Cáceres, Fundão and Madrid), mortality rates were positively associated with GO predictions for all methods, with 95% credible intervals not overlapping zero for most methods (Figure 5). With respect to the two common gardens under milder Atlantic climate, mortality rates in Bordeaux were also positively associated with GO predictions for all methods except RDA (with the 95% credible intervals of LFMM and control and all SNPs overlapping zero), whereas in Asturias, no association between GO predictions and mortality rates was found (Figure 5). CTDs based on temperature seasonality (*bio4*) were highly correlated with GO predictions from candidate SNPs (Figure S95) and were therefore also good predictors of mortality rates in all common gardens except Asturias (Figure S100). This similarity stems from the identification of temperature seasonality as the main climatic driver explaining genomic variation of candidate SNPs in all four GO methods (e.g., Figure S15 for GF and Figure S27 for GDM), and therefore the major weight of this variable when generating GO predictions. The other CTDs showed less consistent associations with mortality rates across common gardens (Figure S100).

**Figure 5:**
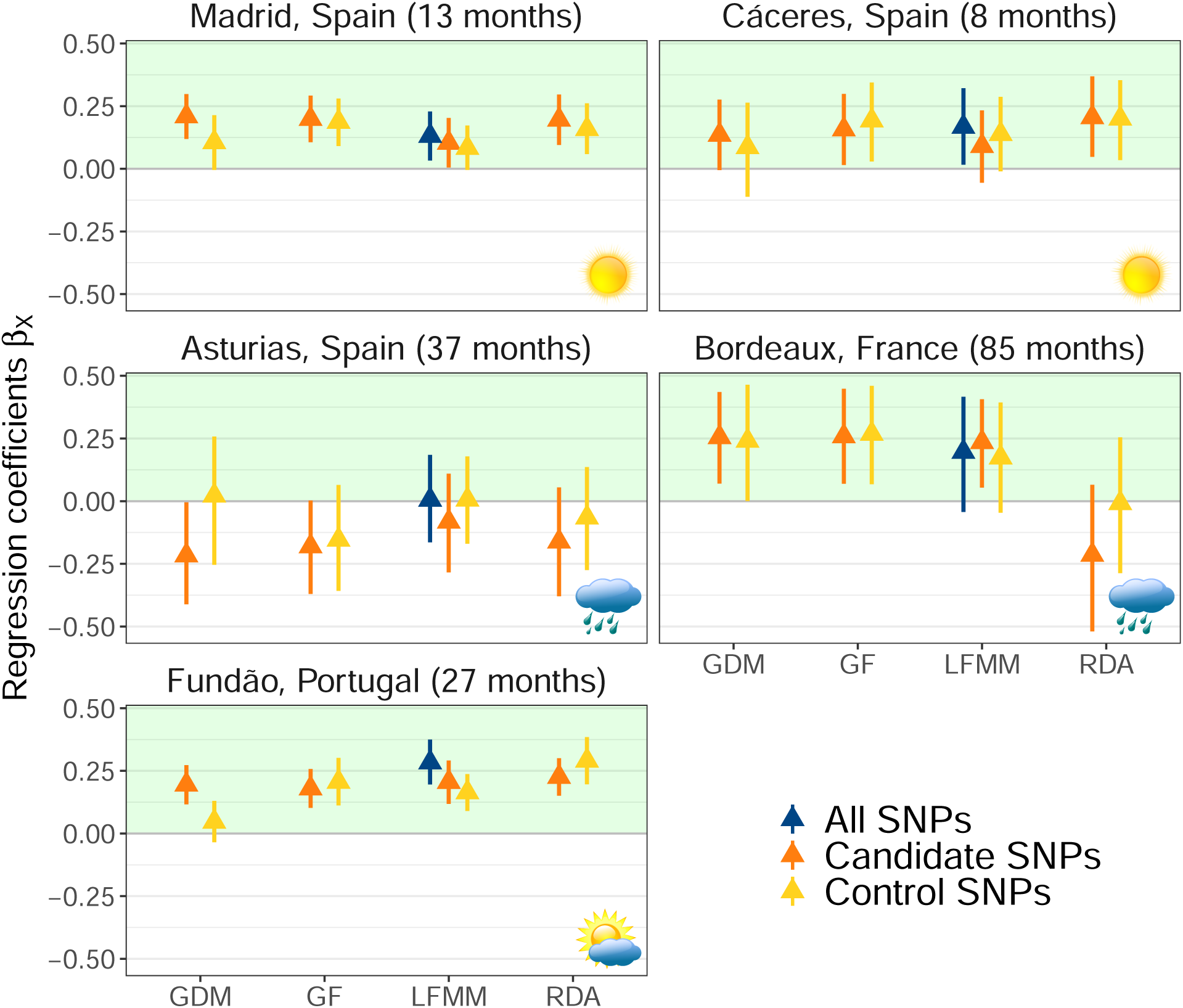
Estimated mean and 95% credible intervals of the regression coefficients 𝛽_𝑋_ standing for the association between mortality rates and GO predictions in the five common gardens. Graph titles include the time in months corresponding to the age at which height and survival were recorded. Coefficients in the green area have the expected sign, reflecting higher mortality rates in populations with higher GO predictions. The weather icons represent the climatic conditions in the five common gardens between the planting and measurement dates: Atlantic climates with mild winters and relatively wet summers in Asturias and Bordeaux, Mediterranean climates with severe summer droughts in Cáceres and Madrid, and intermediate climates with wet winters but dry summers in Fundão.

Tree height in the five common gardens was strongly associated with the proxy for initial tree height at the planting date (Figure S112). GO predictions (as well as CTDs) were not associated with height in Asturias and Fundão (Figures S113-S115, S119-120 & S124). In Cáceres and Madrid, some of the GO predictions (e.g., from GF and control SNPs or from RDA and candidate SNPs) had negative linear regression coefficients (Figure S113) but their overall estimated relationship with height (after accounting for the quadratic coefficients, Figure S114) had unexpected inverted U-shaped forms (Figures S116 & S118). Overall, GO predictions from RDA (with both control and candidate SNPs) in Bordeaux were the only GO predictions with a linear negative relationship with tree height (Figures S113 & S117). However, the strength of these associations remained small and the proportion of variance explained by the height models including these GO predictions could almost not be differentiated from the proportion of variance explained by a model not including them (Figure S125). CTDs also showed generally weak and inconsistent relationships with tree height across the five common gardens; but notice the consistent and relatively stronger associations of the CTDs for precipitation seasonality and annual precipitation in the dry sites (Cáceres, Fundão and Madrid), and to a lesser extent in Bordeaux (Figures S113-114 and S120-S124).

## 4 Discussion

The GO concept has gained considerable traction in recent years, and is often suggested as a tool for predicting population performance under climate change *in situ* or at a potential location for transplantation (e.g., Rhoné et al. 2020, Lachmuth et al. 2023). However, GO predictions have only been validated under a limited number of scenarios (mostly with height measurements from common gardens; e.g., Fitzpatrick et al. 2021) and species, e.g., balsam poplar (*Populus balsamifera*) in Fitzpatrick et al. (2021) and lodgepole pine (*Pinus contorta*) in Capblancq et al. (2018). We evaluated the consistency and empirical validity of GO predictions from four methods (GF, GDM, LFMM and RDA), two sets of loci (control and candidate SNPs) and five GCMs using data from 34 populations (454 genotypes) of maritime pine, a forest tree species with strong population genetic structure. We found substantial variability in GO predictions and demonstrated the importance of combining different sources of data to assess their empirical validity. GO predictions from the two nonparametric approaches, GF and GDM, and the candidate SNPs showed the strongest and most consistent associations with mortality rates in common gardens and natural populations. GO predictions from candidate and control SNPs were similarly associated with mortality rates in common gardens, while only GO predictions from candidate SNPs were able to predict mortality rates in natural populations. We observed almost no linear negative associations between GO predictions and tree height in the common gardens, which most likely originates from the strong effect of the population genetic structure on tree height in maritime pine (Archambeau et al. 2022). Our study demonstrates the imperative to validate GO predictions using multiple data sources, before these predictions can be used as informative metrics in conservation or management strategies.

### 4.1 High variability in genomic offset (GO) predictions across methods

Variability in GO predictions across methods was somehow expected given their different theoretical assumptions (Gain et al. 2023). RDA and LFMM assume linear adaptive gradients and Gaussian selection within populations (Capblancq and Forester 2021, Gain and François 2021), and under some conditions, Gain et al. (2023) demonstrated that their predictions should be identical. GF and GDM are nonparametric approaches that make no a priori assumption about the shape of the gene-climate relationships and the selection gradient within populations (Fitzpatrick and Keller 2015, Gain et al. 2023). These methodological differences may explain why we observed a clear distinction among GO predictions based on GDM or GF and candidate SNPs and the other approaches (Figure 3). As GDM and GF using candidate SNPs showed better empirical validity, it suggests that the turnover in allele frequency of the candidate SNPs along the climatic gradients is nonlinear, which may not be the case for the control SNPs. The nonlinearity of adaptive gradients may be common in forest trees, e.g., it has been found in red spruce (*Picea rubens*, Mahony et al. 2020, Capblancq et al. 2023). Interestingly, simulations have shown that GF turnover functions better capture the turnover patterns for non-monotonic adaptive gradients than for linear ones (Láruson et al. 2022). Evaluating the shape of adaptive gradients beforehand might therefore prove relevant in making the GO approach more robust and trustworthy.

Although both GDM and GF methods using candidate SNPs identified the populations of Armayán (ARM) and Alto de la Llama (ALT) as being the most at risk of maladaptation, there was still substantial variability between the two approaches (Figure 3). One of the most striking differences was that the three populations predicted to have the lowest risk of maladaptation in the GDM approach, i.e., Oria (ORI), Valdemaqueda (VAL) and Arenas de San Pedro (ARN), were among the twelve populations predicted to be at highest risk with the GF approach (Figure 3b). This prediction variability may originate from using different algorithms to estimate the turnover functions, integrating the geographical distance among populations in the GDM approach or accounting for correlation between predictors in the GF approach (Fitzpatrick and Keller 2015).

Demonstrating the high variability in GO predictions across methods is not new. Fitzpatrick and Keller (2015) pointed out the differences between GF and GDM predictions in the paper that launched the GO concept and advised the use of the two approaches in tandem. Lind et al. (2023) found high variability among predictions from GF and RDA in Douglas fir (*Pseudotsuga menziesii*), but not in jack pine (*Pinus banksiana*). By showing the high inconsistency across GO predictions from four methods, our results therefore provide further evidence that relying on the predictions of a single method without any knowledge of its empirical validity for the species under study must be avoided.

### 4.2 The role of population genetic structure

To date, no consensus has been reached on how to account for population genetic structure and demographic history in landscape genomics. This also applies to GO approaches in which the first step is to identify a set of loci potentially involved in climate adaptation (Capblancq et al. 2023, Eckert and Neale 2023). In species for which climatic gradients that drive local adaptation covary with neutral genetic variation, correcting for population structure in GEA reduces the number of false positives but at the expense of a higher number of false negatives (Lotterhos and Whitlock 2015). In red spruce, a species in which climate adaptation strongly covaries with neutral population structure and geographic distance, Capblancq et al. (2023) showed that there was almost no overlap among outliers identified by GEA methods correcting for population genetic structure and those that did not, and that selecting outliers only from those methods that did correct resulted in the removal of most of the adaptive signal. Therefore, to obtain a set of candidate SNPs representative of climate adaptation, Capblancq et al. (2023) originally used a combination of four GEA methods, two with correction for population structure and two without, and considered as candidates for local adaptation loci that were identified by at least two of them. Given the highly confounded patterns of population genetic structure and climate adaptation in our sampled populations (Figure S5), and more generally in maritime pine (Alberto et al. 2013, Jaramillo-Correa et al. 2015, Archambeau et al. 2022), we followed the approach of Capblancq et al. (2023) and identified a set of 69 candidate SNPs, among which 12 were identified only by GEA methods not correcting for population genetic structure (Figure 2). The unambiguous higher associations of mortality rates in natural populations with GO predictions based on candidate SNPs compared to GO predictions based on control SNPs (Figure 4c) suggest that our set of candidate SNPs successfully included loci located within or nearby genes involved in climate adaptation, and thus captured part of the adaptive signal. It also suggests that climate maladaptation in populations with a high population structure would be better predicted using GO predictions based on pre-selected candidate SNPs.

The four methods evaluated in our case study also account in different ways for the neutral population genetic structure. We used GF without correction for population genetic structure as it has been successfully validated with simulations (Láruson et al. 2022) and common garden data (Lind et al. 2023). Correcting GF turnover functions for population genetic structure has been explored with Moran’s eigenvector maps (Fitzpatrick and Keller 2015) or allele frequencies corrected for population relatedness based on the population covariance matrix (Fitzpatrick et al. 2021). However, the first approach still lacks empirical validation, while the second has only been validated in core populations of balsam poplar with low genetic structure. GDM was performed with correction for the population genetic structure to estimate the gene-climate relationships (using population geographic coordinates) and no correction in the prediction model, whereas we applied the RDA without correcting for population structure at either stage. LFMM is the only method that incorporates the population genetic structure both in the estimation of the gene-climate relationships and in the prediction model (Rellstab et al. 2021), via latent factors included as covariates and representing the population genetic relatedness (Gain and François 2021). This may explain the more consistent associations with height and mortality data across SNP sets obtained with LFMM compared to the other GO methods. However, LFMM predictions showed weaker associations with mortality rates in the NFI plots compared to methods using candidate SNPs (Figure 4c) and were also less robust than GDM and GF predictions using candidate SNPs in the empirical validation based on mortality data from common gardens (Figure 5). These results suggest that correcting for population genetic structure in the prediction model of GO methods may hinder the ability to capture climate maladaptation in maritime pine. Whether this observation could be generalized to other forest tree species with strong population genetic structure will require further investigation.

Most importantly, our study has revealed the major effect that the population genetic structure can have on the validation of GO predictions in common gardens. Previous height growth models fitted on the five CLONAPIN common gardens showed that the genetic-by-environment interaction (G*E) explained very little variance, and that the genetic component of height growth was mainly explained by the gene pools of origin (Archambeau et al. 2022). In our study, we used the population intercepts from these models, which represent the average tree height of each population across the five common gardens, to account for the initial tree height at the planting date in the mortality models and the neutral genetic component in the height models. This approach worked well in the mortality models as most detected associations between GO predictions and mortality rates were in the expected direction, which was not the case without correcting for the initial tree height (results not shown). However, this approach did not allow us to observe negative associations between GO prediction and height consistently across the different common gardens, which may suggest that the effect of population genetic structure on height variation is too strong in maritime pine and prevents any use of height as a fitness proxy in studies comparing populations of this species.

### 4.3 Validation of genomic offset (GO) predictions using different data sources

Our results first demonstrate the value of combining different traits (mortality and height in our study) to validate GO predictions. GO predictions have mainly been validated with height data from common gardens until now (e.g., Capblancq et al. 2018, Fitzpatrick et al. 2021). However, for species with strong population genetic structure and low G*E, our study suggests that using tree height to validate GO predictions may not be appropriate, and that mortality data are probably more reliable. Greater reliability of mortality relative to height (and DBH) for validating GO predictions has also been found in jack pine (Lind et al. 2023). When both mortality and height data are available, an alternative option would be to estimate the association between height and GO predictions, setting the height of dead individuals to zero, as done in Capblancq et al. (2023).

Second, our study also suggests that robust validation of GO predictions requires the use of different data sources, ideally including a mix of observational data (if possible on a large scale) and experimental data (if possible in contrasted environments). While mortality rates in common gardens were almost equally well predicted by candidate or control SNPs (Lind et al. 2023 and our study), the use of mortality data from natural populations (NFIs) allowed us to demonstrate the greater empirical validity of candidate SNPs over control SNPs. To our knowledge, differences in the empirical validity of GO predictions from candidate or control SNPs have not been demonstrated so far, although predicting GO from a set of candidate SNPs is generally advised (Fitzpatrick and Keller 2015, Fitzpatrick et al. 2018, Capblancq et al. 2020).

The use of mortality rates in natural populations as a means of validating GO predictions also provides insights into the evolutionary dynamics that may be at play in a natural setting. On the one hand, higher mortality rates have the potential to accelerate adaptive evolution by enhancing turnover within populations (Kuparinen et al. 2010). On the other hand, higher predicted GO in populations with higher mortality rates suggests that these populations would have to undergo more significant changes to remain adapted to the changing climatic conditions, therefore counterbalancing the positive effects of mortality rates on adaptive evolution. Care should however be taken when interpreting this validation approach. First, although we corrected for some potential confounding factors (competition among trees, different sampling schemes across countries and differences in mean tree age across plots), we cannot assert that the remaining mortality events were caused by climate change alone. Indeed, mortality is a multifactorial stochastic process which is hard to model and whose causes are difficult to determine (Franklin et al. 1987, Lines et al. 2010). Second, one of the steps to predict GO at the locations of the NFI plots is to project the genomic composition across the landscape based on the estimated gene-climate relationships. Although our population sample represents the climatic space of the NFI plots relatively well (Figures 4a, S59 and S60), this step remains a spatial extrapolation, based on the assumption that the gene-climate relationships estimated in the studied populations hold true across the entire area covered by the NFI plots (Rellstab et al. 2021, Lind et al. 2023).

Finally, a better understanding of the conditions under which GO predictions are valid would greatly benefit from the use of other data sources, such as experiments specifically designed to assess the ability of populations to grow outside their range (e.g., the REINFFORCE arboretum network; https://reinfforce.iefc.net/en/), or regeneration and reproduction data in natural stands, and from the comparison of GO predictions with predictions from other types of models such as ecophysiological (e.g., Ruffault et al. 2022) or eco-evolutionary models (e.g., Cotto et al. 2017).

### 4.4 Risk of maladaptation in maritime pine

Maritime pine populations located in northwestern Iberian Peninsula and northern Brittany (France) were predicted to be at higher risk of maladaptation based on GF and GDM and the can-didate SNPs, i.e., the two methods that best predicted mortality rates in the validation steps (Figure 6). The risk of climate maladaptation for populations in central and eastern Spain remained much more uncertain, as the predictions of the two methods contradict each other: higher risk with GDM predictions vs. lower risk with GF predictions (Figure 6). Both methods identified the populations of Armayán (ARM) and Alto de la Llama (ALT) in northern Spain as the two populations with highest risk of climate maladaptation (Figure 3b). Compared to the other populations sampled, ALT and ARM do not currently experience particularly high exposure to climate change (Figure S126) nor show divergent genomic composition for control and candidate SNP sets (Figure S128). This illustrates how genomic and climatic information are combined with the GO approach to produce a novel metric of population maladaptation that differs from previous metrics based solely on climatic information (e.g., CTDs). However, we may note that GO predictions from candidate SNPs were strongly driven by temperature seasonality (*bio4*) in our study, thus resulting in high correlations between GO predictions from candidate SNPs and the CTDs based on temperature seasonality in common gardens (Figure S95), and ultimately similar associations with mortality rates (Figure S100).

**Figure 6:**
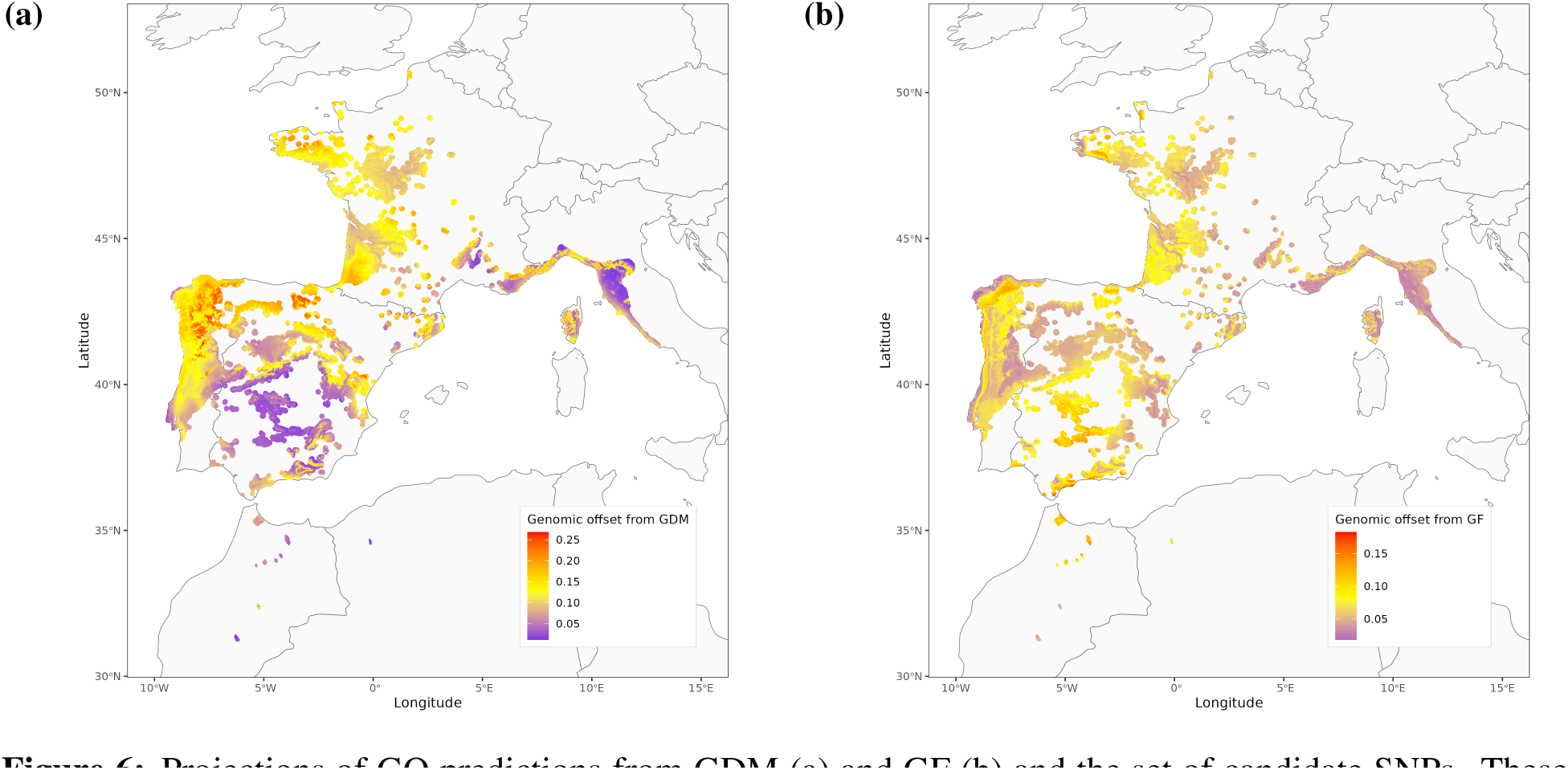
Projections of GO predictions from GDM (a) and GF (b) and the set of candidate SNPs. These predictions correspond to the averaged GO predictions across the five climate general circulation models (GCMs), using the period 1901-1950 for the reference climates, and the period 2041-2060 for the future climates under the shared socio-economic pathway (SSP) 3.7-0.

Importantly, we must remain cautious when interpreting GO predictions, especially when the aim is to use them to inform conservation or management strategies. The limits of the GO approach have already been discussed in depth (Hoffmann et al. 2021, Rellstab et al. 2021, Ahrens et al. 2023), so we will mention here just a few arguments that we consider should be given particular attention in maritime pine, and more generally in forest trees. First, climate change exposure is integrated within the GO approach using distances between long-term means of past and future climatic conditions. However, population vulnerability to climate change will also strongly depend on the variation and correlation of the climatic variables, and the magnitude and probability of climatic extremes (Parmesan et al. 2000, Nadeau and Fuller 2015). For example, in forest trees, die-offs have been attributed to increasingly frequent and intense drought events (Allen et al. 2010, 2015). Second, vulnerability to climate change is not only determined by the risk of maladaptation but also by the ability of a population to respond to the changes. This intraspecific response capacity relies on three mechanisms: phenotypic plasticity, dispersal ability and adaptive potential (IPCC 2007, Foden et al. 2019). Most tree species show high phenotypic plasticity (Nicotra et al. 2010, Benito Garzón et al. 2019), long-distance gene flow facilitating the spread of beneficial alleles (Kremer et al. 2012), and high levels of genetic variation within populations (Scotti et al. 2016). Since these components of population response capacity are not integrated into the GO approach, any credible interpretation of its predictions must take this context into account.

## 5 Conclusion

Our study confirms the great potential value of GO predictions, even in tree species with strong population genetic structure. However, the empirical validity of GO approaches is variable, which will continue to undermine confidence in GO predictions until the conditions under which they are robust are properly understood. Towards that end, developing a more solid theoretical framework around the GO concept appears necessary. Progress has already been made in defining a quantitative theory for GO statistics (Gain et al. 2023) and identifying limitations of the GO approach (Hoffmann et al. 2021, Rellstab et al. 2021, Ahrens et al. 2023). Empirical validation based on multiple data sources (e.g., natural populations vs common gardens, multi-environment common gardens, multiple traits, multiple species, etc) will also be needed before the GO concept can be confidently applied in conservation and management strategies as a metric of maladaptation.

## Acknowledgments

We thank A. Saldaña, F. del Caño, E. Ballesteros and D. Barba (INIA) and the ‘Unité Expérimentale Forêt Pierroton’ (UEFP, INRAE; https://doi.org/10.15454/1.54832646991 93726E12) for field assistance (plantation and measurements). Data used in this research are part of the Spanish Network of Genetic Trials (GENFORED, http://www.genfored.es). We thank all persons and institutions linked to the establishment and maintenance of field trials used in this study. We are especially grateful to Ricardo Alía, Christophe Plomion and Juan Majada who initiated and supervised the establishment of the CLONAPIN network (i.e., the five common gardens used in the validation part). We thank Paloma Ruiz-Benito, Thibaut Capblancq and Sylvain Schmitt for their constructive comments. This study was funded by the European Union’s Horizon 2020 research and innovation programme under grant agreements No 862221 (FORGENIUS) and No 101081774 (OPTFORESTS), and by the newLEAF project (NE/V019813/1) under the UK’s ‘Future of UK Treescapes’ programme, which was led by UKRI-NERC, and jointly funded by UKRI-AHRC & UKRI-ESRC, with contributions from Defra and the Welsh and Scottish governments. Views and opinions expressed are however those of the author(s) only and do not necessarily reflect those of the European Union. Neither the European Union nor the granting authority can be held responsible for them.

## Author contributions

SCG-M designed the experiment and supervised the curation of field data. MdM cleaned and formatted the phenotypic data in the common gardens. MM extracted the climatic values at the location of the populations and the National Forest Inventory (NFI) plots, and provided the raster files to project the genomic offset predictions across the species range. F Bagnoli, SCG-M and GGV did the DNA extractions, production, cleaning and checking of the genomic data. AC cleaned and formatted mortality data from the NFI plots. SCG-M, JA and MBG conceived the paper methodology. JA and F Barraquand built the Bayesian models and codes in the validation steps. JA conducted the data analyses. JA led the writing of the manuscript. All authors interpreted the results, contributed to the writing of the manuscript and gave final approval for publication.

## Data and code availability

Data and code were deposited in the Dryad repository at https://doi.org/10.5061/dryad.bnzs7h4jt. The scripts are also available in the following Github repository: https://github.com/JulietteArchambeau/GOPredEvalPinpin.

