## Supplementary Information for "Evaluating genomic offset predictions in a forest tree with high population genetic structure"

### Supplementary Information of the paper: 'Evaluating genomic offset predictions in a forest tree with high population genetic structure'

2024-05-16

**References** **139**

### 1 Glossary

**ClimateDT**: Climate Downscaling Tool. A geo-web service providing a large number of climatic variables downscaled at 1-km resolution between 1901 and 2098.

**CTD**: Climatic transfer distance. An index calculated independently for each climatic variable and which is equal to the absolute difference between the values of the climatic variable at the location of the populations and at the location of the common gardens.

**dbMEMs**: Distance-based Moran's Eigenvector Maps. Orthogonal vectors calculated based on a geographic distance matrix and maximizing the spatial autocorrelation (measured by Moran's coefficient) of the sampled locations.

**GCM**: General circulation model (also called Global Climate Model). Numerical models representing physical processes in the atmosphere, ocean, cryosphere and land surface and simulating the response of the global climate system to increasing greenhouse gas concentrations. They are used for forecasting climate change under different scenarios of greenhouse gas emissions. In the present study, we used the five GCMs available in ClimateDT and based on UKCP18 projections, namely GFDL-ESM4, IPSL-CM6A-LR, MPI-ESM1-2-HR, MRI-ESM2-0 and UKESM1-0-LL.

**GDM**: Generalized Dissimilarity Modelling. One of the methods used to generate GO predictions at the location of the populations, the common gardens and the NFI plots (section Section 7.4).

**GF**: Gradient Forest. One of the methods used to estimate the gene-climate relationships (section Section 5.2) and to generate GO predictions at the location of the populations, the common gardens and the NFI plots (section Section 7.2).

**GO**: Genomic offset. An index which is expected to provide an estimate of the adaptive shift required to track climate change.

**LFMM**: Latent factor mixed models. One of the methods used to estimate the gene-climate relationships (section Section 5.3) and to generate GO predictions at the location of the populations, the common gardens and the NFI plots (section Section 7.3).

**NFI**: National Forest Inventory. In the present paper, we used mortality data from the French and Spanish NFIs harmonized in Changenet et al. (2021) (section Section 8).

**PCA**: Principal component analysis.

**RDA**: Redundancy Analysis. One of the methods used to estimate the gene-climate relationships (section Section 5.1) and to generate GO predictions at the location of the populations, the common gardens and the NFI plots (section Section 7.1).

### 2 Genomic data

#### 2.1 Sample size

Table 1: Number of genotypes in each population.

| Population | Number of genotypes per population |
|---|---|
| ALT | 8 |
| ARM | 7 |
| ARN | 14 |
| BAY | 13 |
| BON | 8 |
| CAD | 8 |
| CAR | 5 |
| CAS | 8 |
| CEN | 9 |
| COC | 15 |
| COM | 3 |
| CUE | 23 |
| HOU | 25 |
| LAM | 9 |
| LEI | 20 |
| MAD | 1 |
| MIM | 17 |
| OLB | 20 |
| OLO | 23 |
| ORI | 22 |
| PET | 22 |
| PIA | 12 |
| PIE | 9 |
| PLE | 16 |
| PUE | 7 |
| QUA | 16 |
| SAC | 8 |
| SAL | 10 |
| SEG | 19 |
| SIE | 7 |
| STJ | 24 |
| TAM | 12 |
| VAL | 10 |
| VER | 24 |

In the present study, we used 34 populations and 454 genotypes. There were between 1 and 25 genotypes per population, with an average of about 13.4 genotypes per population.

#### 2.2 Population neutral genetic structure

The population neutral genetic structure was inferred with a PCA on a set of 10350 SNPs including SNPs with minor allele frequencies (Figure S1). We did not remove SNPs with minor allele frequencies because small genetic variation can be informative to differentiate the neutral genetic groups (i.e. hereafter 'gene pools'). We retained the first three PCs of the PCA as proxy of population evolutionary history in the RDA.

Figure 1: Principal component analysis of the genomic data. Each point corresponds to a population and the colors correspond to the main gene pool of each population. No minor allele frequency filter was applied because small genetic variation can be informative to differentiate the neutral genetic groups.

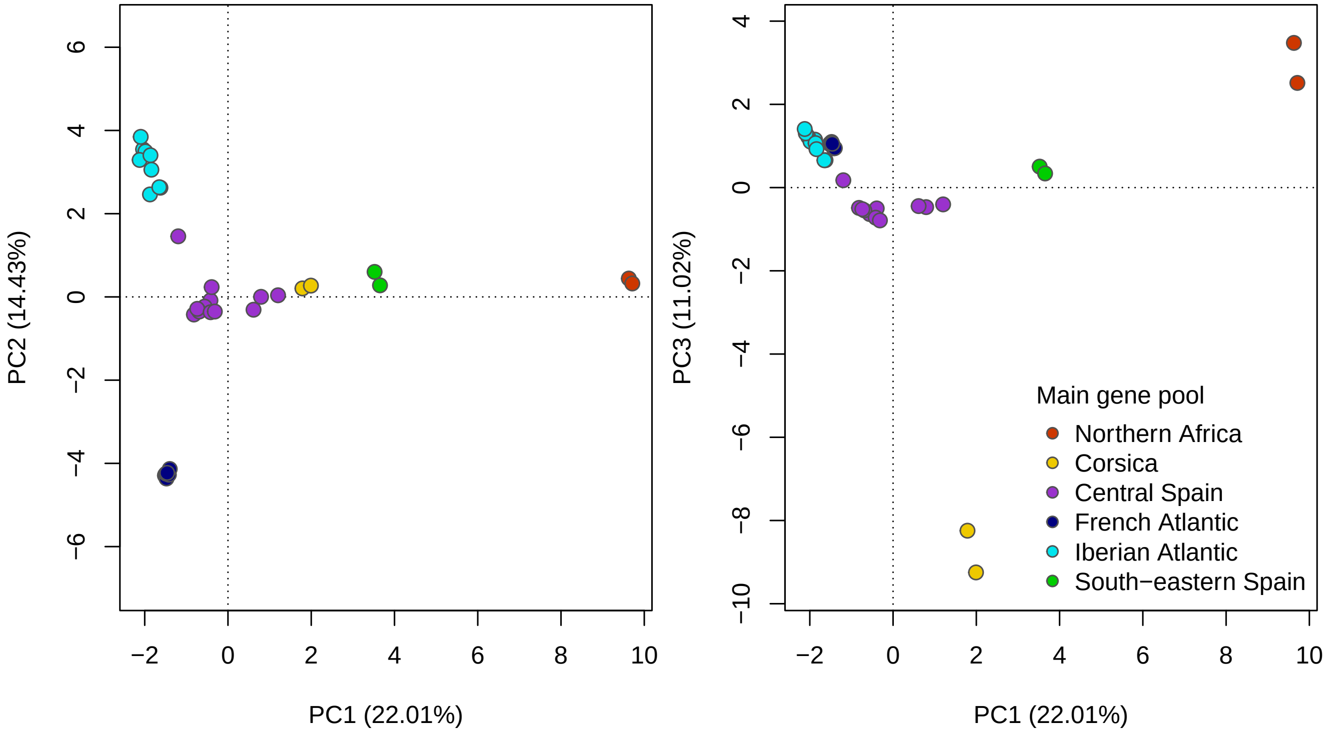

### 3 Selection of the climatic variables

We extracted the climatic data from the Climate Downscaling tool (ClimateDT) developed within the framework of the B4EST project. We used the time period 1901-1950 as reference period, i.e. which is expected to capture the climatic conditions under which the populations evolved and are currently locally adapted. We used predictions of future climates for the period 2041-2060 under the shared socio-economic pathway 3-7.0 (SSP3-7.0) and from five different GCMs, namely: GFDL-ESM4, IPSL-CM6A-LR, MPI-ESM1-2-HR, MRI-ESM2-0 and UKESM1-0-LL.

#### 3.1 Preselection step

In a preselection step, we removed climatic variables that (i) had reduced biological interest for the study goals, (ii) were too highly correlated with any other variable ($\rho > 0.95$), or (iii) that had no meta-information on the ClimateDT website. The remaining climatic variables are described in Table S2.

Table 2: Preselected climatic variables

| Label | Description | Unit |
|---|---|---|
| bio1 | Mean annual temperature | Celsius degrees (°C) |
| bio3 | Isothermality (bio2/bio7) (×100) | Index |
| bio4 | Temperature seasonality (standard deviation ×100) | Celsius degrees (°C) |
| bio5 | Max temperature of warmest month | Celsius degrees (°C) |
| bio6 | Min temperature of coldest month | Celsius degrees (°C) |
| bio8 | Mean temperature of wettest quarter | Celsius degrees (°C) |
| bio9 | Mean temperature of driest quarter | Celsius degrees (°C) |
| bio12 | Annual precipitation | Millimeters (mm) |
| bio15 | Precipitation seasonality (coefficient of variation) | Index |
| Eref | Hargreaves reference evaporation | Millimeters (mm) |
| SP | Summer precipitation | Millimeters (mm) |
| MWMT | Mean warmest month temperature | Celsius degrees (°C) |
| AHM | Annual heat moisture index | °C / mm |
| SHM | Summer heat moisture index | °C / mm |

Figure 2: Correlation matrix including the geographical coordinates (i.e. latitude and longitude), elevation and climatic variables at the location of the populations. The variables noted as MEM$X$ correspond to the distance-based Moran's eigenvector maps (dbMEMs) with positive eigenvalues calculated based on the geographical coordinates of the populations with the function `dbmem` of the `adespatial` R package v0.3.23 (**adespatial2022a?**).

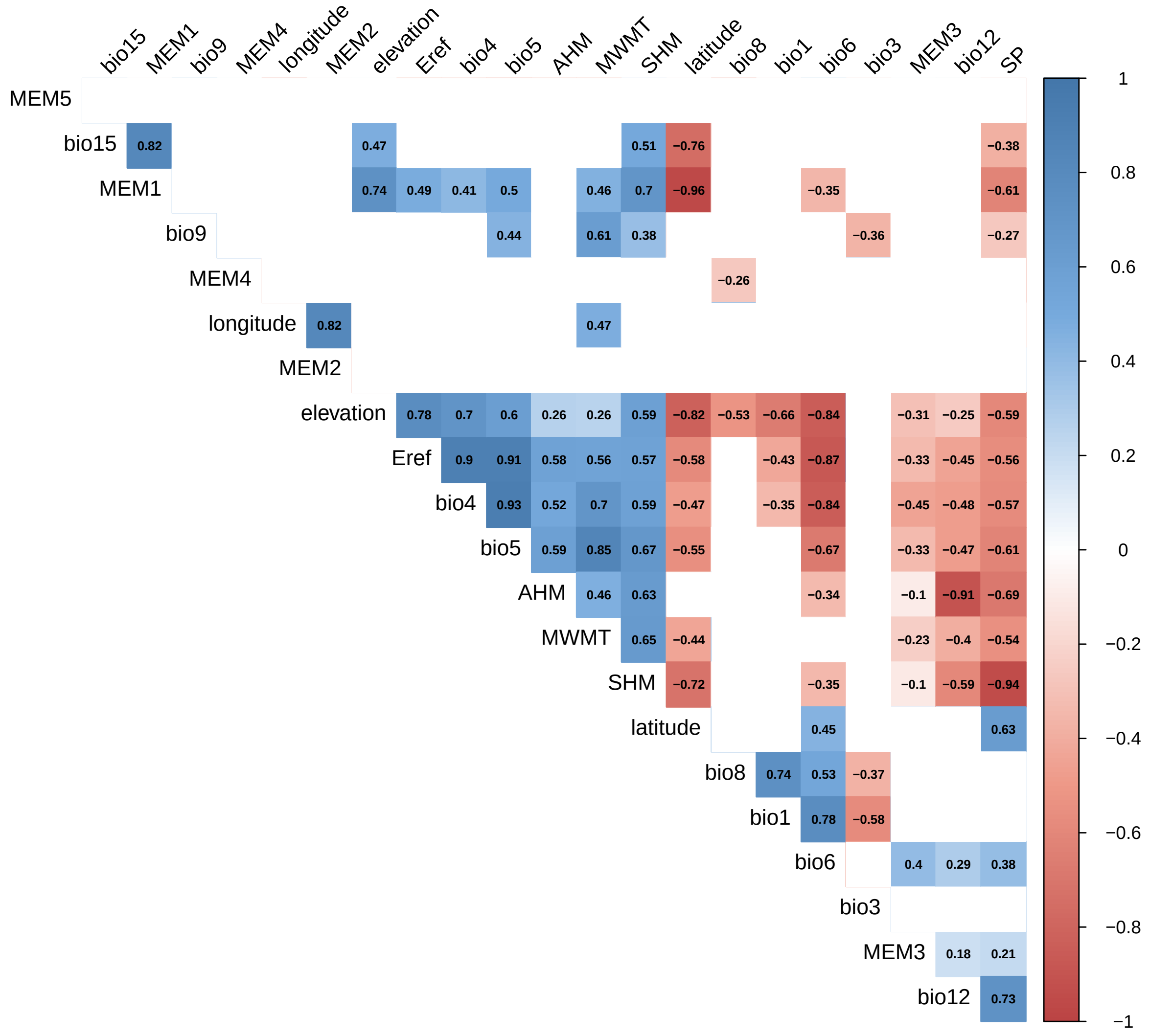

#### 3.2 Selection of the climatic variables

Based on the reduced set of climatic variables from Table S2, we then selected the final set of climatic variables based on three criteria:

1. biological relevance
2. contribution to genomic variation.

3. magnitude of the differences between the current and future values of the climatic variables.

The final set of climatic variables is shown in Table S3 and their distributions are shown in Figure S3.

Table 3: Climatic variables selected for the gene-environment association analyses and the GO predictions.

| Label | Description | Unit |
|---|---|---|
| bio1 | Mean annual temperature | Celsius degrees (°C) |
| bio3 | Isothermality (bio2/bio7) (×100) | Index |
| bio4 | Temperature seasonality (standard deviation ×100) | Celsius degrees (°C) |
| bio12 | Annual precipitation | Millimeters (mm) |
| bio15 | Precipitation seasonality (coefficient of variation) | Index |
| SHM | Summer heat moisture index | °C / mm |

Figure 3: Distributions of the climatic variables selected for the gene-environment association analyses and the GO predictions.

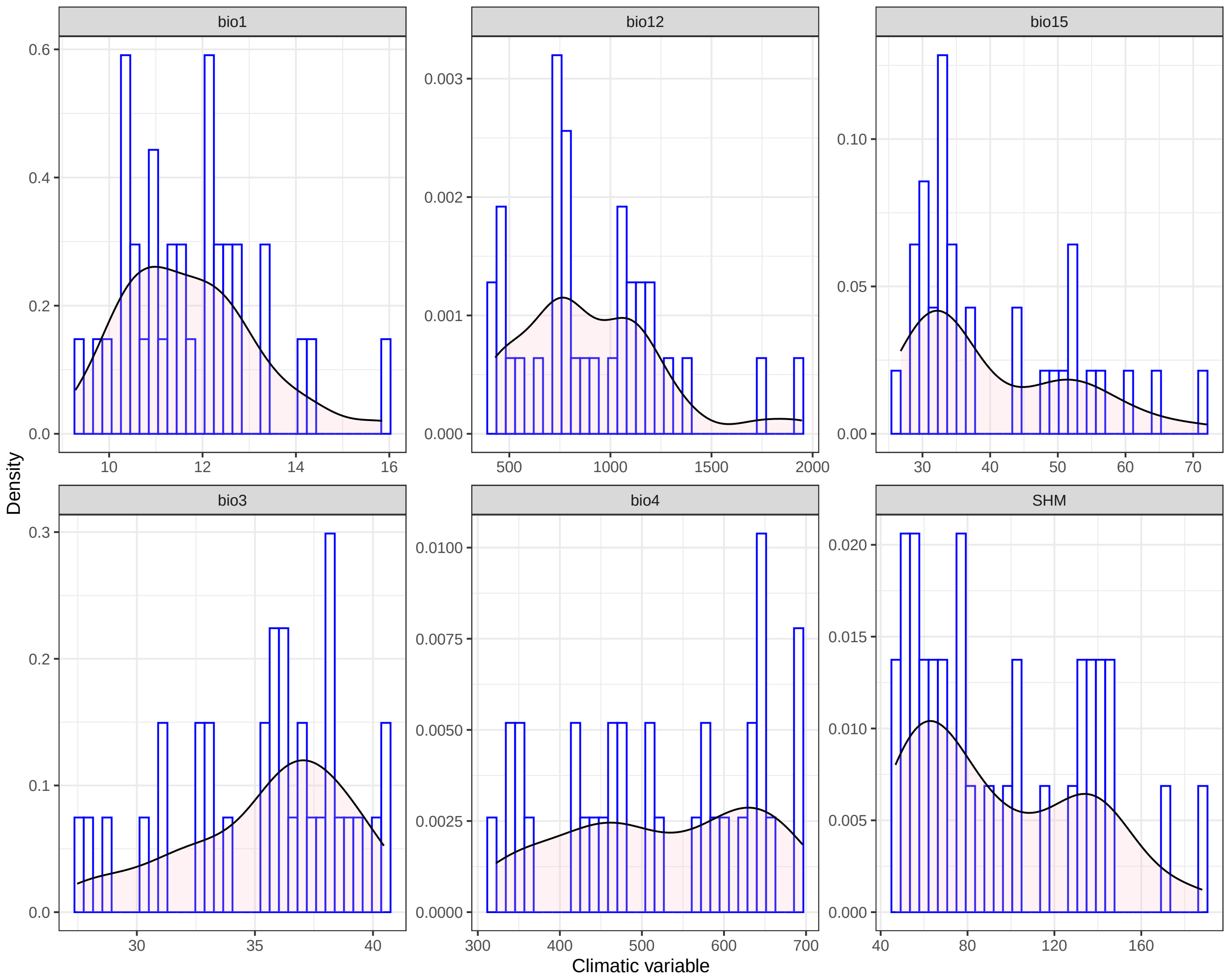

##### 3.2.1 Criteria 1: biological relevance

We aimed to provide GO predictions capturing both the changes in annual climatic conditions (e.g. mean annual temperature or precipitation) and in seasonal climatic conditions (e.g. summer droughts).

**Winter cold temperatures:** Previous studies found that maritime pine populations show strong patterns of adaption to temperatures, and especially cold temperatures (Archambeau et al. 2023; Grivet et al. 2011). We did not include climatic variables related to winter cold temperatures because, in the context of climate change, the expected increase in winter cold temperatures may benefit maritime pine populations. In such a scenario, populations undergoing strong variations in winter cold conditions will have a high GO, which will not reflect a potential maladaptation but will, on the contrary, inform about a potential increase in their fitness under climate change. Increased winter cold temperatures may also impact the survival of young trees, the reproductive ability of adult trees or the dynamics of pests and pathogens. However, it is not clear how this negative impacts may counterbalance the positive effects of cooler winter, and that is why we did not include climatic variables related to winter cold temperatures in the set of climatic variables used to make GO predictions.

##### 3.2.2 Criteria 2: contribution to genomic variation.

We used a RDA-based stepwise selection procedure to identify the set of climatic variables maximizing the genetic variance explained (see for instance Capblancq et al. 2021). We used the `ordiR2step` function of the package `vegan`, in which we have to specify two models:

- a *null* model where genomic variation is explained only by an intercept.
- a *full* model including as predictors all the climatic variables shown in Table S2.

For including new variables during the selection procedure, we used the default stopping criteria of the `ordi2step` function: variable significance of $p < 0.01$ using 1000 permutations and the comparison of adjusted variation ($R^2_{adj}$) explained by the selected variables to $R^2_{adj}$ explained by the full model. This selection criteria means that if the new variable is not significant or the $R^2_{adj}$ of the model including the new variable does not exceed the $R^2_{adj}$ of the full model, the selection procedure stops.

We performed 100 iterations of the stepwise selection procedure and counted the number of times each climate variable was selected. The results are reported in Table S4.

Table 4: Number of times that each climatic variable was selected among the 100 iterations of the RDA-based stepwise selection procedure.

| variable | count |
|---|---|
| bio15 | 100 |
| bio3 | 89 |
| bio4 | 100 |

##### 3.2.3 Criteria 3: exposure to climate change

To compare the values of the climatic variables under current and future climates, we calculated the relative climatic distance $D$ for each climatic variable $x$:

$$D_x = \frac{\mu_{fut} - \mu_{ref}}{\mu_{ref}}$$

where $\mu_{ref}$ is the average of the climatic variable of interest over the reference period (i.e. 1901-1950) and $\mu_{fut}$ is the average of the predictions for the climatic variable over the future period 2041-2070 and under the shared socio-economic pathway (SSP) 3-7.0.

Figure 4: Relative climatic distances for the six climatic variables used to make GO predictions. The relative climatic distances were calculated for five different GCMs, each corresponding to a different panel.

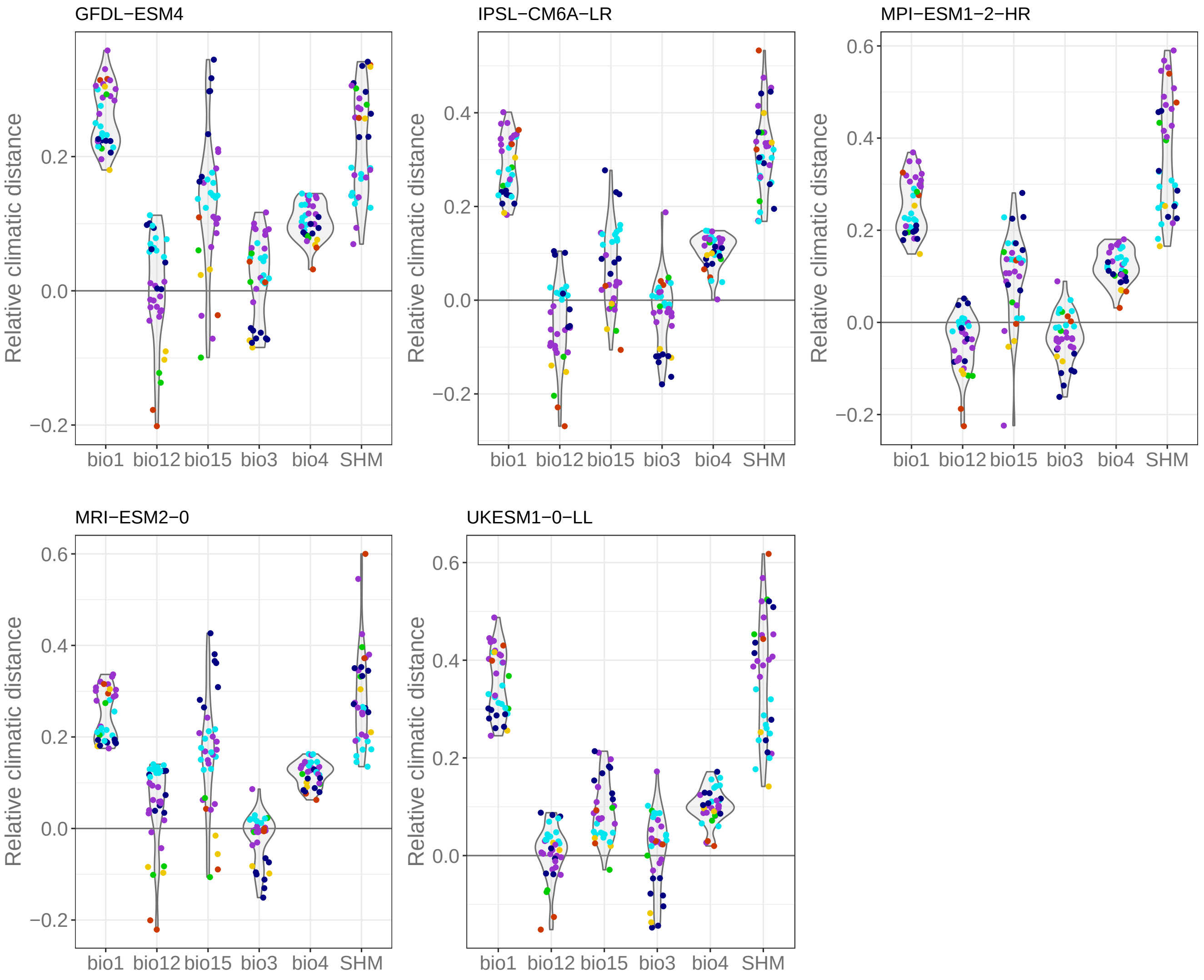

### 4 Variance partitioning of the genetic variation

Following Capblancq & Forester (2021), we used a combination of RDA and partial RDA (pRDA) to estimate the proportion of genetic variation that can be or cannot be uniquely attributed to climate, neutral population structure and geography. The response variable was the allele frequencies of the populations. The climate was represented by the six climatic variables that were selected in the previous section (Table S3), i.e. the climatic variables that are then used to identify the loci potentially involved in local adaptation (see the gene-environment association analyses in the section below) and to make GO predictions. The neutral population genetic structure was accounted for with the first three PCs of the PCA based on the genomic data not filtered for MAF (Figure S1). The geography was either accounted for with the geographical coordinates of the populations or with the distance-based Moran's eigenvector maps (dbMEMs) with positive eigenvalues and calculated based on the geographical coordinates of the populations with the function `dbmem` of the `adespatial` R package v0.3.23 (**adespatial2022a?**).

We first used an RDA model including all explanatory variables (i.e. climate, neutral genetic structure and geography), and named the 'full model'. The full model provides the total amount of genetic variance (i.e. inertia) explained by the explanatory variables together. We then used three pRDA models, named the 'pure models', to estimate the contribution of genetic variance that can be uniquely attributed to each explanatory variable. For that, for the three pRDA models, we used either the climate, geography and neutral population structure as explanatory variables, while conditioning on the remaining two variables.

These analyses were conducted using the function `rda` of the `vegan` R package v2.6.4 (Oksanen et al. 2022).

Table 5: Variance partioning using geographical coordinates to account for geography.

| RDA models | Total exp. variance | Relative exp. variance | P-value |
|---|---|---|---|
| Full model: Y ~ clim + geo + pgs. | 0.76 | 1.00 | 0.00 |
| Pure climate model: Y ~ clim \| (geo + pgs) | 0.10 | 0.14 | 0.00 |
| Pure pop. gen. structure model: Y ~ pgs \| (geo + clim) | 0.22 | 0.30 | 0.00 |
| Pure geography model: Y ~ geo \| (pgs + clim) | 0.04 | 0.05 | 0.01 |

Table 6: Variance partioning using dbMEMs with positive eigenvalues to account for geography.

| RDA models | Total exp. variance | Relative exp. variance | P-value |
|---|---|---|---|
| Full model: Y ~ clim + geo + pgs. | 0.80 | 1.00 | 0.00 |
| Pure climate model: Y ~ clim \| (geo + pgs) | 0.08 | 0.10 | 0.04 |
| Pure pop. gen. structure model: Y ~ pgs \| (geo + clim) | 0.14 | 0.17 | 0.00 |
| Pure geography model: Y ~ geo \| (pgs + clim) | 0.08 | 0.10 | 0.00 |

In Tables S5 and S6, `Y` refers to the genomic data (i.e. the population allele frequencies filtered for MAF), `clim` refers to climate, `pgs` refers to the population neutral genetic structure and `geo` refers to the geography.

As dbMEMs explain more genetic variation than the geographical coordinates, we only provide the proportions of variance explained obtained with dbMEMs in the manuscript and figure below.

Figure 5: Variance partitioning of genomic variation using a combination of RDA and pRDA models. The response variable was the population allele frequencies and the explanatory variables were climate (the five climatic variables used for GO predictions), geography (distance-based Moran's eigenvector maps) and the neutral genetic structure (first three axes of a PCA based on genomic data).

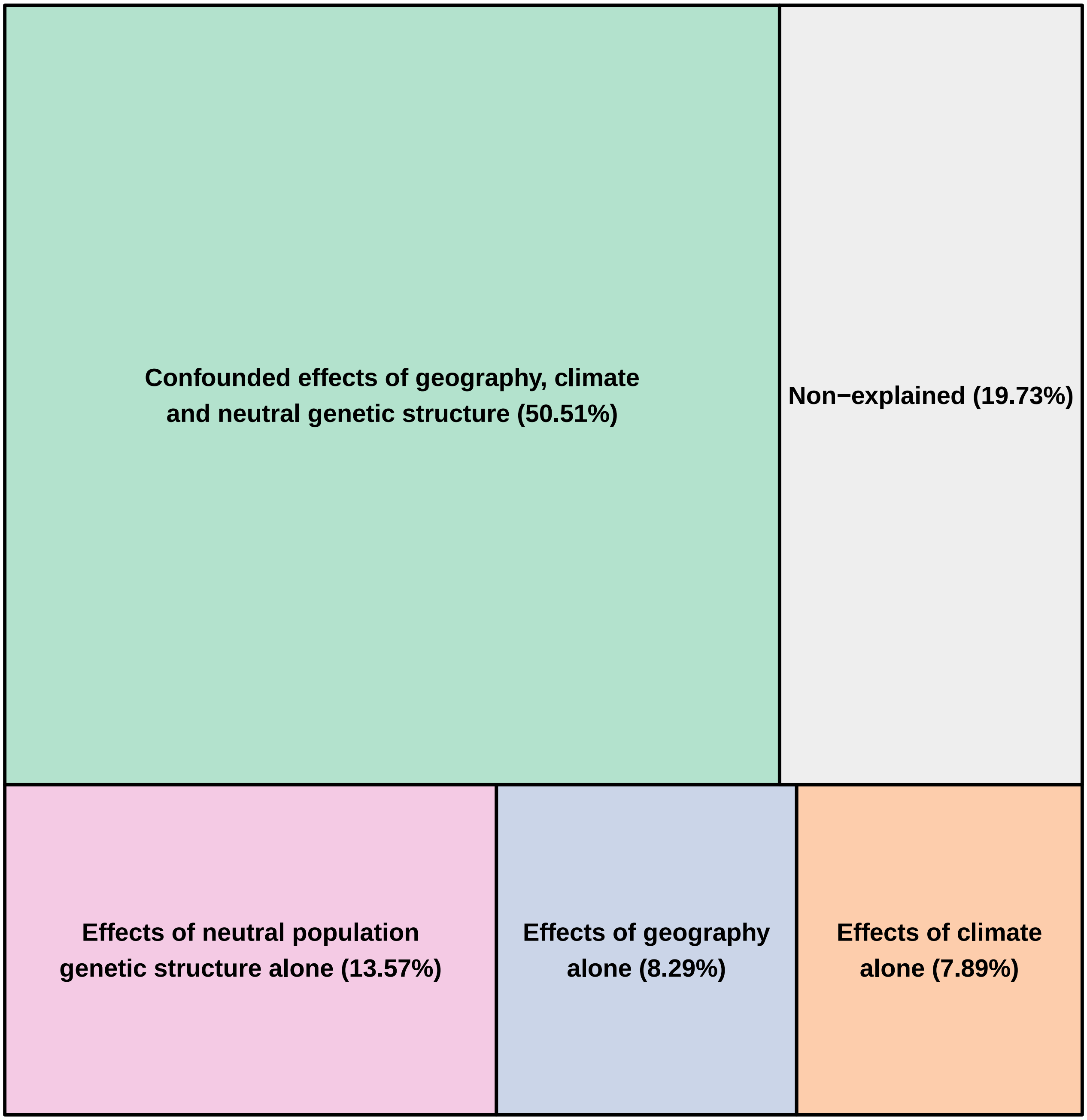

### 5 Gene-environment association methods

#### 5.1 Redundancy analysis (RDA)

RDA was performed with the `vegan` R package v2.6.4 (Oksanen et al. 2022). We identify the outlier SNPs based on:

- a RDA not corrected for population structure, hereafter referred as *RDA*.
- a partial RDA corrected for population structure (using the first three PC scores, see Figure S1), hereafter referred as *pRDA*.

Following Capblancq et al. (2018), outlier SNPs were identified based on their extremeness along a distribution of Mahalanobis distances estimated between each locus and the center of the RDA space using the first two RDA axes, i.e. the two axes that retain most of the genetic variation in our study. The $p$-values and $q$-values of the Mahalonobis distances were calculated using the `radapt` function from the Github repository associated with Capblancq & Forester (2021). The `radapt` function relies on the `covRob` function from the `robust` R package v0.7.4 (Wang et al. 2024) and the `qvalue` function from the `qvalue` R package (Storey et al. 2024). We used a false discovery threshold (FDR) of 5% to identify the outliers, which means that 5% or less of the identified outliers may be false positives.

Figure 6: Visualization of the outlier SNPs identified with the RDA. Left panels: RDA space. Right panels: Manhattan plots. The gray line in the Manhattan plots corresponds to the Bonferroni threshold, i.e. SNPs with $p$-values < 0.01 / number of SNPs.

(a) RDA

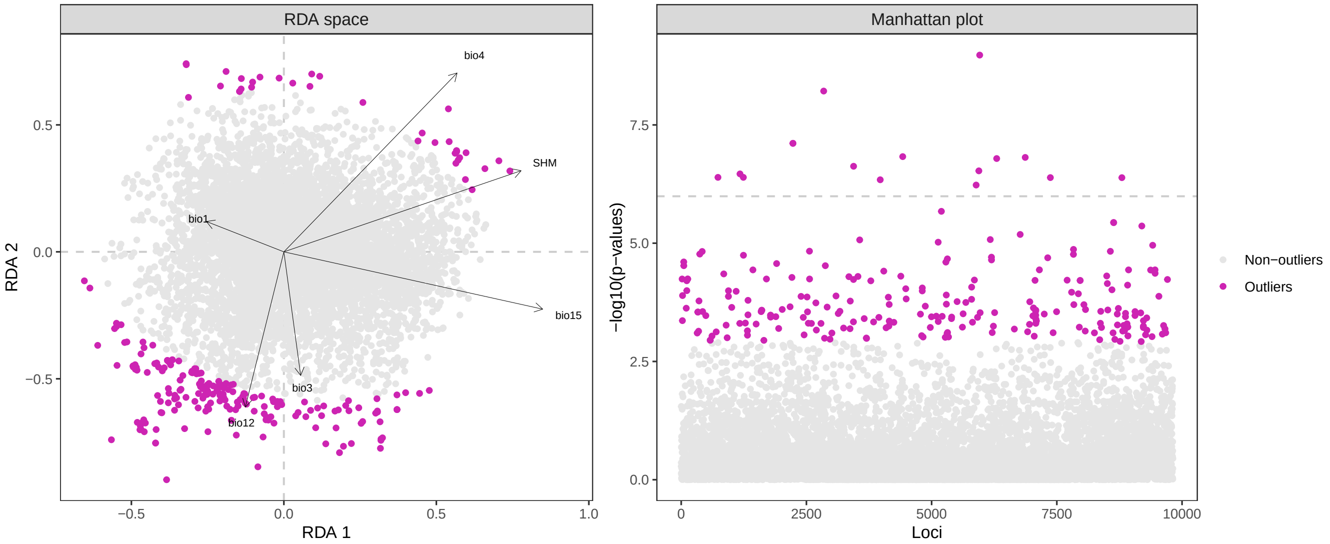

(b) pRDA

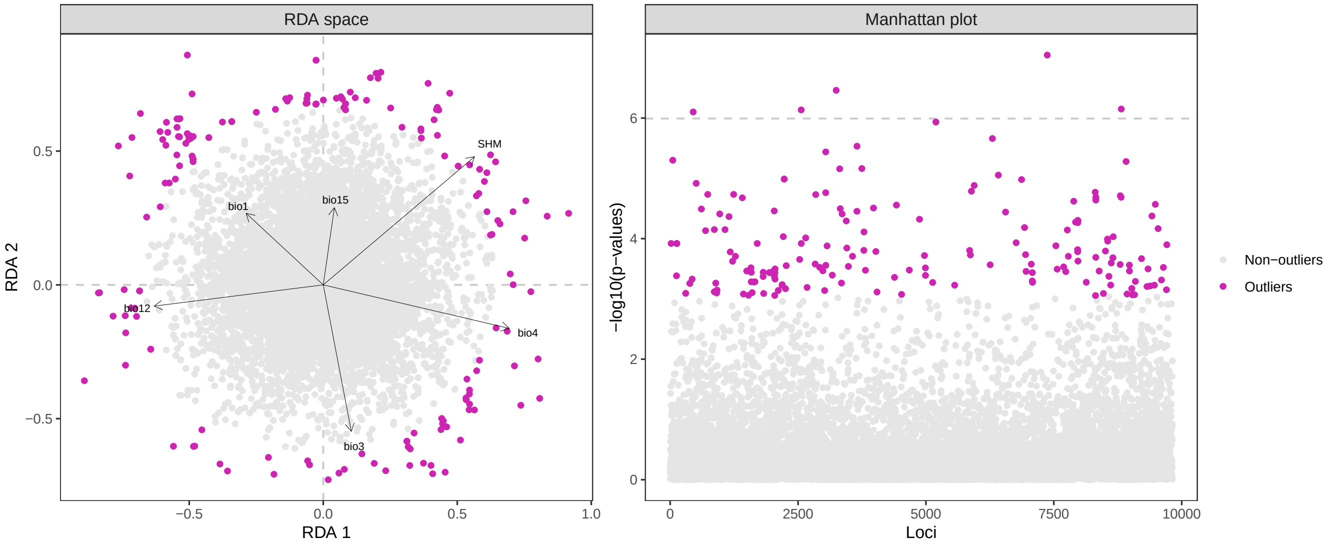

#### 5.2 Gradient Forest (GF)

We identified outlier SNPs with the GF algorithm following Fitzpatrick et al. (2021) and Capblancq et al. (2023). GF models were fitted to each locus individually. We then compared the importance ($R^2$) value of each locus to the $R^2$ distribution of all SNPs by calculating empirical $p$-values, an approach described in Lotterhos et al. (2014). More specifically, the empirical $p$-value $\hat{p}$ of each locus $l$ is calculated as follows:

$$\hat{p}_l = 1 - (r_l/N)$$

with $r_l$ being the rank of the $R^2$ value of locus $l$ within the distribution of $R^2$ values of the all SNPs ($N$ SNPs).

Note that selecting the SNPs based on the rank of their empirical $p$-value is equivalent to selecting them based on the rank of their $R^2$ value, i.e., the 0.5% of the SNPs with the highest $R^2$ should match the 0.5% of the SNPs with the lowest empirical $p$-value.

We ran three independent GF runs and identified the SNPs that were in the 0.5% SNPs with the lowest empirical $p$-values across the three runs.

Figure 7: Venn diagram of the number of SNPs that were in the 0.5% SNPs with the lowest empirical $p$-values across three independant runs of the GF-based GEA.

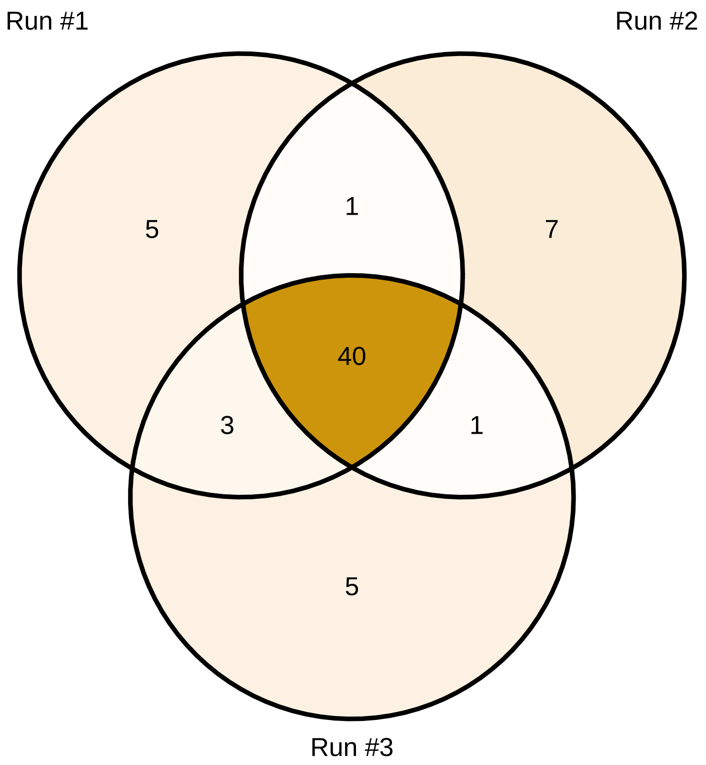

40 SNPs were identified as outliers in the three runs and were kept for the following analyses.

**Comment:** To our knowledge, GF has only been used once as a genome scan (Capblancq et al. 2023) and its ability to identify loci underlying local adaptation has still not been evaluated and confronted to other GEAs. We performed several runs of GF models as a genome scan and there was little overlap in the outliers identified across runs. This is in part why we only considered SNPs as outliers if they were identified by at least two GEA methods. Like that, if a SNP was identified as an outlier only with the GF algorithm, it was not included in the set of outliers used to estimate the GO.

GF models were implemented with the `gradientForest` R package (Ellis et al. 2012) and the scripts available from the Github repository associated with Fitzpatrick et al. (2021).

#### 5.3 Latent factor mixed models (LFMM)

A latent factor mixed model (LFMM) is a multivariate mixed regression model that estimates simultaneously the effects of environmental variables (fixed effects) and unobserved confounders called latent factors (Caye et al. 2019; Frichot et al. 2013). We used the function `lfmm2` of the `lea` R package to run LFMM and we set the number of latent factors $K$ to 6 as the sampled populations derive from six gene pools (Figure S8; Jaramillo-Correa et al. (2015)).

Figure 8: Cross-entropy criterion based on the genomic data not filtered for minor allele frequencies (MAF).

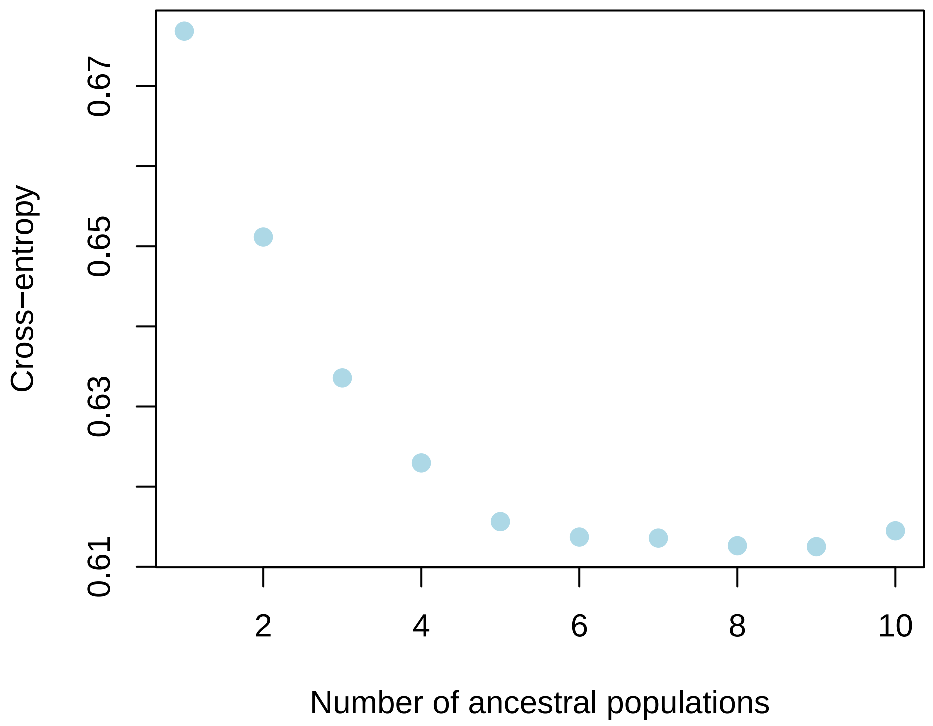

We used the function `lfmm2.test` to calculate the $p$-values with the default options `genomic.control=TRUE` (i.e. $p$-values are recalibrated after correction for confounding) and `full=TRUE` (i.e. $p$-values are calculated for the full set of climatic variables using Fisher tests). Outlier SNPs were then identified with a FDR threshold of 5%.

Figure 9: Manhattan plot showing the outlier SNPs identified with LFMM (SNPs with a red circle). The orange line corresponds to the Bonferroni threshold.

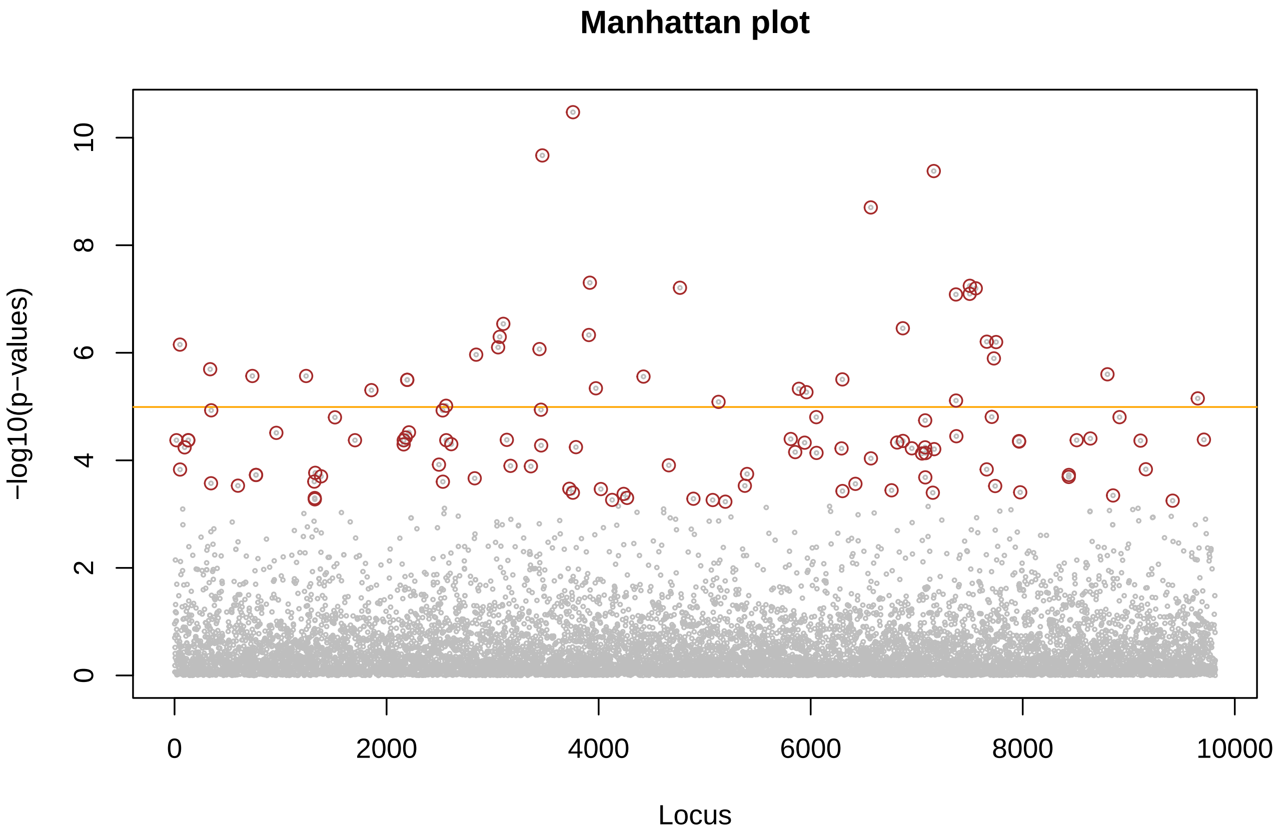

#### 5.4 BayPass

We identified candidate SNPs with the BayPass software, using the standard covariate model and the Important Sampling (IS) approximation.

Using the core model without covariate from BayPass, we first estimated the population covariance matrix with a genomic dataset not filtered for minor allele frequencies, as the latter can be particularly informative to infer the population neutral genetic structure.

Figure 10: Heatmap of the population covariance matrix estimated with BayPass

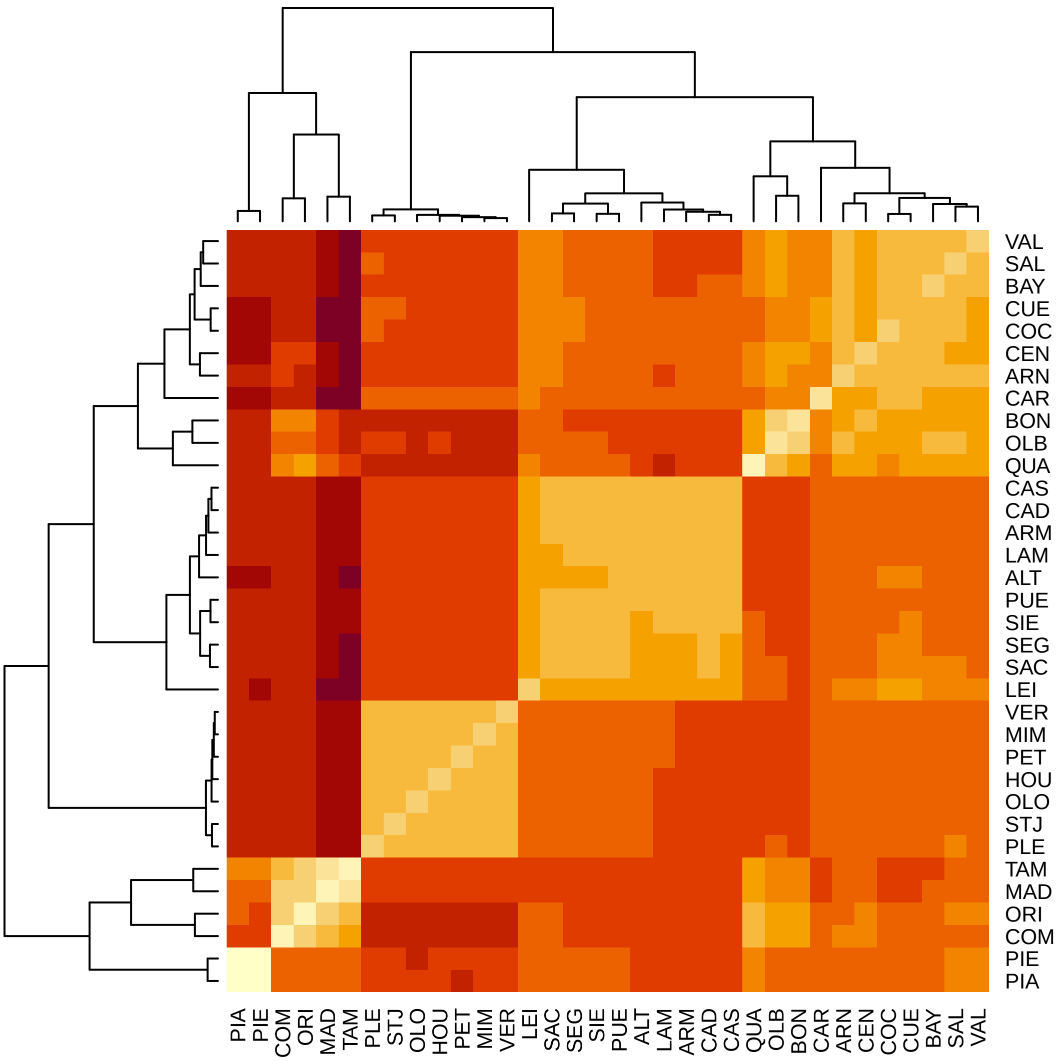

#### 6 Set of control and candidate SNPs

Common outliers across GEAs:

- 240 outliers were identified with the RDA.
- 178 outliers were identified with the pRDA.
- 40 outliers were identified with GF.

- 120 outliers were identified with LFMM.
- 46 outliers were identified with BayPass.

For the following analyses, we kept the 85 outlier SNPs identified by at least two GEAs. Our goal was to get a set of candidate SNPs enriched for loci involved in adaptation to climate, with a good balance between not being too conservative (e.g. by selecting SNPs in common across all methods) and not including too many false positives (e.g. by selecting SNPs identified by at least one GEA). Selecting SNPs identified by at least two GEAs also allowed us to select candidate SNPs that can only be identified with GEAs that correct for population structure (pRDA, LFMM, BayPass), or candidate SNPs that can only be identified with methods that do not correct for population structure (RDA and GF).

Among the 85 outlier SNPs, we identified outlier SNPs located on the same scaffold/contig (a scaffold/contig measuring approximately between 400 and 10,000 bp). If two (or more) outlier SNPs were located on the same scaffold/contig, we kept the SNP with the lowest $p$-value in the RDA.

After this filtering step, there were 69 candidate SNPs left.

Last, we randomly sampled a set of control SNPs (with the same number as the candidate SNPs) from SNPs that were not identified by any of the GEA methods.

### 7 GO predictions

We predicted the GO with four methods: Redundancy analysis (RDA), Gradient Forest (GF), latent factor mixed models (LFMM) and Generalized Dissimilarity Modelling (GDM). For each method, GO predictions were generated at the locations of the maritime pine populations using the period 1901-1950 as reference period (i.e., past climates under which the populations evolved) and the projections from five GCMs under the period 2041-2060 for the future climates (SSP3.7-0). The five GCMs used were GFDL-ESM4, IPSL-CM6A-LR, MPI-ESM1-2-HR, MRI-ESM2-0 and UKESM1-0-LL.

#### 7.1 Redundancy analysis (RDA)

The methods to generate GO predictions with RDA are based on Capblancq et al. (2023) and Capblancq & Forester (2021) (and the associated Github repository).

For each set of candidate and control SNPs, a RDA was performed with the set of SNPs as multivariate response and the set of climatic variables as explanatory variables. For the set of candidate SNPs, the RDA space resulting from this analysis can be considered as an 'adaptively enriched genetic space' (Capblancq & Forester 2021; Capblancq et al. 2023).

Figure 11: RDA biplot showing the association between the selected SNPs (i.e. candidate SNPs for adaptation to climate or control SNPs) and the set of climatic variables in the RDA space. The full name and units of the climatic variables can be found in Table S3.

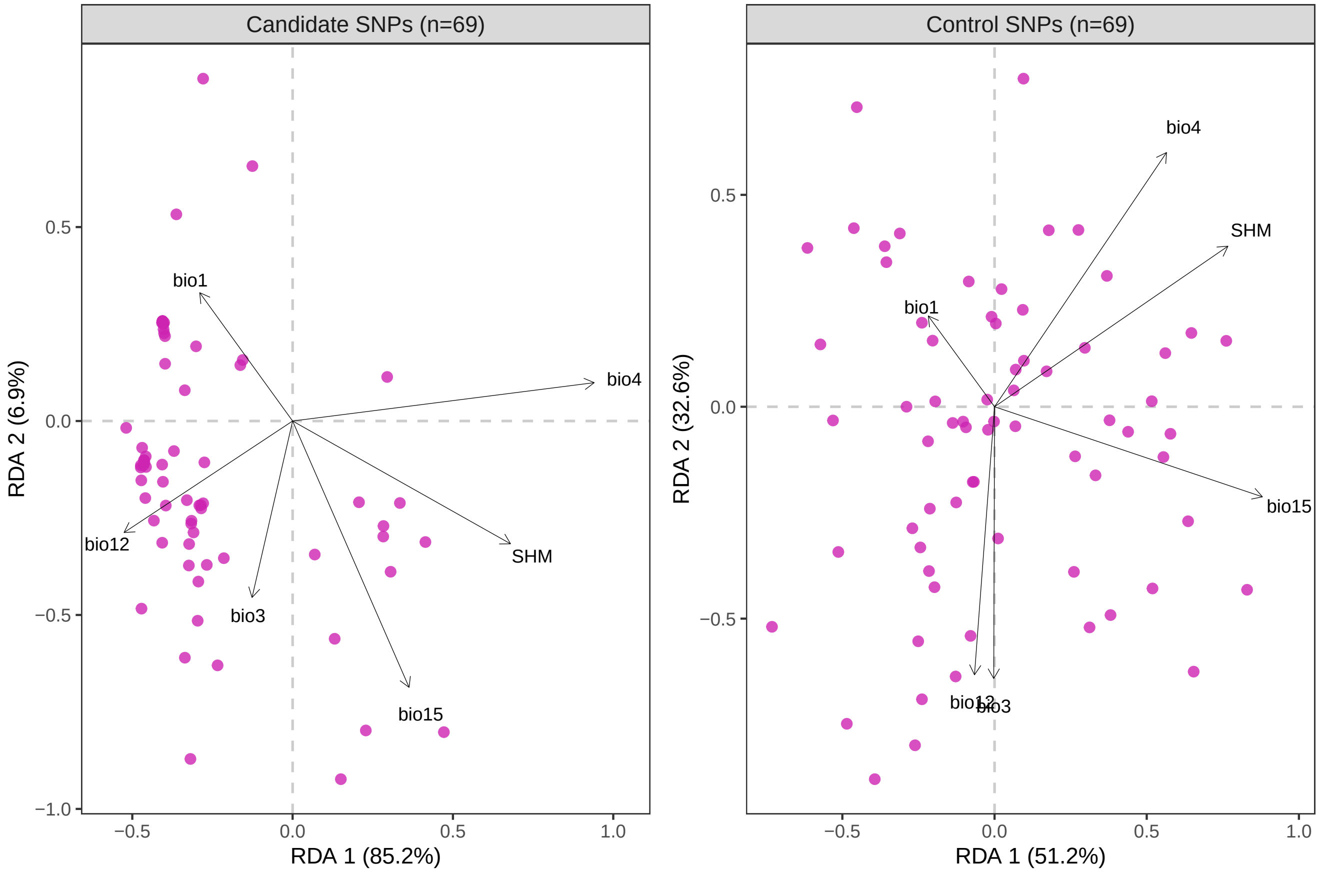

We used the scores of the climatic variables along the RDA axes to calculate a genetic-based index for each pixel of the landscape. For each RDA axis, the index is calculated as follows:

$$\sum_{i=1}^{n} a_i b_i$$

where $i$ is one of the $n$ climatic variables used in the RDA model, $a$ is the score of the climatic variable $i$ along the RDA axis and $b$ is the standardized value for the climatic variable $i$ at the focal pixel.

When calculating based on the set of candidate SNPs, this index can be considered as an adaptive index that provides an estimate of the adaptive genetic similarity or difference of all pixels on the landscape as a function of the values of the climatic predictors at that location.

Figure 12: Spatial projection of the predicted current genomic composition through extrapolation of the RDA model across maritime pine range. Pixels with similar colors are expected to have a similar genetic composition.

(a) Candidate SNPs.

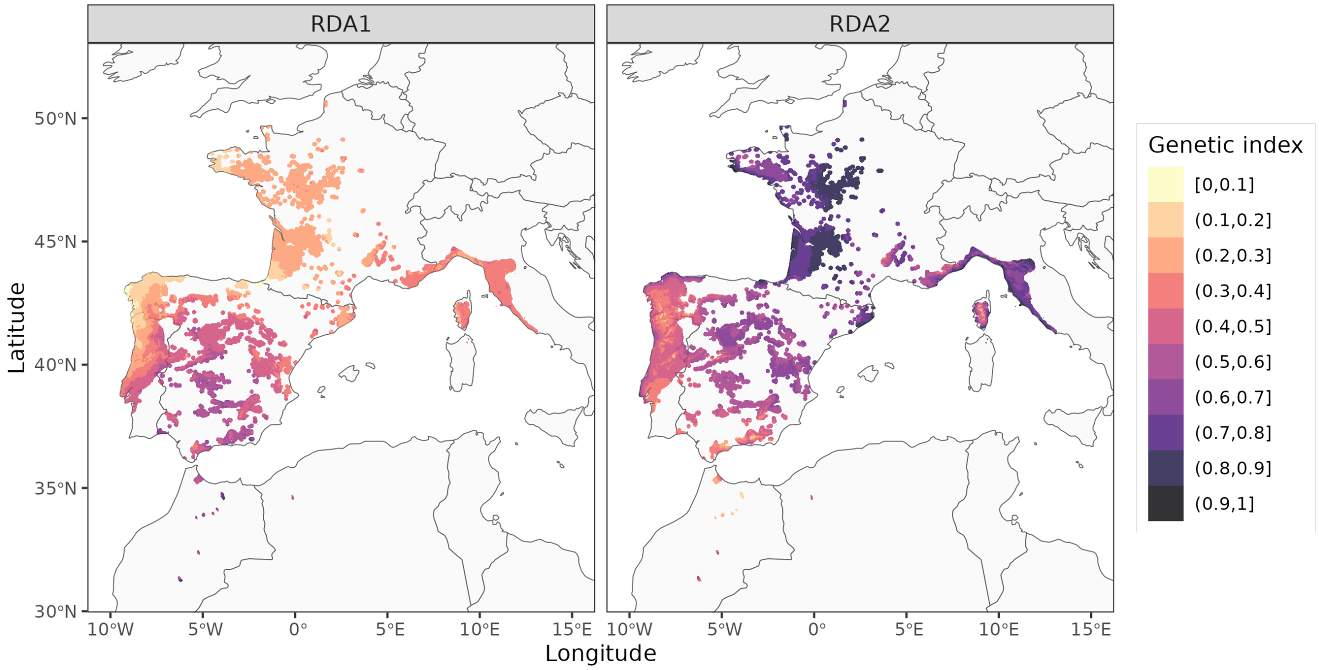

(b) Control SNPs.

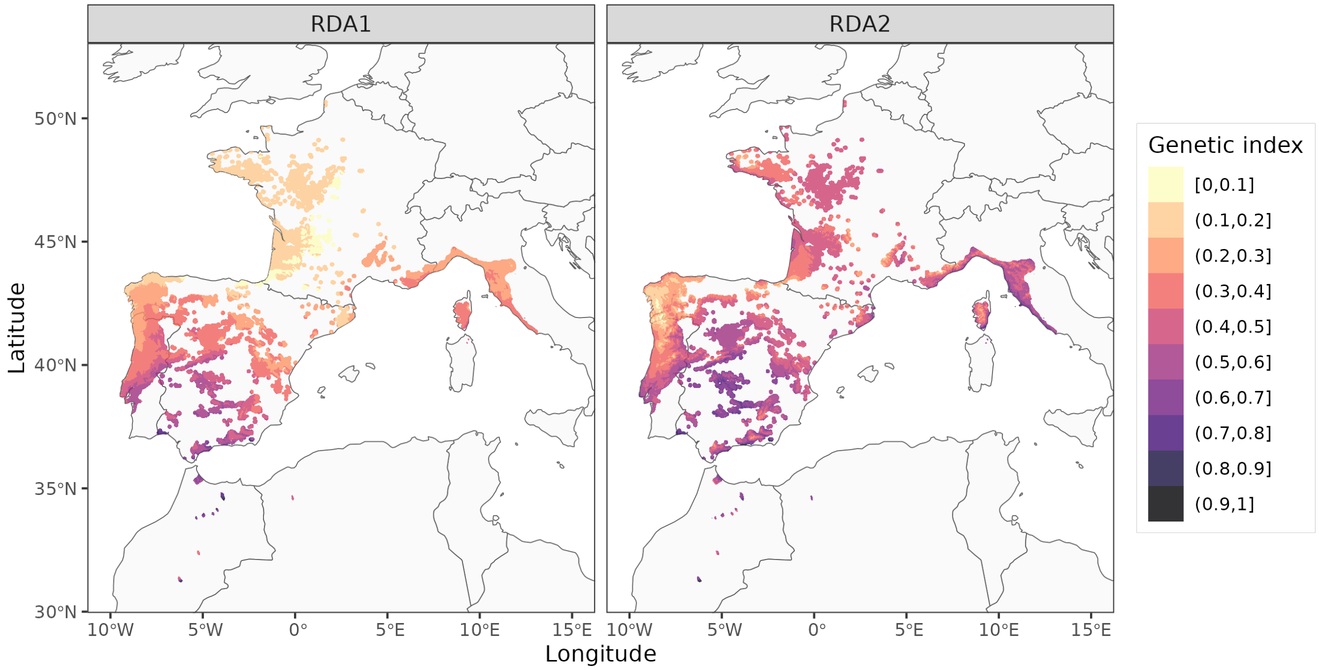

The RDA-based genetic indexes were estimated for both present and future climates across the maritime pine range, and for both control and candidate SNPs. The GO was then calculated for each pixel of the maritime pine range and corresponds to the difference between the genetic indexes calculated for present and future climates.

Figure 13: GO predictions with RDA at the location of the studied populations.

(a) Candidate SNPs.

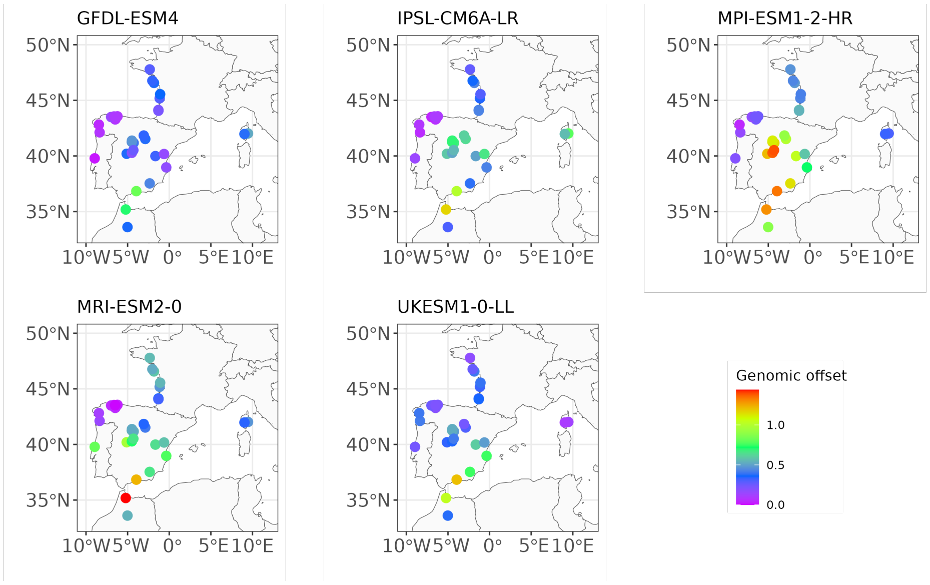

(b) Control SNPs.

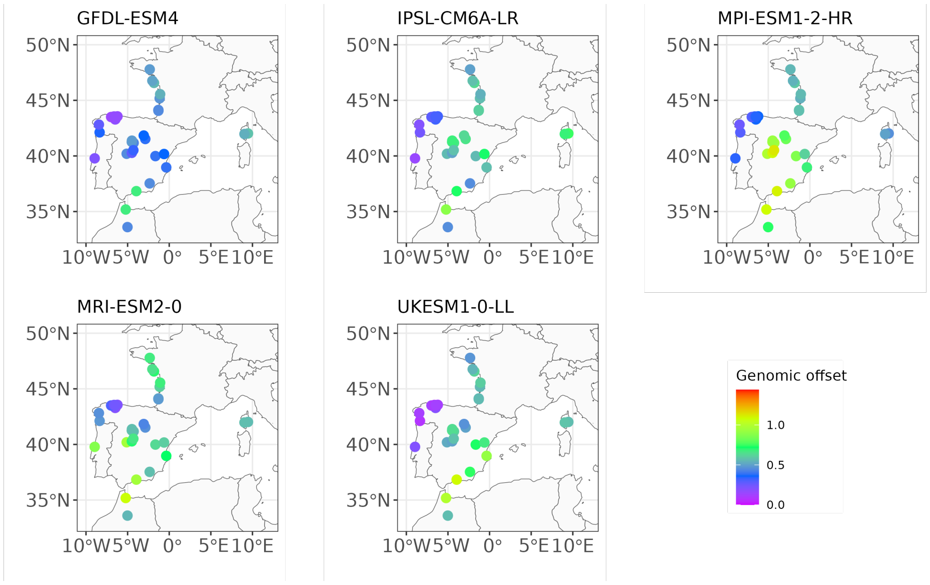

Figure 14: GO predictions with RDA projected across the maritime pine range using the period 1901-1950 as reference period (i.e. past climates under which the populations evolved) and the predictions of five GCMs (one per panel) under the period 2041-2060 for the future climates (SSP3.7-0).

(a) Candidate SNPs.

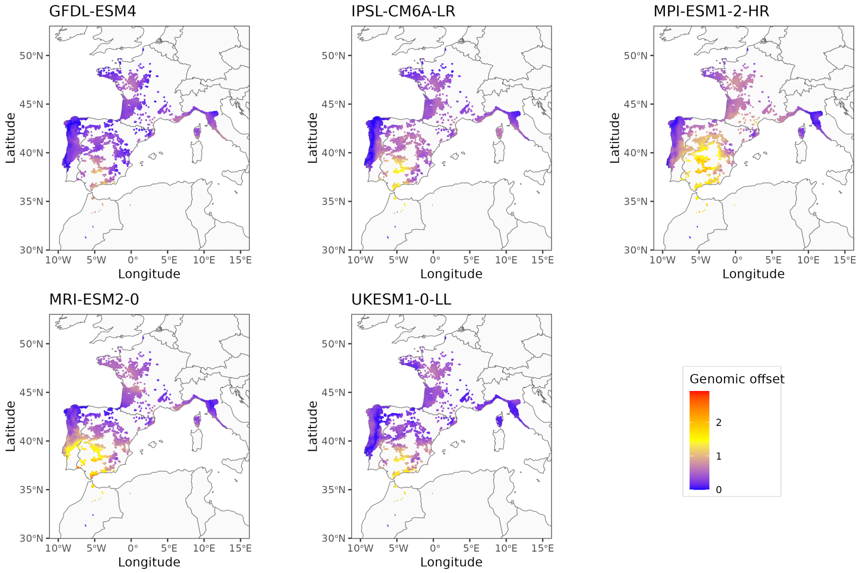

(b) Control SNPs.

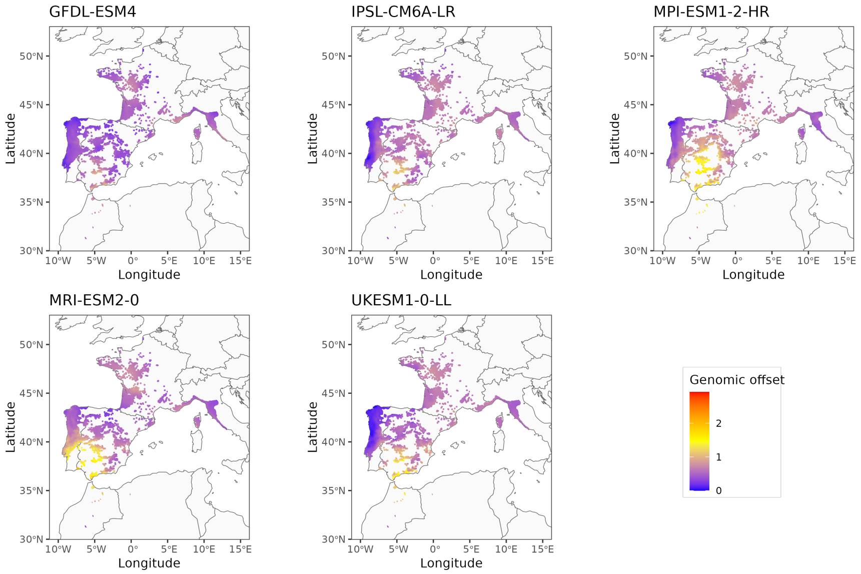

#### 7.2 Gradient Forest (GF)

GO predictions with GF are mostly based on the Github repository associated with Fitzpatrick et al. (2021).

##### 7.2.1 Evaluation of the GF models

We generate a set of plots to evaluate the GF models:

- **Predictor overall importance plots**. These plots show the mean accuracy importance of the climatic variables (left panel) and their mean importance weighted by SNPs $\mathcal{R}^2$ (right panel).
- **Splits density plots**. These plots show the binned split importance and location on each gradient (spikes), kernel density of splits (black lines), of observations (red lines) and of splits standardised by observations density (blue lines). Each distribution integrates to the importance of the climatic variable. These plots capture the rate of change in allele frequency along each climatic gradient, i.e. where important frequency changes of multiple alleles are occurring along the gradient.
- **Allele cumulative plots**. These plots show, for each SNPs, the cumulative importance distributions of splits improvement scaled by $\mathcal{R}^2$ weighted importance, and standardised by density of observations. These turnover functions are expected to show the cumulative change in allele frequency for each SNP along each climatic gradient and therefore to identify where the allele frequency changes are the most important along the gradient (Fitzpatrick et al. (2015)).
- **Predictor cumulative plots**. These plots show, for each climatic variable, the cumulative importance distributions of splits improvement scaled by $\mathcal{R}^2$ weighted importance, and standardised by density of observations, averaged over all SNPs. These turnover functions are expected to show the cumulative change in overall allelic composition along a given climatic gradient. The maximum height of each curve should indicate the total amount of turnover in allele frequencies associated with the climatic variable considered, and by extension, the relative importance of that variable in explaining changes in allele frequency, holding all other variables constant (Fitzpatrick & Keller (2015)). The shape of each turnover function should capture how the rate of change in allele frequencies varies along the climatic gradient, and could therefore be useful to identify and where the most important changes in genetic composition occur on the gradient (Fitzpatrick & Keller (2015)). However, this interpretation was questioned in Láruson et al. (2022), who found with simulations that for linear allele frequency clines, the steepness of the turnover function does not reflect the rate of allele frequency change.
- $\mathcal{R}^2$ **measure of the fit** of the random forest model for each SNPs.

The code to generate those plots comes from 'Example analysis of biodiversity survey data with R package `gradientForest`' by C. Roland Pitcher, Nick Ellis and Stephen J. Smith (pdf available here).

Figure 15: Predictor overall importance plots.

(a) Candidate SNPs.

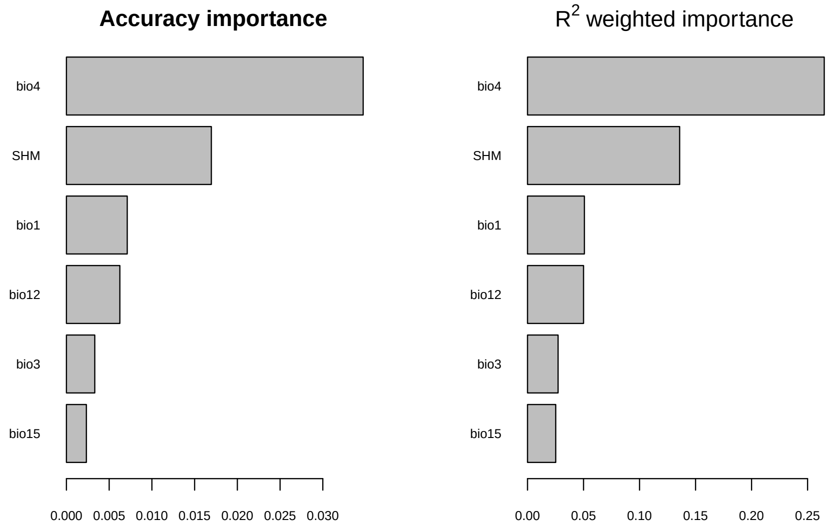

(b) Control SNPs.

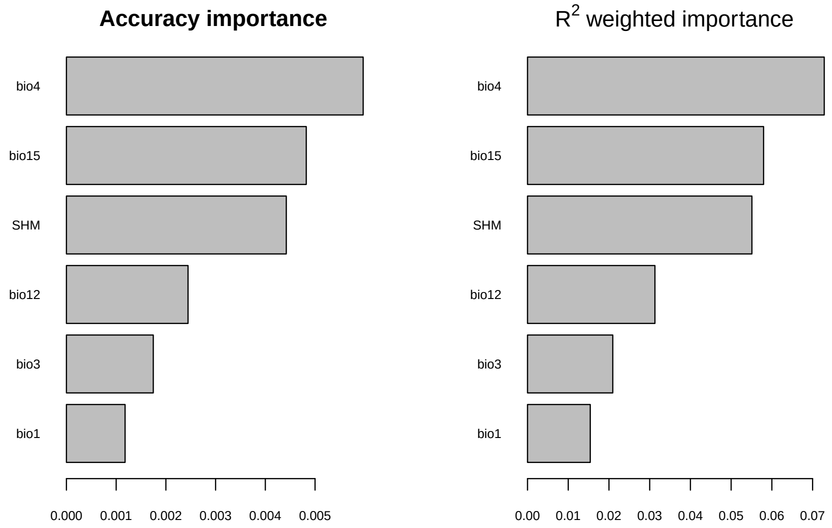

Figure 16: Splits density plots.

(a) Candidate SNPs.

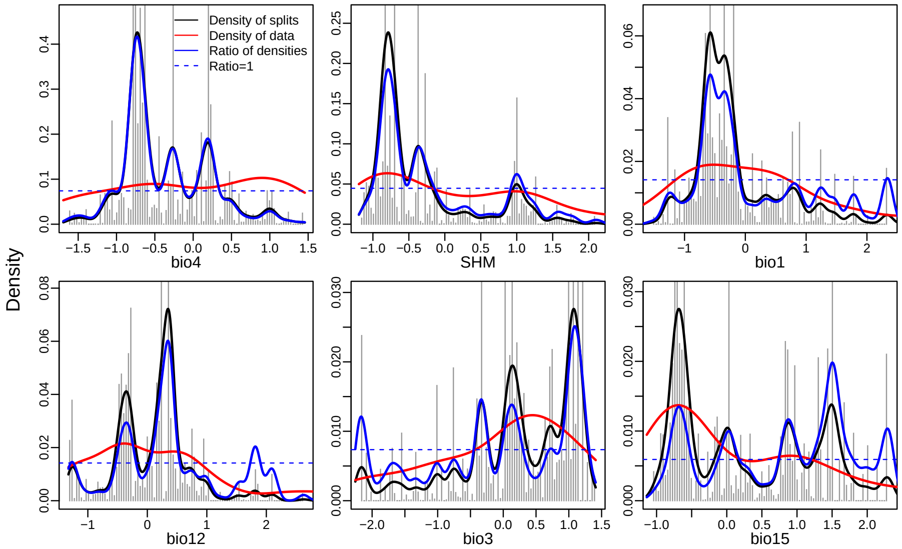

(b) Control SNPs.

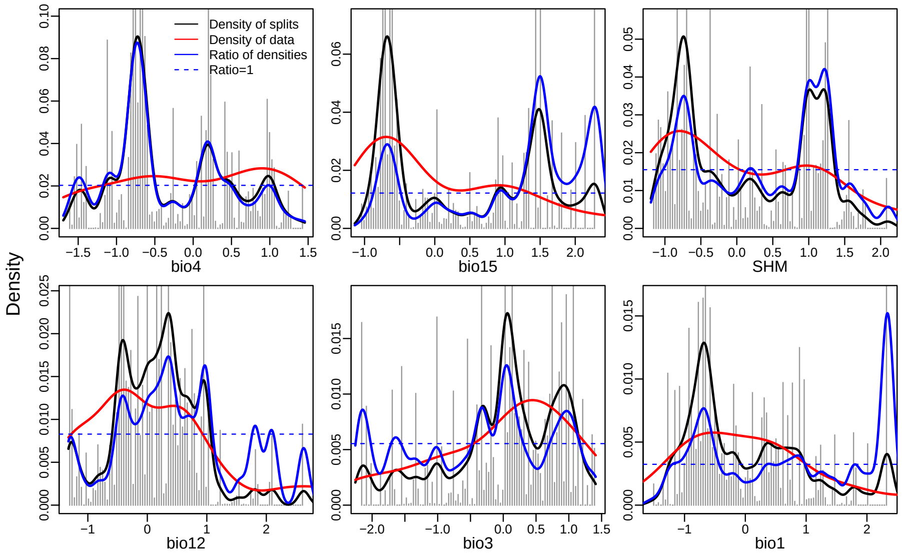

Figure 17: Allele cumulative plots for the candidate SNPs.

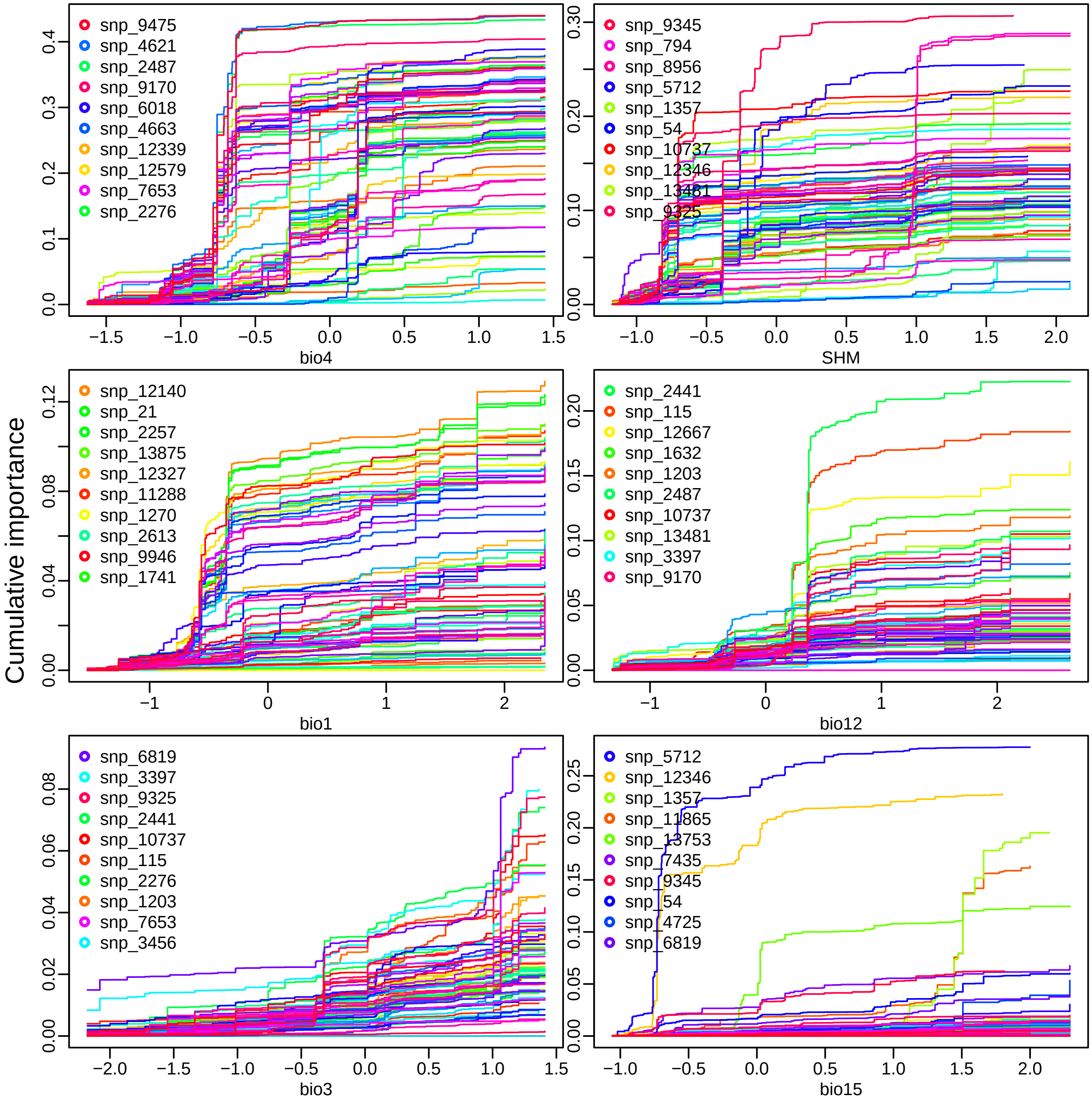

Figure 18: Allele cumulative plots for the control SNPs.

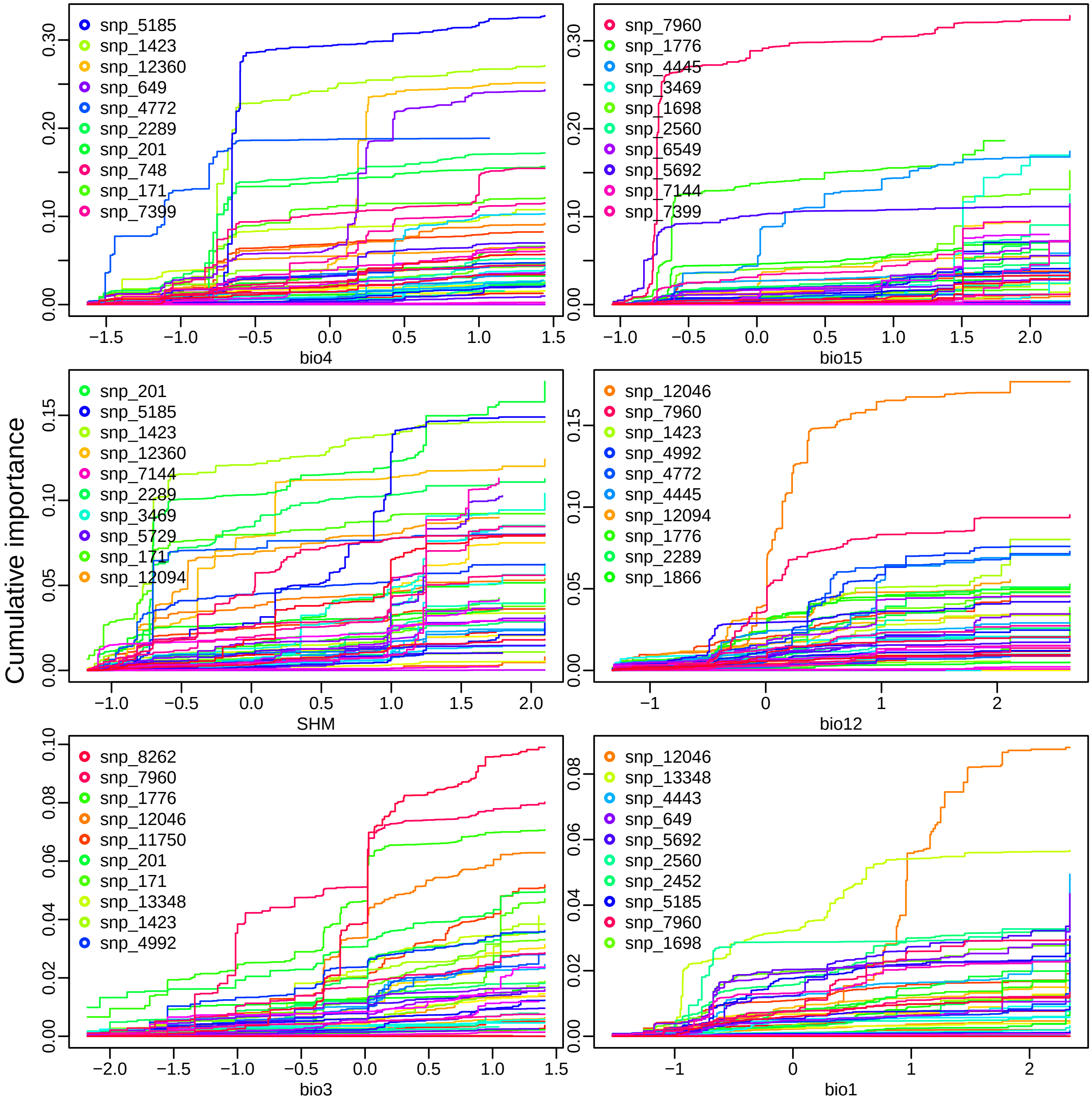

Figure 19: Predictor cumulative plots for candidate SNPs.

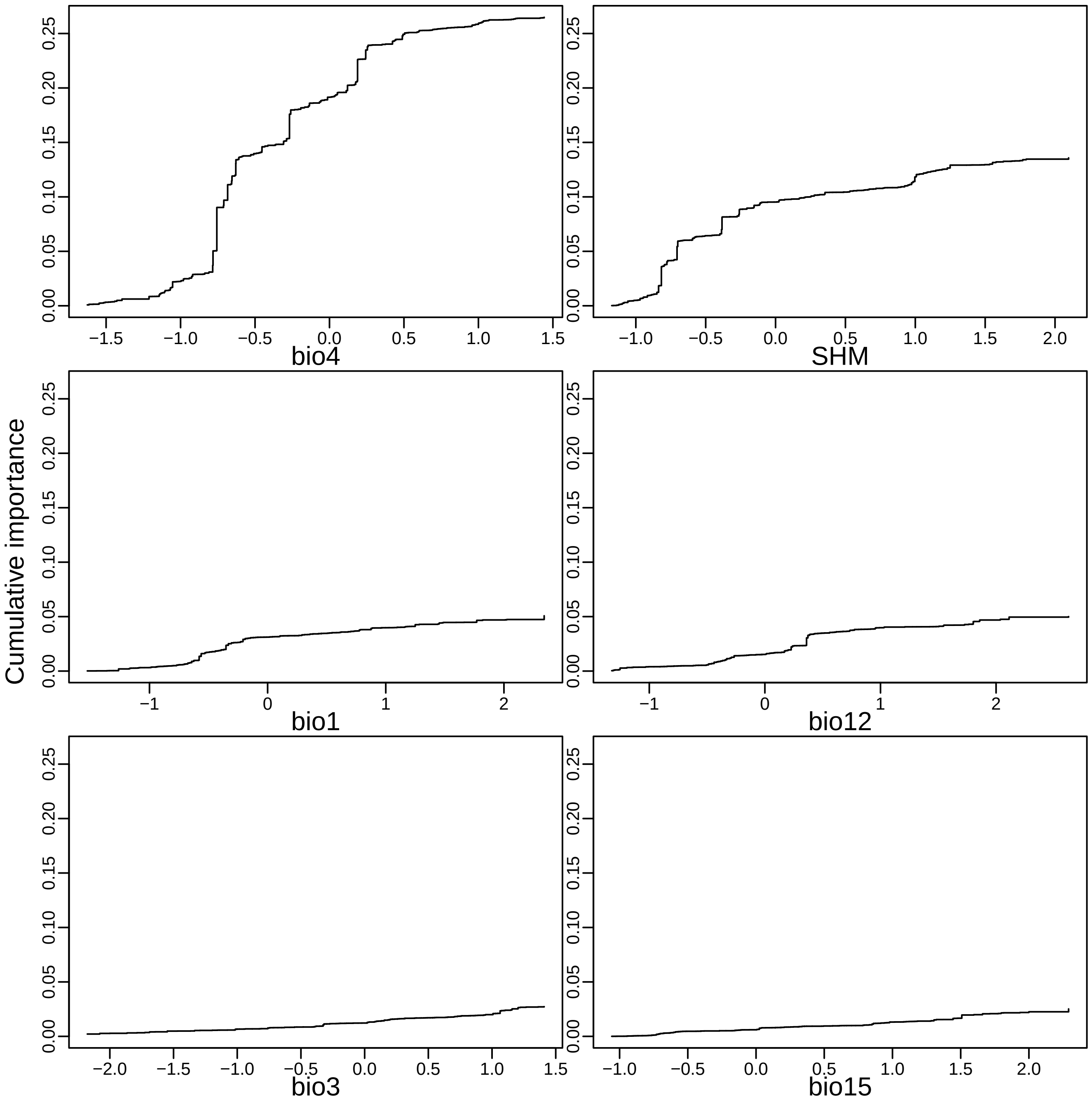

Figure 20: Predictor cumulative plots for control SNPs.

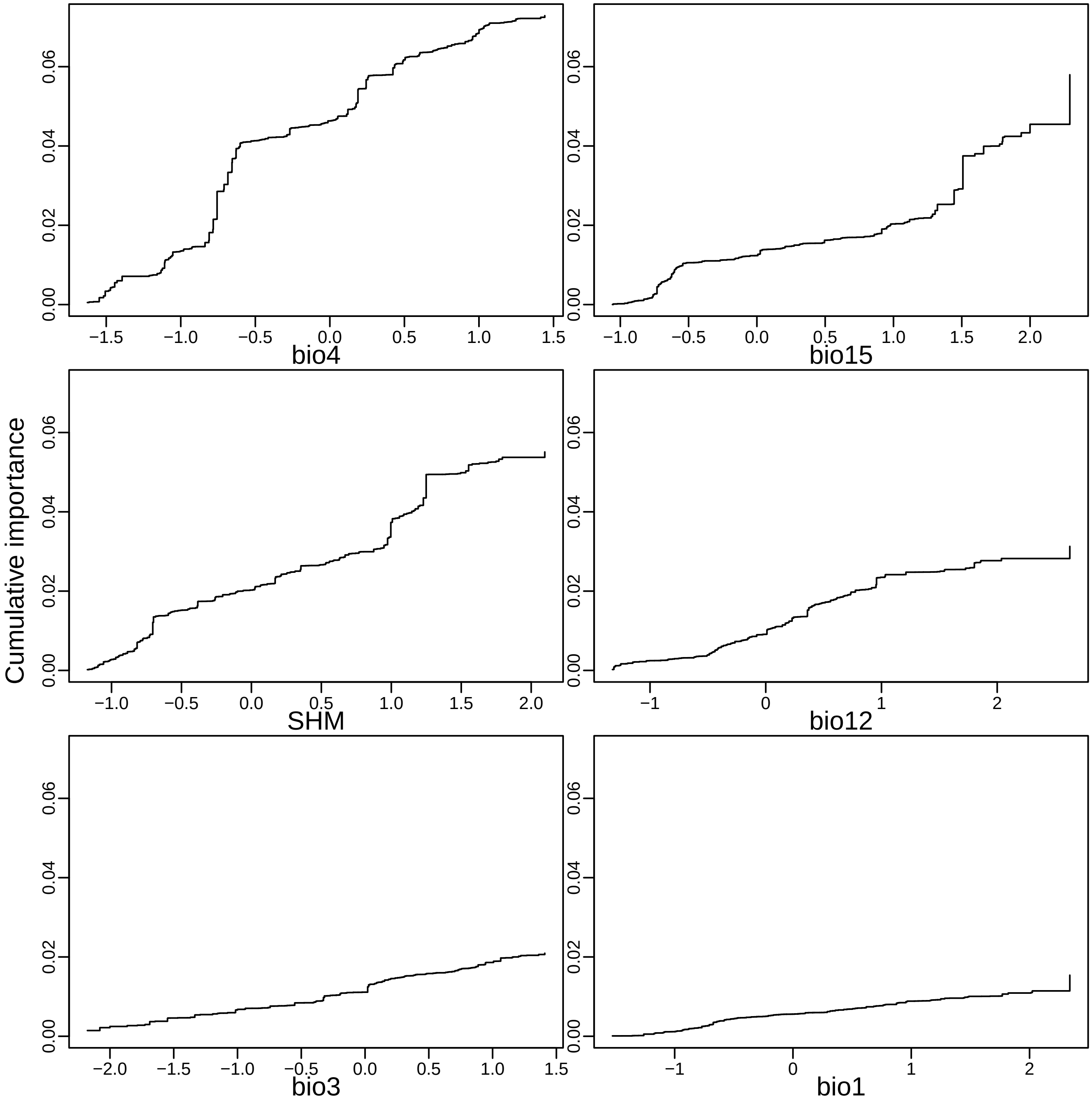

Figure 21: $\mathscr{R}^2$ measure of the fit of the random forest model for each SNPs.

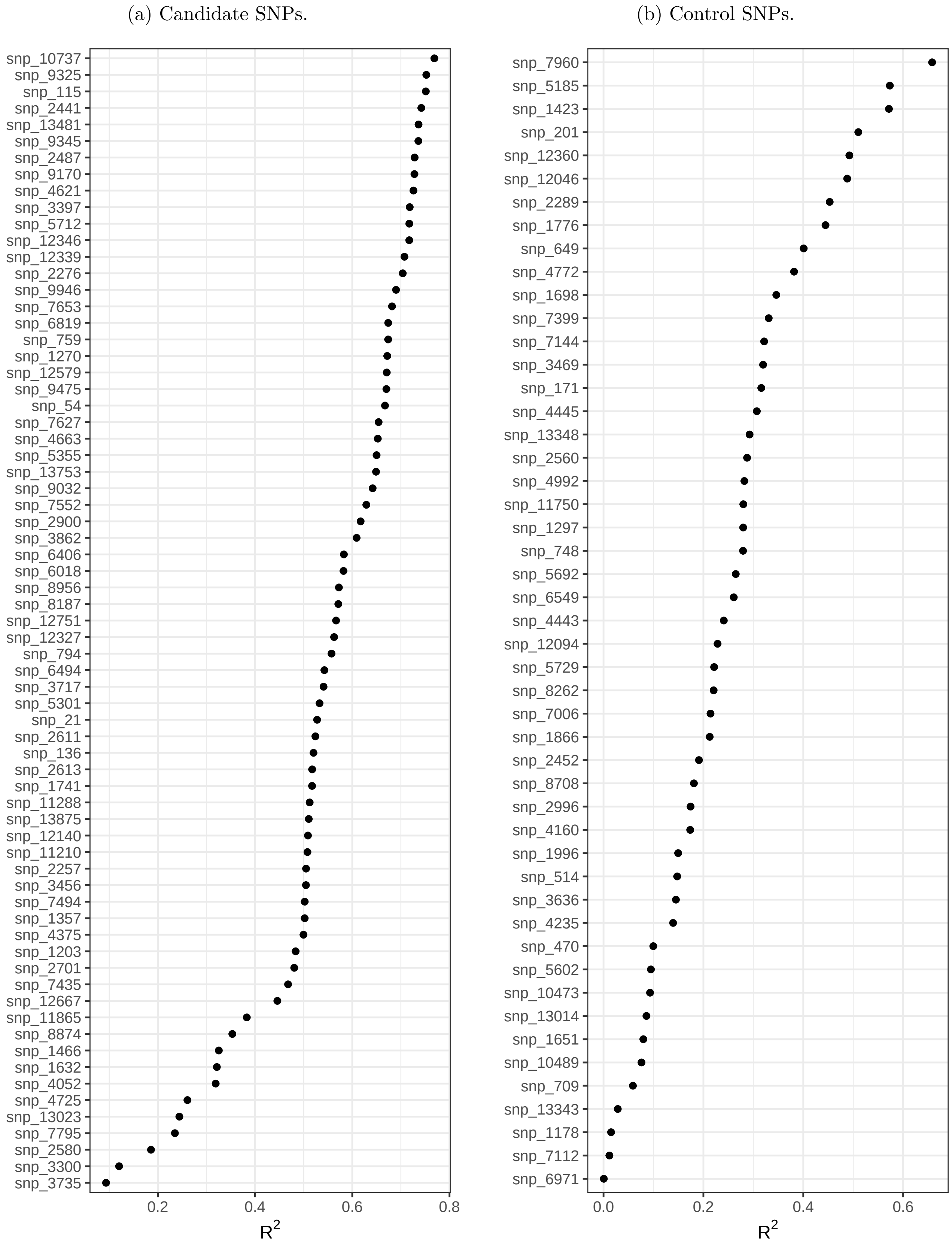

##### 7.2.2 GO predictions

Figure 22: GO predictions with GF at the location of the studied populations.

(a) Candidate SNPs.

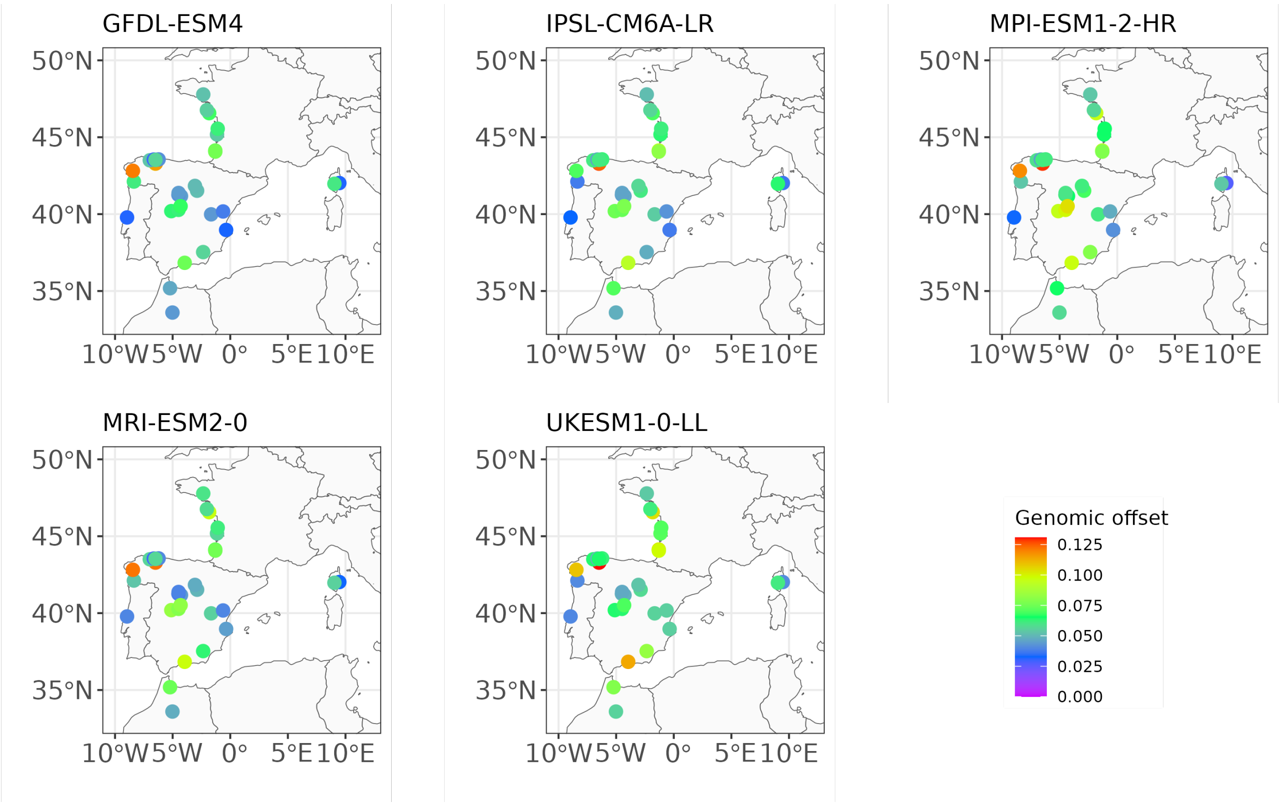

(b) Control SNPs.

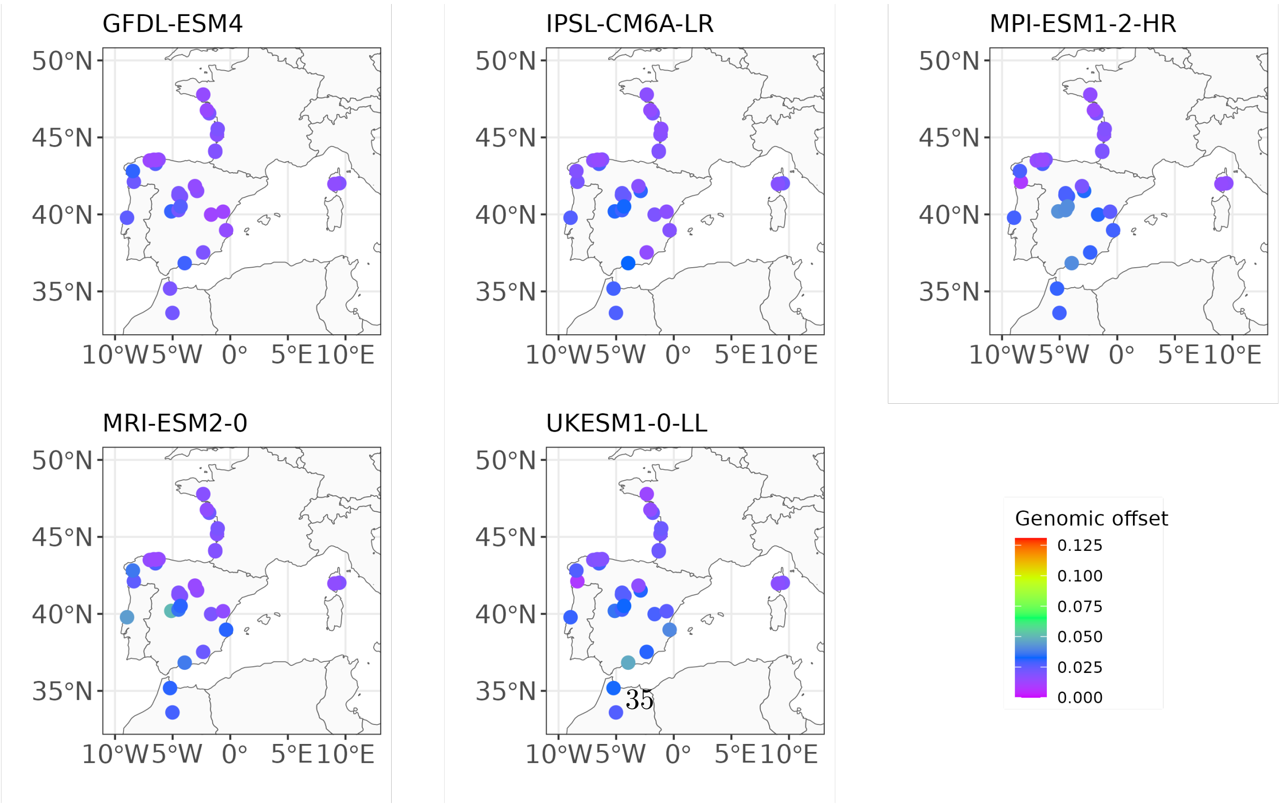

#### 7.3 Latent factor mixed models (LFMM)

Figure 23: GO predictions with LFMM and all SNPs.

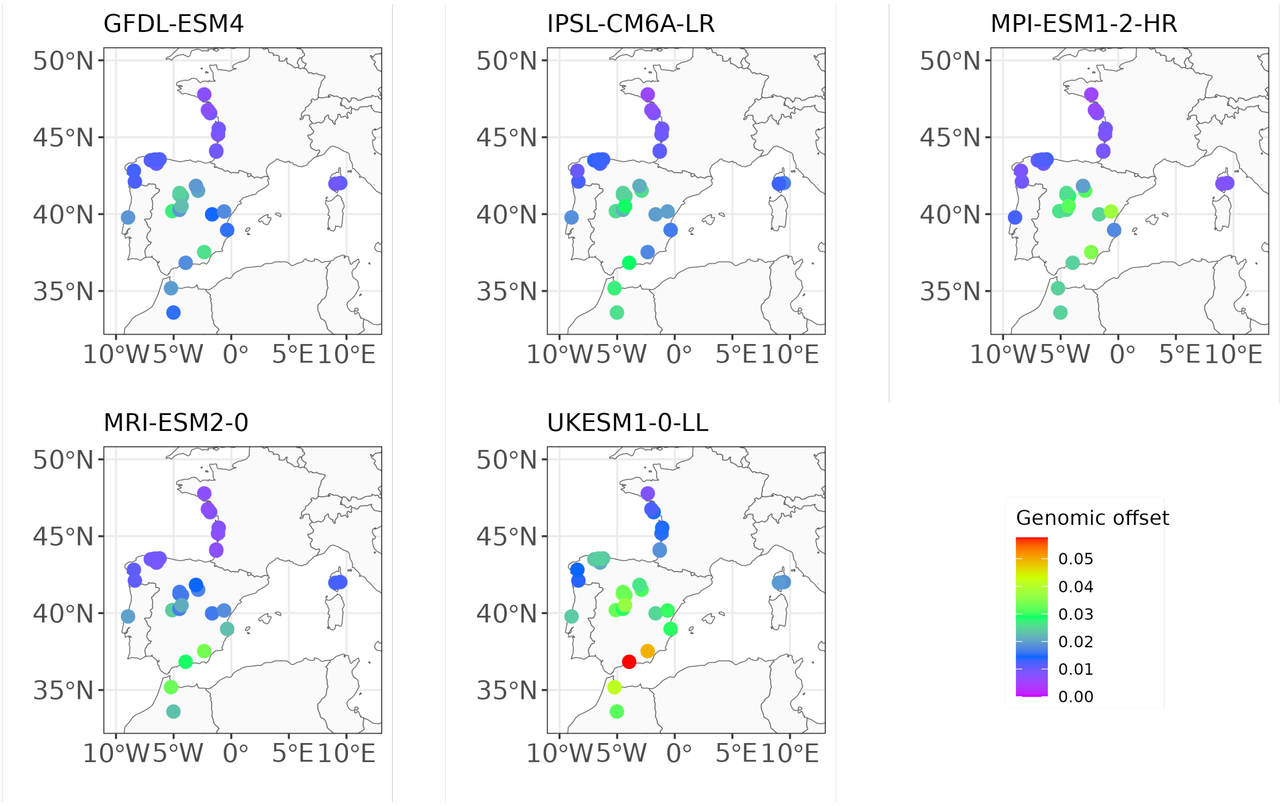

Figure 24: GO predictions with LFMM at the location of the studied populations.

(a) Candidate SNPs.

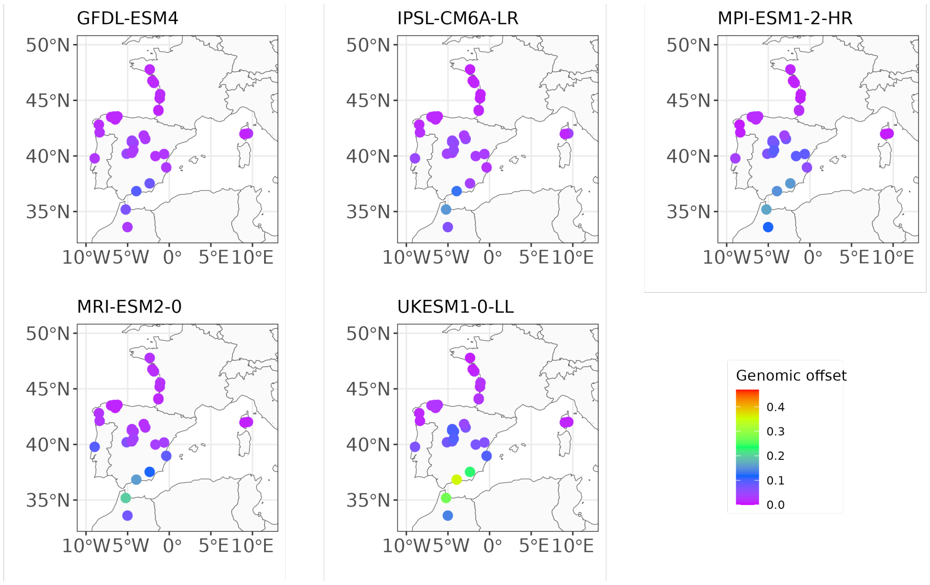

(b) Control SNPs.

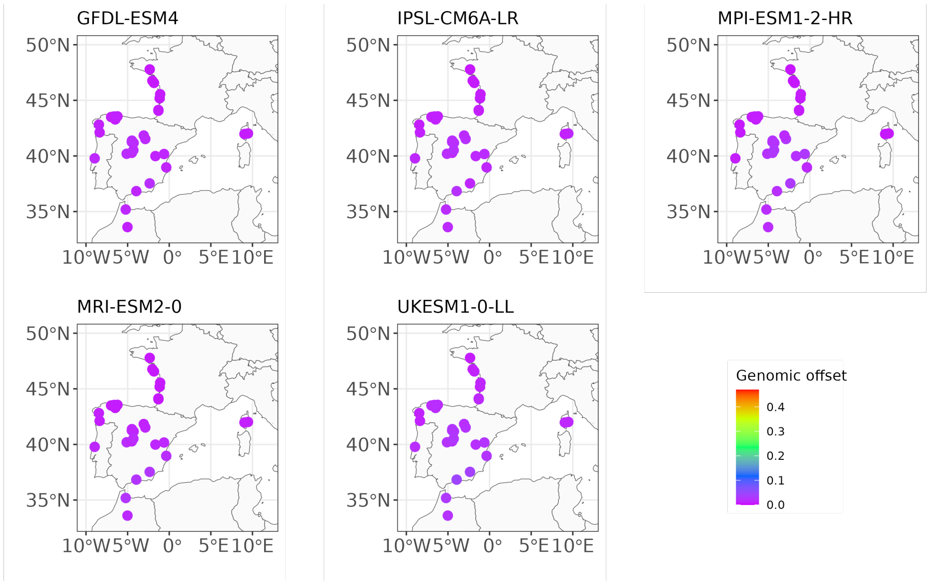

#### 7.4 Generalized Dissimilarity Modelling (GDM)

GDM analyses are based on Ferrier et al. (2007), Mokany et al. (2022), Fitzpatrick & Keller (2015), the GDM website and the GDM Github.

Figure 25: Observed genetic distance against predicted climatic distance.

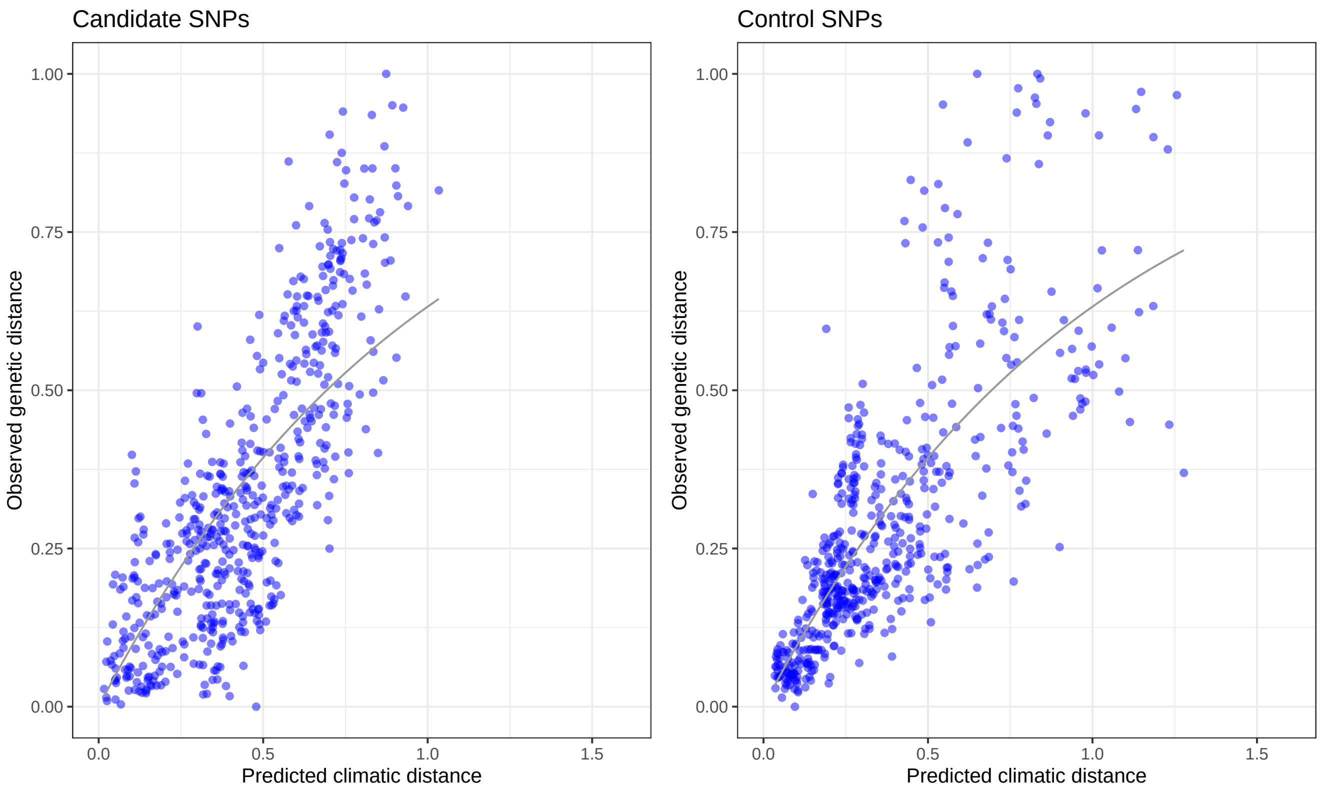

Figure 26: Predicted vs observed genetic distance.

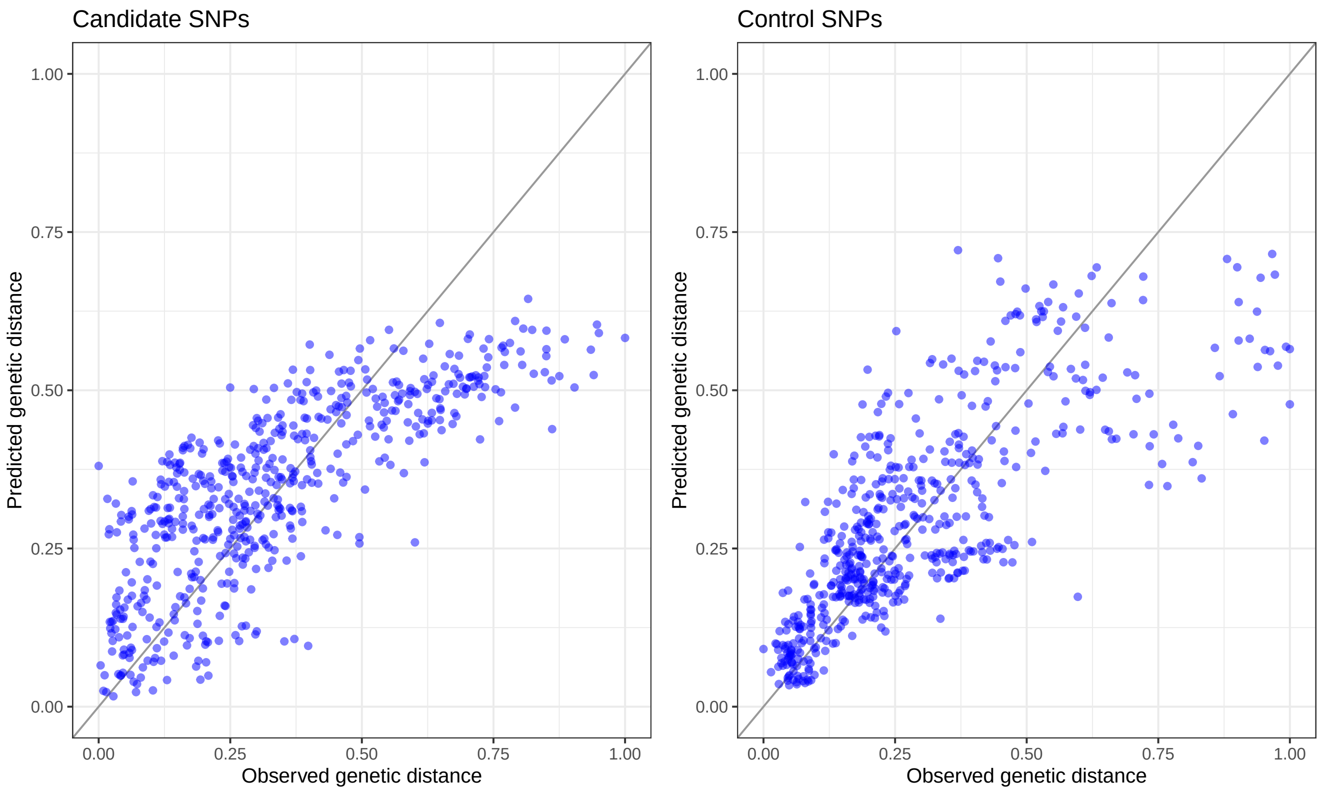

The fitted I-splines in Figure 27 and Figure 28 show how population genetic composition changes along each climatic gradient. They are only showed for climatic variables with non-zero coefficients, i.e. with a relationship with the genetic distance.

Similarly to the interpretation of the GF turnover functions, Fitzpatrick & Keller (2015) suggests that:

- the maximum height of each I-spline should indicate the magnitude of total genetic change along the climatic gradient and thereby should capture the relative importance of the corresponding climatic variable in contributing to allelic turnover while holding all other variables constant (i.e. a partial climatic distance).
- the spline's shape is expected to capture how the rate of genetic change varies with position along the climatic gradient, and thus should provide insight into where along the climatic gradient the genetic changes are the most pronounced.

Figure 27: I-spline plots of the GDM models based on the candidate SNPs.

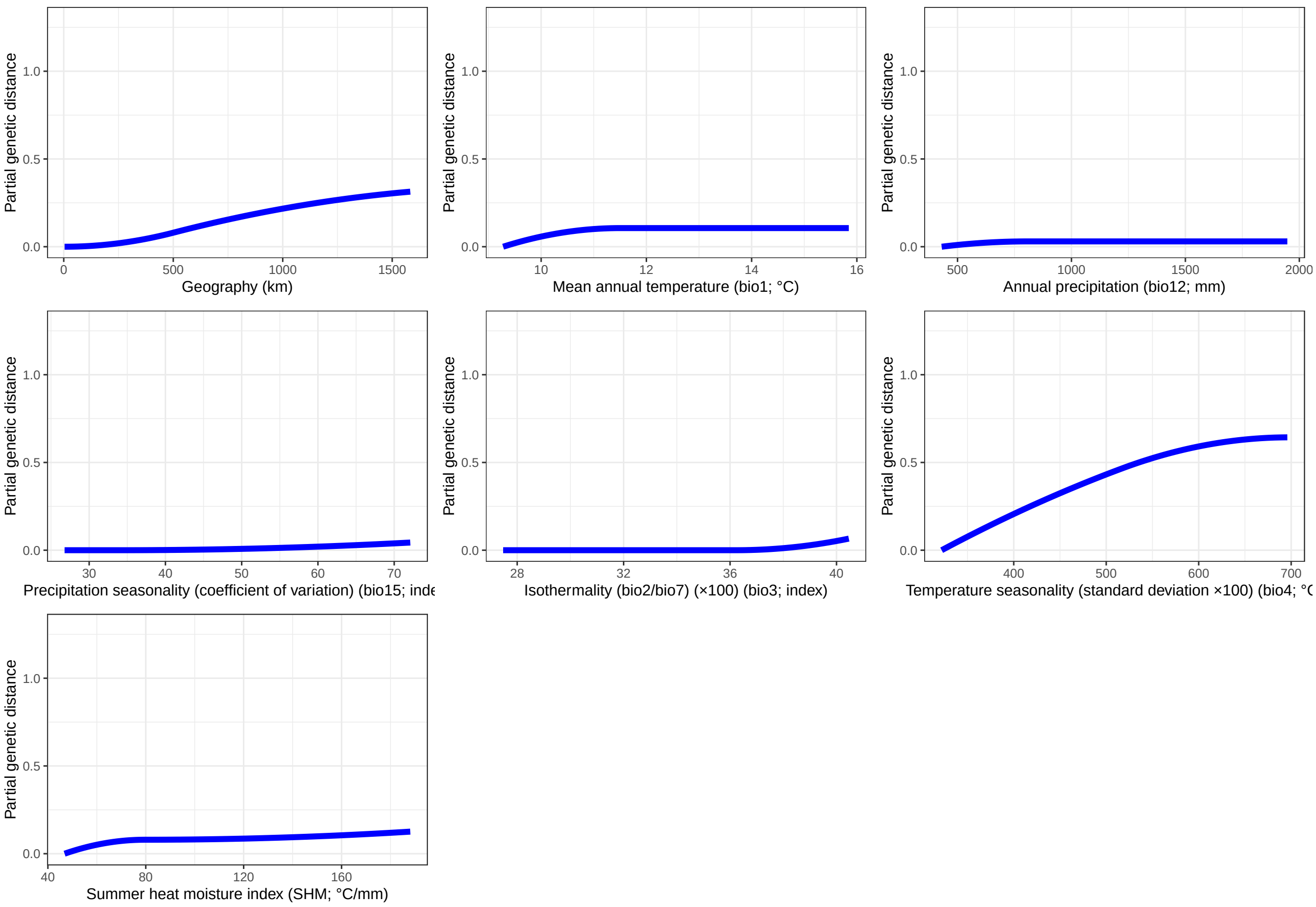

Figure 28: I-spline plots of the GDM models based on the control SNPs.

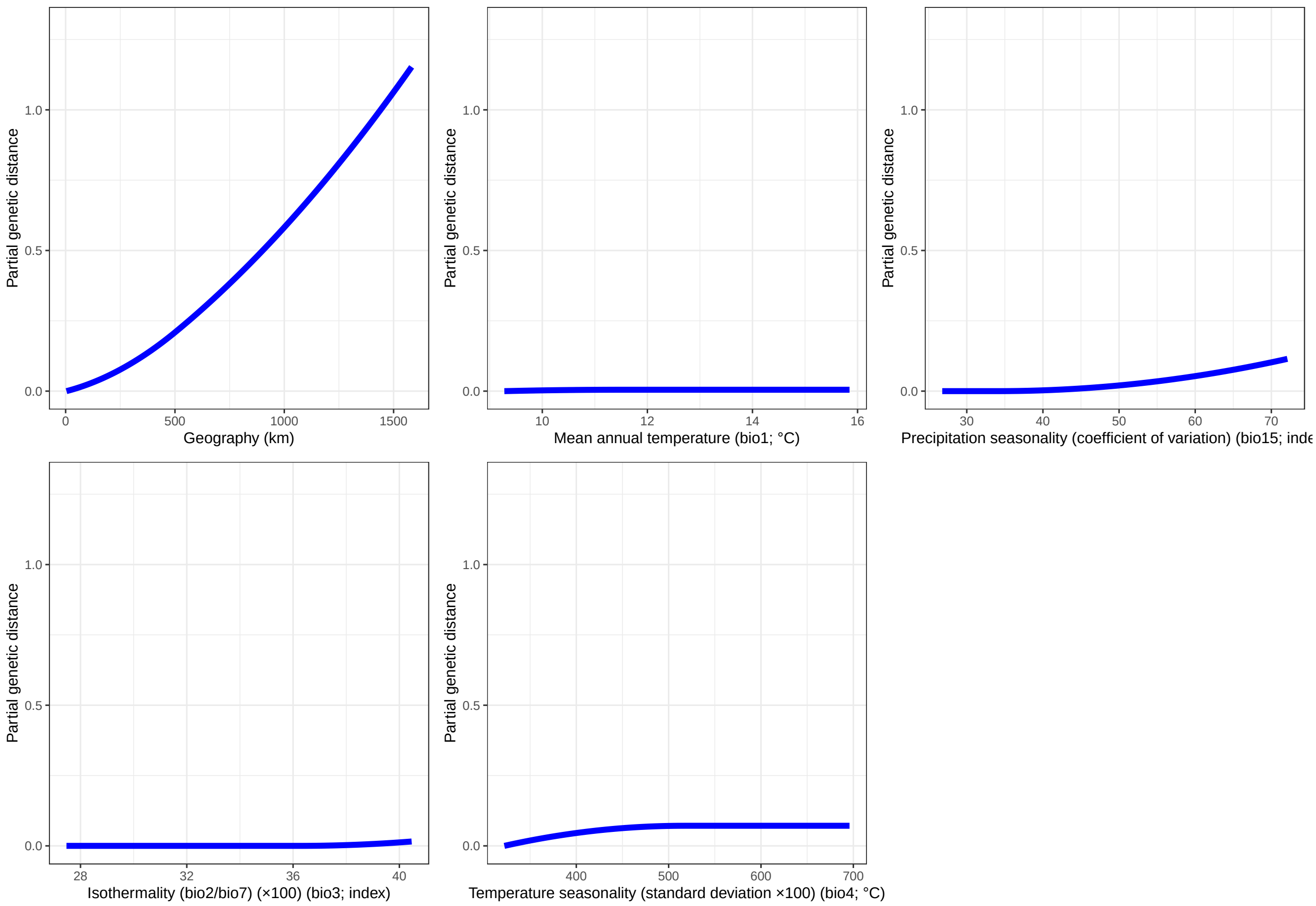

Figure 29: GO predictions with GDM at the location of the studied populations.

(a) Candidate SNPs.

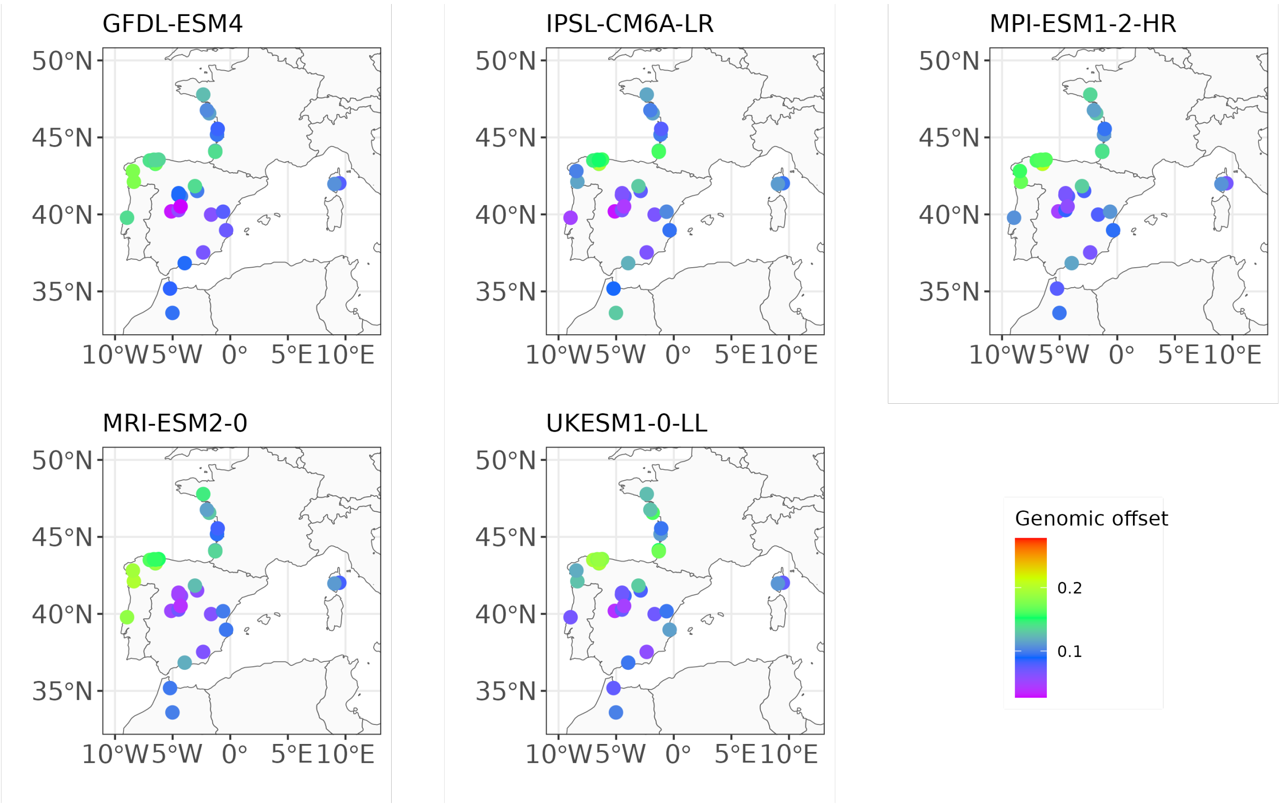

(b) Control SNPs.

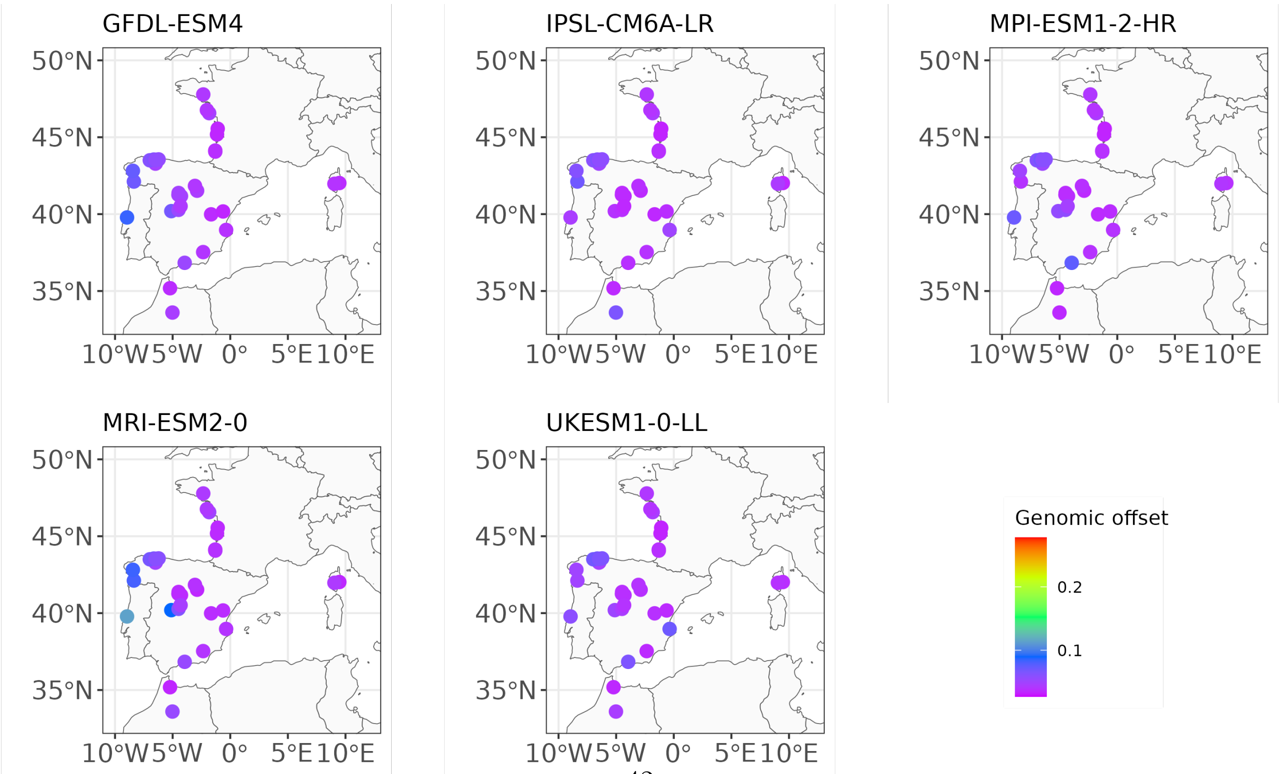

#### 7.5 Relationship with Euclidean distance

In this section, we estimate the relationship between GO predictions and the Euclidean climatic distance $D_p$, calculated for each population $p$ as follows:

$$D_p = \sqrt{\sum_{i=1}^{N}(X_{ref,i,p} - X_{fut,i,p})^2}$$

with $N$ the number of selected climatic variables (in our case, five), $X_{ref,i,p}$ the mean value of the climatic variable $i$ across the reference period 1901-1950 for the population $p$ and $X_{fut,i,p}$ the predicted mean value of the climatic variable $i$ across the future period 2041-2070 for the population $p$.

Figure 30: Linear association between GO predictions with RDA and the control SNPs and the Euclidean climatic distances. Populations are colored based on their main gene pool.

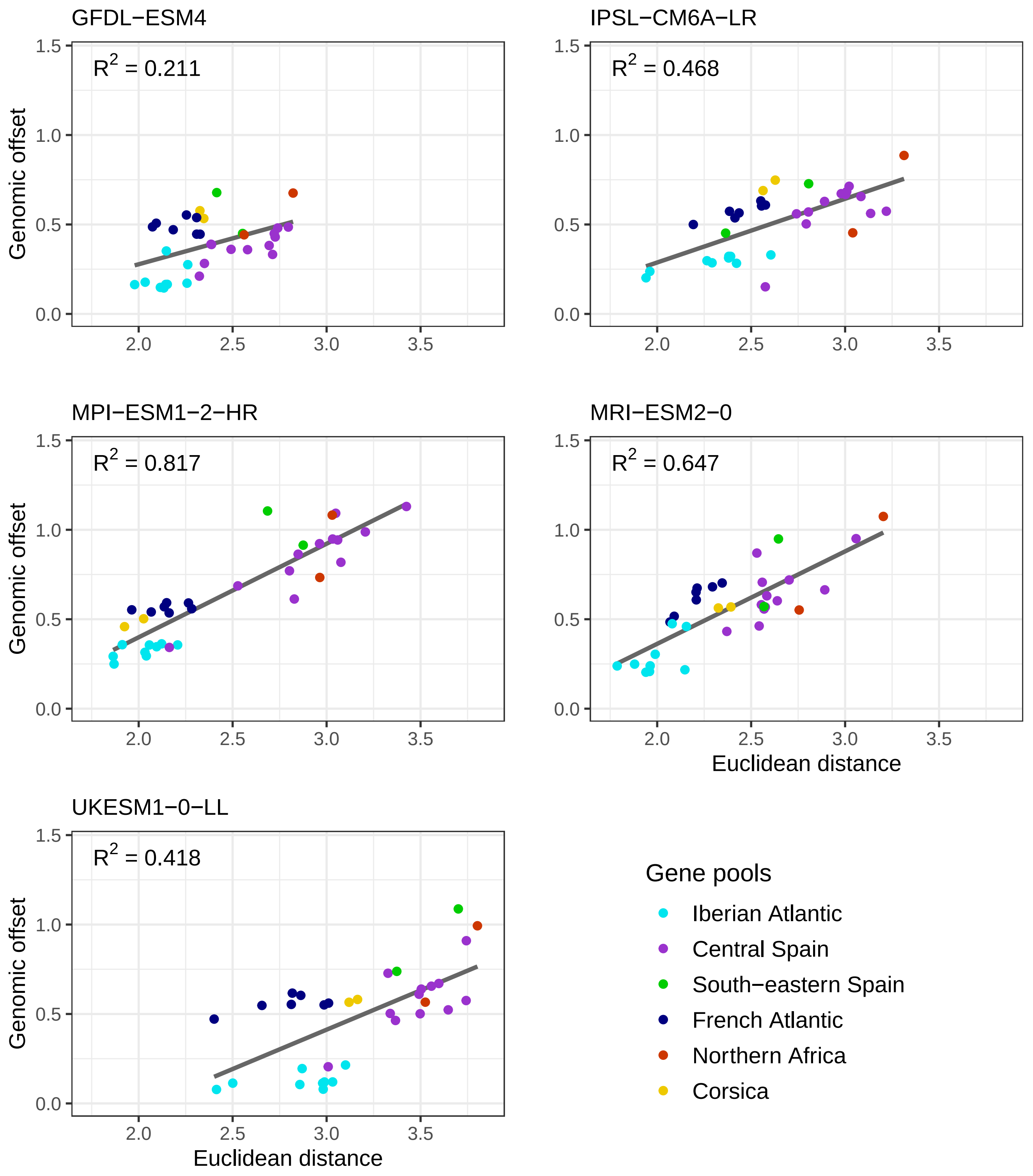

Figure 31: Linear association between GO predictions with RDA and the candidate SNPs and the Euclidean climatic distances. Populations are colored based on their main gene pool.

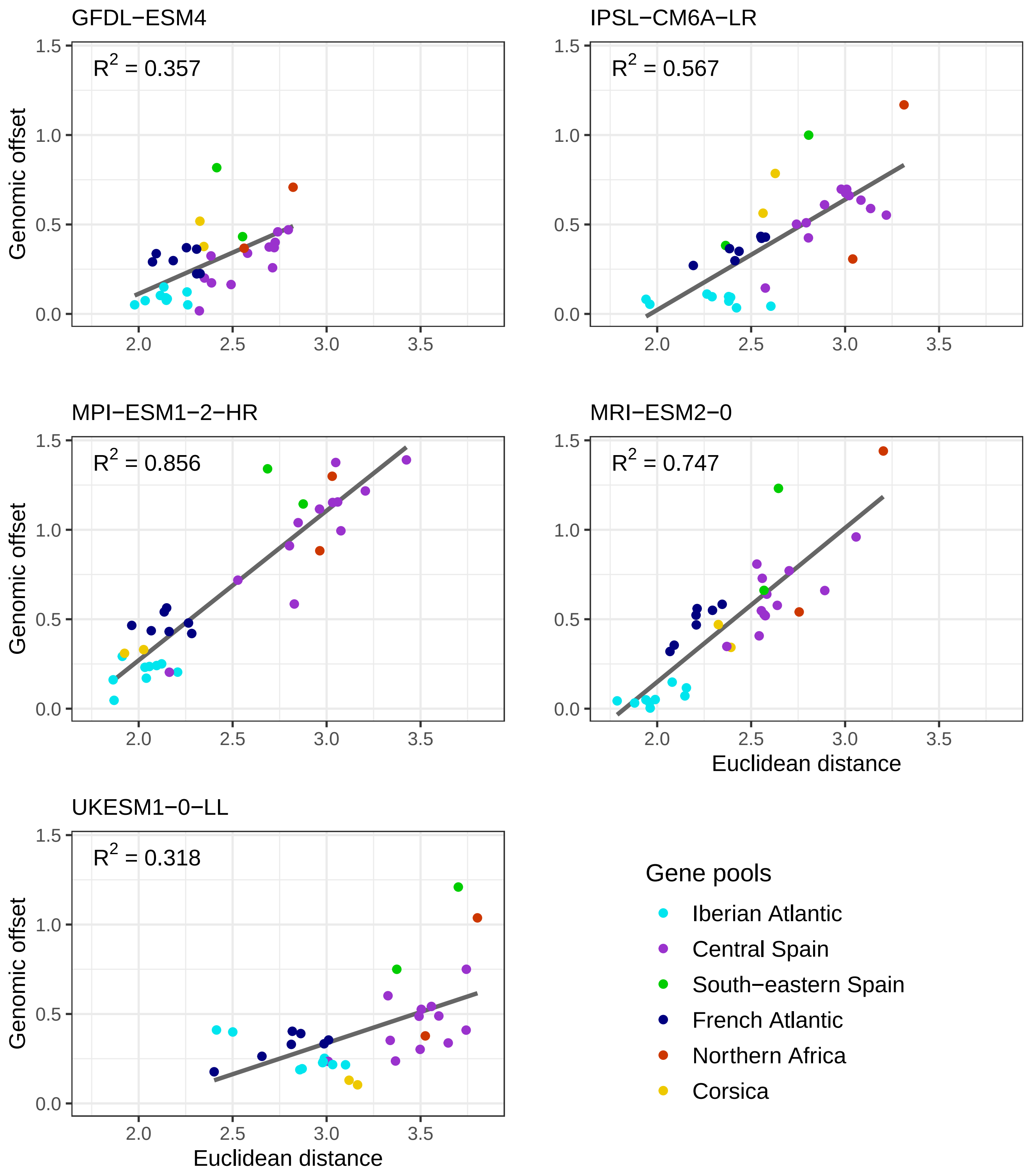

Figure 32: Linear association between GO predictions with GF and the control SNPs and the Euclidean climatic distances. Populations are colored based on their main gene pool.

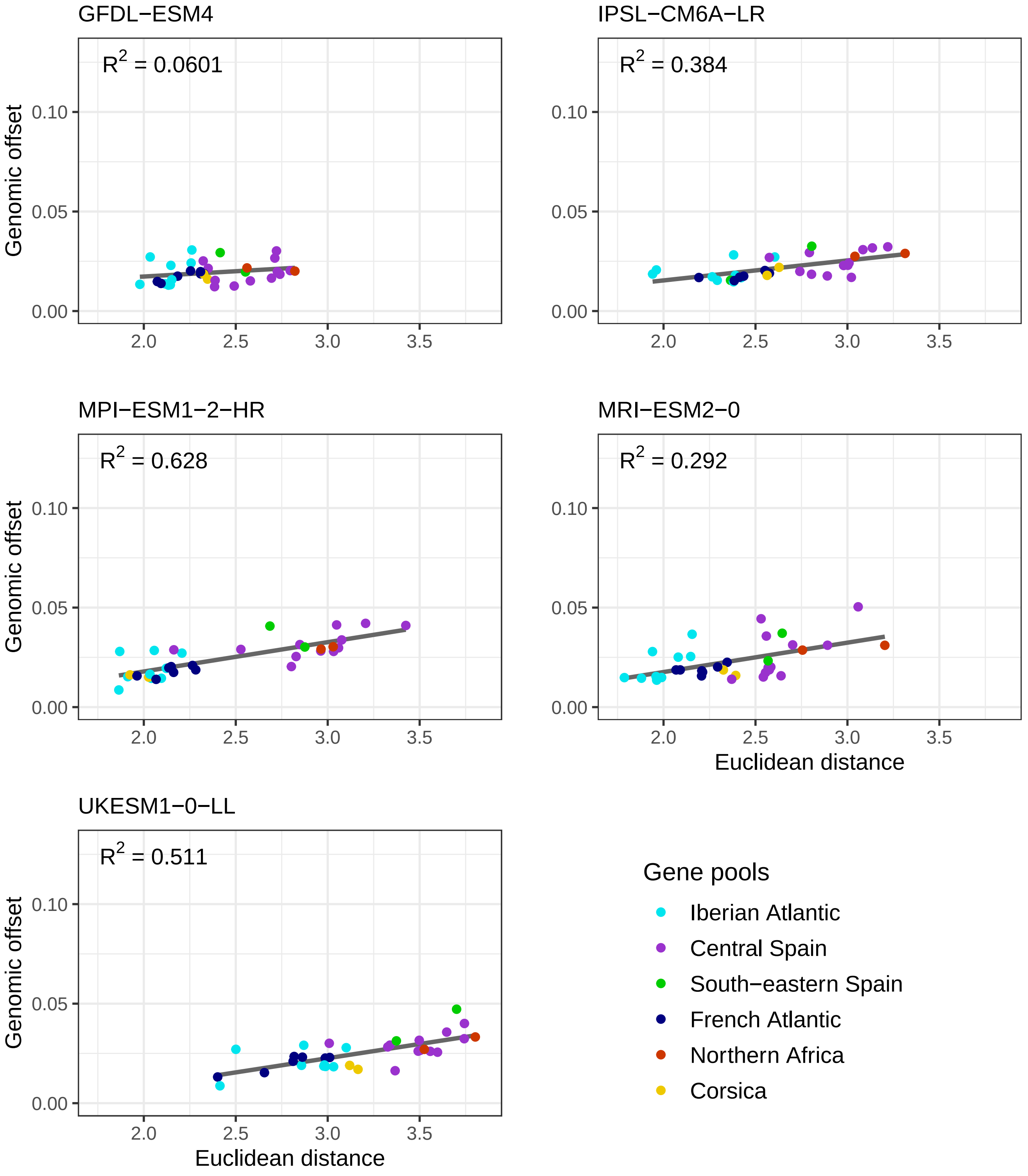

Figure 33: Linear association between GO predictions with GF and the candidate SNPs and the Euclidean climatic distances. Populations are colored based on their main gene pool.

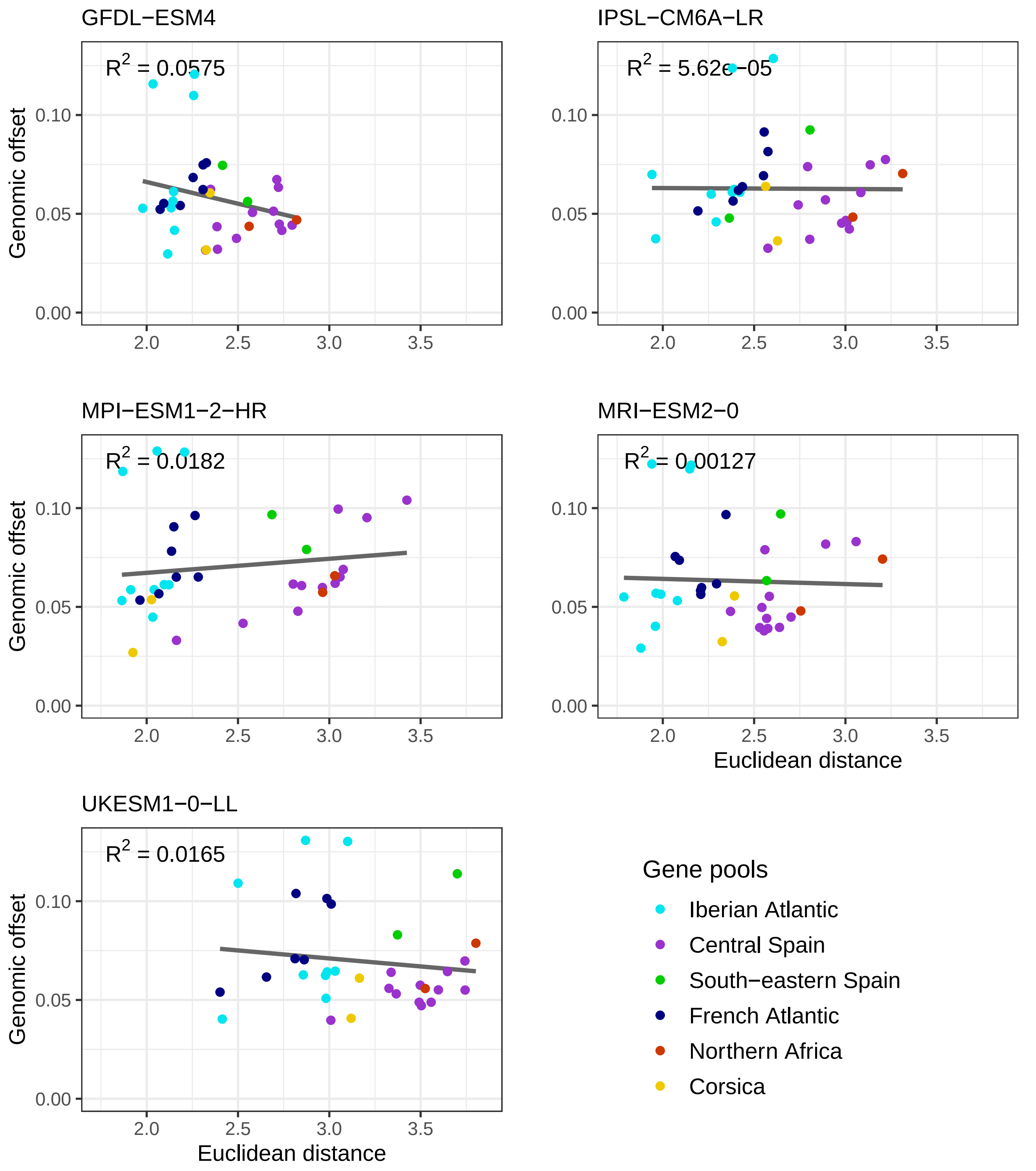

Figure 34: Linear association between GO predictions with LFMM and all SNPs and the Euclidean climatic distances. Populations are colored based on their main gene pool.

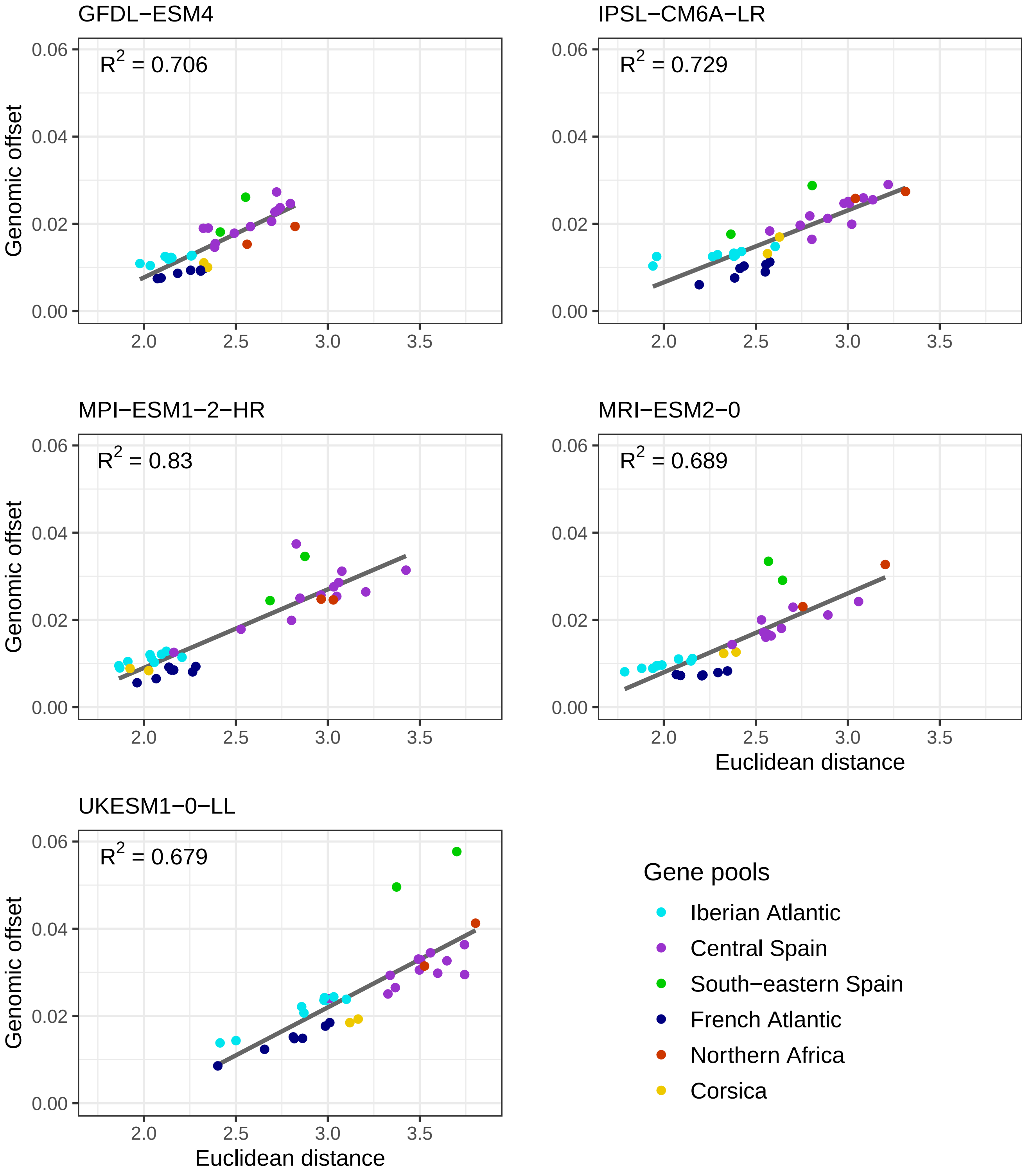

Figure 35: Linear association between GO predictions with LFMM and the control SNPs and the Euclidean climatic distances. Populations are colored based on their main gene pool.

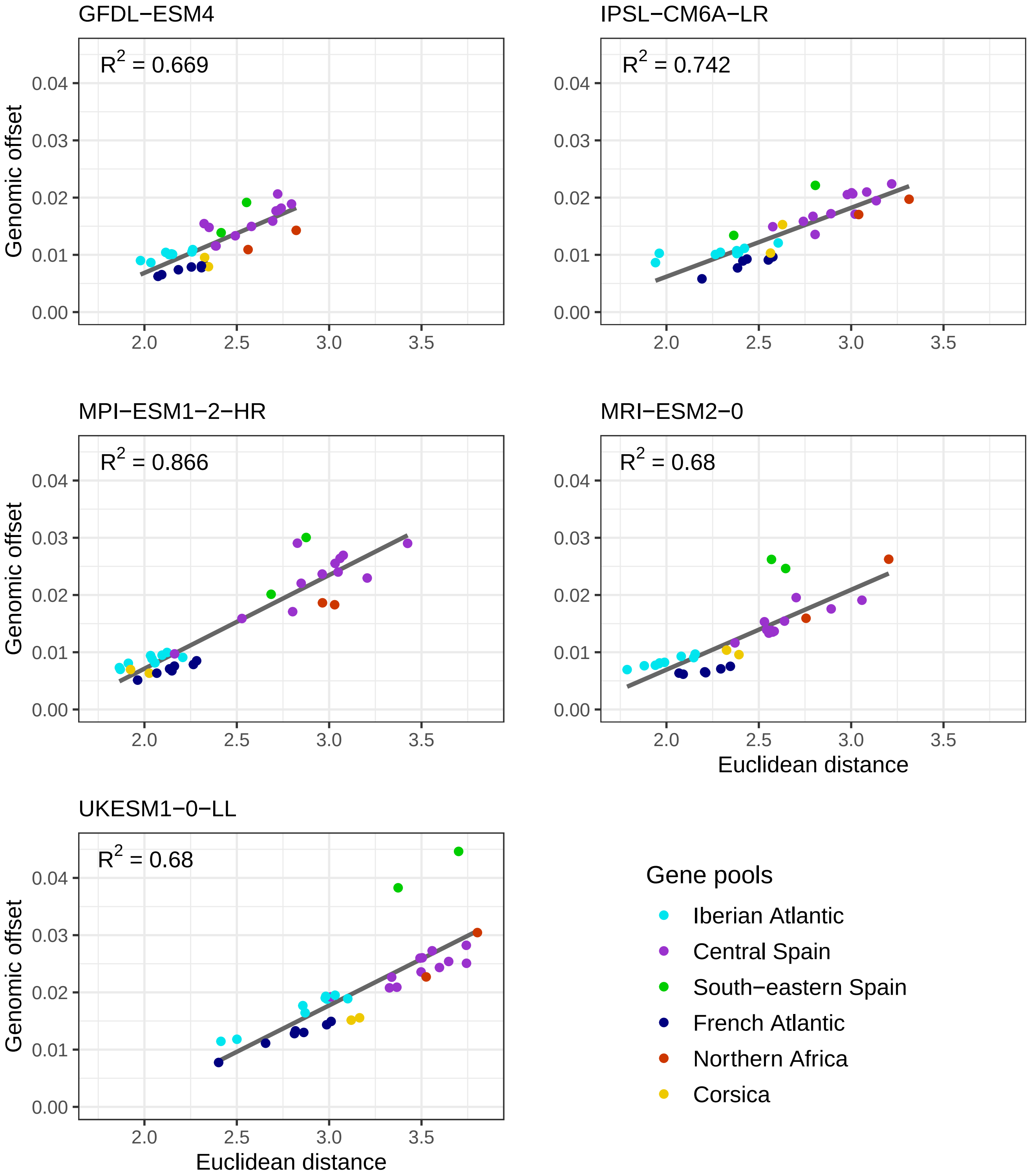

Figure 36: Linear association between GO predictions with LFMM and the candidate SNPs and the Euclidean climatic distances. Populations are colored based on their main gene pool.

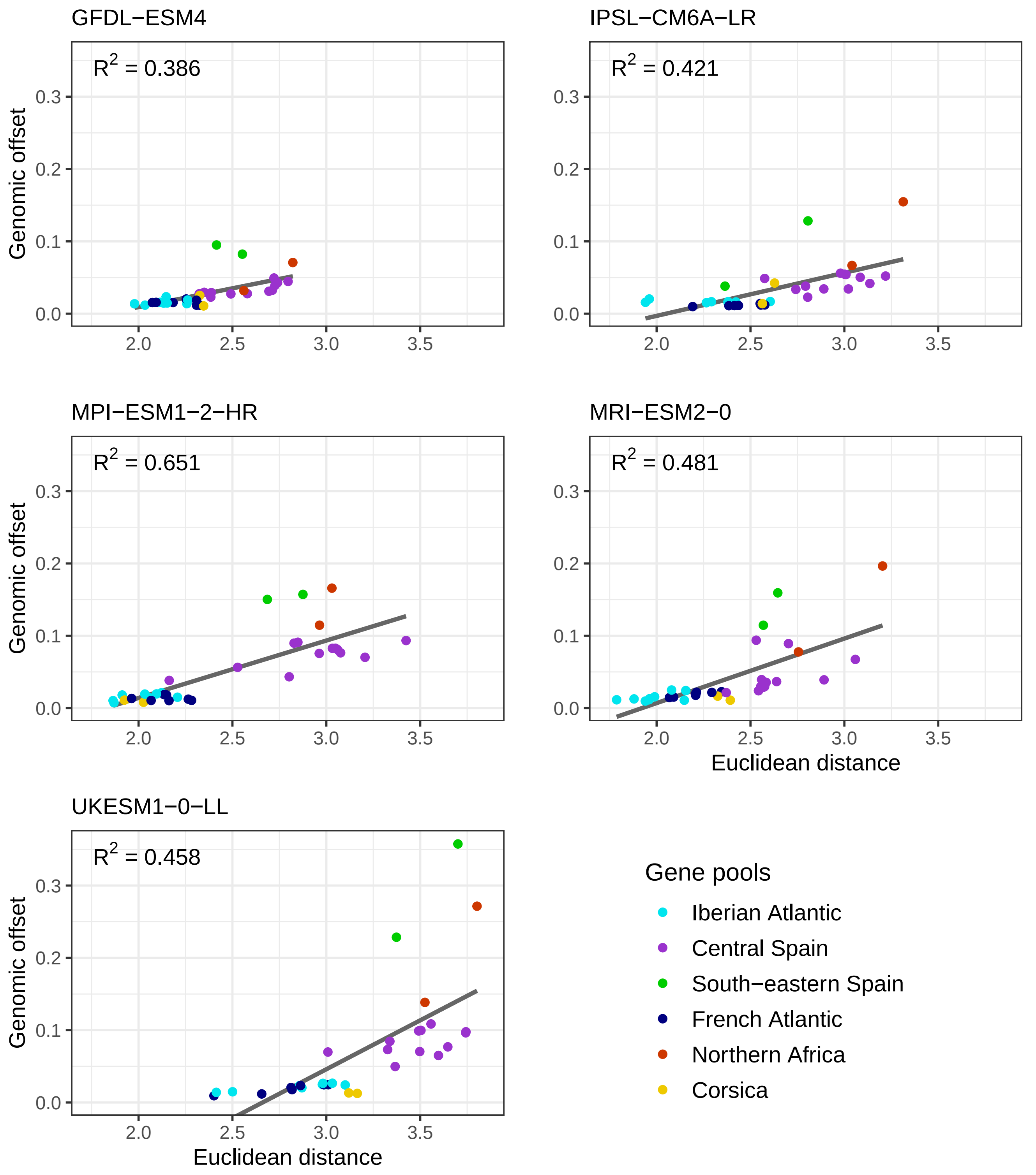

Figure 37: Linear association between GO predictions with GDM and the control SNPs and the Euclidean climatic distances. Populations are colored based on their main gene pool.

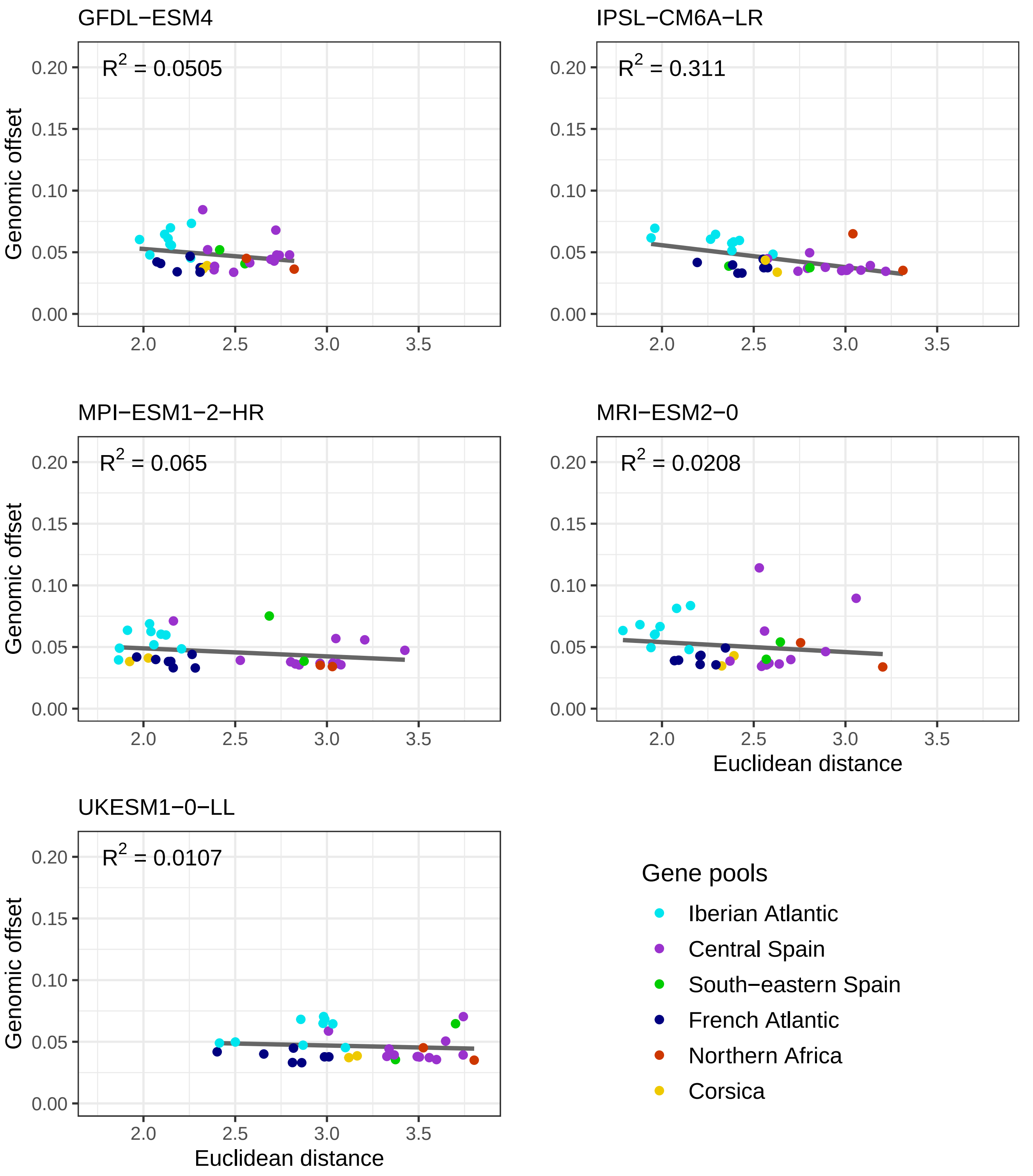

Figure 38: Linear association between GO predictions with GDM and the candidate SNPs and the Euclidean climatic distances. Populations are colored based on their main gene pool.

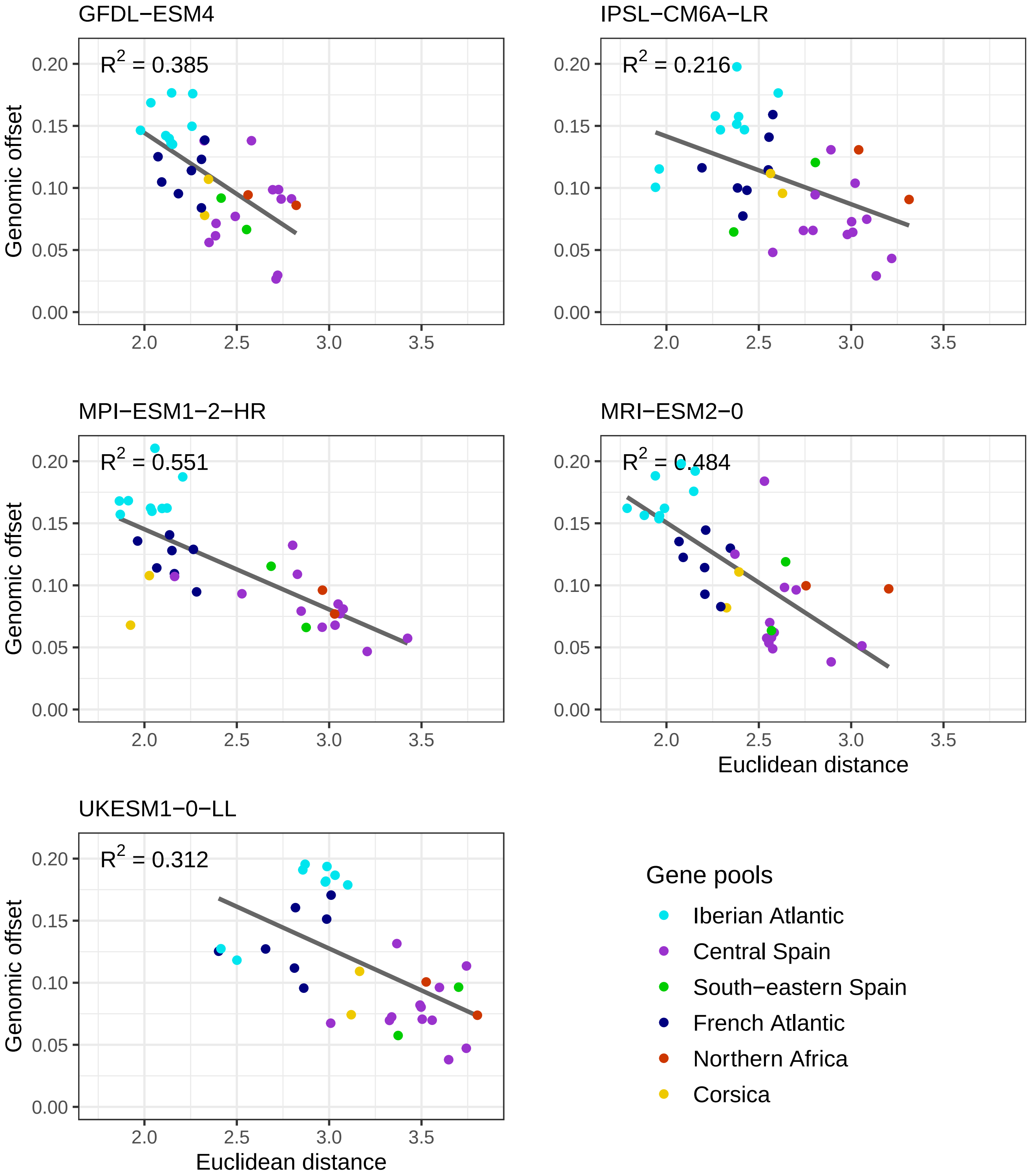

#### 7.6 Variability in GO predictions

##### 7.6.1 Correlations across GO predictions

We calculated Pearson correlations among GO predictions from the four different methods (GF, GDM, RDA and LFMM), three SNP sets (candidates, control and all SNPs for LFMM) and five GCMs.

Figure 39: Correlations among GO predictions from the GFDL−ESM4 GCM.

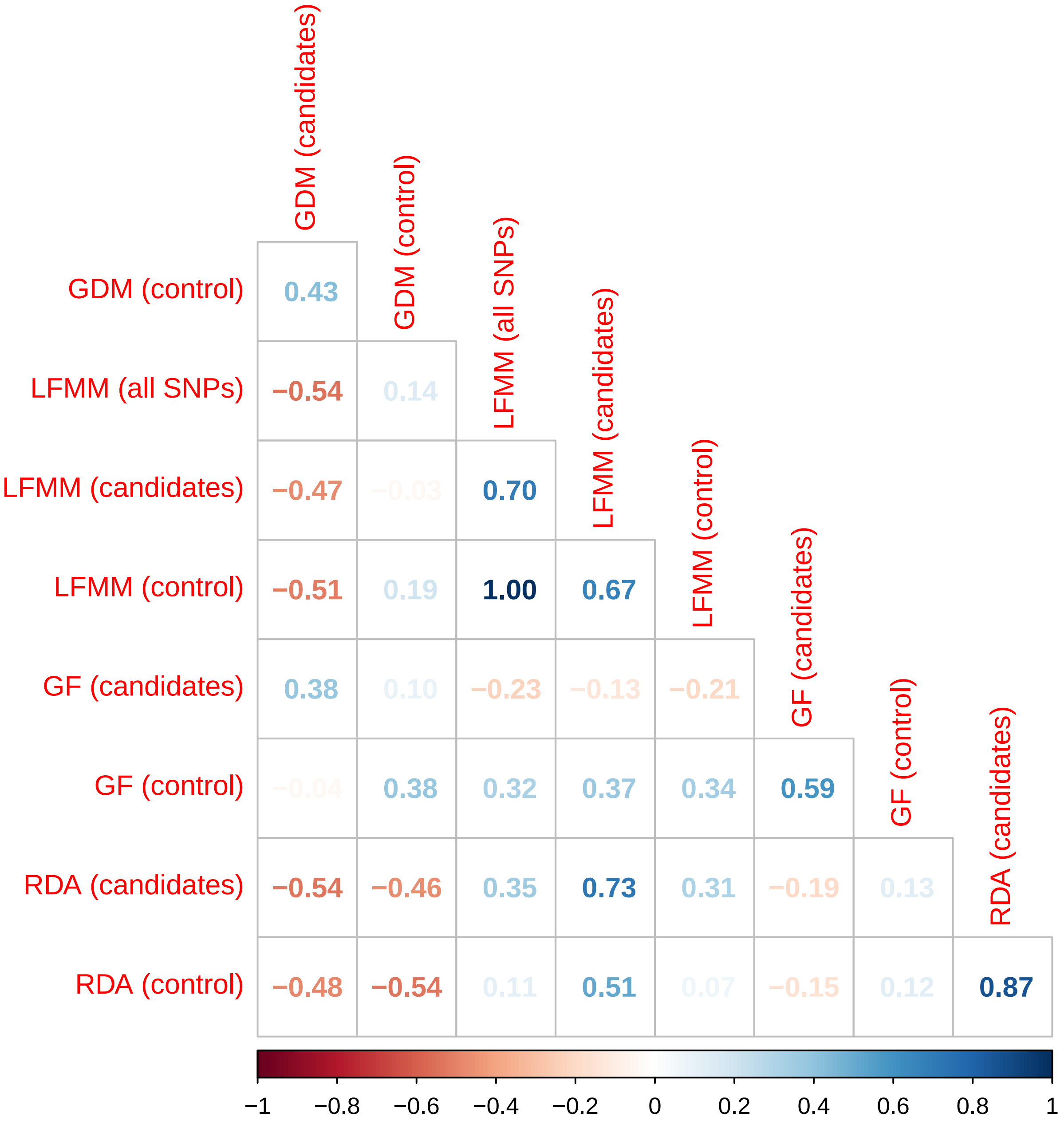

Figure 40: Correlations among GO predictions from the IPSL−CM6A−LR GCM.

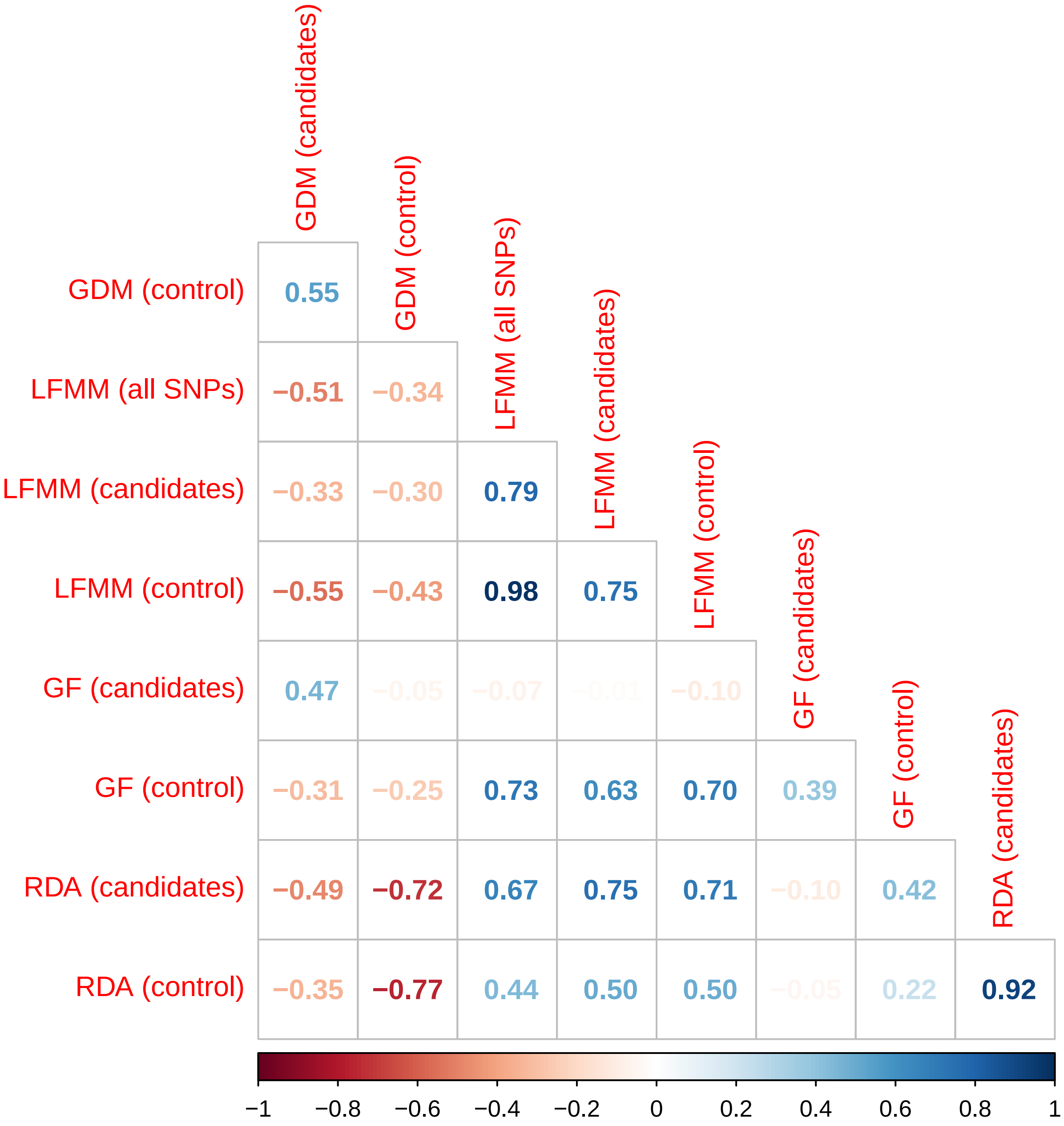

Figure 41: Correlations among GO predictions from the MPI−ESM1−2−HR GCM.

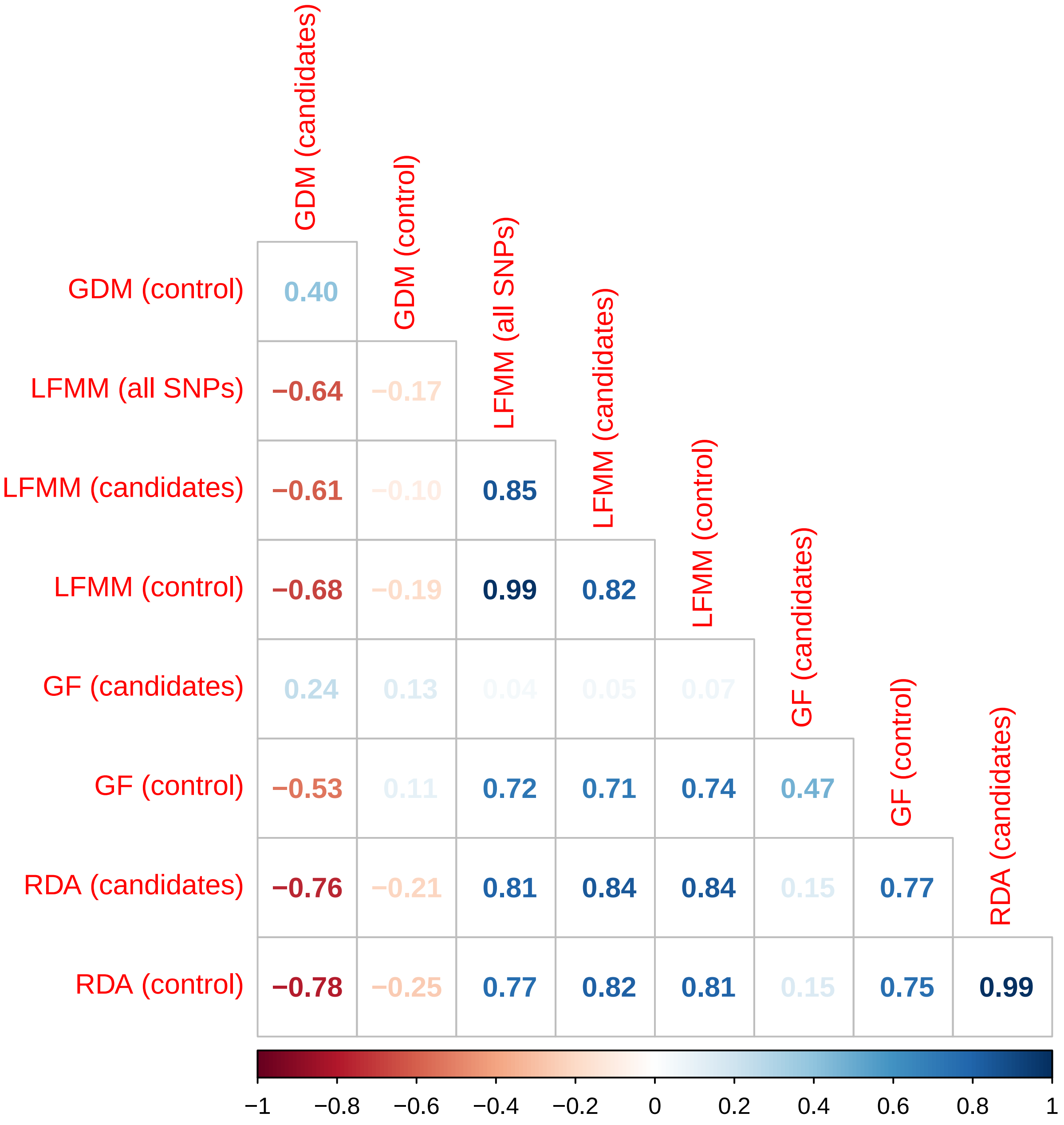

Figure 42: Correlations among GO predictions from the MRI−ESM2−0 GCM.

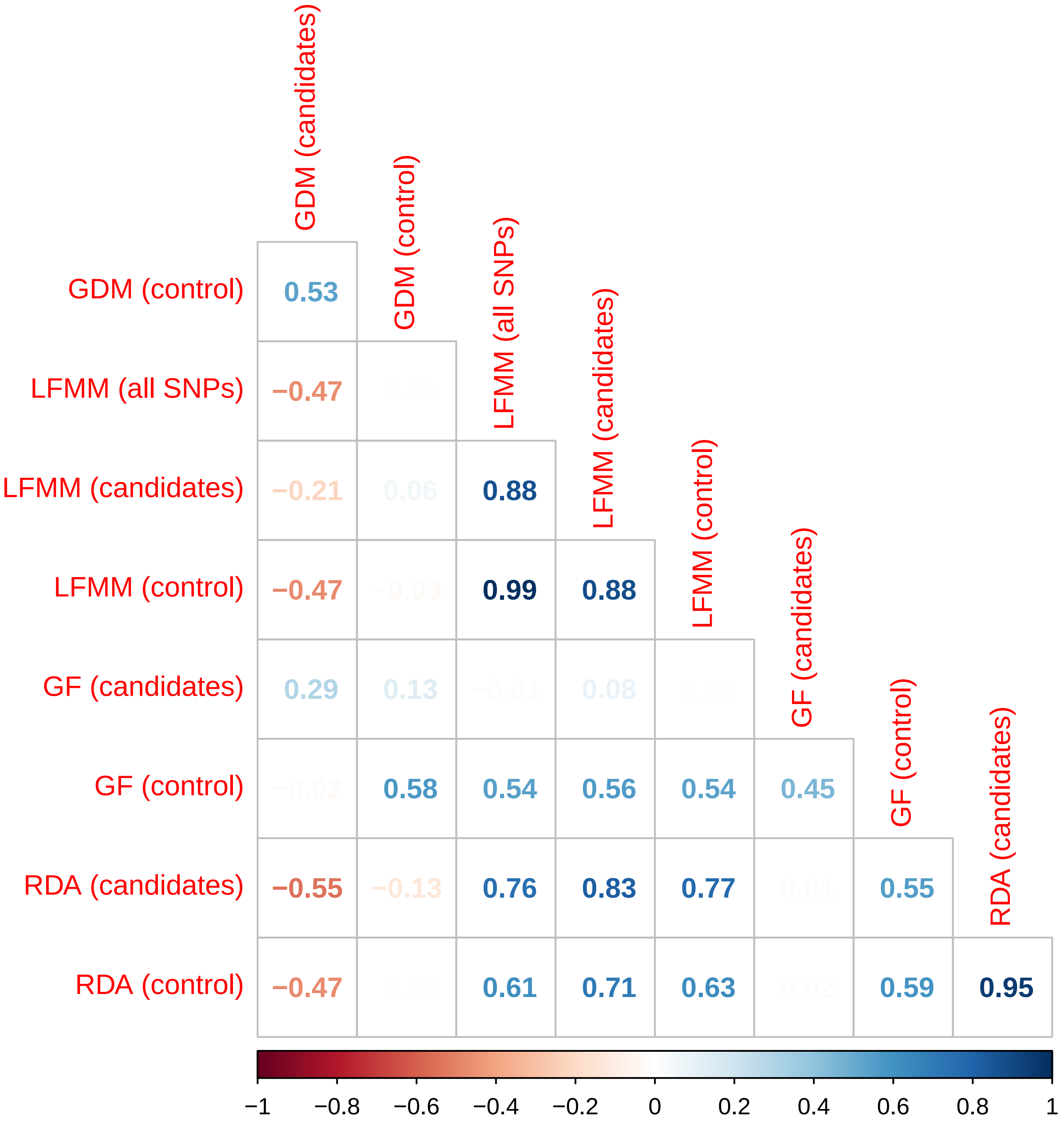

Figure 43: Correlations among GO predictions from the UKESM1-0-LL GCM.

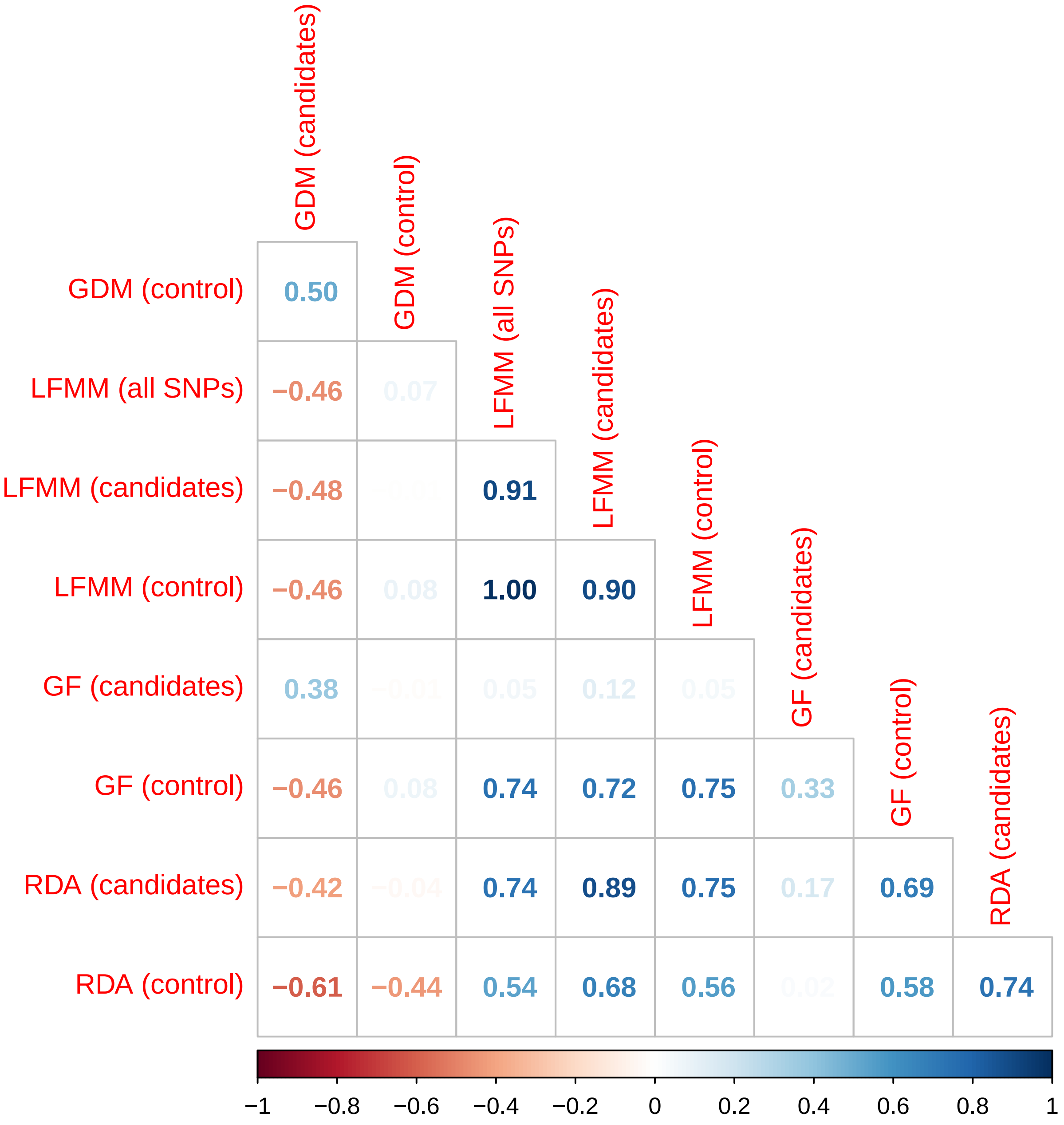

##### 7.6.2 Comparing population ranks

In this section, we compare the population ranks based on their GO predictions from the different methods (GF, GDM, RDA and LFMM), SNP sets (candidates, control and all SNPs for LFMM) and GCMs.

For that, for each combination of method/SNP set/GCM, we ranked the populations based on the GO predictions, i.e. populations with the lowest rank had the highest GO predictions.

###### 7.6.2.1 Across methods/SNP sets

We first compared the population ranks from the different combinations method/SNP set. For that, we generated one plot per GCM. We also generated a plot in which the population ranks are not based on an unique GCM but are based on the average GO predictions across all GCMs. In the plots below, we colored the populations that were among the three populations with the highest GO in at least one combination method/SNP set.

Figure 44: Variability in the population ranks based on GO predictions from the GCM GFDL−ESM4.

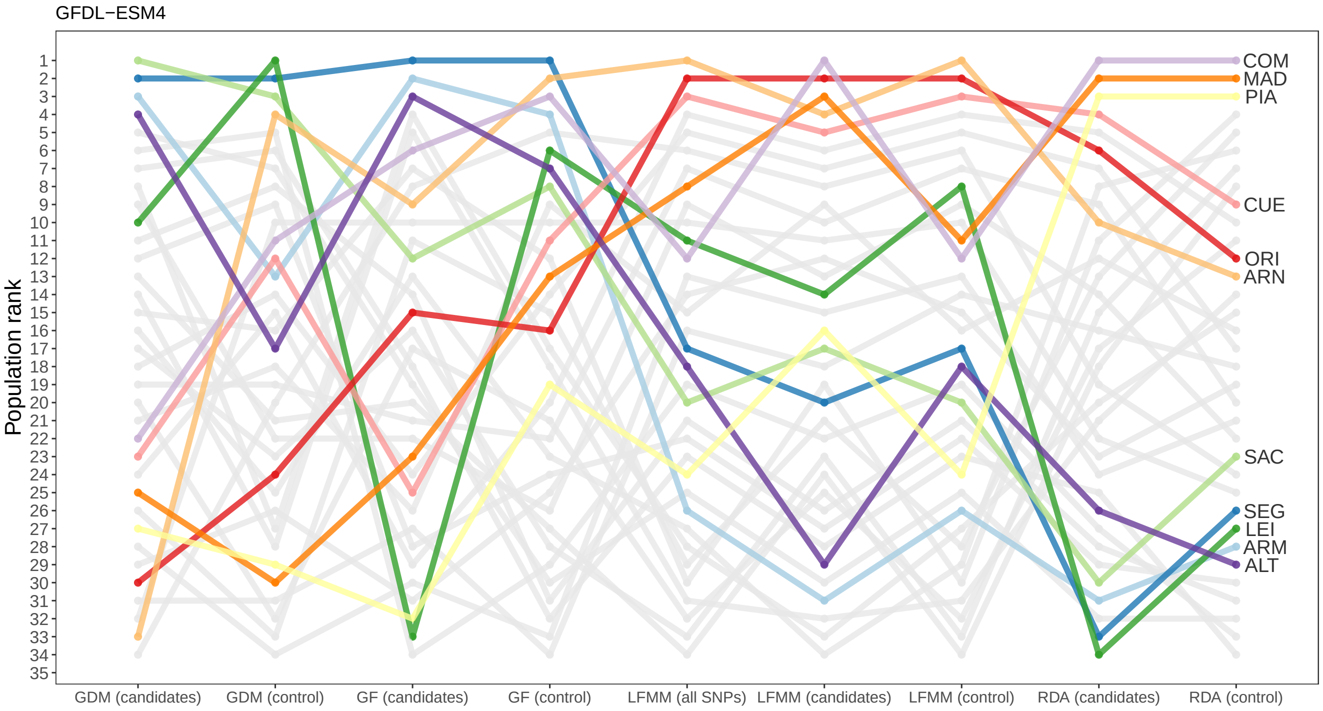

Figure 45: Variability in the population ranks based on GO predictions from the GCM IPSL−CM6A−LR.

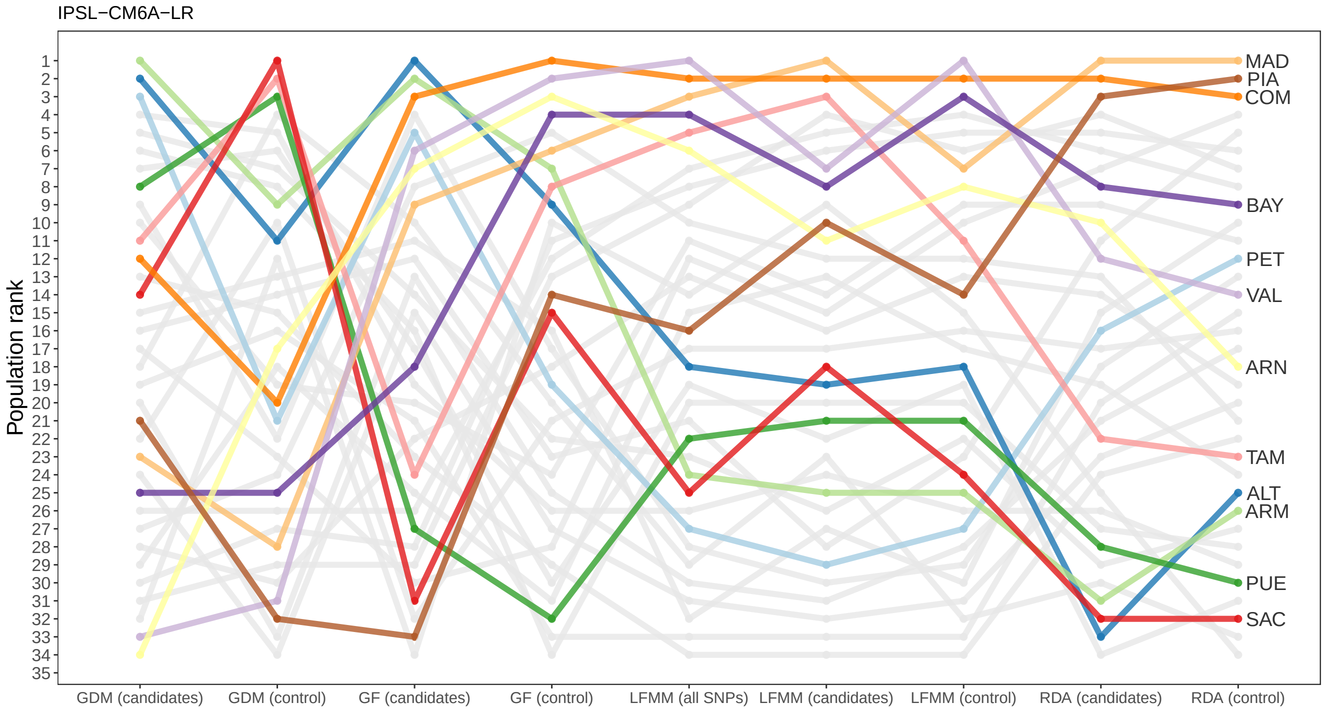

Figure 46: Variability in the population ranks based on GO predictions from the GCM MPI−ESM1−2−HR.

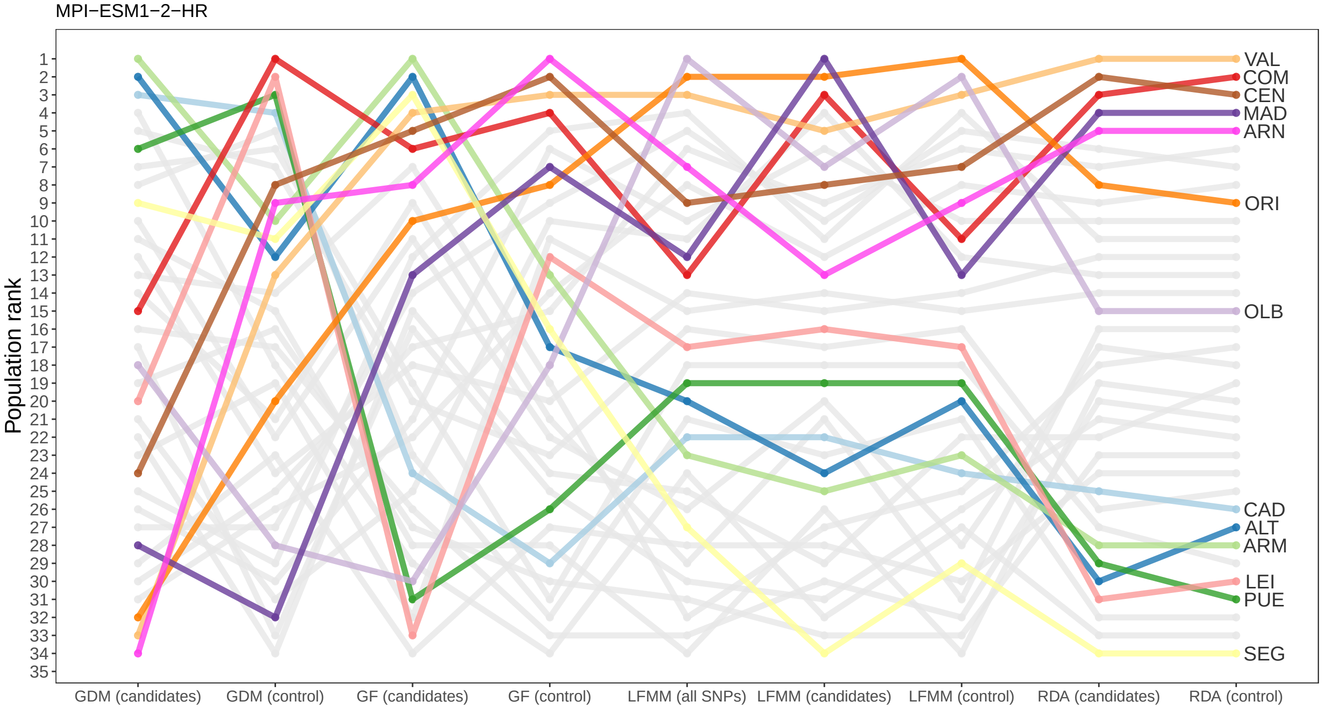

Figure 47: Variability in the population ranks based on GO predictions from the GCM MRI−ESM2−0.

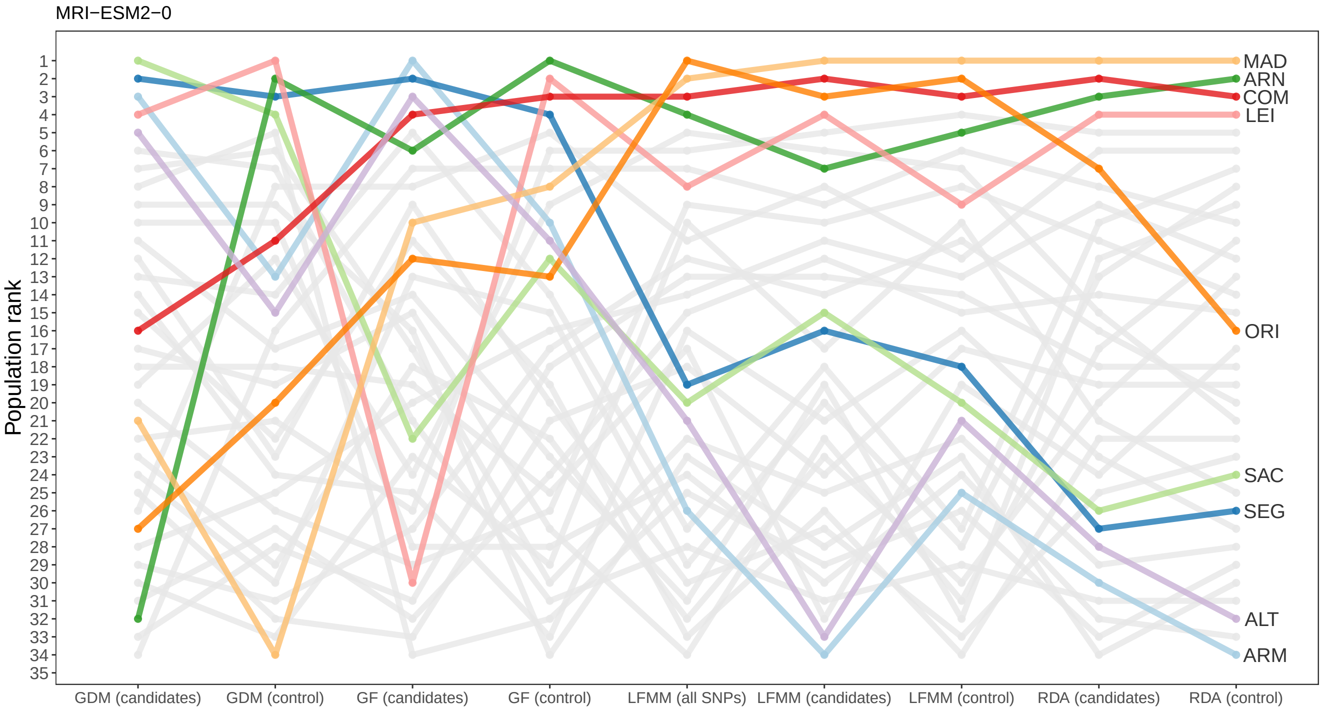

Figure 48: Variability in the population ranks based on GO predictions from the GCM UKESM1−0−LL.

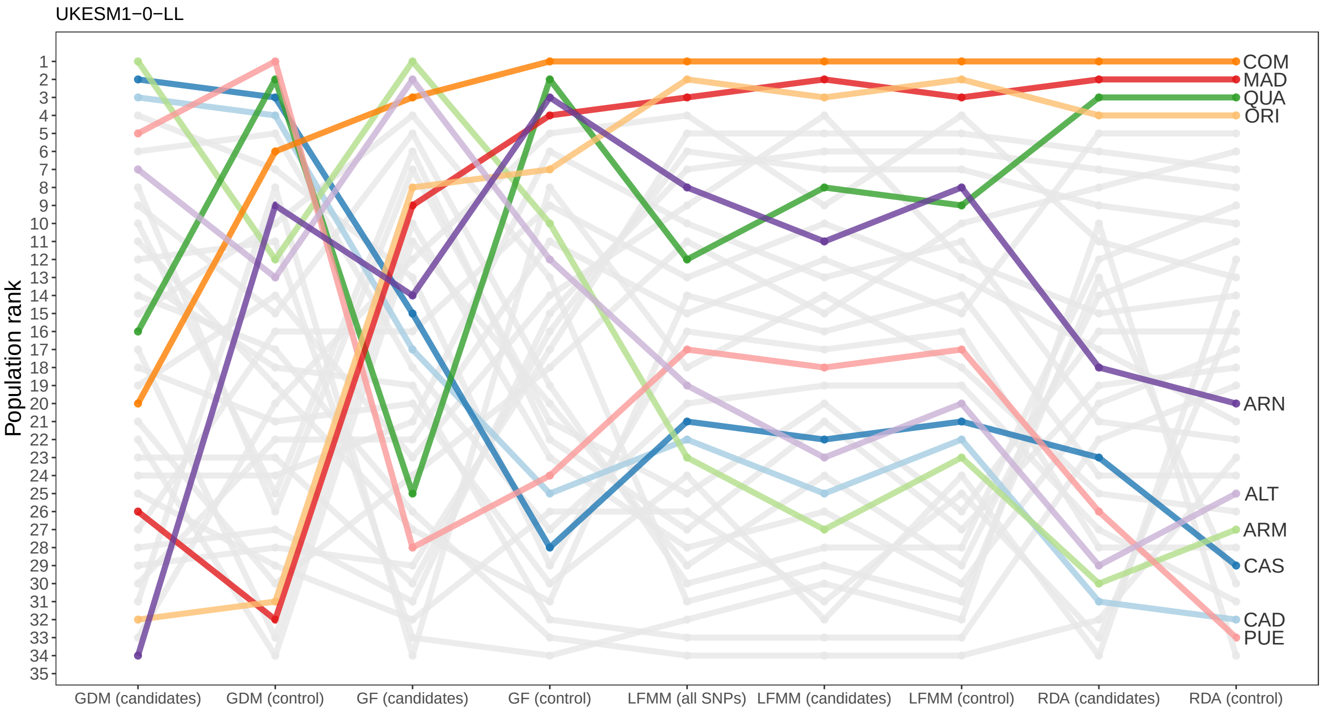

Figure 49: Variability in the population ranks based on the average GO predictions across the five GCMs.

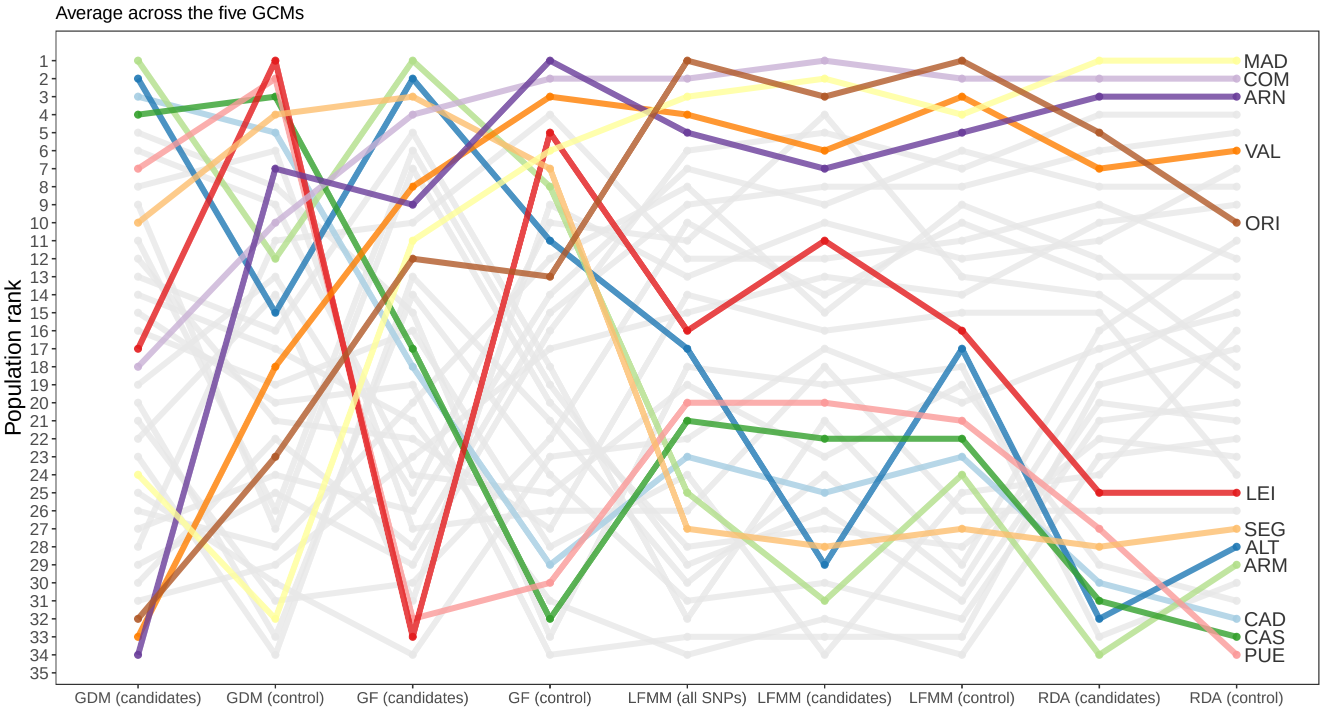

###### 7.6.2.2 Across GCMs

We then compared the population ranks from the different GCMs (or from the average GO predictions across the five GCMs). For that, we generated one plot per combination method/SNP set. We also generated a plot in which the population ranks are not based on an unique combination method/SNP set but are based on the average GO predictions across all combinations method/SNP set.

In the plots below, we colored the populations that were among the three populations with the highest GO in at least one GCM.

Figure 50: Variability in the population ranks based on GO predictions from the GDM approach and the set of candidate SNPs.

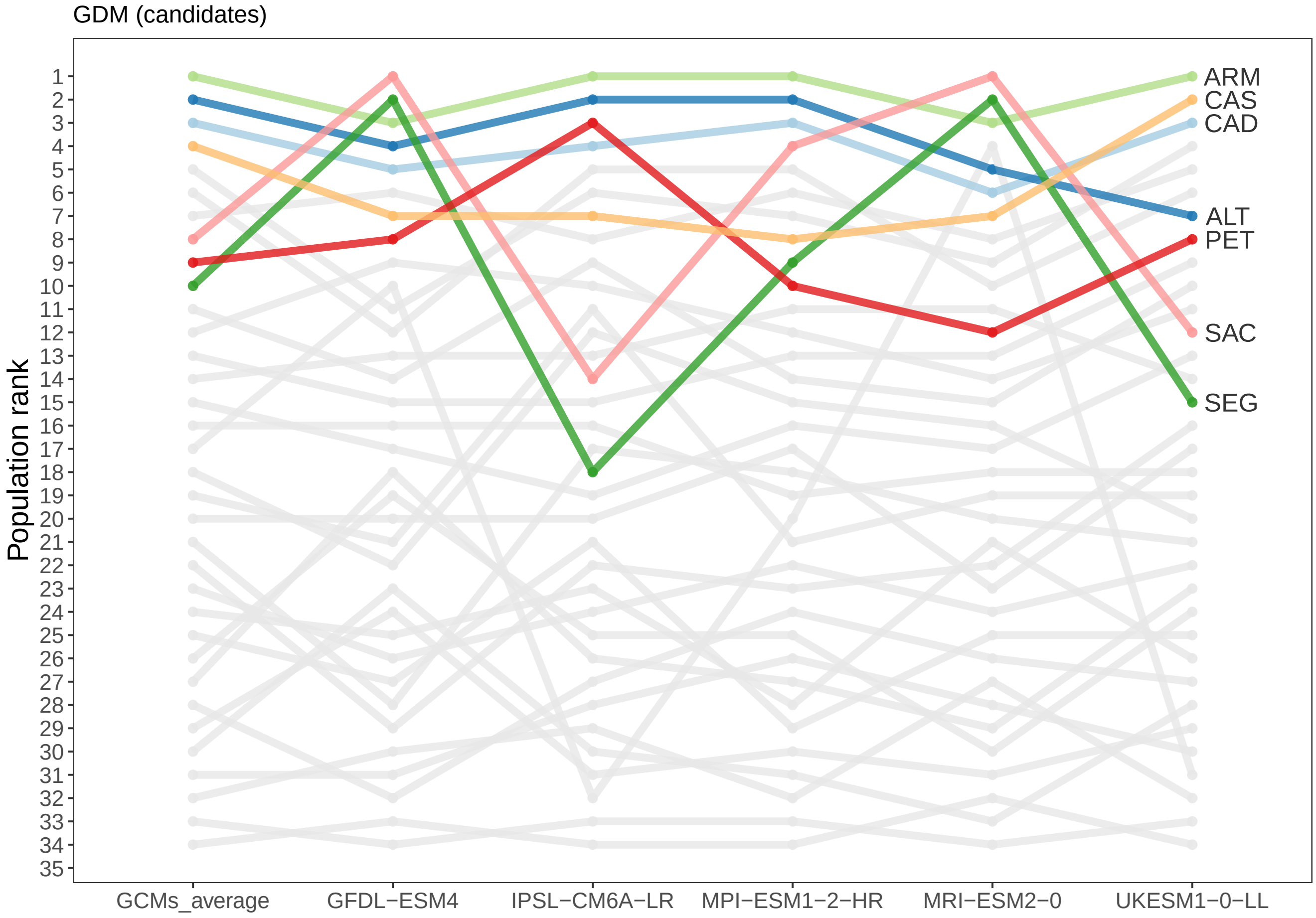

Figure 51: Variability in the population ranks based on GO predictions from the GDM approach and the set of control SNPs.

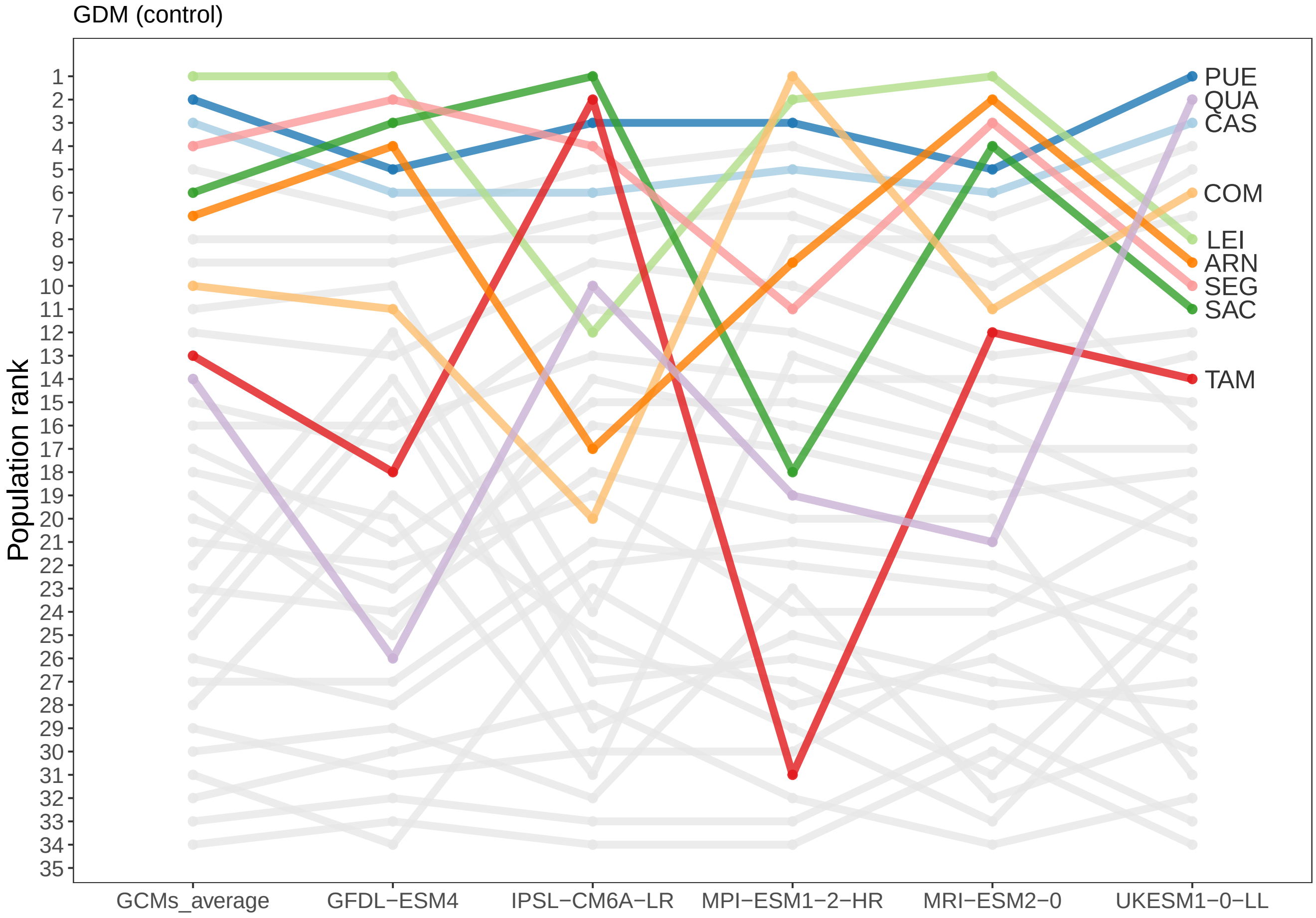

Figure 52: Variability in the population ranks based on GO predictions from the GF approach and the set of candidate SNPs.

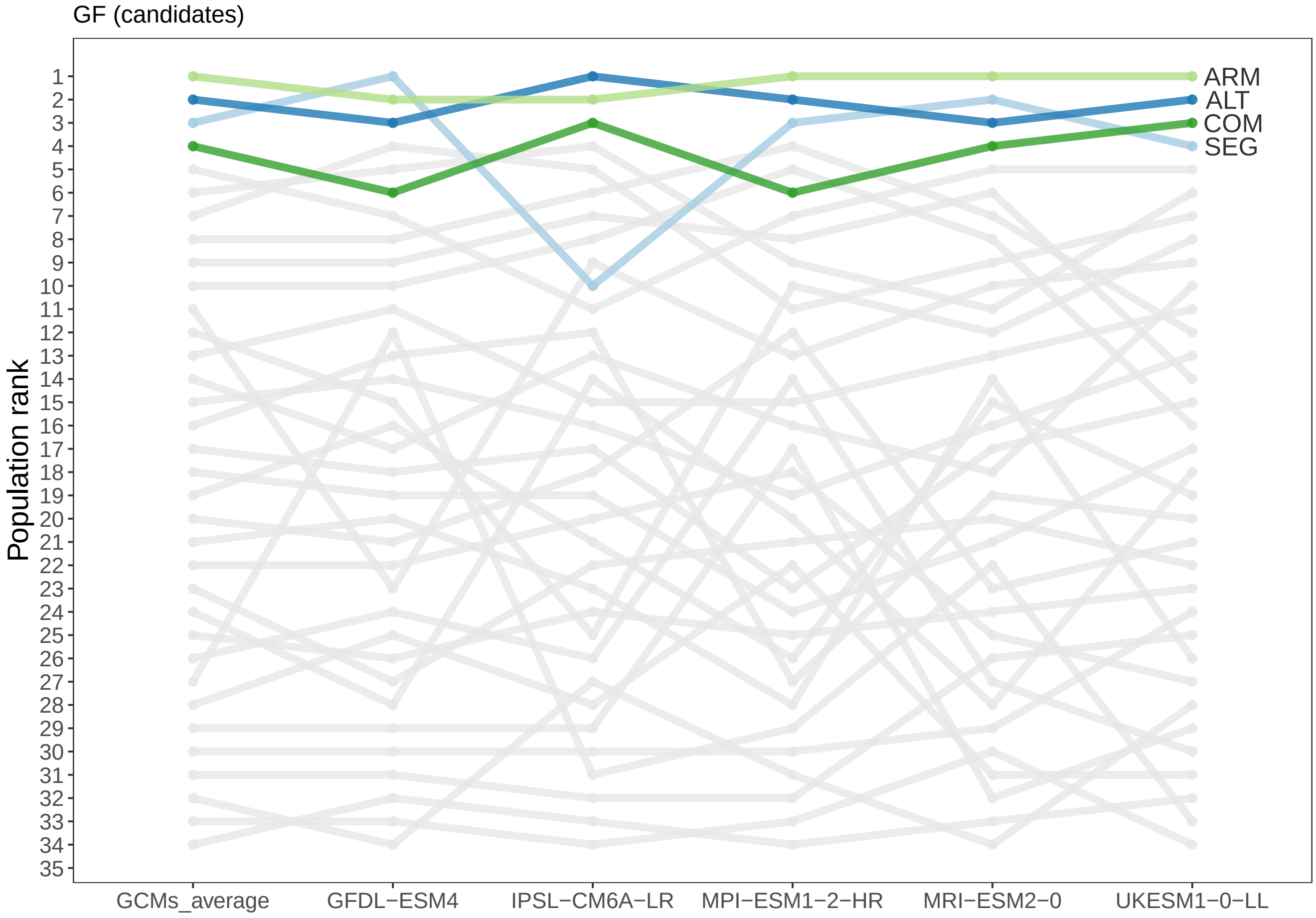

Figure 53: Variability in the population ranks based on GO predictions from the GF approach and the set of control SNPs.

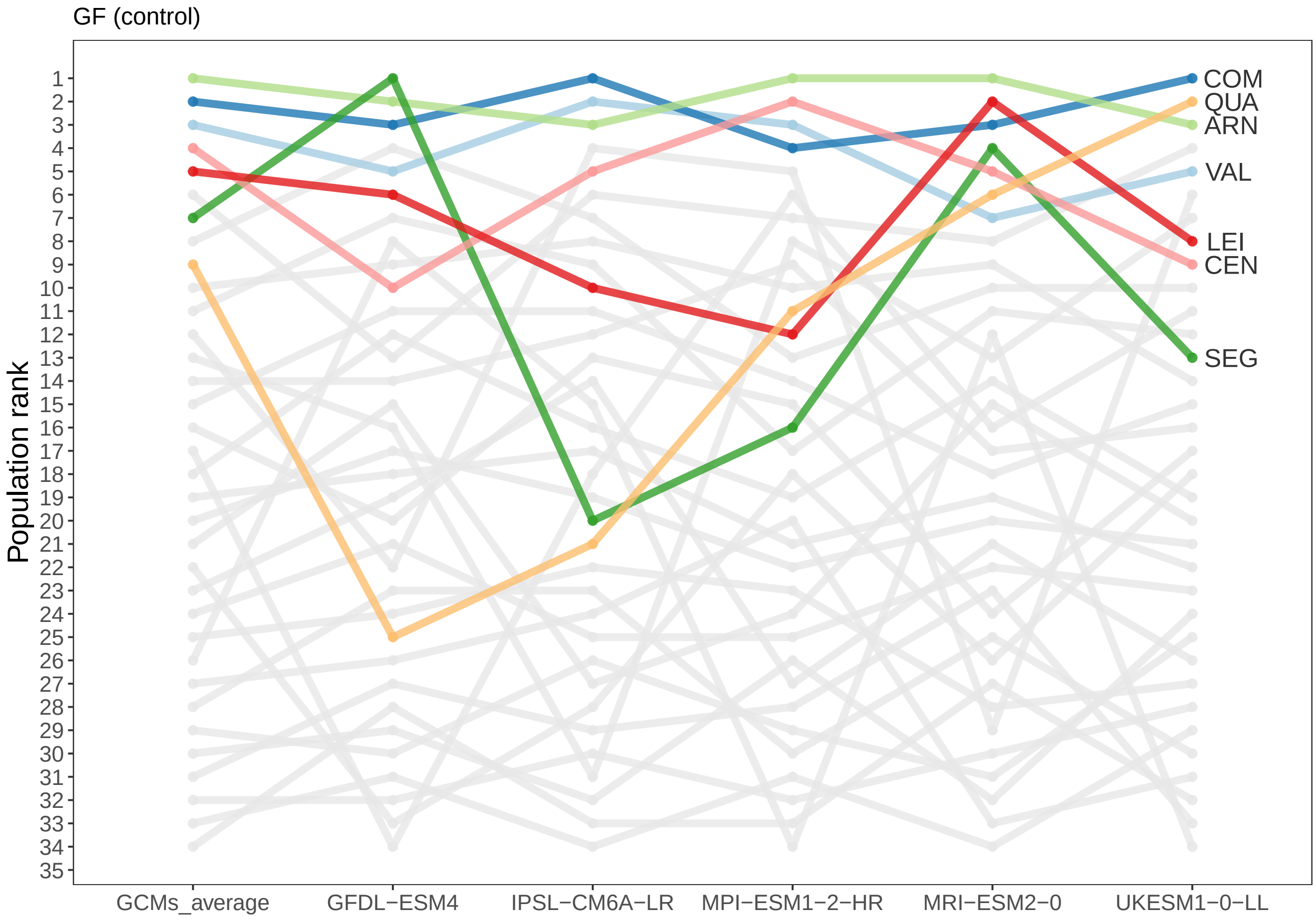

Figure 54: Variability in the population ranks based on GO predictions from the LFMM approach and the set of candidate SNPs.

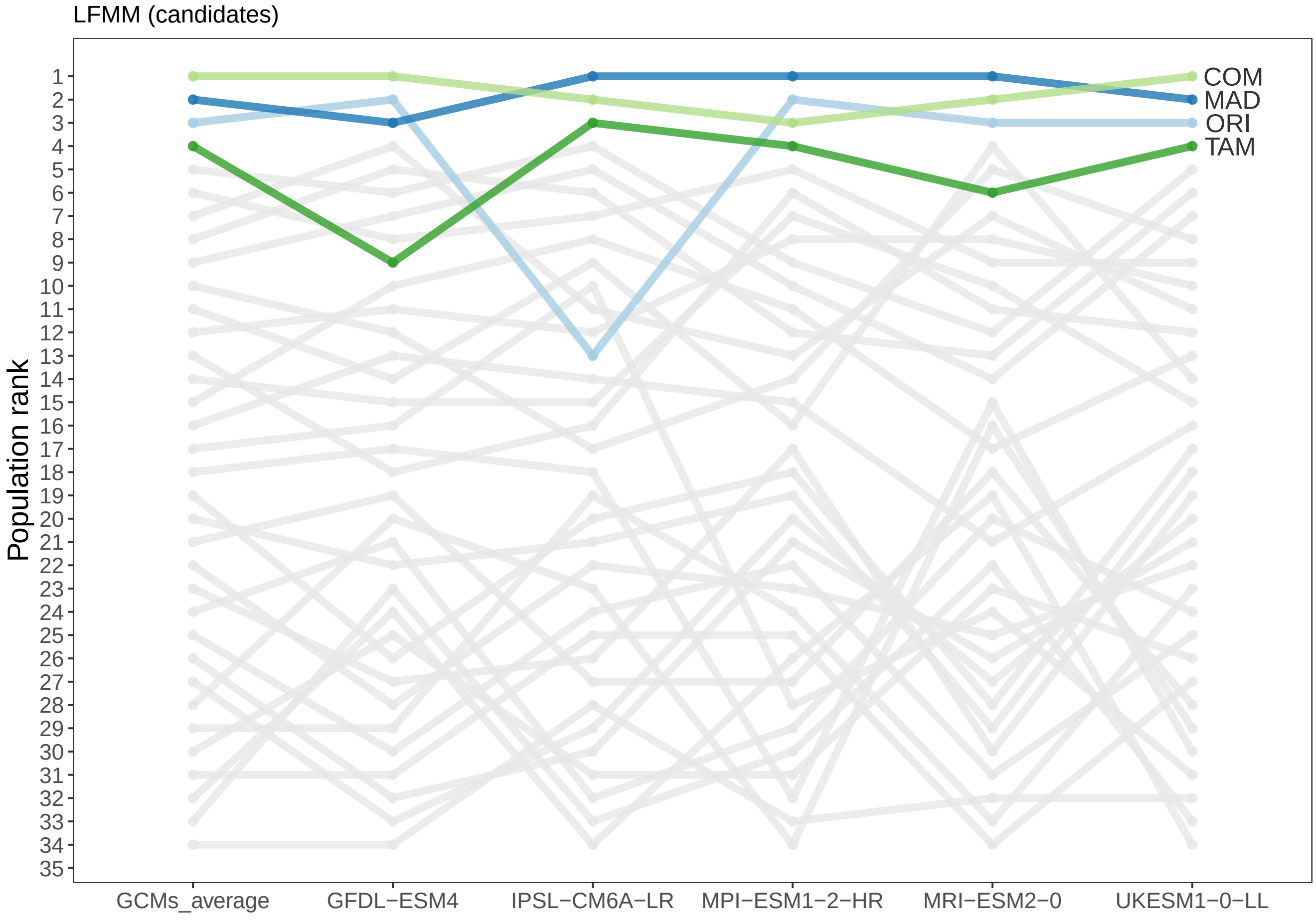

Figure 55: Variability in the population ranks based on GO predictions from the LFMM approach and the set of control SNPs.

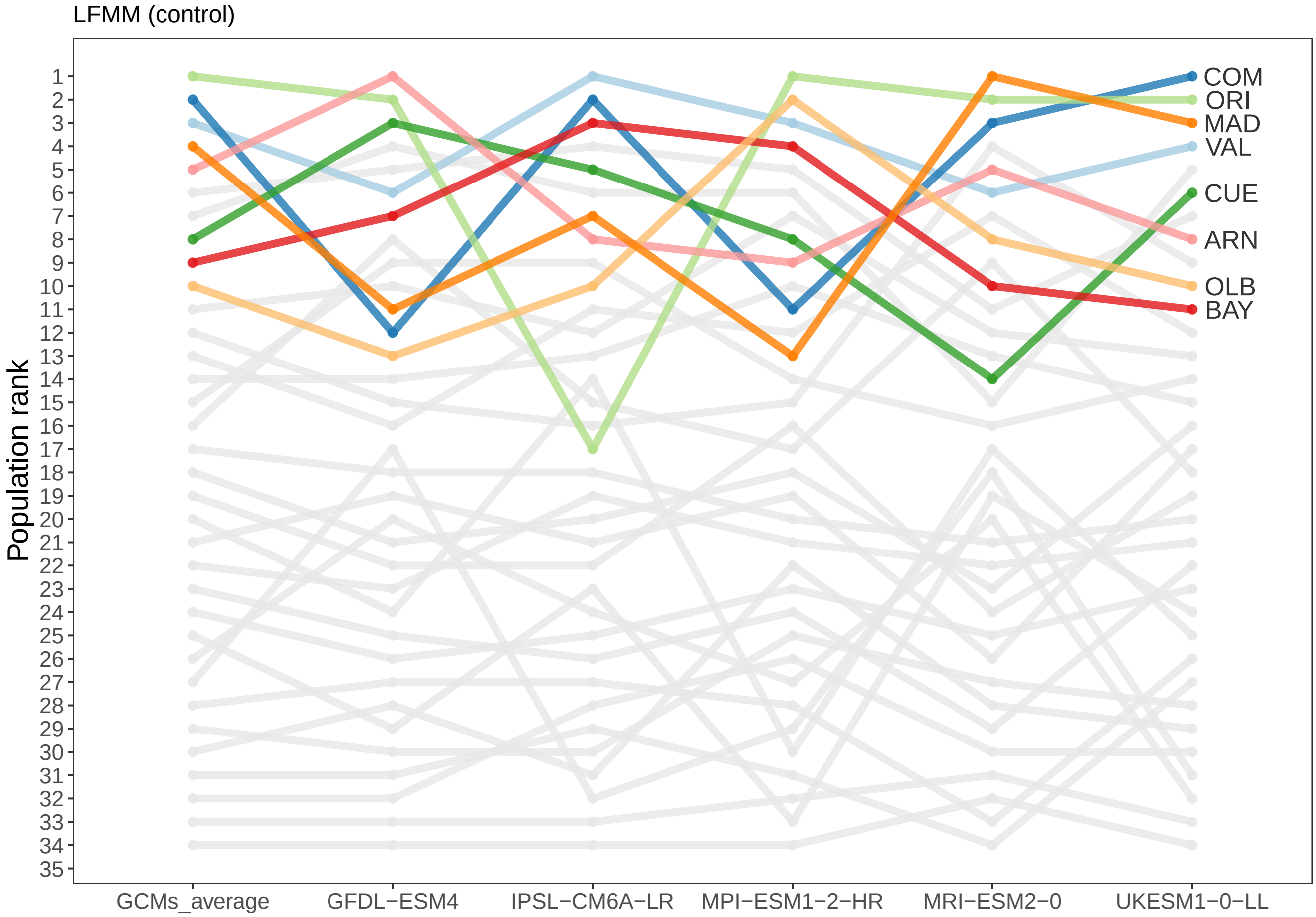

Figure 56: Variability in the population ranks based on GO predictions from the LFMM approach and all SNPs.

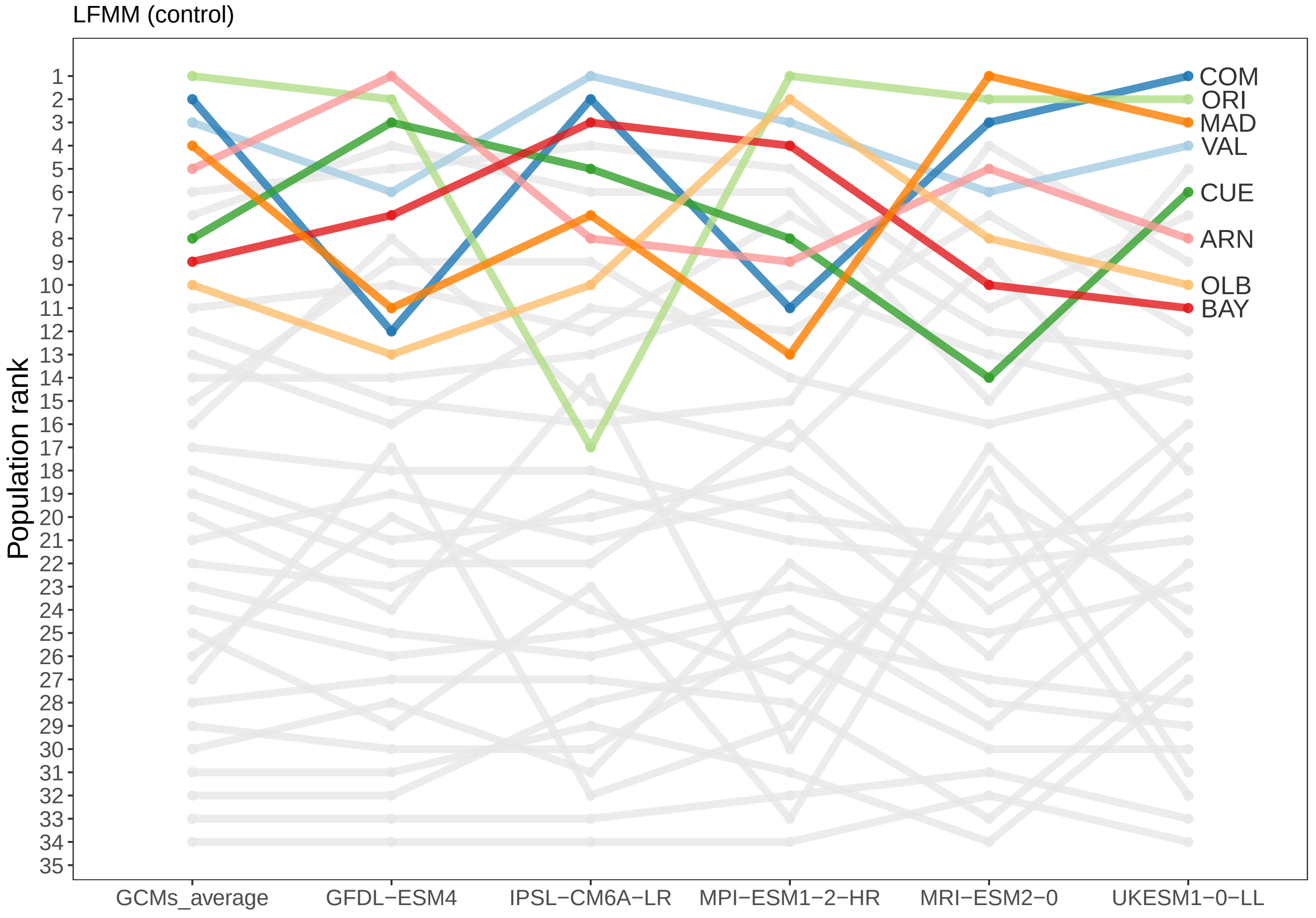

Figure 57: Variability in the population ranks based on GO predictions from the RDA approach and the set of candidate SNPs.

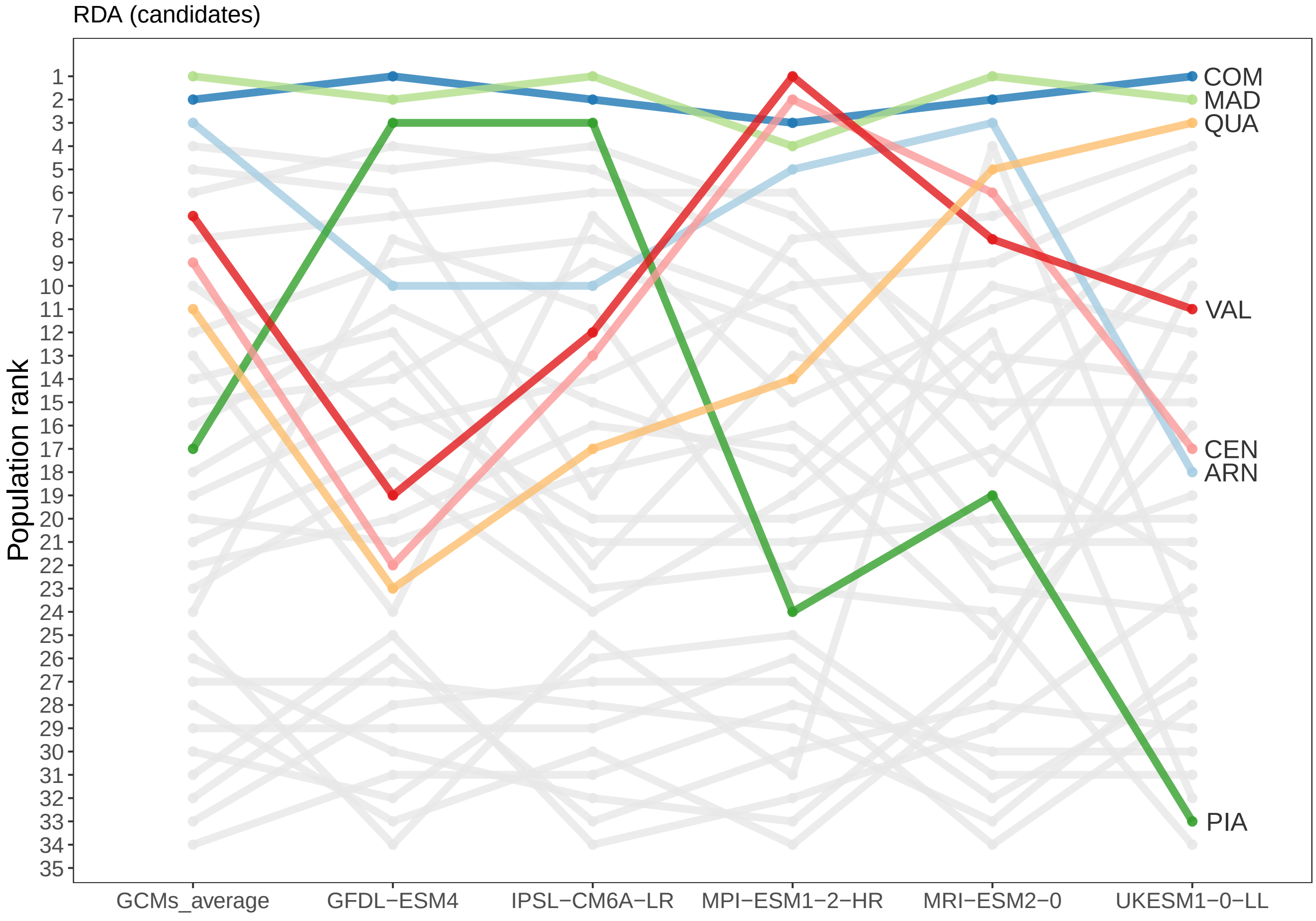

Figure 58: Variability in the population ranks based on GO predictions from the RDA approach and the set of control SNPs.

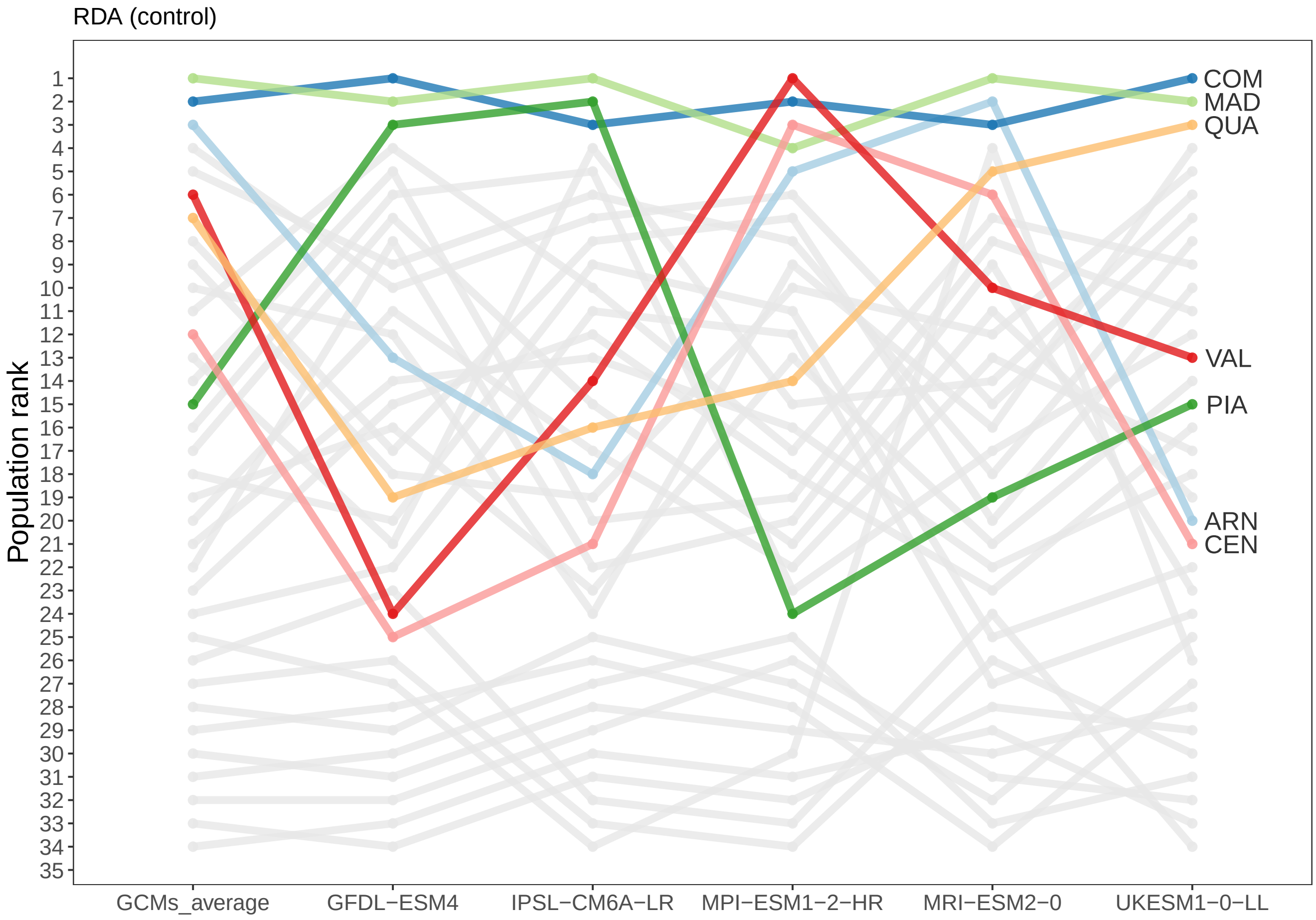

### 8 Validation in the NFI plots

#### 8.1 GO predictions at the location of the NFI plots

##### 8.1.1 Time periods

To extrapolate GO predictions at the location of the NFI plots, we used:

- the average climates under the time period 1901-1950, which are expected to represent the past climates under which the populations have evolved. The time period 1901-1950 is the same reference period as the one used to predict GO at the location of the populations under future climates.
- the average climates for the period during which mortality rates were recorded in the NFI plots and the five preceding years (to account for potential lag climatic effects on tree death). In the

French NFI, a tree was considered dead if its death was estimated within the five years preceding the inventory date. Consequently, for the French inventory, the period used to make GO predictions covered the ten years preceding the inventory date. For the Spanish inventory, the period used to make GO predictions covered the five years preceding the first inventory date and the period between the two inventories.

##### 8.1.2 Climatic space

Projections of the GO predictions outside of the climatic space covered by the sampled populations have to be taken with caution. The figures below (and Figure 4a in the main manuscript) show the climatic space covered by the NFI plots and the 34 sampled populations for the reference period 1901-1950.

Figure 59: Annual precipitation and mean annual temperature at the location of the NFI plots (in gray) and the 34 sampled populations (colored according their main gene pool). Climatic values correspond to average climates over the reference period 1901-1950.

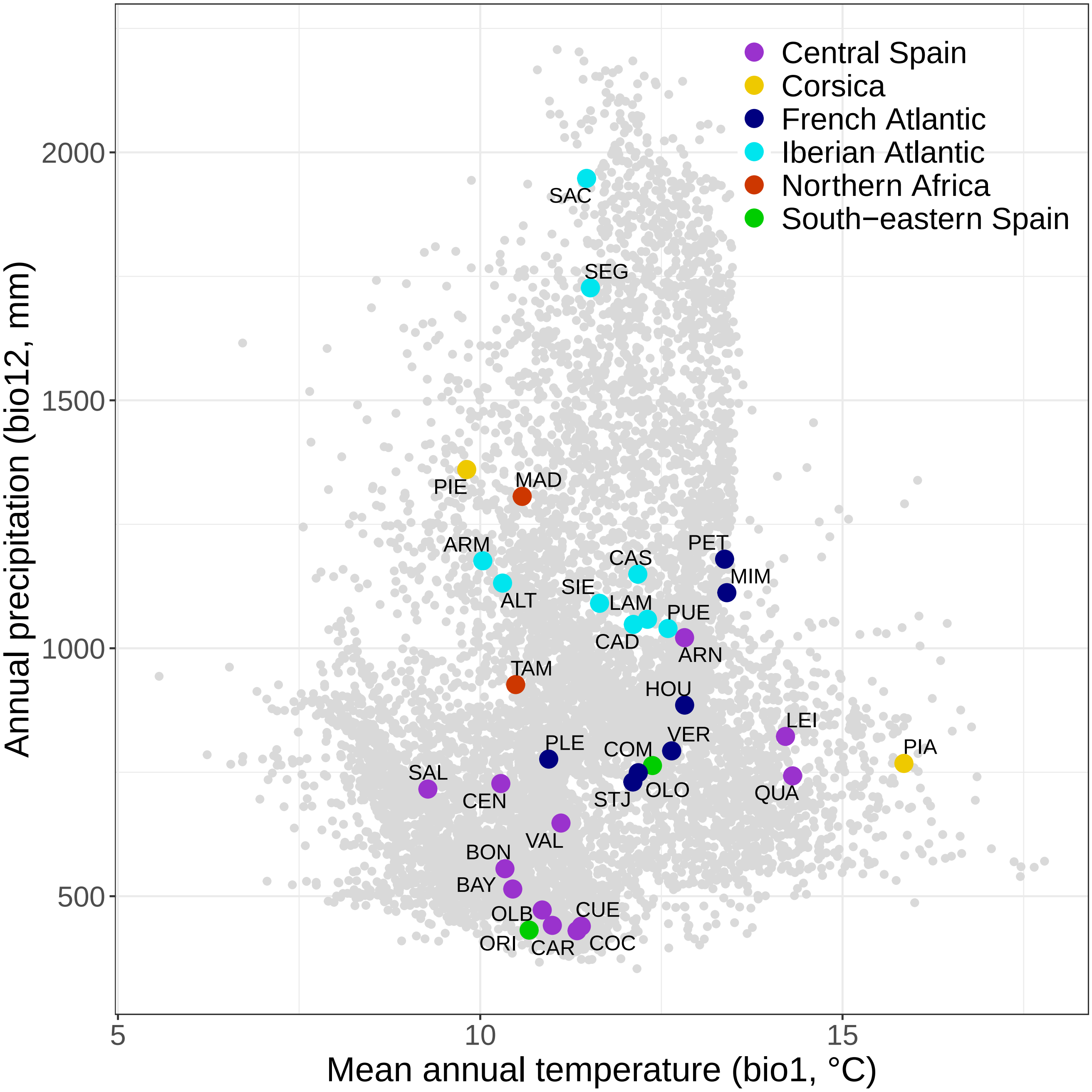

Figure 60: Isothermality and precipitation seasonality at the location of the NFI plots (in gray) and the 34 sampled populations (colored according their main gene pool). Climatic values correspond to average climates over the reference period 1901-1950.

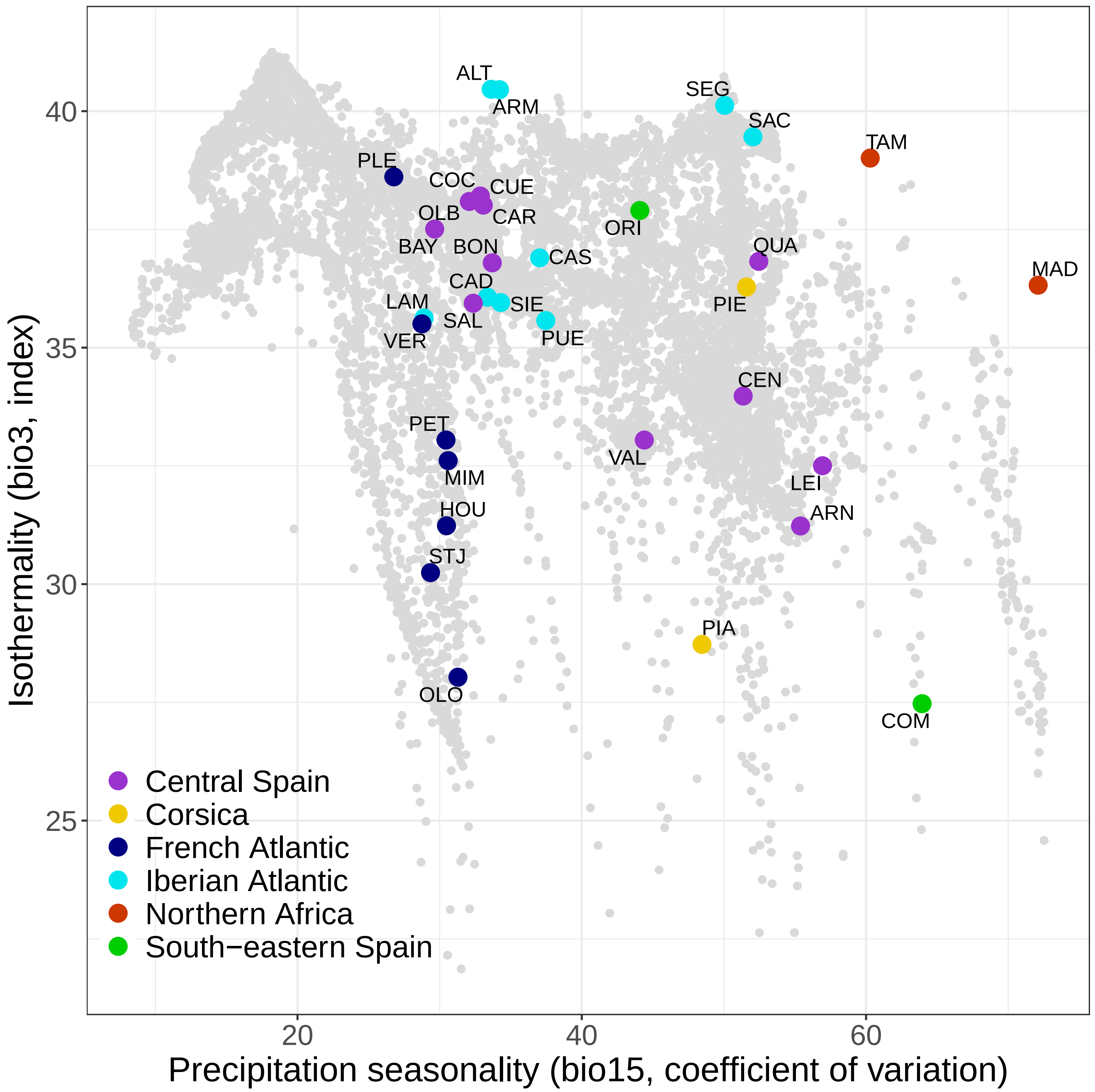

##### 8.1.3 GO predictions with RDA

Figure 61: GO predictions with RDA and the candidate SNPs at the location of the NFI plots.

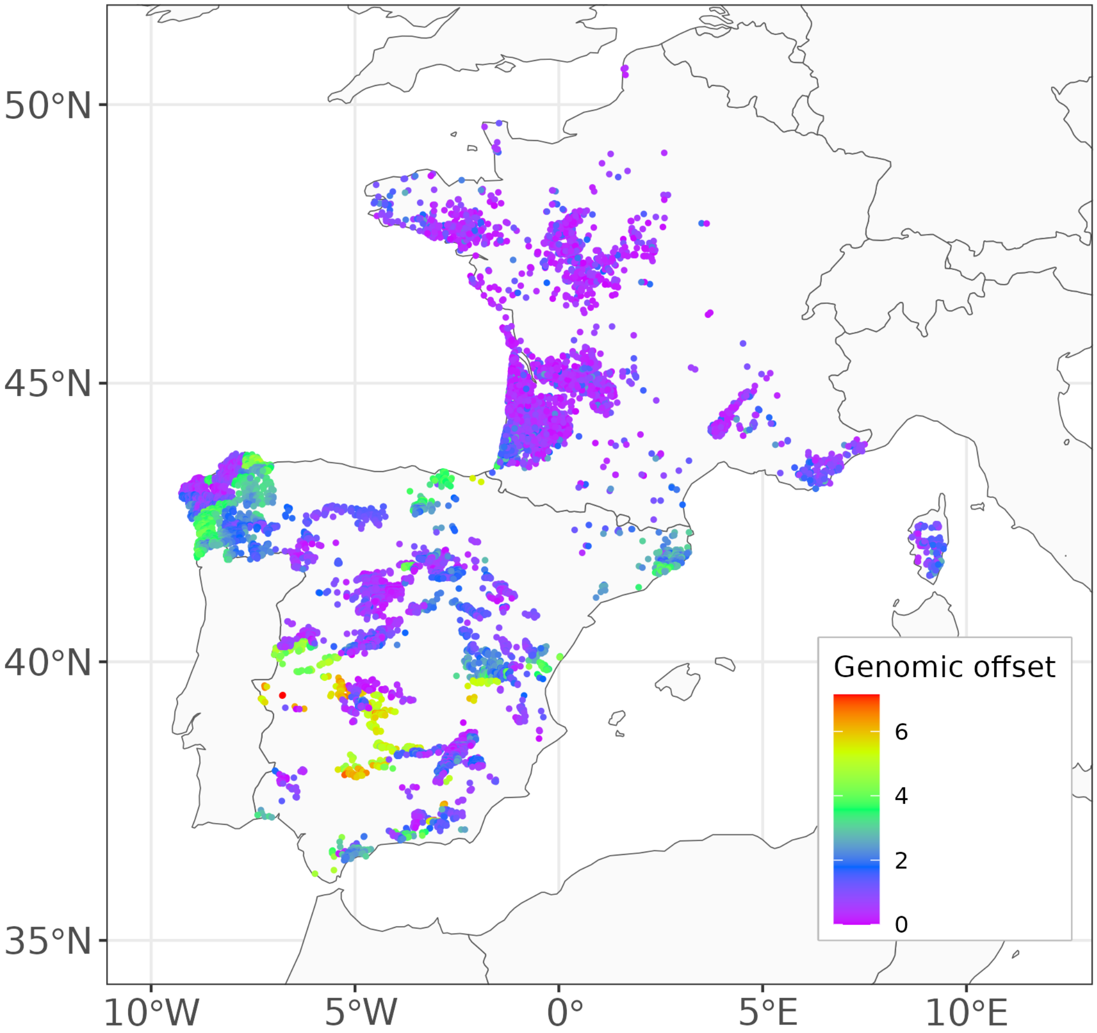

Figure 62: GO predictions with RDA and the control SNPs at the location of the NFI plots.

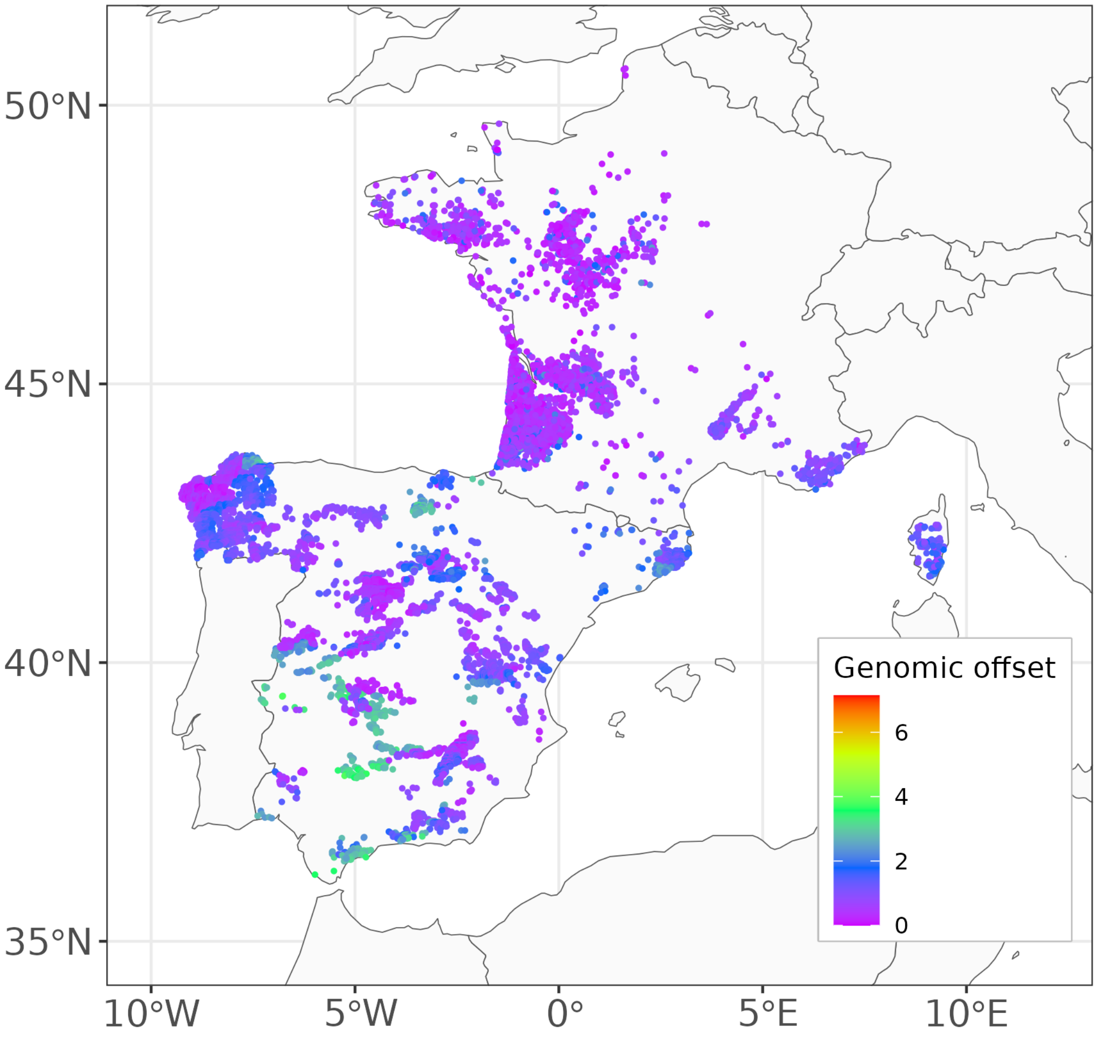

##### 8.1.4 GO predictions with GF

GO predictions with GF and the candidate SNPs are shown in Figure 4c of the main manuscript.

Figure 63: GO predictions with GF and the control SNPs at the location of the NFI plots.

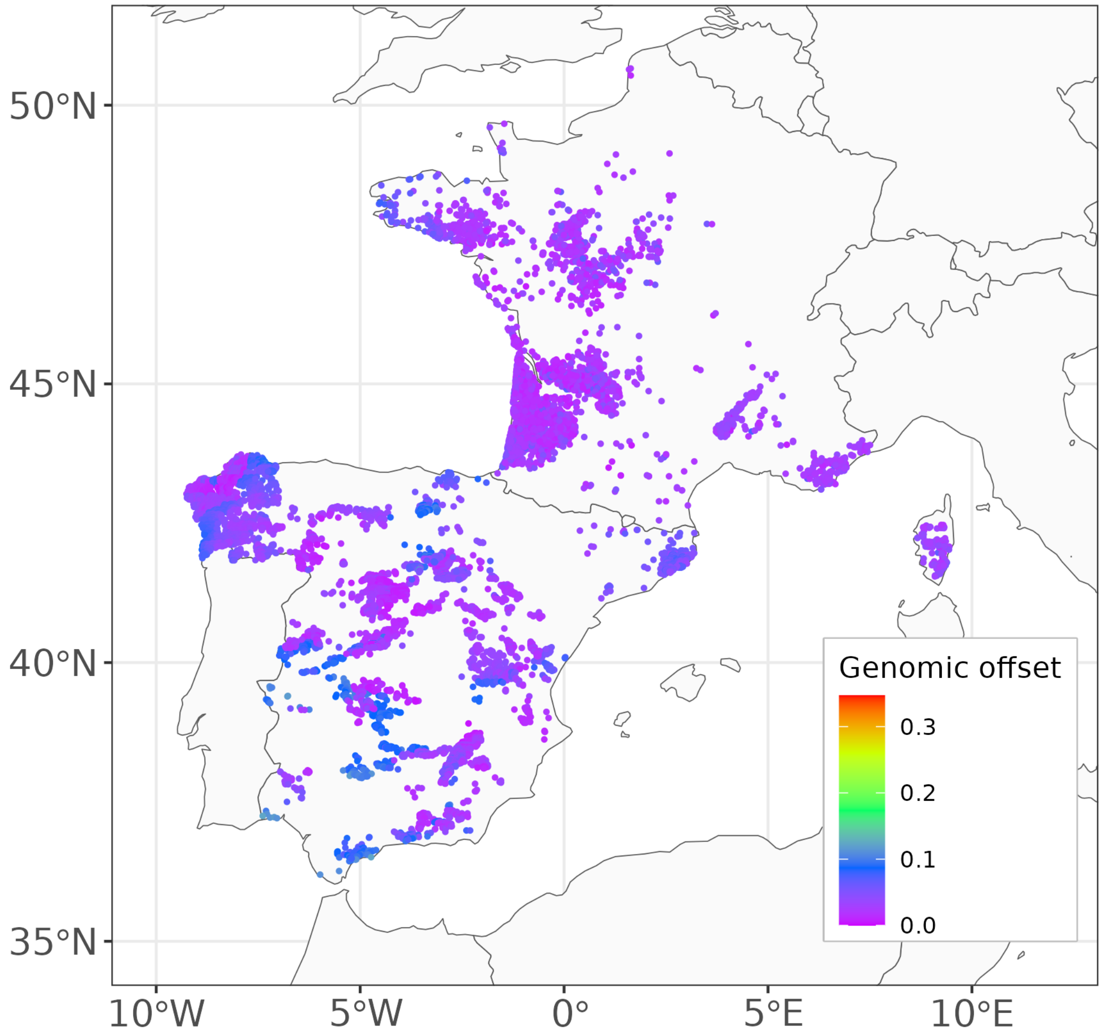

###### 8.1.5 GO predictions with LFMM

Figure 64: GO predictions with LFMM and all SNPs at the location of the NFI plots.

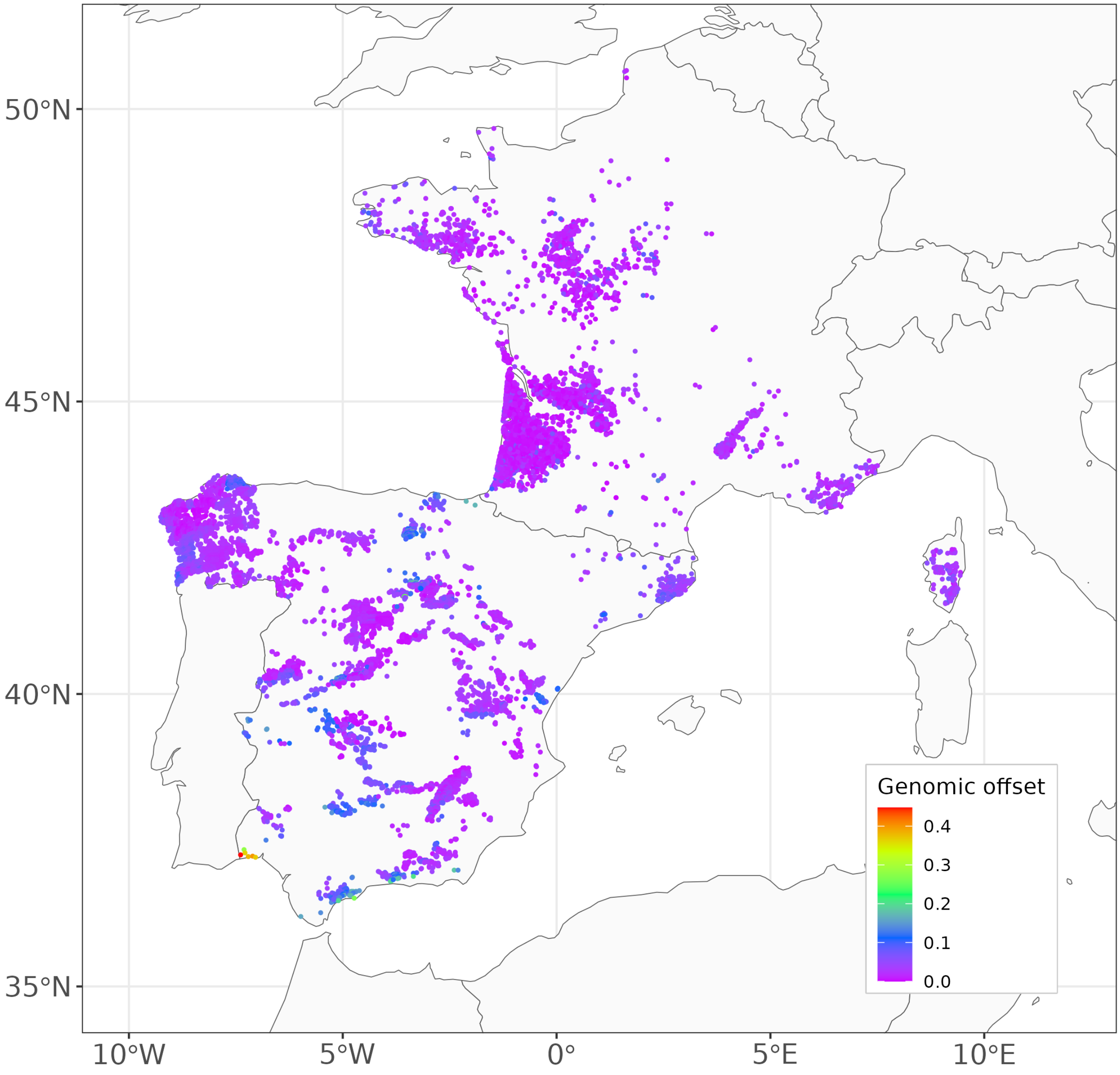

Figure 65: GO predictions with LFMM and the candidate SNPs at the location of the NFI plots.

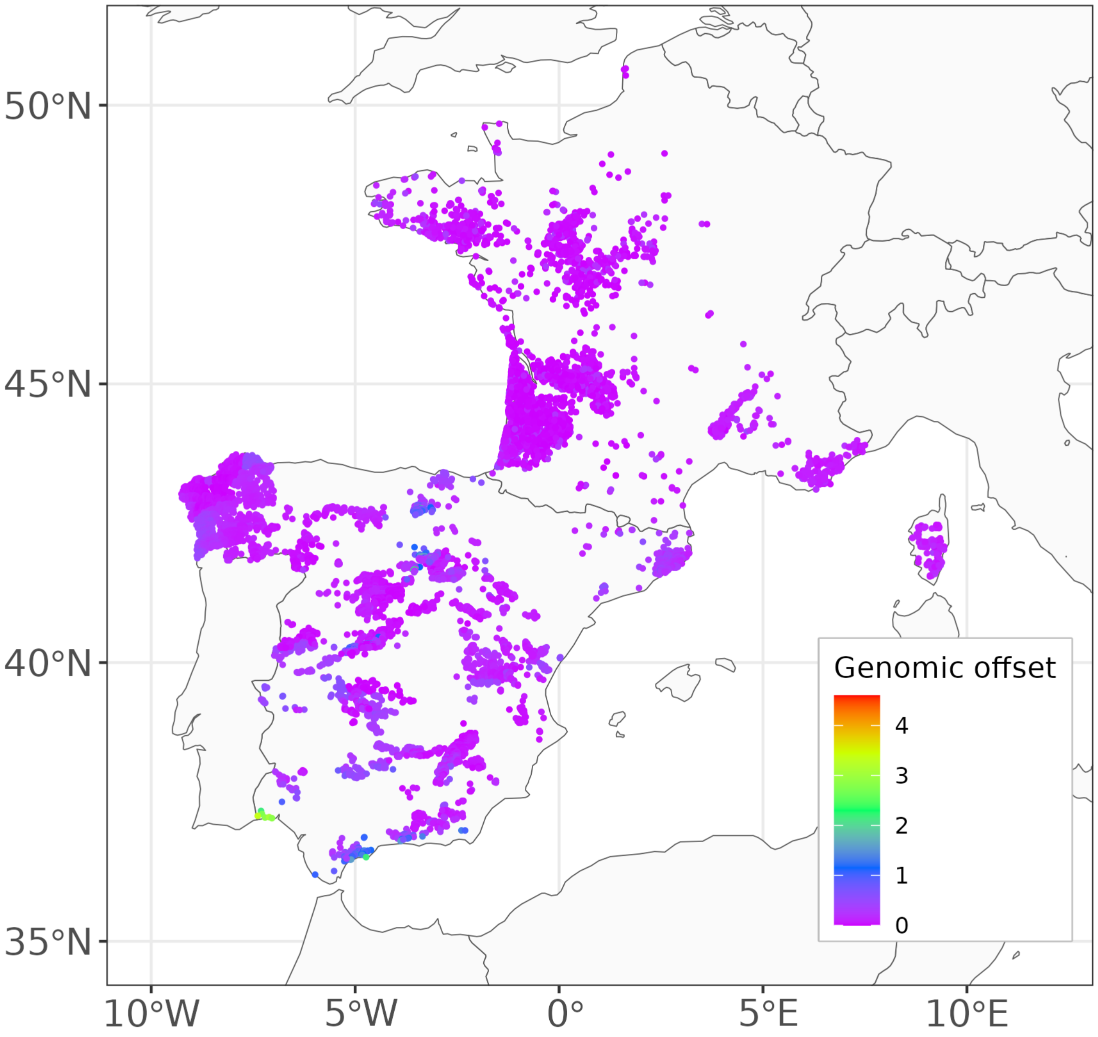

Figure 66: GO predictions with LFMM and the control SNPs at the location of the NFI plots.

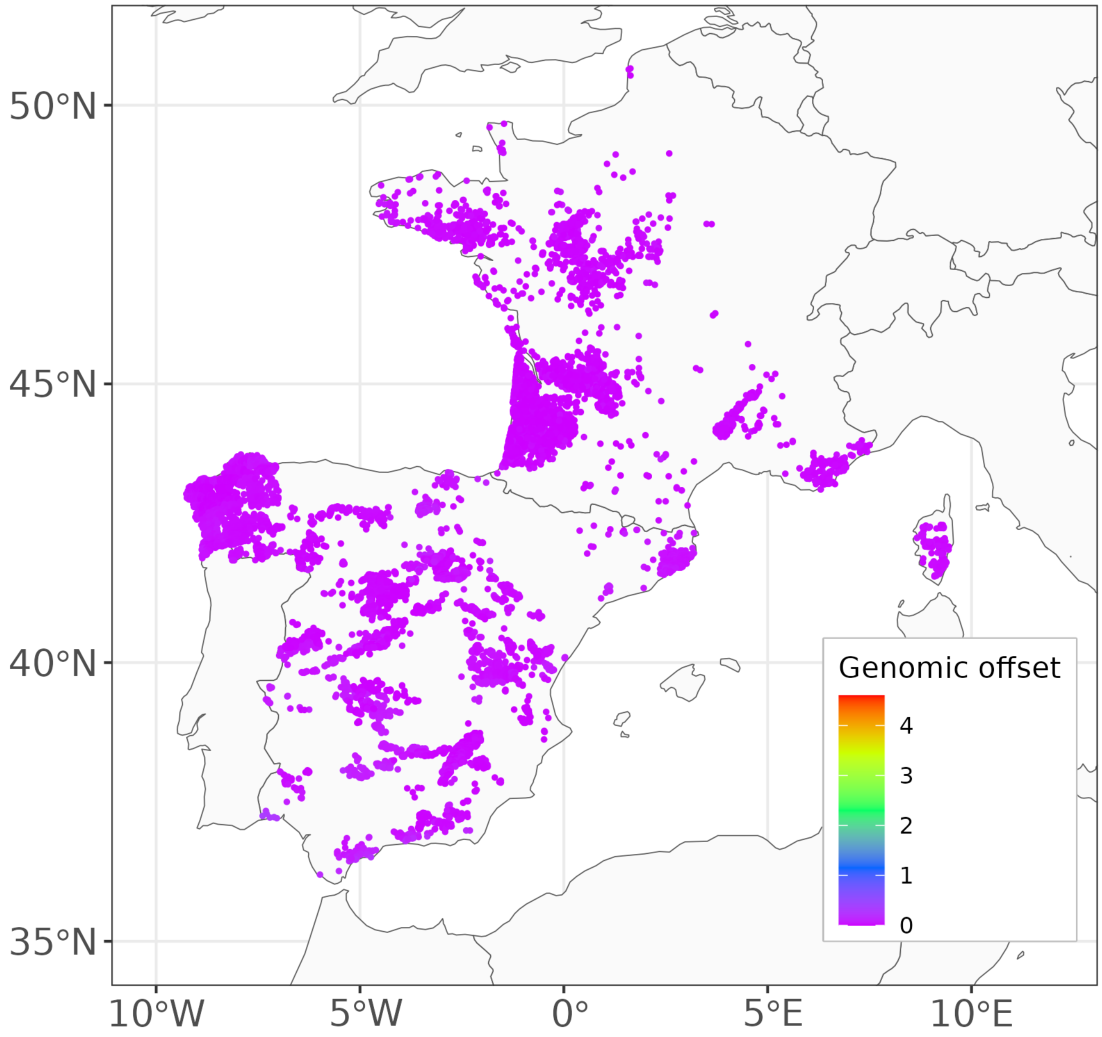

##### 8.1.6 GO predictions with GDM

Figure 67: GO predictions with GDM and the candidate SNPs at the location of the NFI plots.

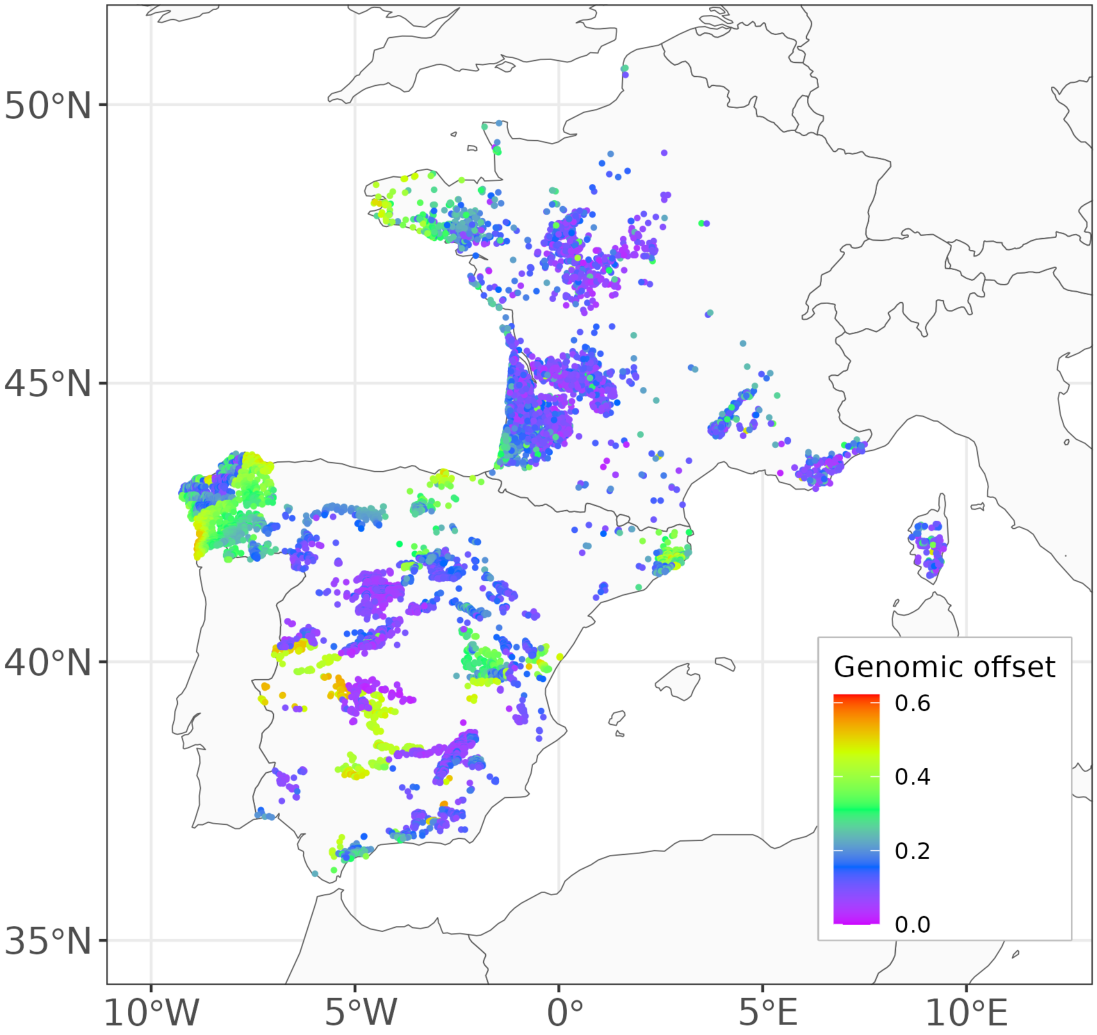

Figure 68: GO predictions with GDM and the control SNPs at the location of the NFI plots.

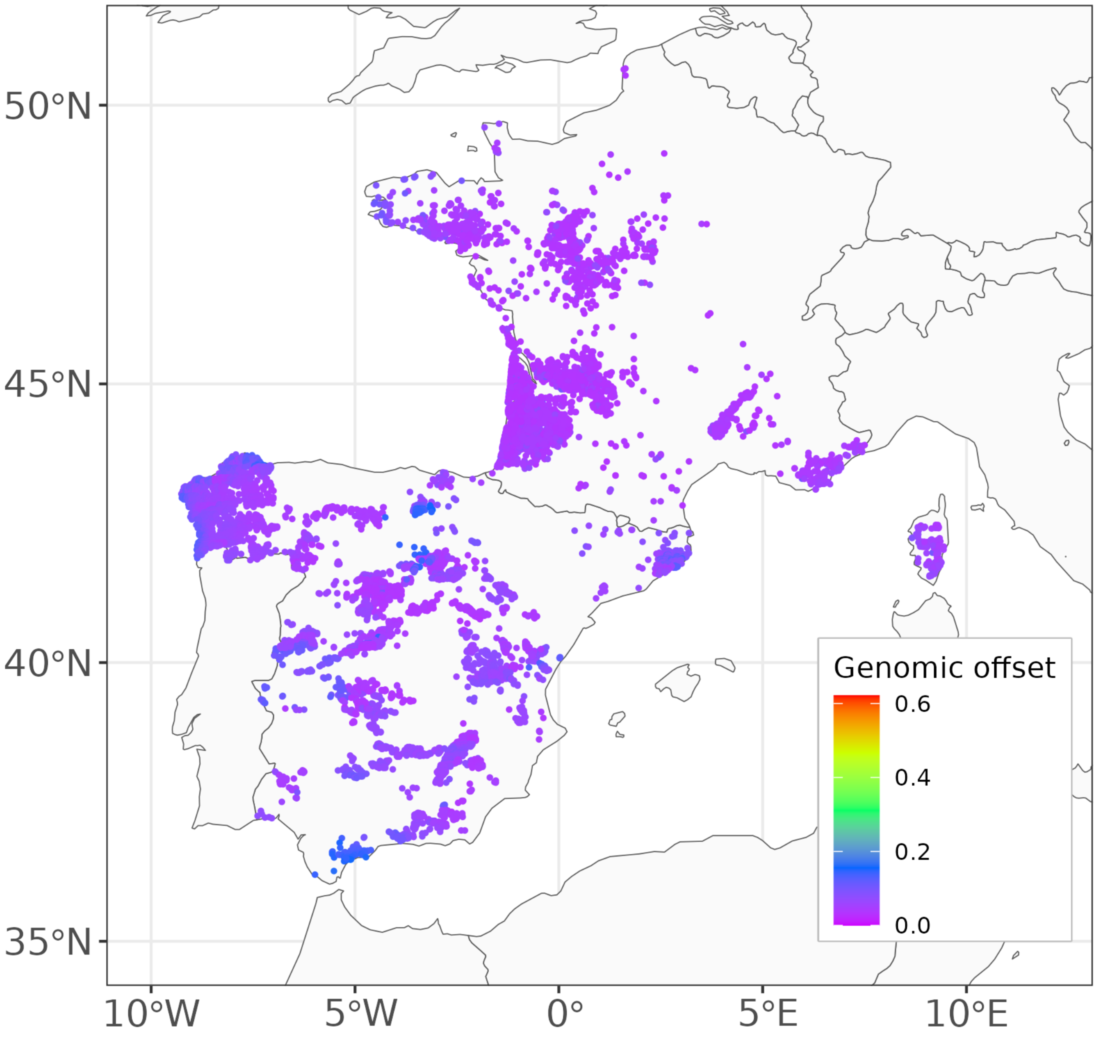

#### 8.2 Modelling approach

The goal of the present validation step was to estimate the association between GO predictions and mortality rates in natural populations, used as a proxy of the absolute fitness of the populations. A positive association between GO predictions and the mortality rates might suggest that the GO metric captures maladaptation in the face of demographic complexity (i.e. the effects of other processes on the spatial patterns in allele frequencies, such as demographic history and gene flow) as well as the genetic architecture of climate adaptation (polygenic trait architectures, G×E interactions, non-additive genetic variance).

We used mortality data from the French and Spanish National Forest Inventories (NFI) harmonized in Changenet et al. (2021). The French inventory data consists of temporary plots sampled between 2005 and 2014 while the Spanish inventory data consists of permanent plots sampled during the second (from 1986 to 1996) and third NFIs (from 1997 to 2008). A tree was recorded as dead if its death was dated at less than 5 years ago in the French NFI or if it was alive in the second inventory but dead in the third one in the Spanish NFI.

In order to account for the different census intervals between inventories, we modeled the proportion $p_i$ of maritime pines that died in the plot $i$ during the census interval with the complementary log-log link and an offset on the logarithm of the census interval $\Delta_i$ for the plot $i$, as follows:

$$
\begin{aligned}
m_i &\sim \text{Binomial}(N_i, p_i) \\
\log(-\log(1 - p_i)) &= \beta_{0,c} + \beta_C C_i + \beta_{DBH} DBH_i + \beta_{C*DBH} C_i DBH_i + \beta_{GO} GO_i + \log(\Delta_i)
\end{aligned}
$$

- $N_i$ is the total number of maritime pines in the plot $i$,
- $m_i$ is the number of maritime pines that died during the census interval $\Delta_i$ in the plot $i$,
- $C_i$ is the basal area of all tree species confounded in the plot $i$ (to account for the competition among trees) and $\beta_C$ is ts associated coefficients,
- $DBH_i$ is the mean diameter at breast height (DBH) of the maritime pines in the plot $i$ (including adults, dead trees and recruitment trees) and $\beta_{DBH}$ is its associated coefficient,
- $GO_i$ is the estimated GO in the plot $i$.
- $\beta_{0,c}$ are country-specific intercepts that account for the methodological differences between the French and Spanish inventories that may bias the estimations.

We used the following weakly informative priors:

$$
\begin{bmatrix} \beta_{0,c} \\ \beta_C \\ \beta_{GO} \\ \beta_{DBH} \\ \beta_{C*DBH} \end{bmatrix} \sim \text{Normal}(0, 1)
$$

##### 8.3 Estimated regression coefficients

Figure 69: Regression coefficients $\beta_C$ standing for the association between mortality rates and the basal area $C$ of all tree species confounded in the NFI plots (i.e., a proxy of the competition among trees). The mean and 95% credible intervals are shown.

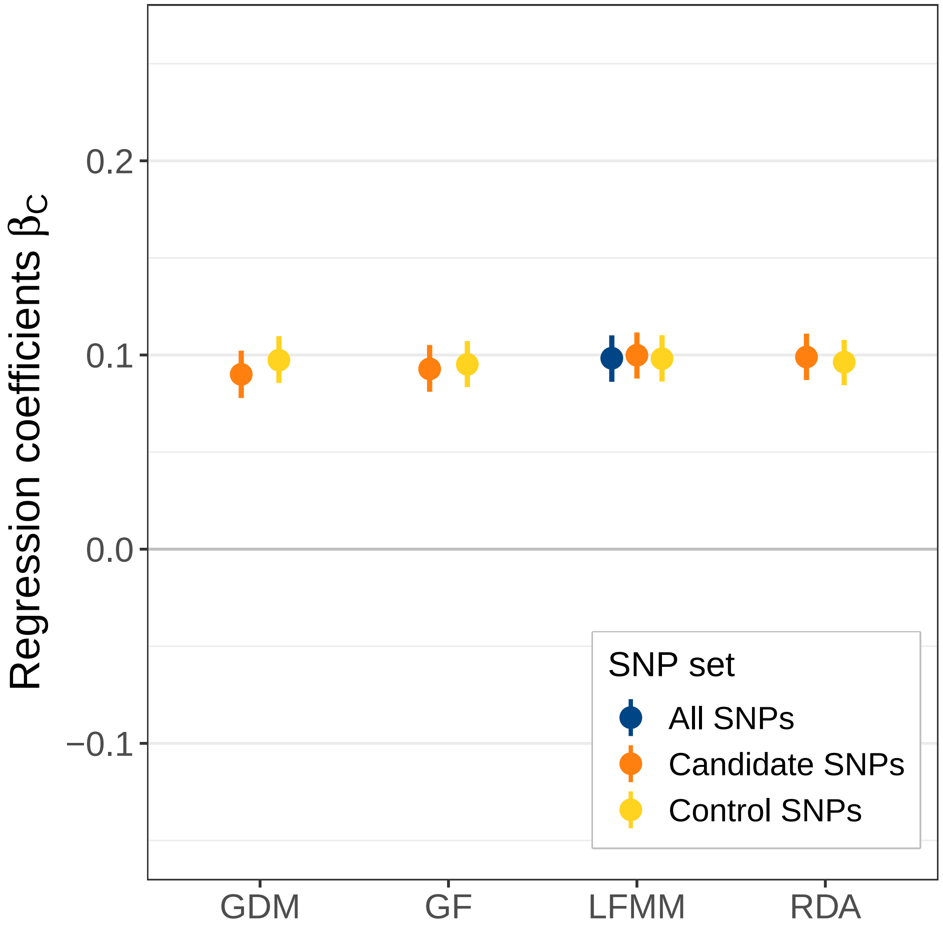

Figure 70: Regression coefficients $\beta_{DBH}$ standing for the association between mortality rates in the NFI plots and the mean diameter at breast height (DBH) of the maritime pines in the NFI plots (including adults, dead trees and recruitment trees). The mean and 95% credible intervals are shown.

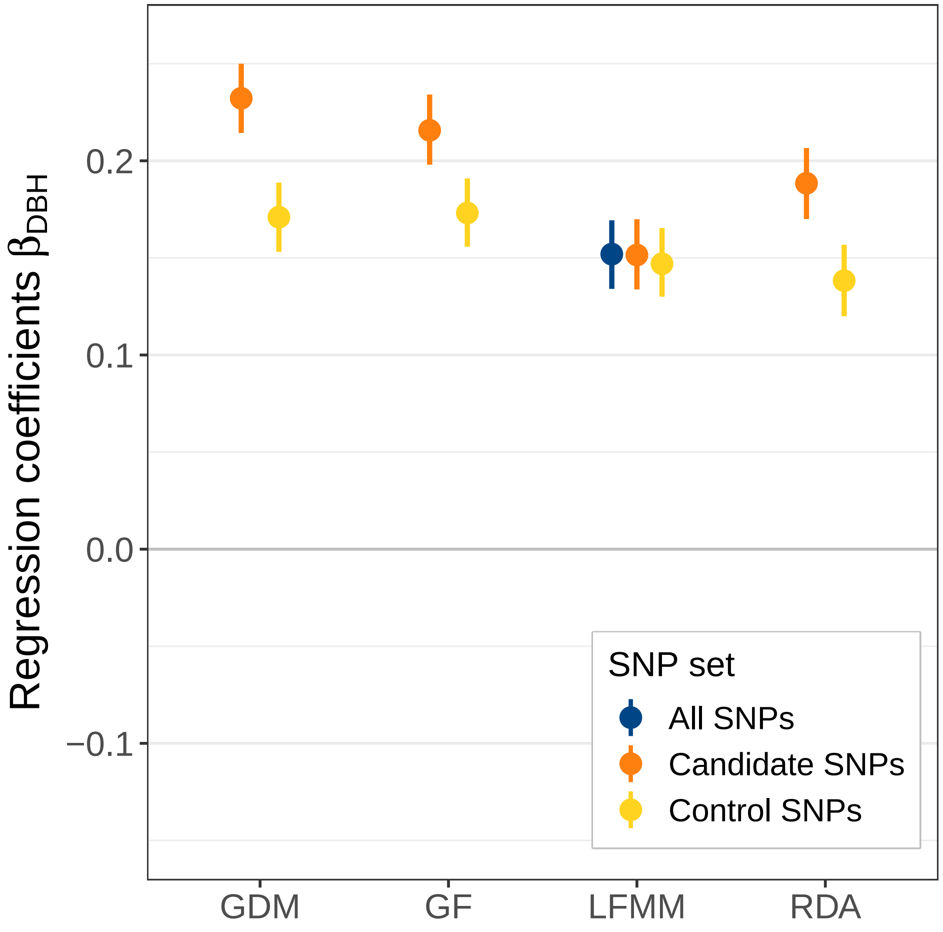

Figure 71: Regression coefficients $\beta_{C*DBH}$ standing for the association between mortality rates and the interaction between the basal area $C$ and the mean diameter at breat height $DBH$ in the NFI plots. The mean and 95% credible intervals are shown.

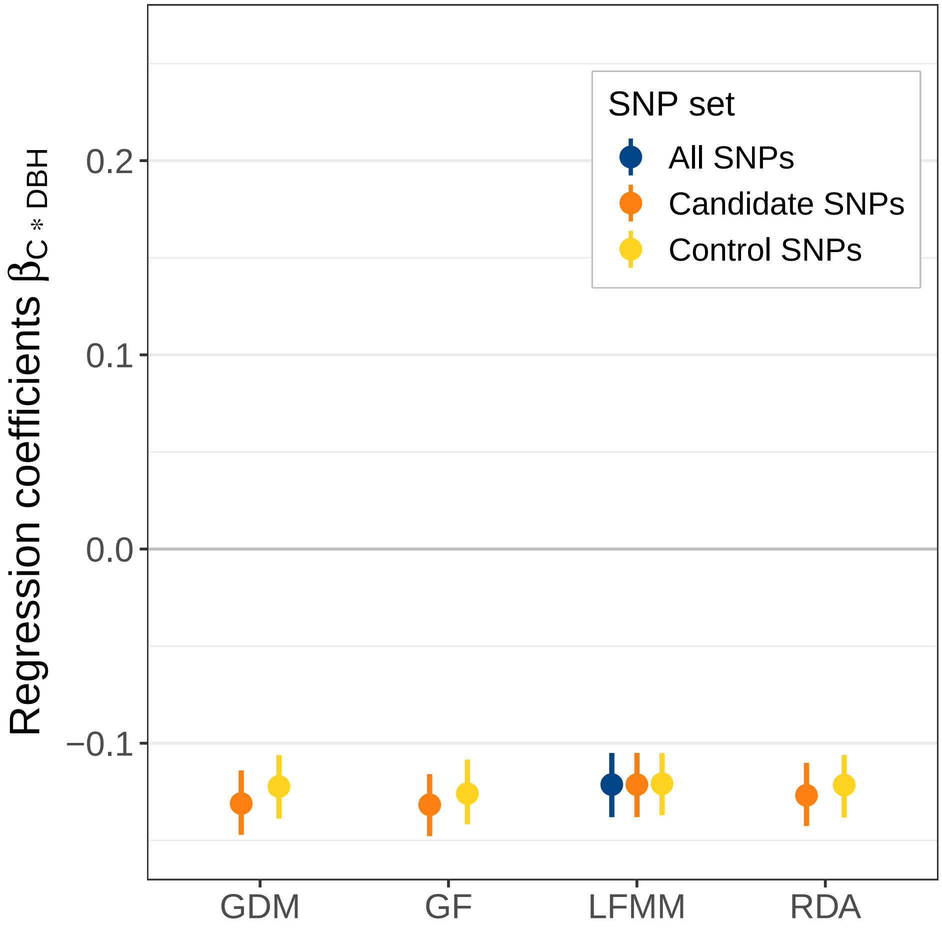

#### 8.4 Predictions of tree mortality probability

The plots below show the estimated association between tree mortality probability $p$ in the NFI plots and GO predictions. Tree mortality probability was predicted with the basal area $C$ and the diameter at breast height $DBH$ fixed to their mean value. The line corresponds to the mean estimate of the probability of tree mortality $p$ and the uncertainty interval corresponds to the estimated 90% credible interval (highest posterior density interval).

Figure 72: Predicted probability of tree mortality as a function of GO predictions in the NFI plots.

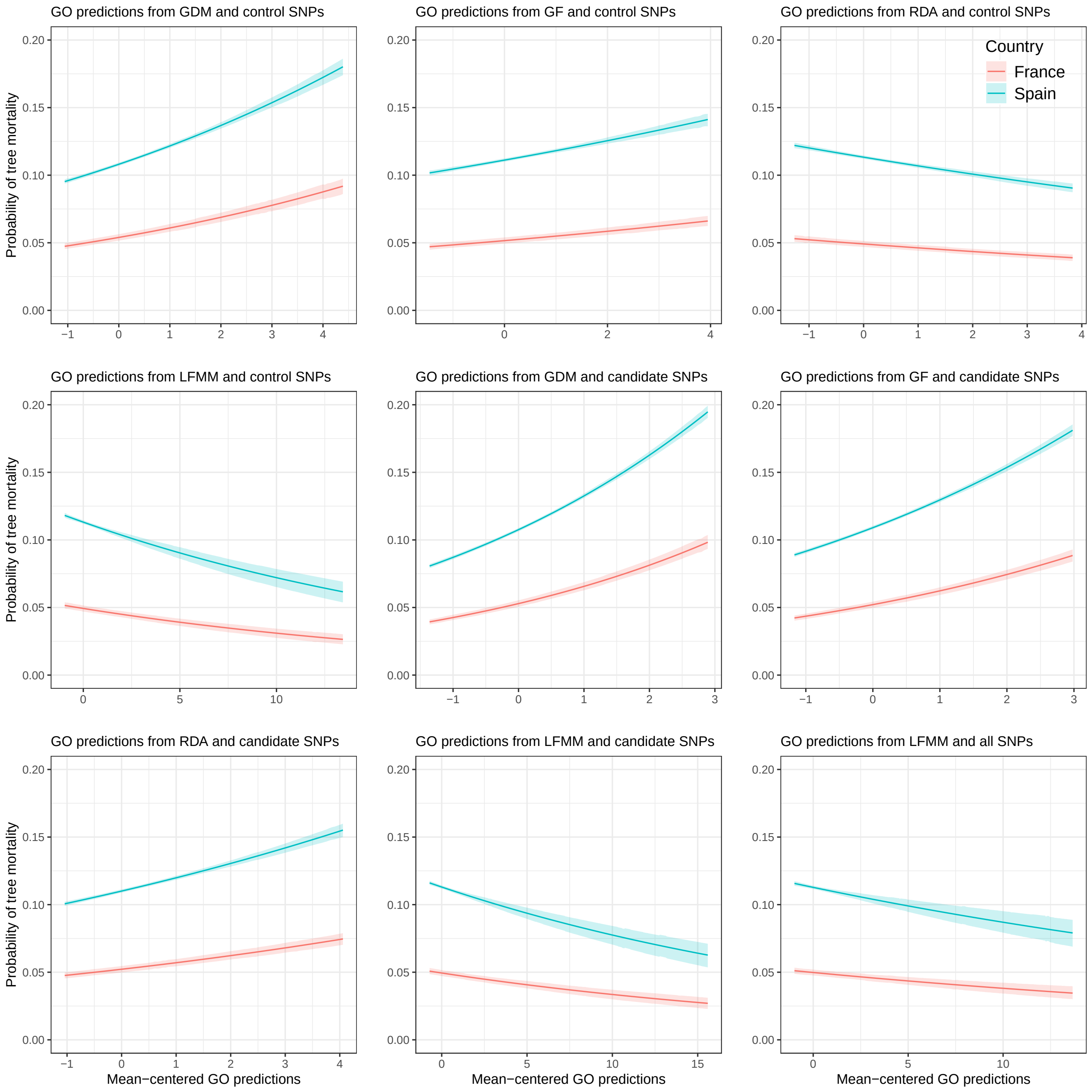

The table below provides:

- the predictions of tree mortality probability $p$ for different values of GO predictions ($GO$={-1, 0, 1}), with $C$ and $DBH$ fixed to their mean value.

- the percent change of $p$ for a one-unit increase ($+\Delta_p$) or decrease ($-\Delta_p$) in $GO$ (which corresponds to a one-standard deviation increase/decrease as $GO$ is mean-centered).

Table 7: Predictions of tree mortality probability in the NFI plots.

| Method | Country | p(GO=-1) | p(GO=0) | p(GO=1) | +Δp | -Δp |
|---|---|---|---|---|---|---|
| GO predictions from GDM and control SNPs | Spain | 0.096 [0.094;0.098] | 0.108 [0.107;0.109] | 0.122 [0.12;0.123] | 12.963% | -11.111% |
| GO predictions from GDM and control SNPs | France | 0.048 [0.046;0.05] | 0.054 [0.052;0.056] | 0.061 [0.058;0.064] | 12.963% | -11.111% |
| GO predictions from GF and control SNPs | Spain | 0.105 [0.103;0.106] | 0.111 [0.11;0.112] | 0.118 [0.117;0.12] | 6.306% | -5.405% |
| GO predictions from GF and control SNPs | France | 0.048 [0.046;0.051] | 0.052 [0.049;0.054] | 0.055 [0.052;0.057] | 5.769% | -7.692% |
| GO predictions from RDA and control SNPs | Spain | 0.12 [0.118;0.122] | 0.113 [0.112;0.115] | 0.107 [0.105;0.108] | -5.31% | 6.195% |
| GO predictions from RDA and control SNPs | France | 0.052 [0.05;0.055] | 0.049 [0.047;0.051] | 0.046 [0.044;0.048] | -6.122% | 6.122% |
| GO predictions from LFMM and control SNPs | Spain | 0.118 [0.117;0.12] | 0.113 [0.112;0.115] | 0.108 [0.107;0.11] | -4.425% | 4.425% |
| GO predictions from LFMM and control SNPs | France | 0.052 [0.049;0.054] | 0.049 [0.047;0.051] | 0.047 [0.045;0.049] | -4.082% | 6.122% |
| GO predictions from GDM and candidate SNPs | Spain | 0.087 [0.086;0.089] | 0.108 [0.106;0.109] | 0.133 [0.131;0.134] | 23.148% | -19.444% |
| GO predictions from GDM and candidate SNPs | France | 0.043 [0.041;0.045] | 0.053 [0.051;0.055] | 0.066 [0.063;0.069] | 24.528% | -18.868% |
| GO predictions from GF and candidate SNPs | Spain | 0.092 [0.09;0.093] | 0.109 [0.108;0.11] | 0.13 [0.128;0.131] | 19.266% | -15.596% |
| GO predictions from GF and candidate SNPs | France | 0.044 [0.042;0.045] | 0.052 [0.05;0.054] | 0.062 [0.059;0.065] | 19.231% | -15.385% |
| GO predictions from RDA and candidate SNPs | Spain | 0.101 [0.099;0.103] | 0.11 [0.109;0.111] | 0.12 [0.118;0.121] | 9.091% | -8.182% |
| GO predictions from RDA and candidate SNPs | France | 0.048 [0.046;0.05] | 0.052 [0.05;0.055] | 0.057 [0.054;0.06] | 9.615% | -7.692% |
| GO predictions from LFMM and candidate SNPs | Spain | 0.117 [0.116;0.119] | 0.113 [0.112;0.114] | 0.109 [0.107;0.11] | -3.54% | 3.54% |
| GO predictions from LFMM and candidate SNPs | France | 0.051 [0.049;0.054] | 0.049 [0.047;0.052] | 0.048 [0.045;0.05] | -2.041% | 4.082% |
| GO predictions from LFMM and all SNPs | Spain | 0.116 [0.114;0.117] | 0.113 [0.112;0.114] | 0.11 [0.108;0.111] | -2.655% | 2.655% |
| GO predictions from LFMM and all SNPs | France | 0.051 [0.049;0.053] | 0.05 [0.048;0.052] | 0.048 [0.046;0.051] | -4% | 2% |

#### 8.5 Evaluating the probability of obtaining such results by chance

We evaluated the probability that the estimated relationship between the mortality rates and GO predictions could be obtained by chance. For that, we ran again the model on the NFI data but we replaced GO predictions by a randomly generated variable $X$ such as $X \sim \mathcal{N}(0,1)$. We repeated this step 100 times and obtained a distribution of $\beta_X$ coefficients capturing the relationship between mortality rates and a randomly generated variable. We then compared this distribution of $\beta_X$ coefficients to the estimated $\beta_{GO}$ coefficients capturing the relationship between the mortality rates and the GO predictions.

Figure 73: Estimated mean and 95% credible intervals of the $\beta_X$ coefficients capturing the relationship between the mortality rates and the GO predictions (predicted GO in the legend) and the $\beta_X$ coefficients capturing the relationship between the mortality rates and a randomly generated variable (random GO in the legend).

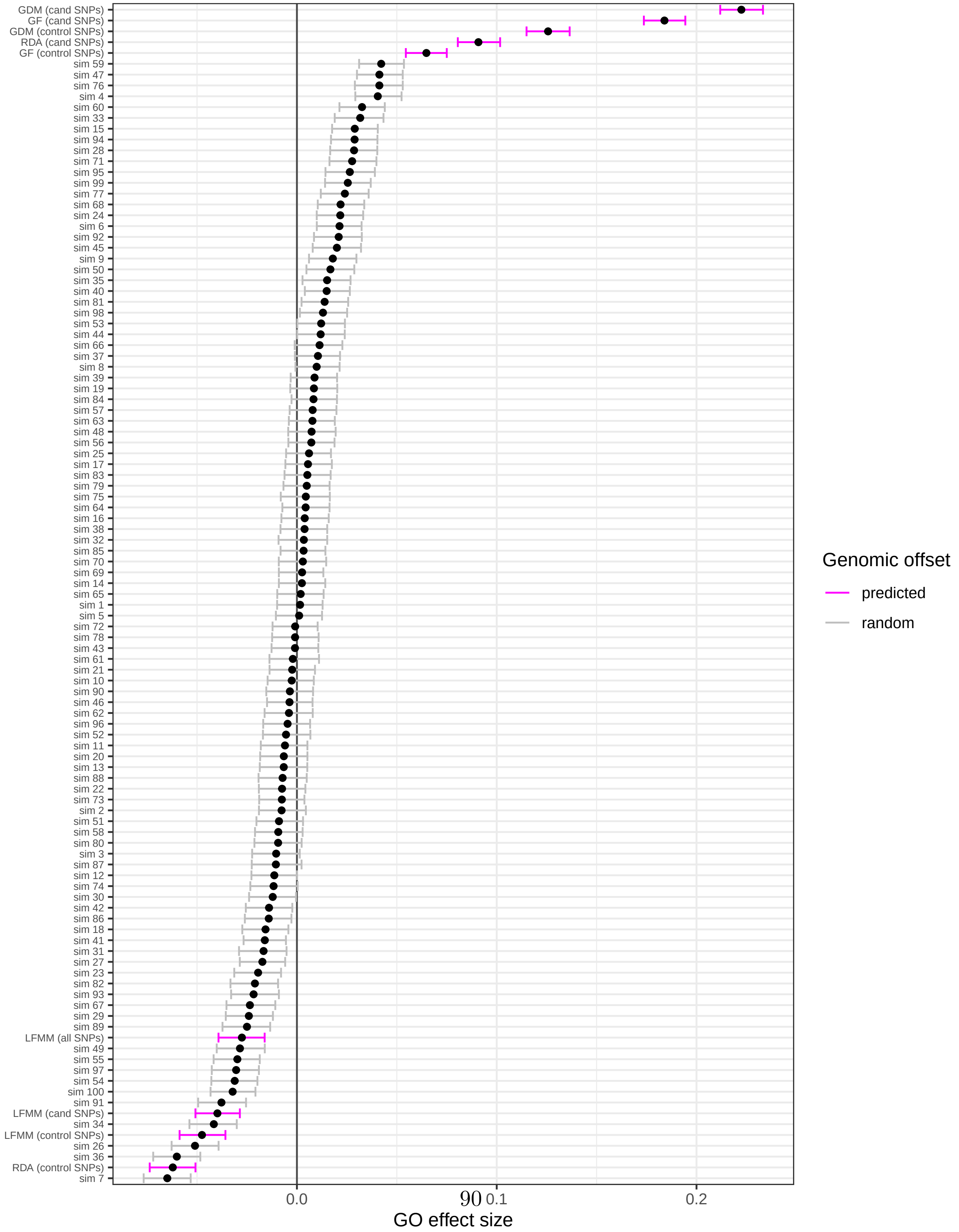

### 9 Validation in the common gardens

#### 9.1 Climate in the common gardens

Table 8: Climate in the five common gardens between the planting and the measurement dates.

| bio1 | bio12 | bio15 | bio3 | bio4 | SHM |
|---|---|---|---|---|---|
| 12.05 | 1148.88 | 43.63 | 40.92 | 406.60 | 69.65 |
| 14.15 | 912.60 | 28.52 | 37.14 | 556.98 | 73.84 |
| 16.62 | 669.00 | 82.03 | 35.95 | 716.42 | 290.50 |
| 14.55 | 460.05 | 47.42 | 36.21 | 754.34 | 217.82 |
| 12.47 | 1272.10 | 66.53 | 38.01 | 611.70 | 122.52 |

See Table S3 for the full names and units of the climatic variables.

#### 9.2 GO predictions

##### 9.2.1 Methodology

To predict the GO at the location of the common gardens, we used:

- the average climates under the time period 1901-1950, which are expected to represent the past climates under which the populations have evolved. The time period 1901-1950 is the same reference period as the one used to predict the GO at the location of the populations and the NFI plots.
- the average climates between the planting date and the measurement date (i.e. height and mortality measurements) at the location of each common garden.

Time period between the planting date and the measurement date in each common garden:

- in Asturias (Spain), trees were planted in February 2011 and measured in March 2014 when the trees were 37 month-old.
- in Bordeaux (France), trees were planted in October 2011 and measured in November 2018 when the trees were 85 month-old.
- in Cáceres (Spain) trees were planted in April 2011 and measured in December 2011 when the trees were 8 month-old. Note that for this common garden, we calculated the climatic variables for the entire year 2011 (instead of calculating the variables only for months between April and December). Indeed, the calculation of the annual climatic variables would be wrong if we do not account for some months, e.g. the mean annual temperature will the overestimated because we do not account for some of the winter months.
- in Madrid (Spain), trees were planted in November 2010 and measured in December 2011 when the trees were 13 month-old.

- in Fundão (Portugal), trees were planted in February 2011 and measured in May 2013 when the trees were 27 month-old.

##### 9.2.2 Mapping GO predictions

In the graphs below, the points correspond the sampled populations of maritime pine and the black asterisk corresponds to the location of the common garden for which the GO was predicted.

###### 9.2.2.1 GO predictions with RDA

Figure 74: GO predictions with RDA in the common garden of Asturias, Spain (black asterisk).

(a) Candidate SNPs.

(b) Control SNPs.

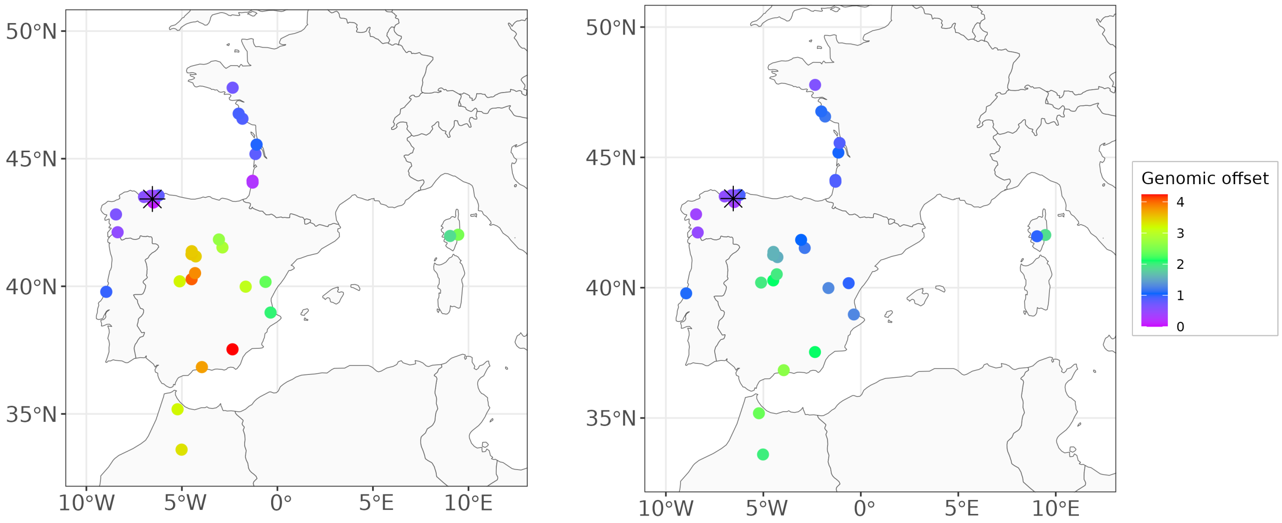

Figure 75: GO predictions with RDA in the common garden of Bordeaux, France (black asterisk).

(a) Candidate SNPs.

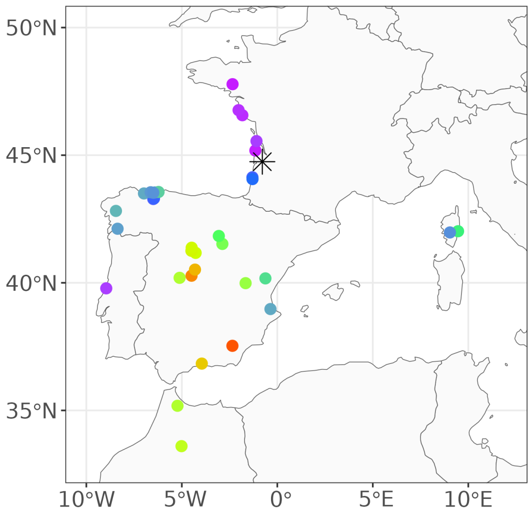

(b) Control SNPs.

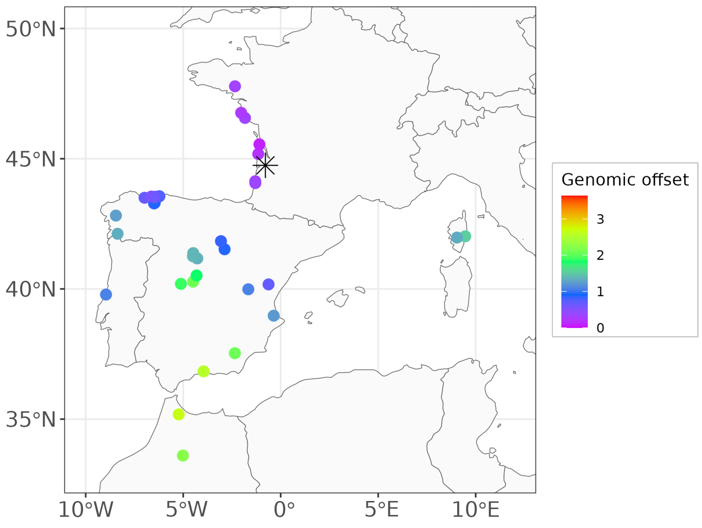

Figure 76: GO predictions with RDA in the common garden of Cáceres, Spain (black asterisk).

(a) Candidate SNPs.

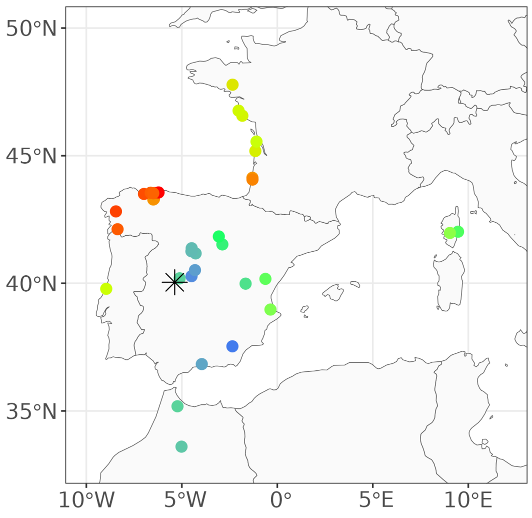

(b) Control SNPs.

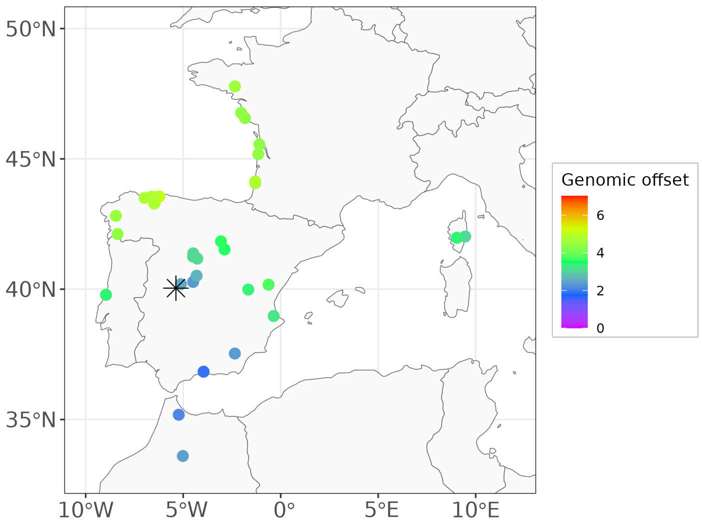

Figure 77: GO predictions with RDA in the common garden of Madrid, Spain (black asterisk).

(a) Candidate SNPs.

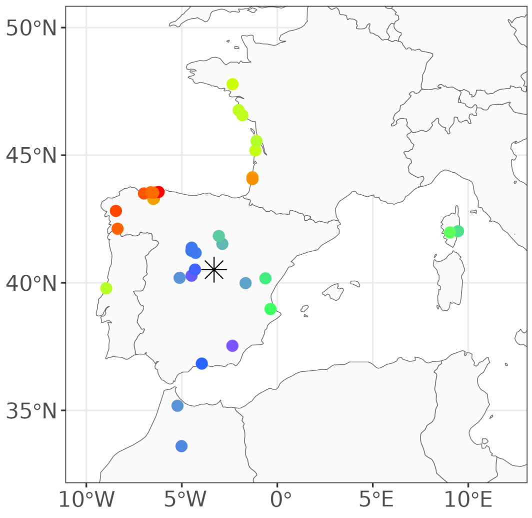

(b) Control SNPs.

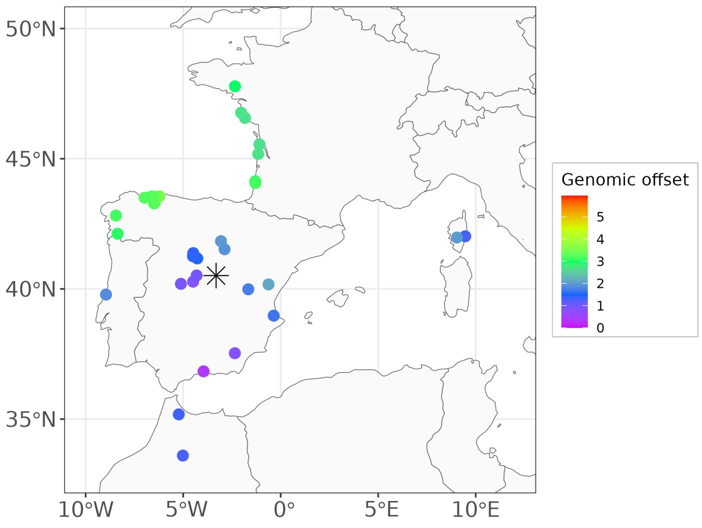

Figure 78: GO predictions with RDA in the common garden of Fundão, Portugal (black asterisk).

(a) Candidate SNPs.

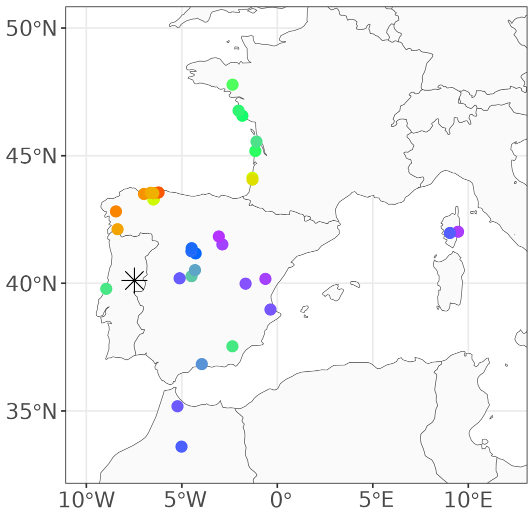

(b) Control SNPs.

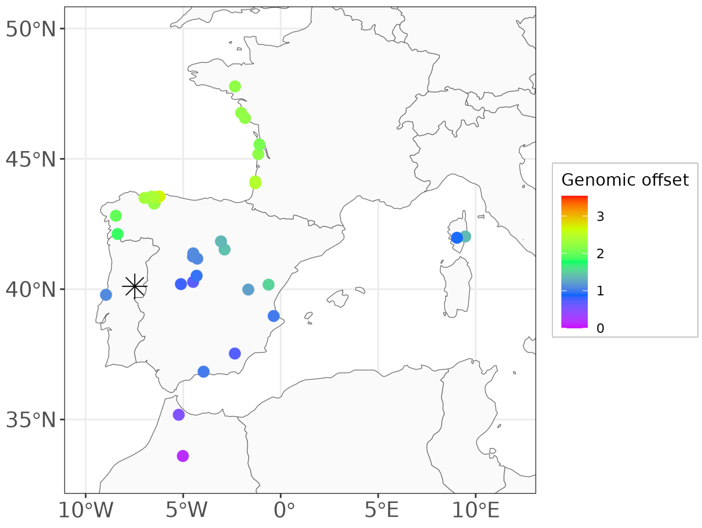

###### 9.2.2.2 GO predictions with GF

Figure 79: GO predictions with GF in the common garden of Asturias, Spain (black asterisk).

(a) Candidate SNPs. (b) Control SNPs.

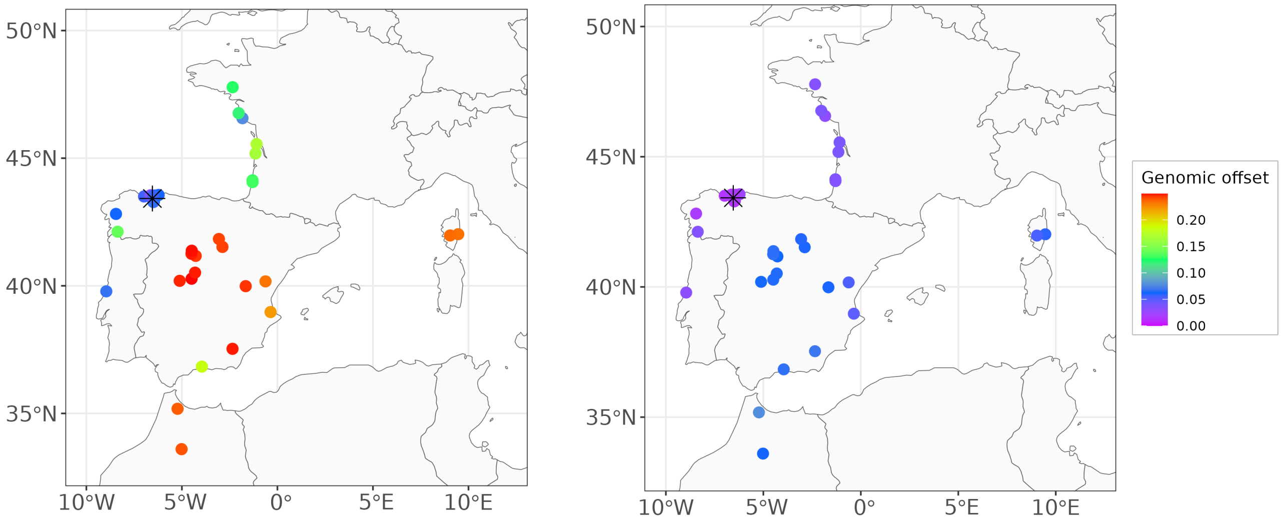

Figure 80: GO predictions with GF in the common garden of Bordeaux, France (black asterisk).

(a) Candidate SNPs. (b) Control SNPs.

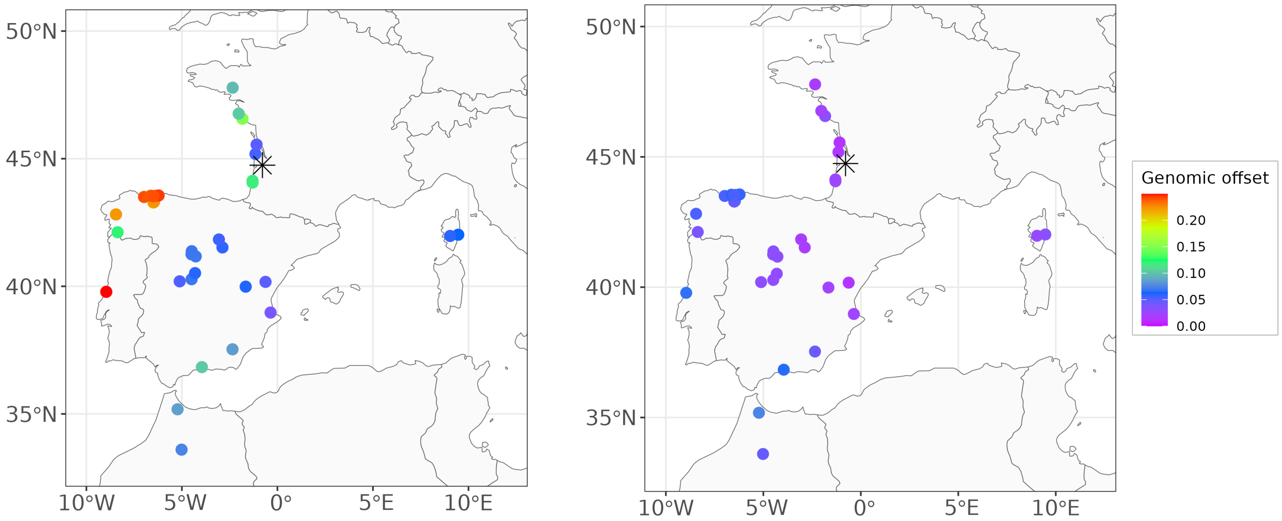

Figure 81: GO predictions with GF in the common garden of Cáceres, Spain (black asterisk).

(a) Candidate SNPs. (b) Control SNPs.

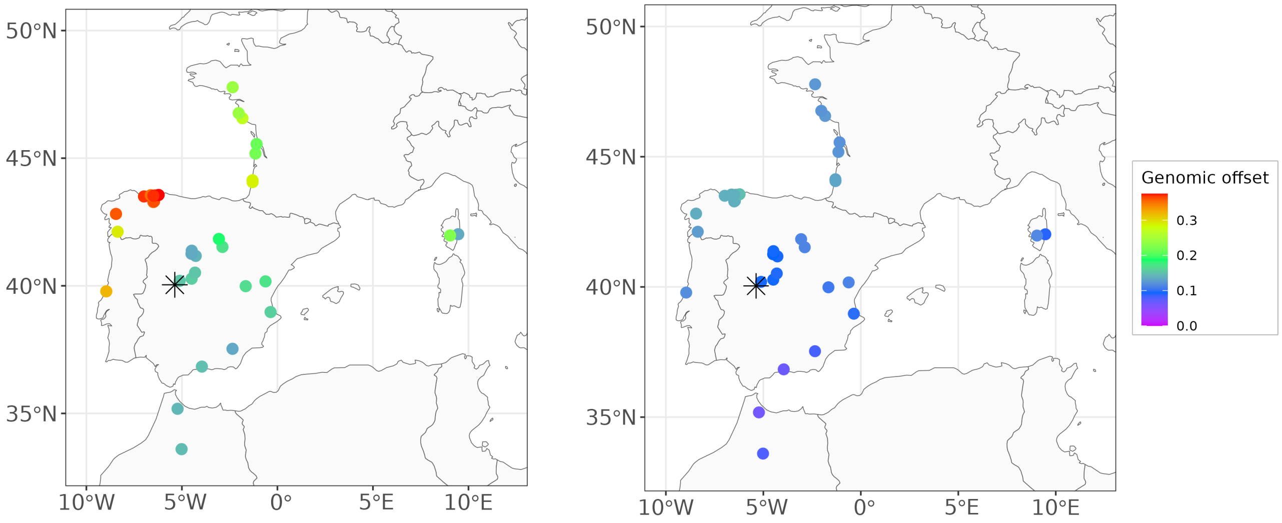

Figure 82: GO predictions with GF in the common garden of Madrid, Spain (black asterisk).

(a) Candidate SNPs. (b) Control SNPs.

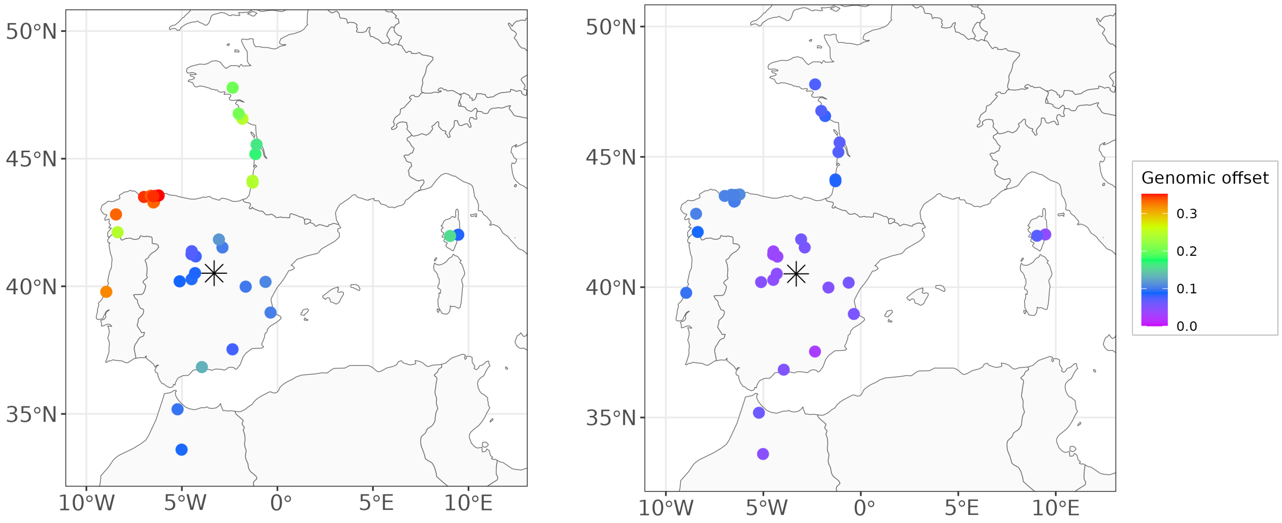

Figure 83: GO predictions with GF in the common garden of Fundão, Portugal (black asterisk).

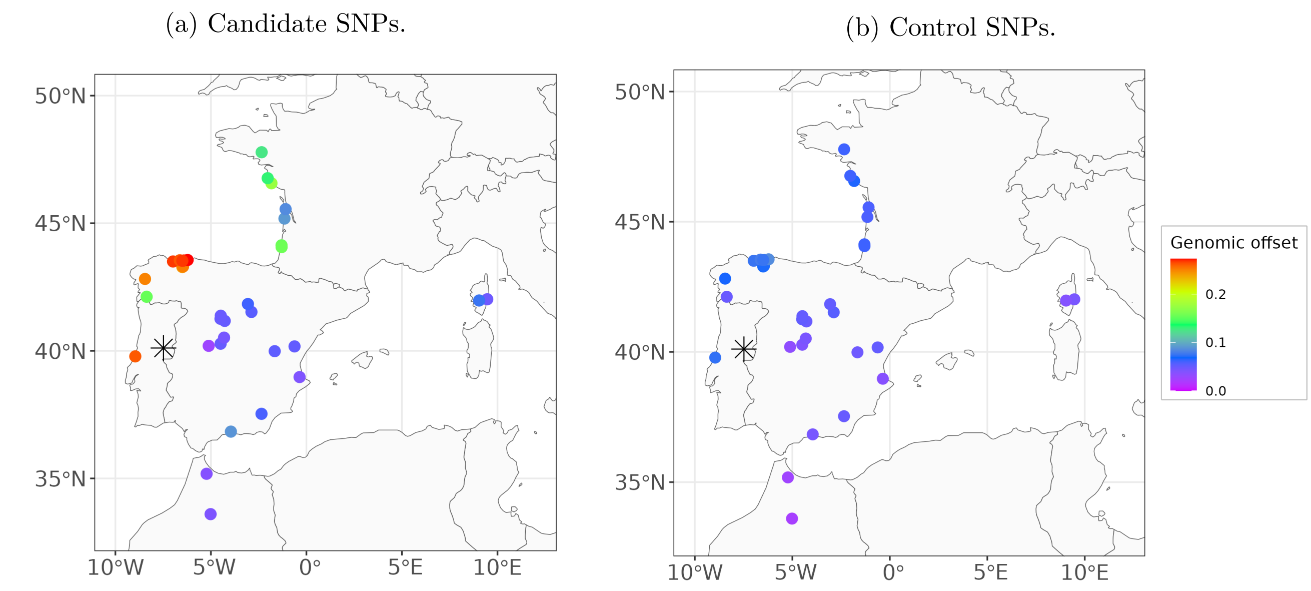

(a) Candidate SNPs.

(b) Control SNPs.

##### 9.2.2.3 GO predictions with LFMM

Figure 84: GO predictions with LFMM and all SNPs for the five common gardens.

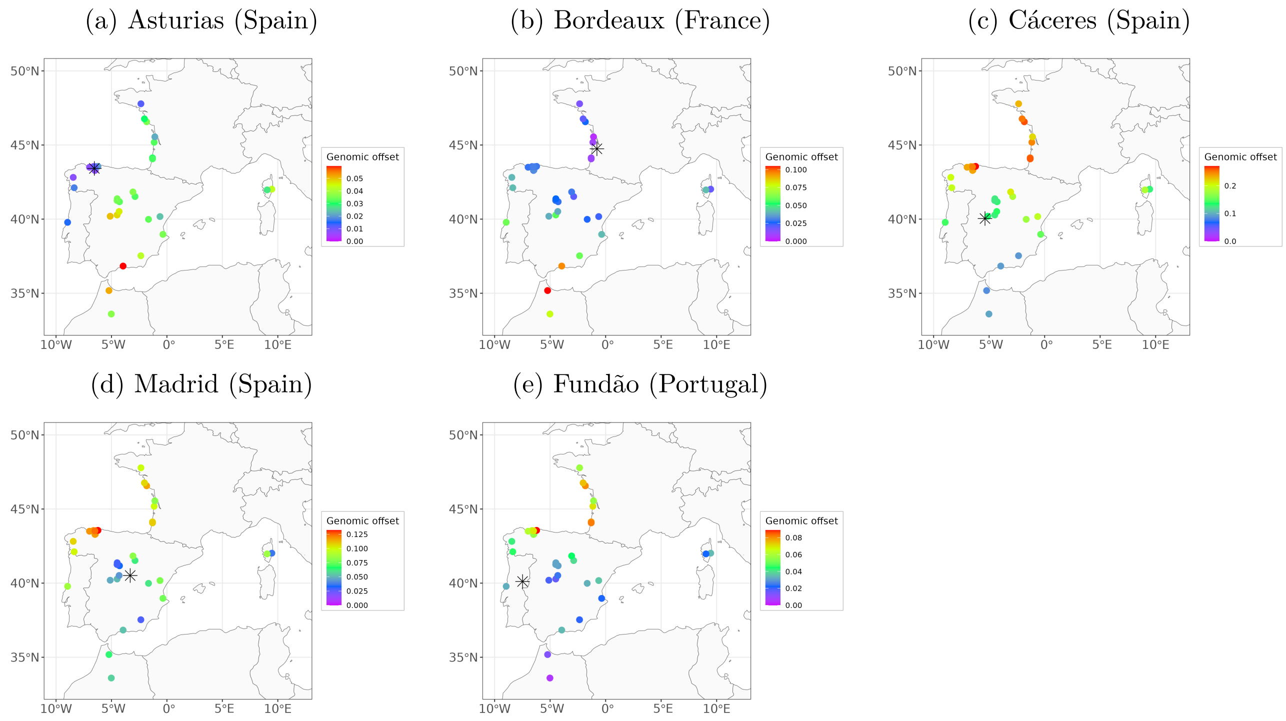

(a) Asturias (Spain)

(b) Bordeaux (France)

(c) Cáceres (Spain)

(d) Madrid (Spain)

(e) Fundão (Portugal)

Figure 85: GO predictions with LFMM in the common garden in Asturias, Spain (black asterisk).

(a) Candidate SNPs. (b) Control SNPs.

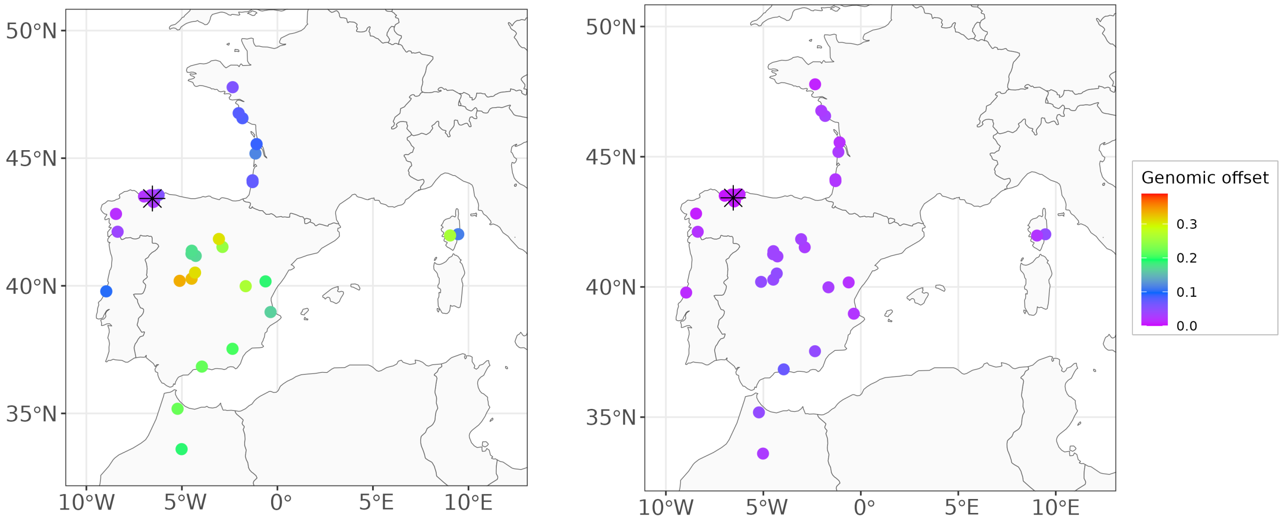

Figure 86: GO predictions with LFMM in the common garden of Bordeaux, France (black asterisk).

(a) Candidate SNPs. (b) Control SNPs.

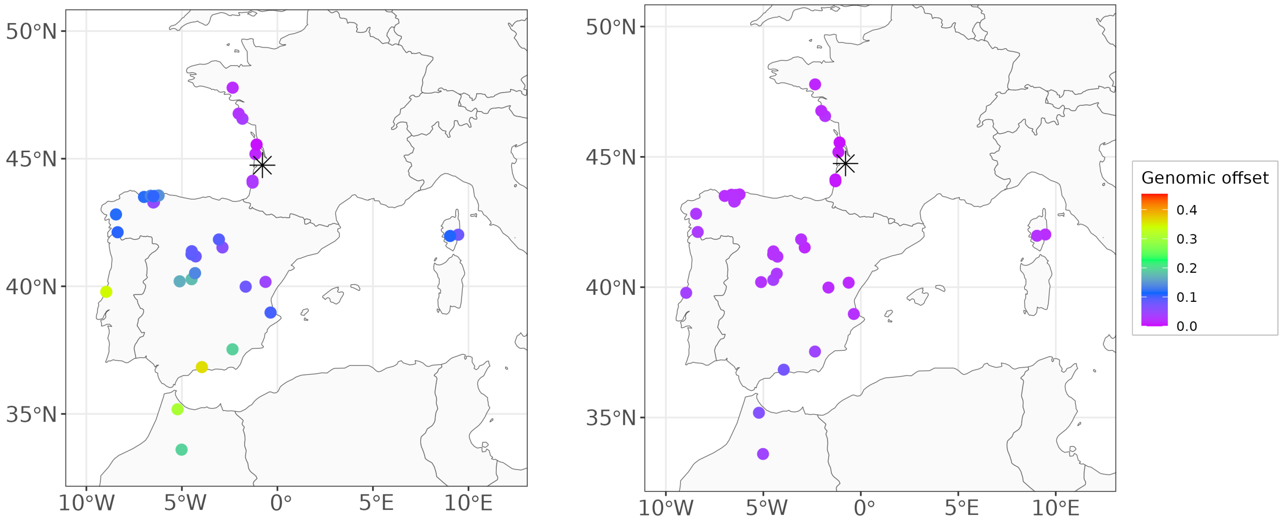

Figure 87: GO predictions with LFMM in the common garden of Cáceres, Spain (black asterisk).

(a) Candidate SNPs.

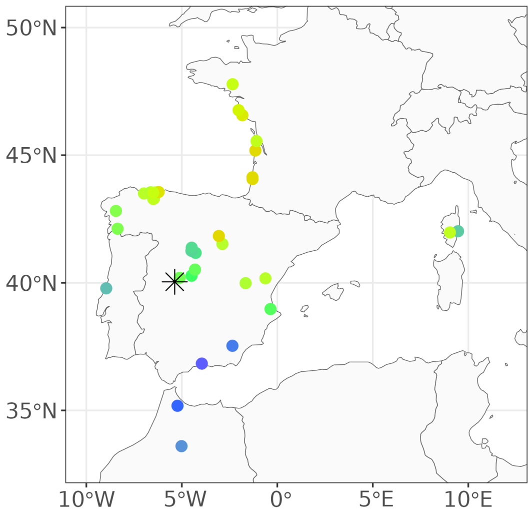

(b) Control SNPs.

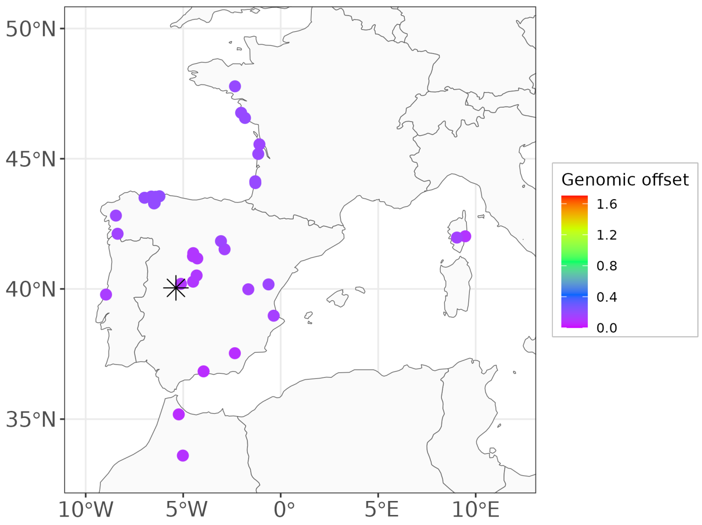

Figure 88: GO predictions with LFMM in the common garden of Madrid, Spain (black asterisk).

(a) Candidate SNPs.

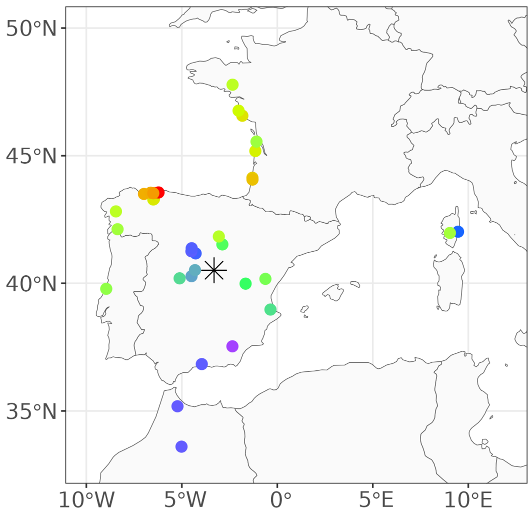

(b) Control SNPs.

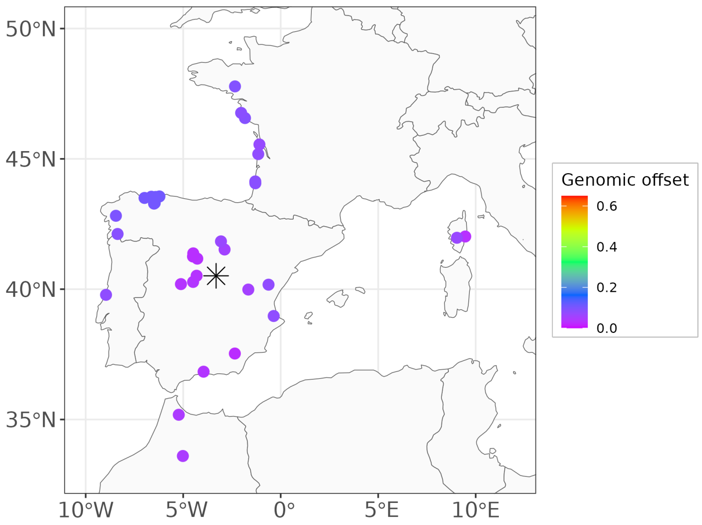

Figure 89: GO predictions with LFMM in the common garden of Fundão, Portugal (black asterisk).

(a) Candidate SNPs. (b) Control SNPs.

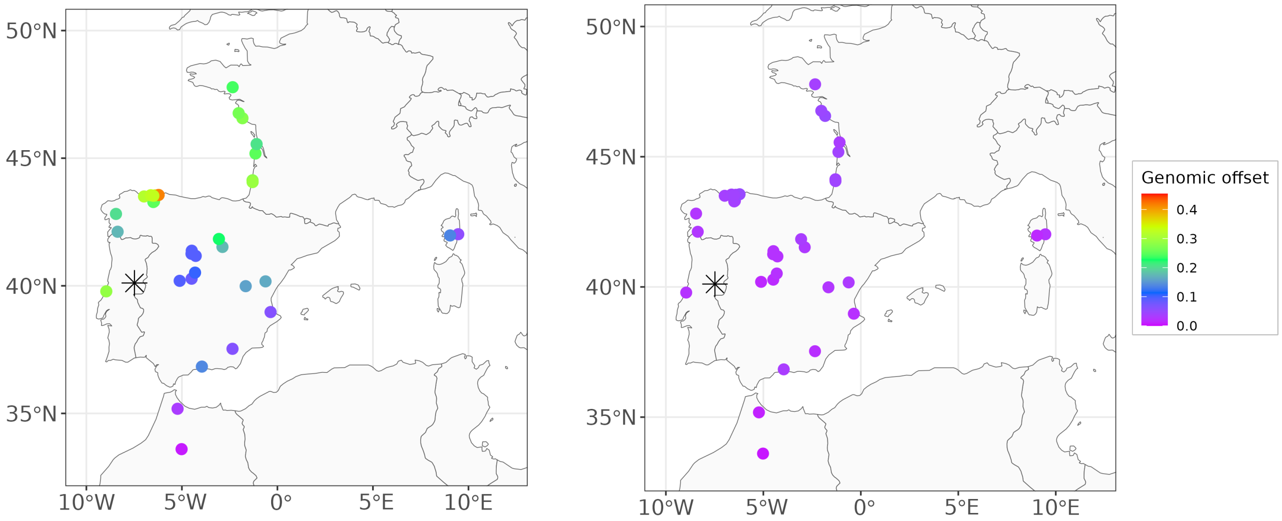

##### 9.2.2.4 GO predictions with GDM

Figure 90: GO predictions with GDM in the common garden of Asturias, Spain (black asterisk).

(a) Candidate SNPs. (b) Control SNPs.

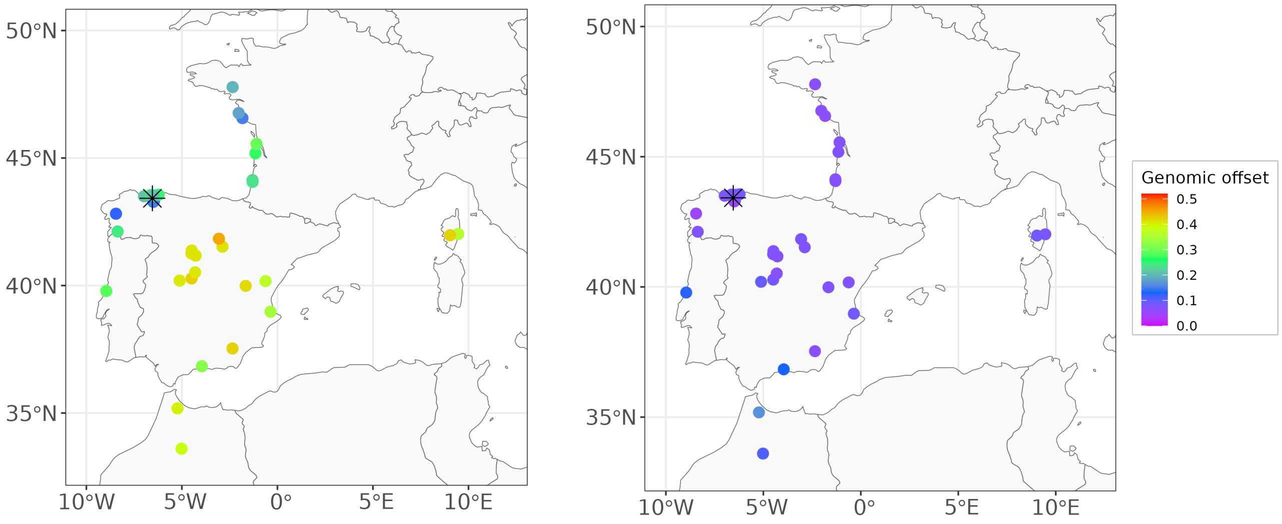

Figure 91: GO predictions with GDM in the common garden of Bordeaux, France (black asterisk).

(a) Candidate SNPs.

(b) Control SNPs.

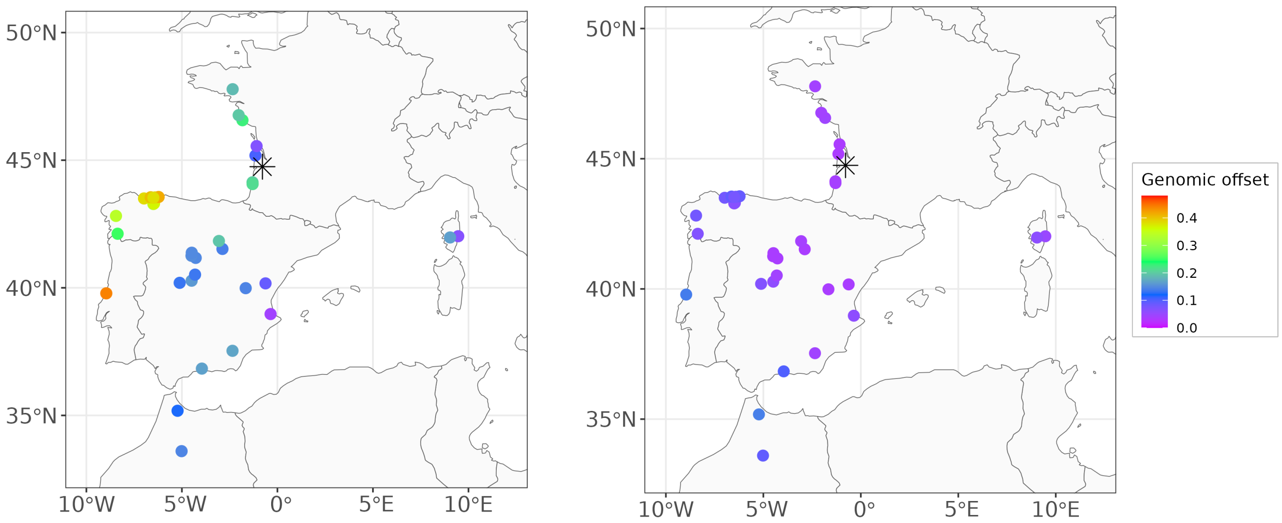

Figure 92: GO predictions with GDM in the common garden of Cáceres, Spain (black asterisk).

(a) Candidate SNPs.

(b) Control SNPs.

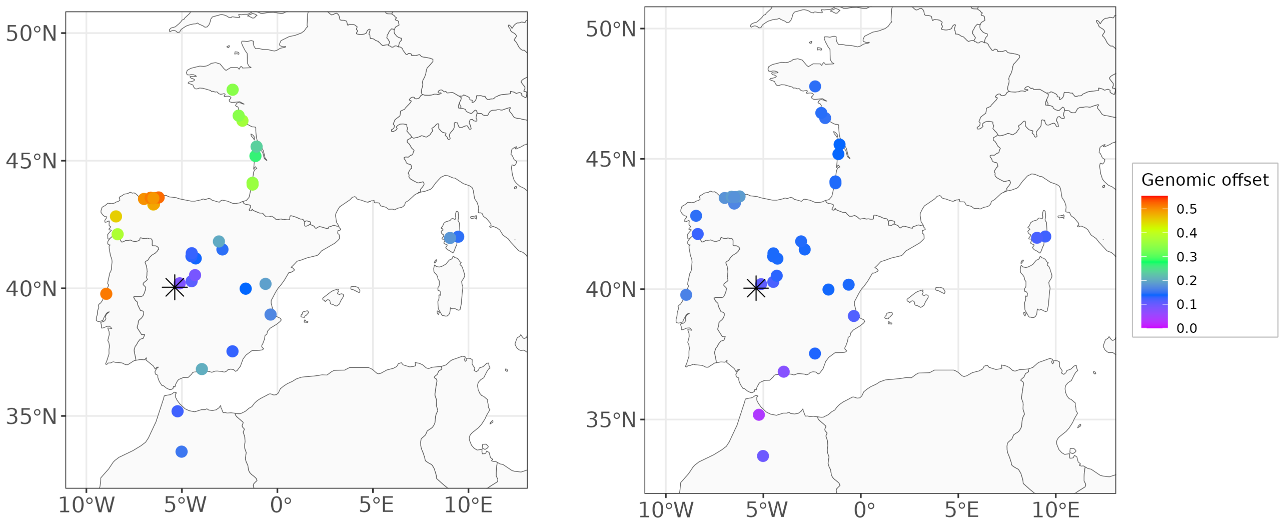

Figure 93: GO predictions with GDM in the common garden of Madrid, Spain (black asterisk).

(a) Candidate SNPs.

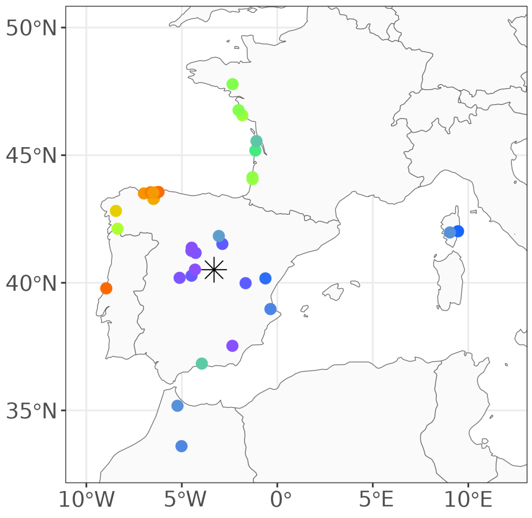

(b) Control SNPs.

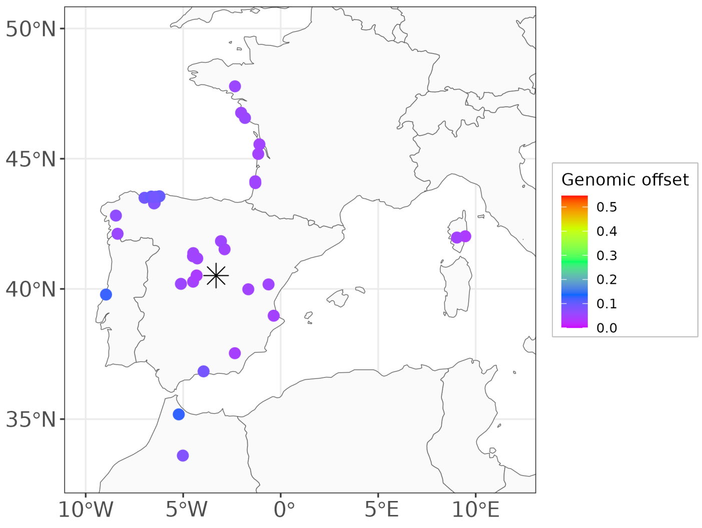

Figure 94: GO predictions with GDM in the common garden of Fundão, Portugal (black asterisk).

(a) Candidate SNPs.

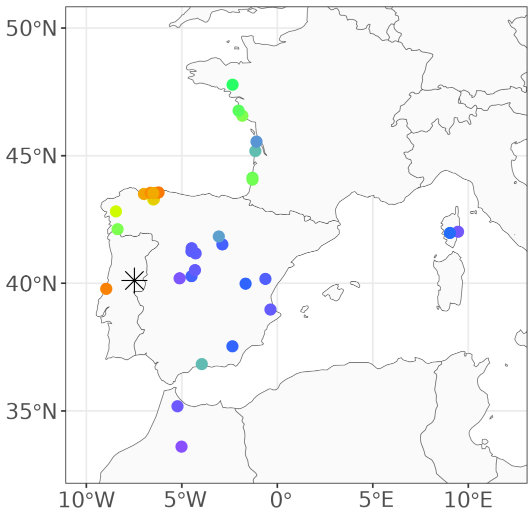

(b) Control SNPs.

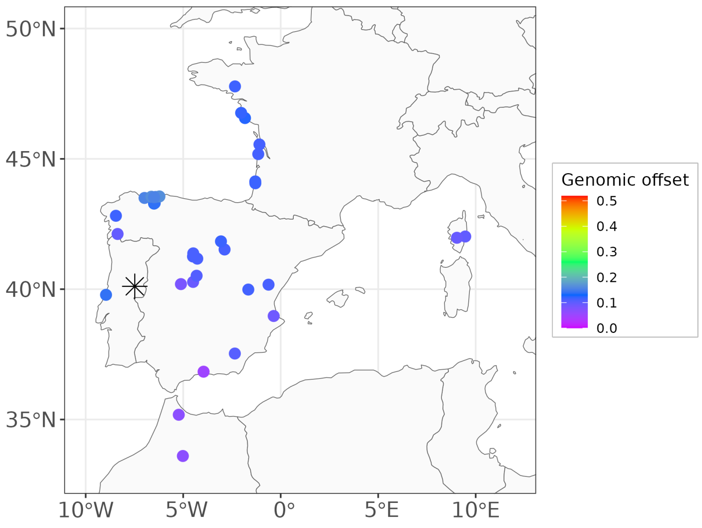

##### 9.2.3 Correlations between GO predictions and CTDs

Figure 95: Correlations between GO predictions and CTDs in Asturias, Spain.

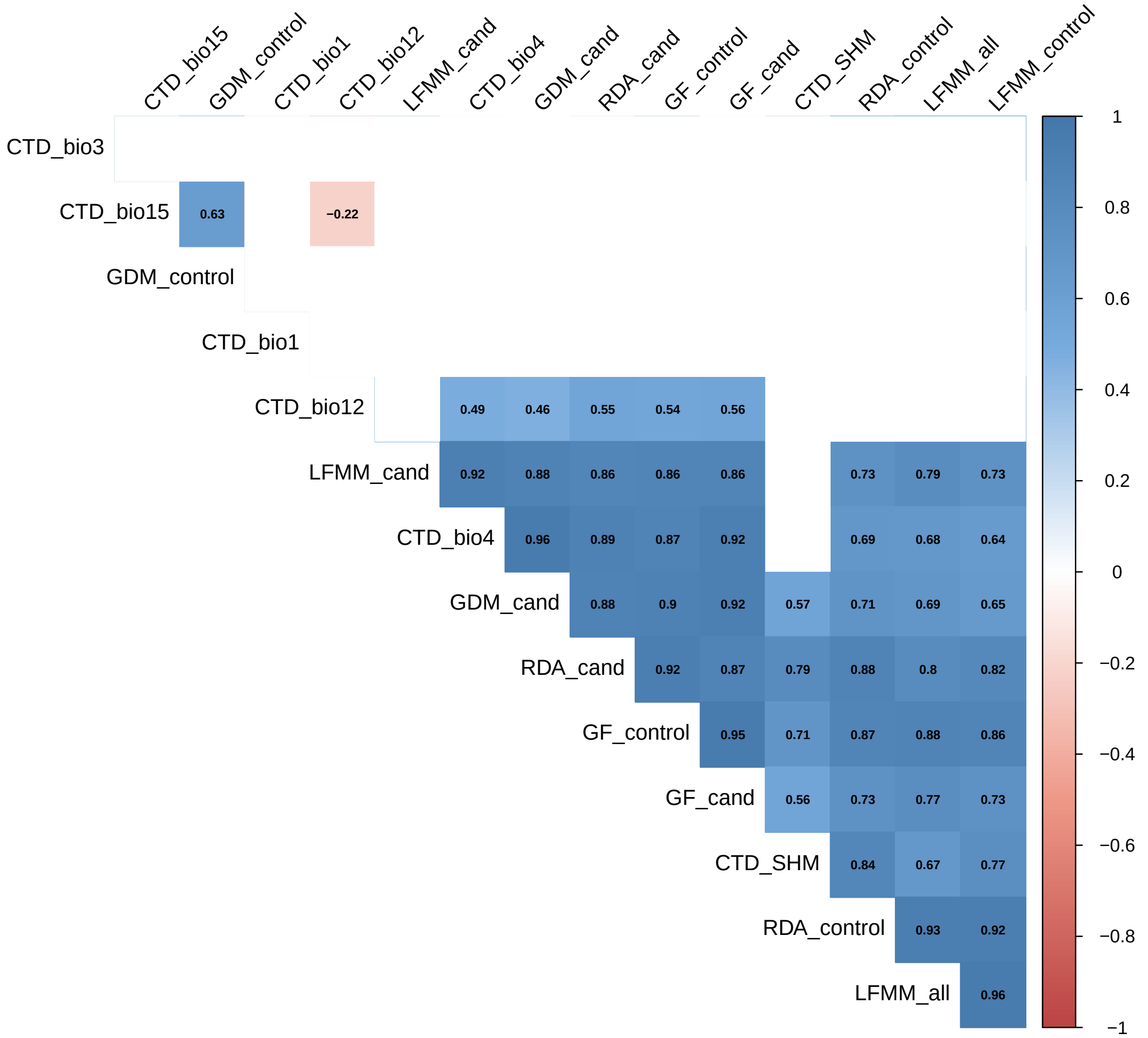

Figure 96: Correlations between GO predictions and CTDs in Bordeaux, France.

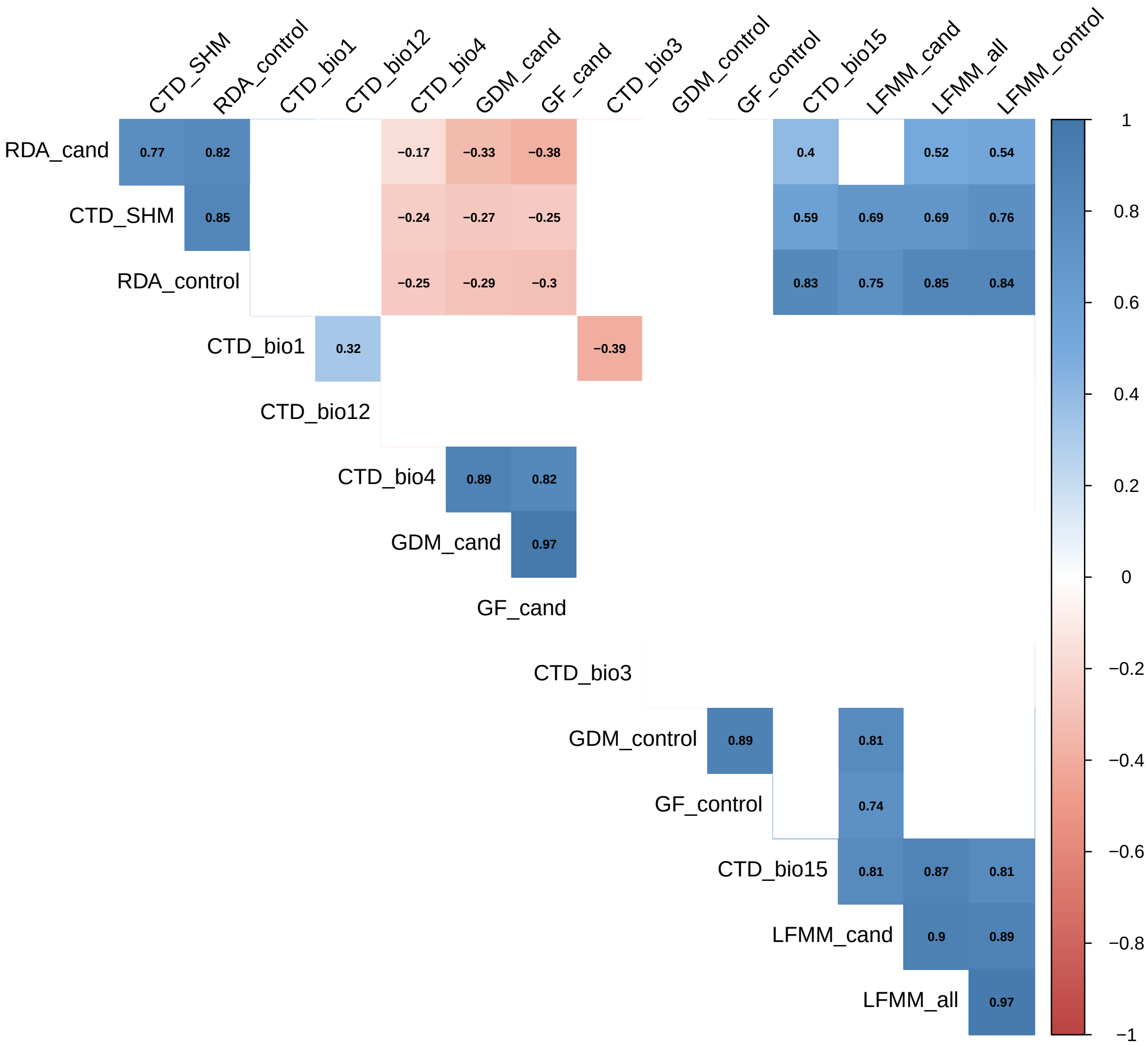

Figure 97: Correlations between GO predictions and CTDs in Cáceres, Spain.

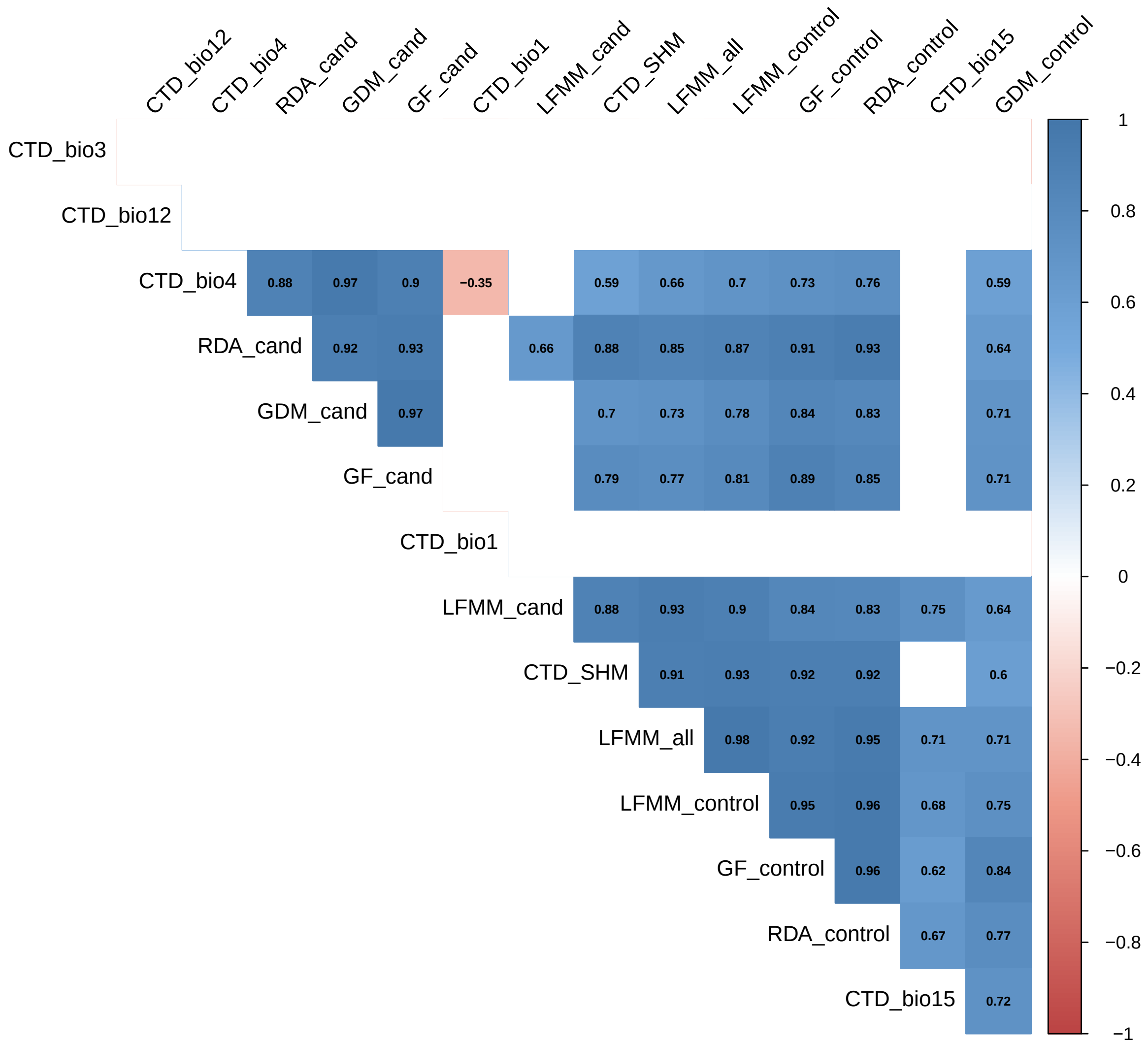

Figure 98: Correlations between GO predictions and CTDs in Madrid, Spain.

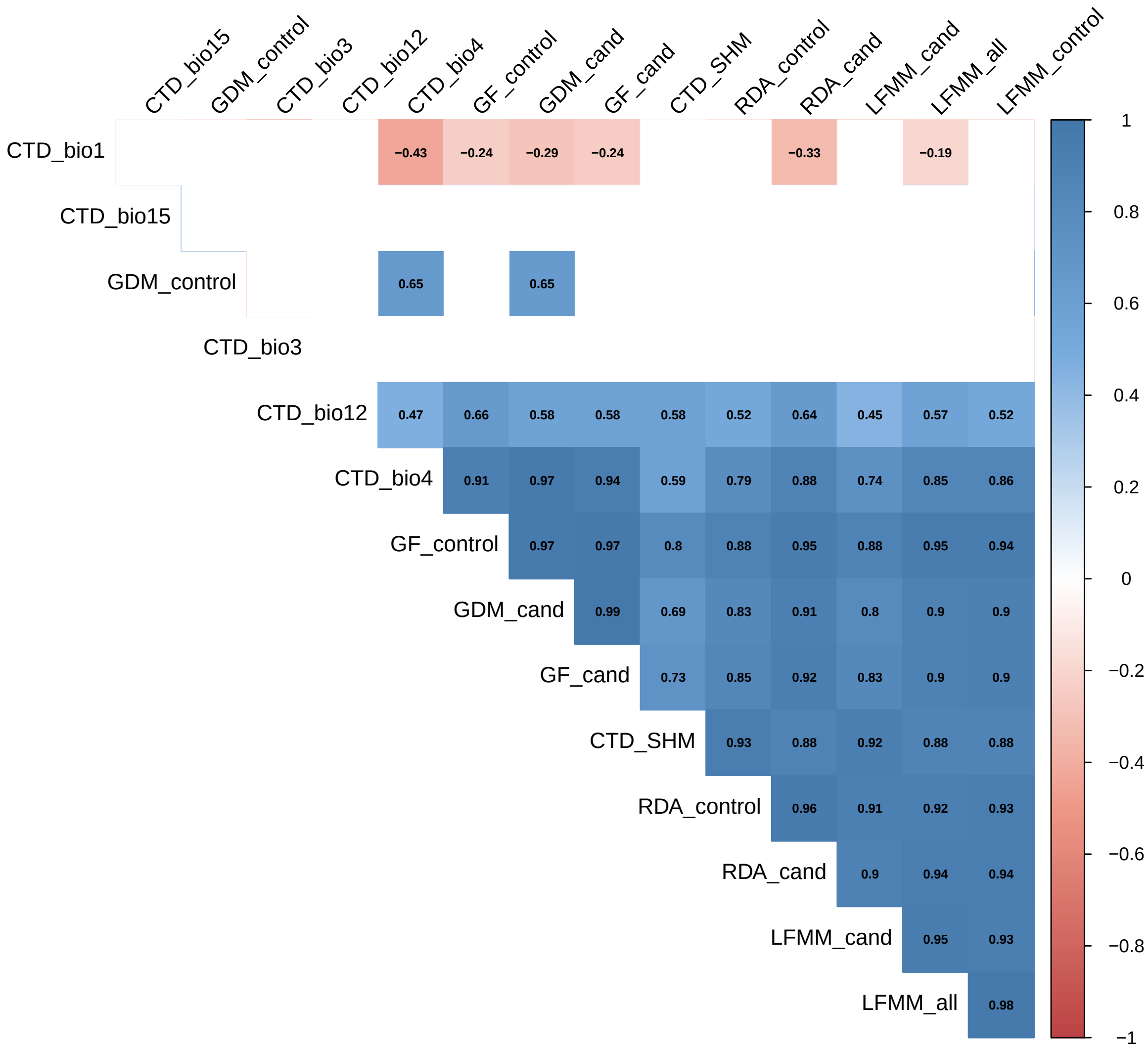

Figure 99: Correlations between GO predictions and CTDs in Fundão, Portugal.

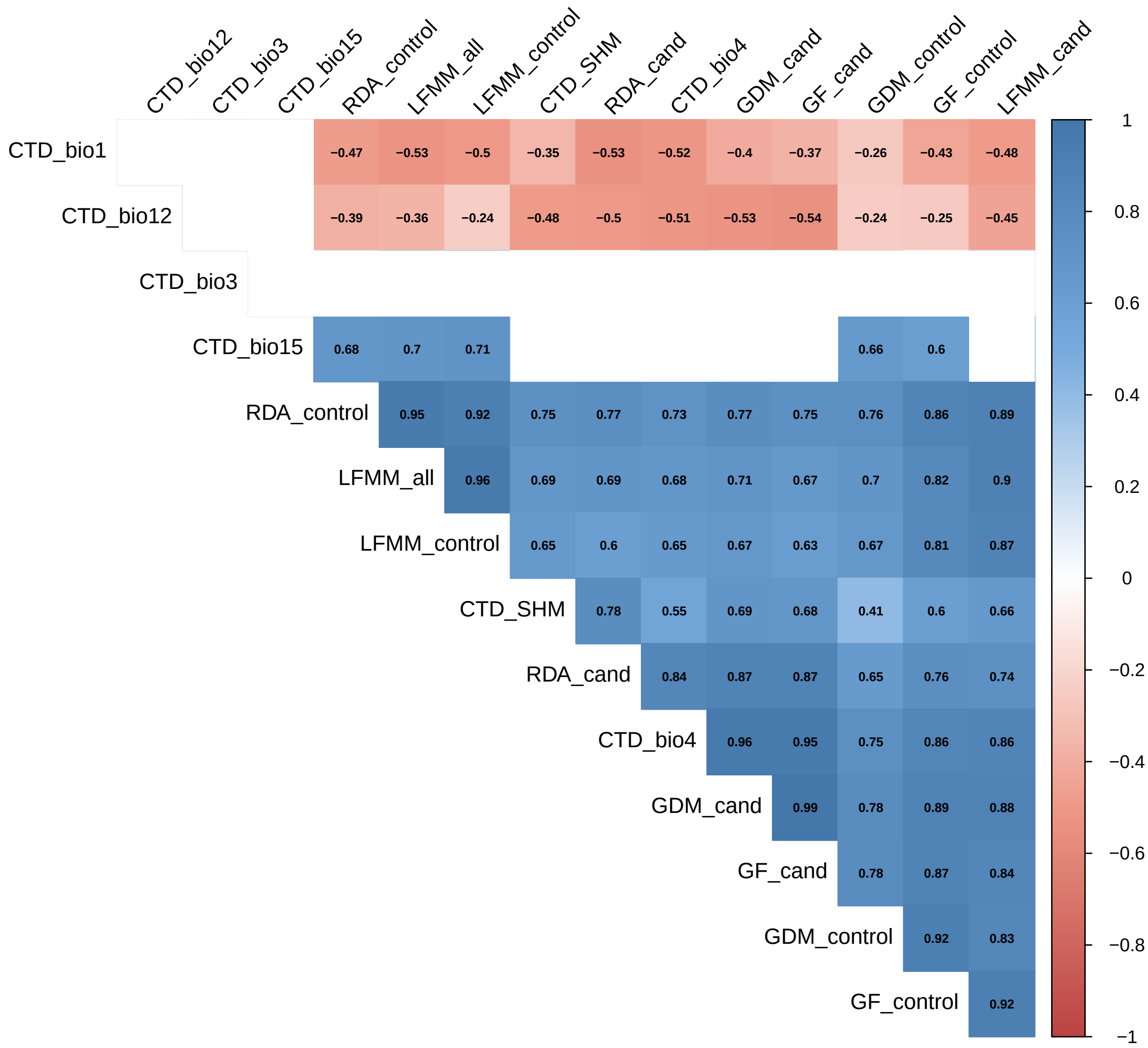

#### 9.3 Sample size

Table 9: Sample sizes in the five common gardens.

| Common garden (tree age) | Number of dead trees | Number of trees alive | Number of height measurements |
|---|---|---|---|
| Asturias, Spain (37 months) | 206 | 3976 | 3976 |
| Bordeaux, France (85 months) | 119 | 3225 | 3217 |
| Cáceres, Spain (8 months) | 3818 | 366 | 340 |
| Madrid, Spain (13 months) | 3138 | 1046 | 1046 |
| Fundão, Portugal (27 months) | 1517 | 2666 | 2665 |

Table 10: Number of dead and alive trees (dead / alive) per population in the five common gardens.

| Population | Asturias | Bordeaux | Cáceres | Madrid | Fundão |
|---|---|---|---|---|---|
| ALT | 3 / 72 | 1 / 53 | 69 / 72 | 52 / 72 | 19 / 72 |
| ARM | 1 / 64 | 1 / 64 | 56 / 64 | 49 / 64 | 23 / 64 |
| ARN | 5 / 136 | 3 / 109 | 129 / 136 | 107 / 136 | 46 / 136 |
| BAY | 8 / 144 | 3 / 142 | 132 / 144 | 113 / 144 | 56 / 144 |
| BON | 2 / 71 | 1 / 48 | 62 / 72 | 48 / 72 | 14 / 72 |
| CAD | 5 / 80 | 3 / 70 | 74 / 80 | 62 / 80 | 22 / 80 |
| CAR | 0 / 48 | 0 / 46 | 44 / 48 | 36 / 48 | 21 / 48 |
| CAS | 4 / 80 | 6 / 78 | 78 / 80 | 66 / 80 | 31 / 80 |
| CEN | 2 / 72 | 1 / 38 | 66 / 72 | 50 / 72 | 18 / 72 |
| COC | 6 / 144 | 3 / 109 | 137 / 144 | 118 / 144 | 53 / 143 |
| COM | 0 / 32 | 3 / 32 | 28 / 32 | 21 / 32 | 10 / 32 |
| CUE | 9 / 224 | 9 / 194 | 211 / 224 | 186 / 224 | 101 / 224 |
| HOU | 11 / 208 | 6 / 160 | 186 / 208 | 145 / 208 | 66 / 208 |
| LAM | 3 / 72 | 4 / 60 | 68 / 72 | 57 / 72 | 27 / 72 |
| LEI | 14 / 184 | 10 / 143 | 162 / 184 | 147 / 184 | 64 / 184 |
| MAD | 0 / 8 | 0 / 8 | 8 / 8 | 5 / 8 | 2 / 8 |
| MIM | 7 / 144 | 4 / 126 | 135 / 144 | 116 / 144 | 67 / 144 |
| OLB | 3 / 176 | 4 / 116 | 149 / 176 | 113 / 176 | 27 / 176 |
| OLO | 22 / 191 | 3 / 132 | 173 / 192 | 145 / 192 | 73 / 192 |
| ORI | 4 / 208 | 2 / 183 | 196 / 208 | 162 / 208 | 67 / 208 |
| PET | 11 / 192 | 3 / 147 | 171 / 192 | 136 / 192 | 75 / 192 |
| PIA | 8 / 128 | 1 / 110 | 111 / 128 | 91 / 128 | 43 / 128 |
| PIE | 3 / 72 | 4 / 55 | 67 / 72 | 50 / 72 | 17 / 72 |
| PLE | 8 / 160 | 9 / 138 | 155 / 160 | 131 / 160 | 71 / 160 |
| PUE | 1 / 64 | 5 / 56 | 58 / 64 | 50 / 64 | 30 / 64 |
| QUA | 5 / 136 | 3 / 117 | 123 / 136 | 83 / 136 | 36 / 136 |
| SAC | 1 / 72 | 2 / 51 | 65 / 72 | 56 / 72 | 35 / 72 |
| SAL | 7 / 112 | 0 / 69 | 104 / 112 | 85 / 112 | 55 / 112 |
| SEG | 12 / 168 | 7 / 168 | 154 / 168 | 135 / 168 | 77 / 168 |
| SIE | 2 / 64 | 1 / 47 | 59 / 64 | 40 / 64 | 29 / 64 |
| STJ | 15 / 224 | 3 / 180 | 190 / 224 | 154 / 224 | 70 / 224 |
| TAM | 8 / 120 | 4 / 71 | 114 / 120 | 107 / 120 | 60 / 120 |
| VAL | 5 / 96 | 5 / 64 | 87 / 96 | 69 / 96 | 29 / 96 |
| VER | 11 / 216 | 5 / 160 | 197 / 216 | 153 / 216 | 83 / 216 |

#### 9.4 Mortality models

##### 9.4.1 Mathematical model

Here is the mathematical model that we used to estimate the association between GO predictions or climate transfer distances (CTD) and the proportion of dead trees in the populations, independently in the five common gardens:

$$a_p \sim \text{Binomial}(N_p, p_p)$$
$$\text{logit}(p_p) = \beta_0 + \beta_H H_p + \beta_X X_p$$

- $a_p$ is the count of individuals that died in the population $p$
- $N_p$ is the total number of individuals in the population $p$ (=number of individuals that were initially planted in the common garden)
- $p_p$ is the estimated probability of mortality in the population $p$
- $X_p$ is the GO or CTD for the population $p$ and $\beta_X$ its associated coefficient (see Figure X of the manuscript).
- $H_p$ is the BLUPs for height of the population $p$, i.e. the population varying intercept estimated across all common gardens in the model 1 of Archambeau et al. (2022). $H_p$ is used as a proxy of the initial tree height of the population $p$. Indeed, tree height acts as a confounder in the experiment: trees that were higher at the time of planting were more likely to survive. As expected, this association was particularly strong in Madrid and Cáceres (Spain) where the seedlings experienced an extreme drought event the same year they were planted, resulting in very high mortality rates (Figure S101). More surprisingly, this association was also observed in Fundão, Portugal (Figure S101). However, the initial tree height was not associated with mortality rates in Bordeaux (France) and Asturias (Spain), which benefit from the favorable climates of the Atlantic region and in which the mortality rates were very low (Figure S101).

We used the following weakly informative priors:

$$\begin{bmatrix} \beta_{0,c} \\ \beta_H \\ \beta_X \end{bmatrix} \sim \mathcal{N}(0, 5)$$

##### 9.4.2 Estimated regression coefficients

###### 9.4.2.1 GO predictions and CTDs

Figure 100: Estimated mean and 95% credible intervals of the regression coefficients $\beta_X$ standing for the association between mortality rates and GO predictions or CTDs in the five common gardens. This figure is the same as Figure 5 in the main manuscript, except we added the regression coefficients for the CTDs.

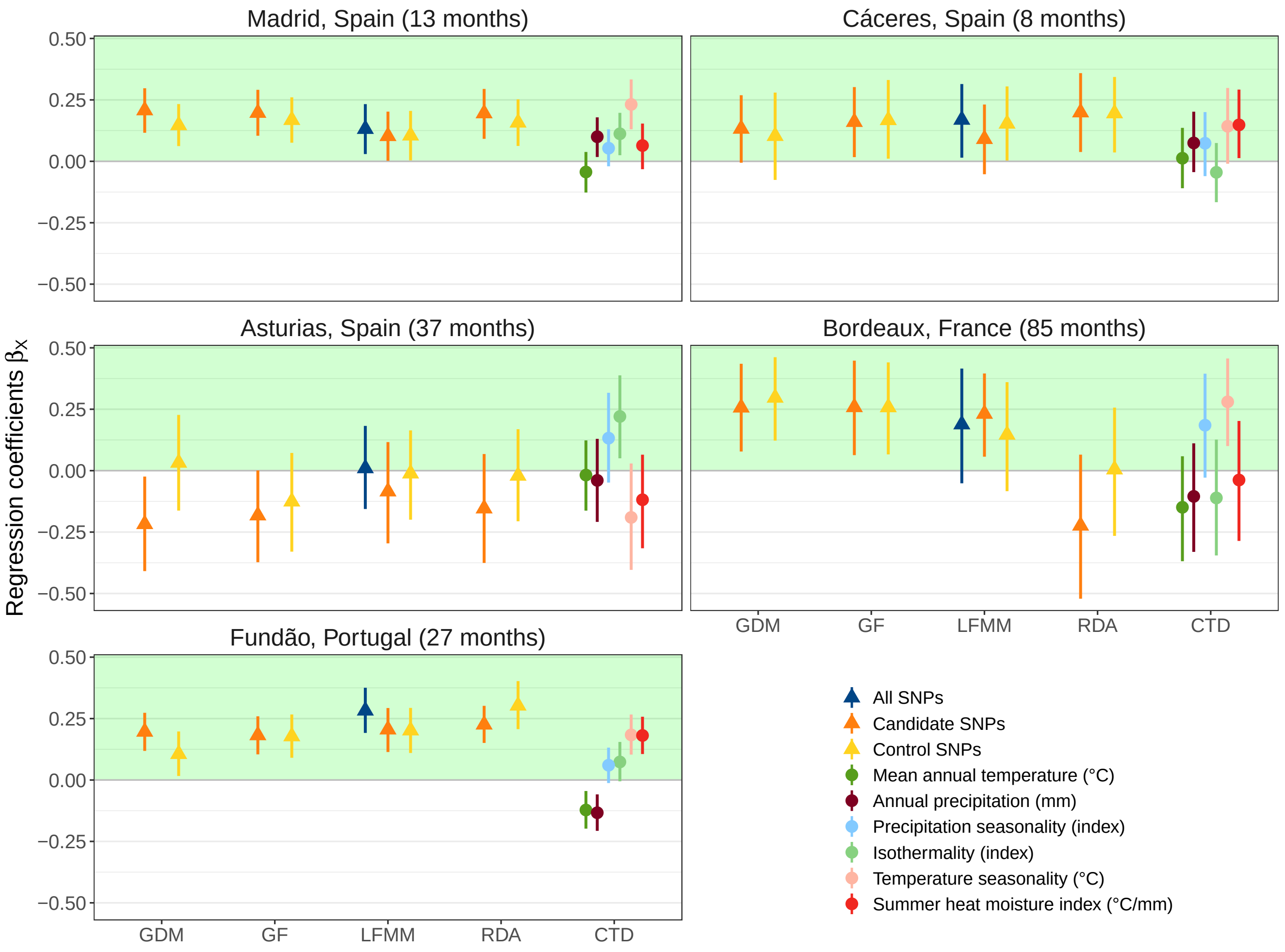

##### 9.4.2.2 Initial tree height

Figure 101: Estimated mean and 95% credible intervals of the regression coefficients $\beta_H$ standing for the association between mortality rates and the initial tree height.

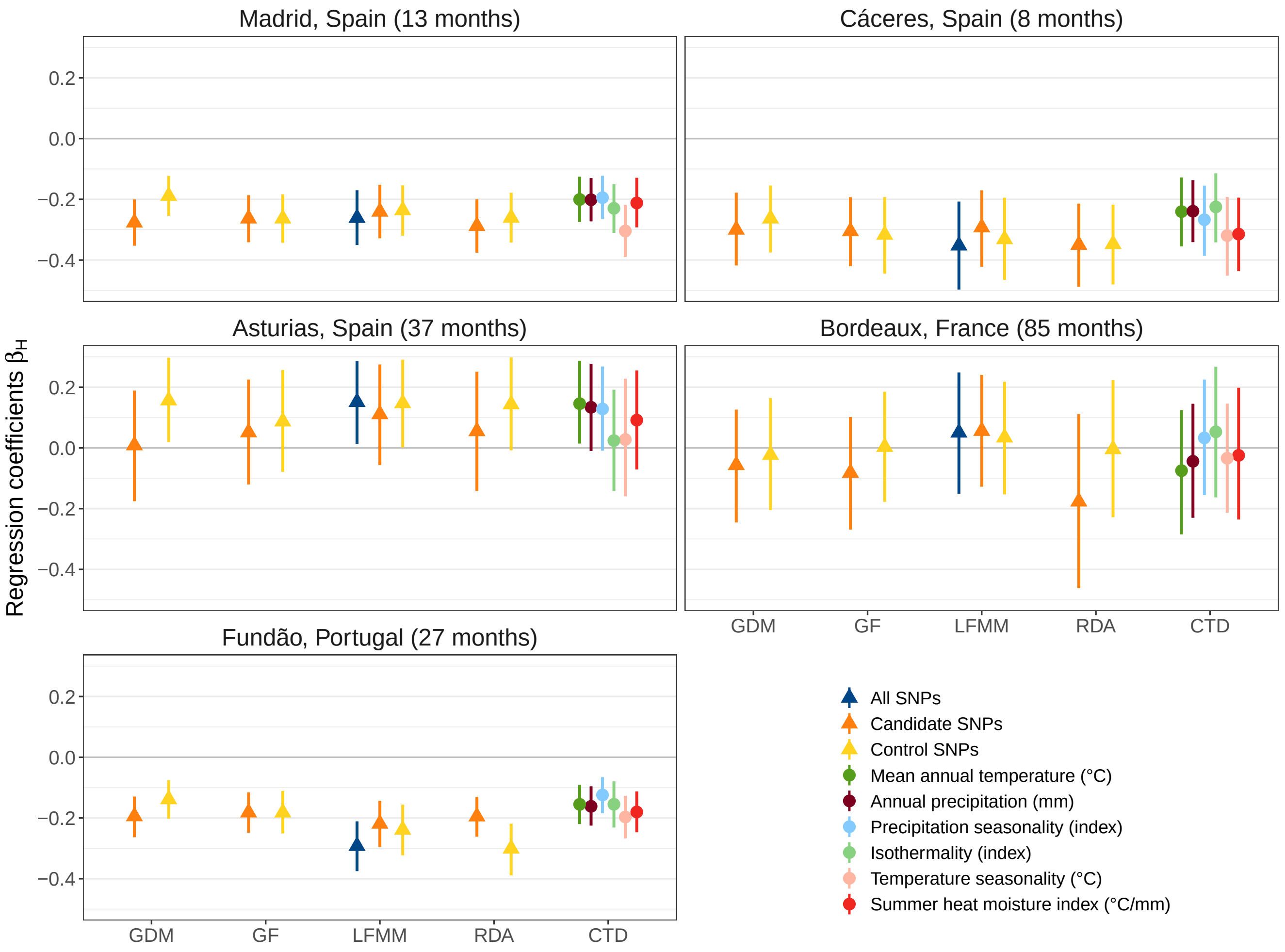

##### 9.4.3 Predictions of tree mortality probability

The plots below show the estimated association between tree mortality probability $p$ and GO predictions or CTDs. Tree mortality probability was predicted with the initial tree height $H$ fixed to its mean value in each common garden. The blue line corresponds to the mean estimate of the probability of tree mortality $p$ and the uncertainty interval corresponds to the its estimated 90% credible interval (highest posterior density interval).

###### 9.4.3.1 Plots with GO predictions as covariates

Figure 102: Predicted probability of tree mortality as a function of GO predictions in Asturias (Spain).

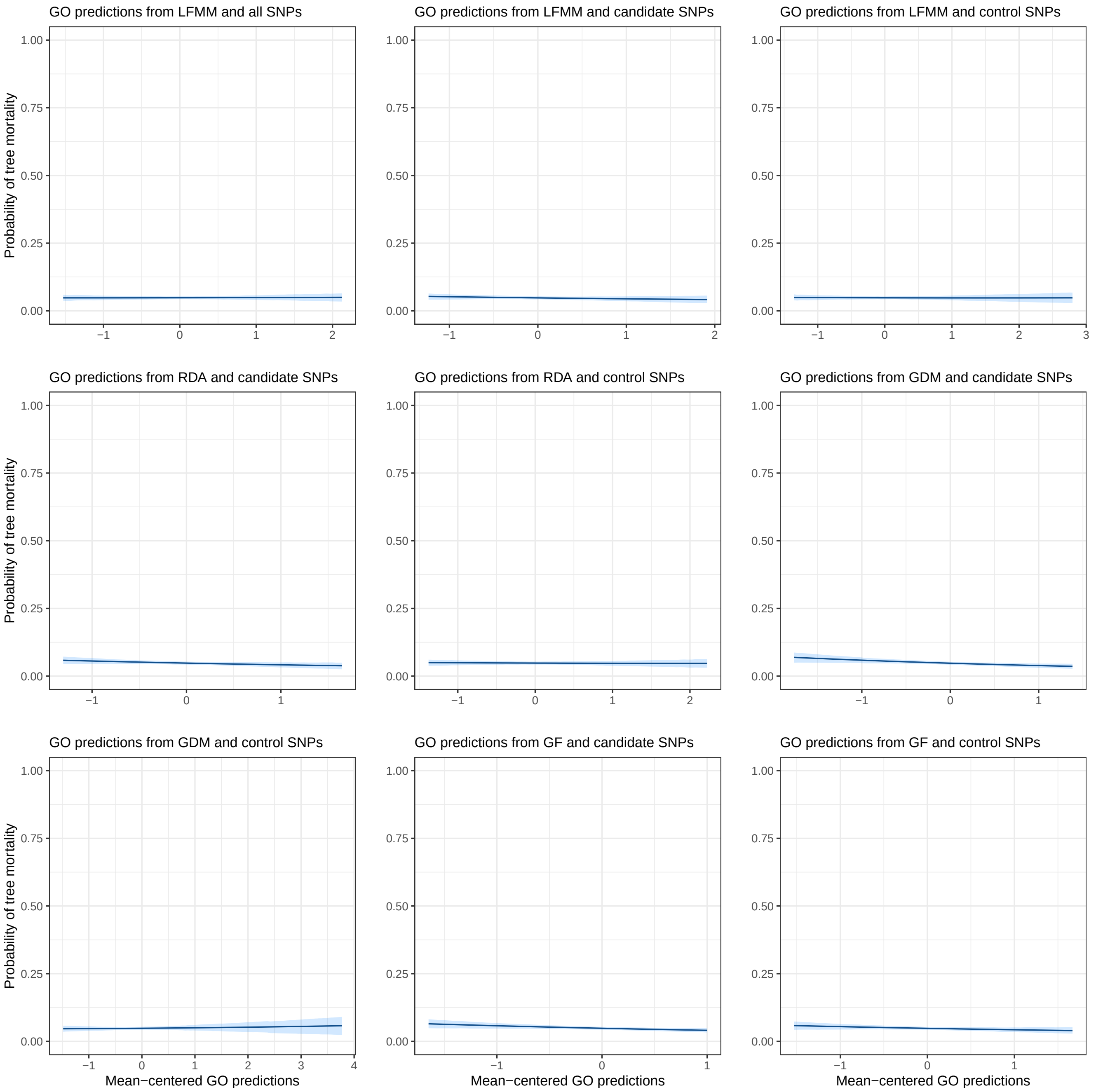

Figure 103: Predicted probability of tree mortality as a function of GO predictions in Bordeaux (France).

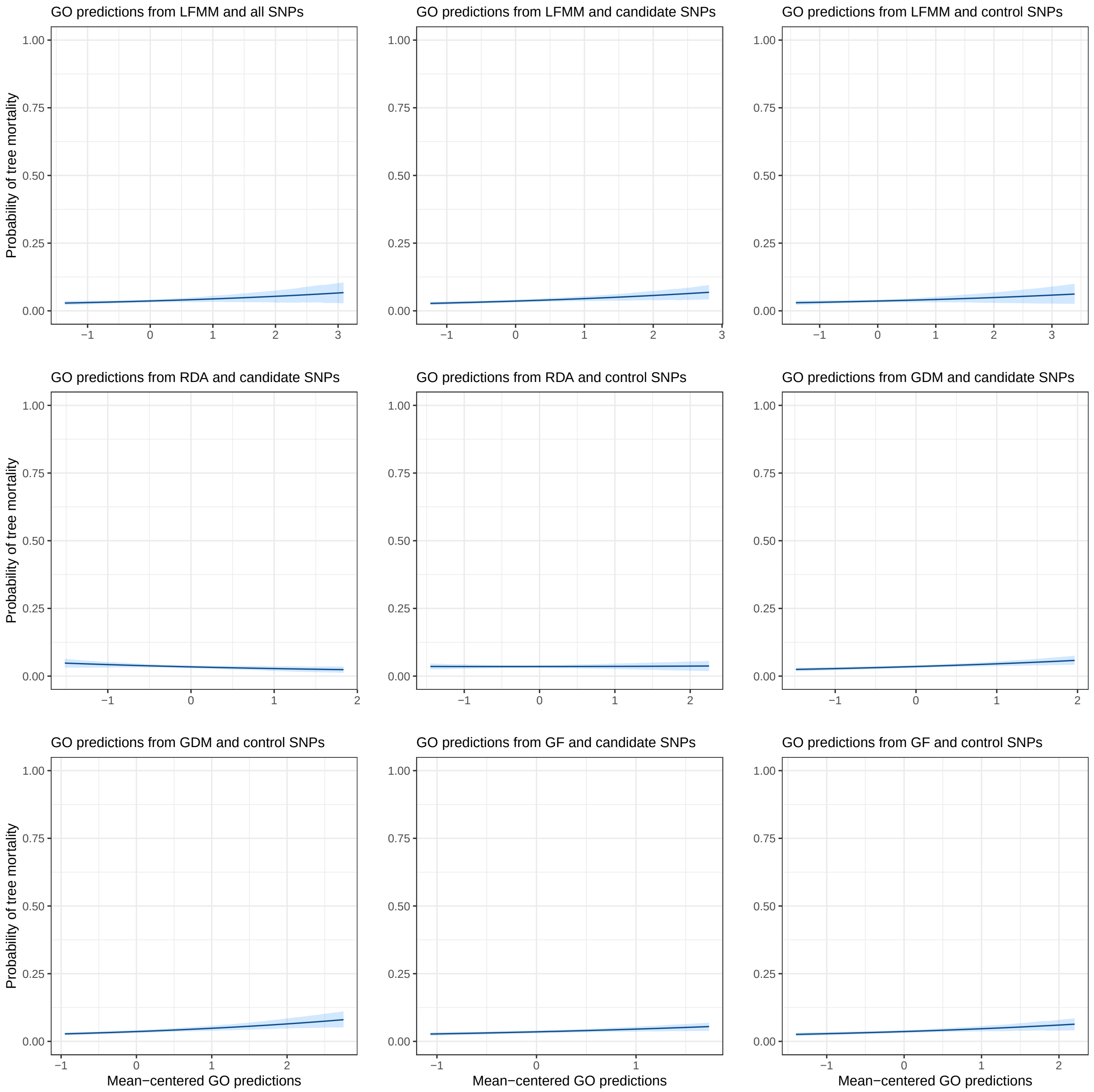

Figure 104: Predicted probability of tree mortality as a function of GO predictions in Cáceres (Spain).

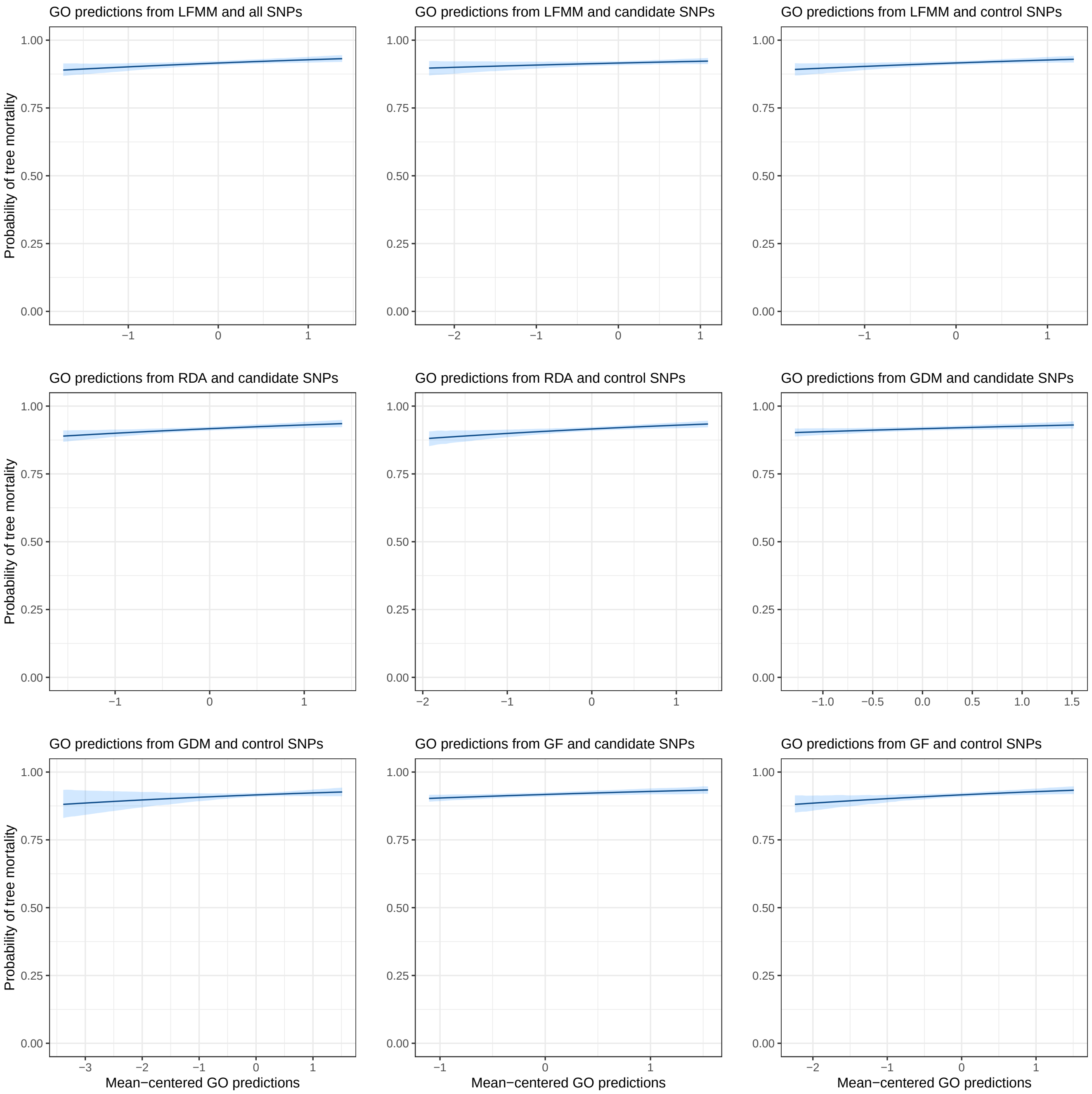

Figure 105: Predicted probability of tree mortality as a function of GO predictions in Madrid (Spain).

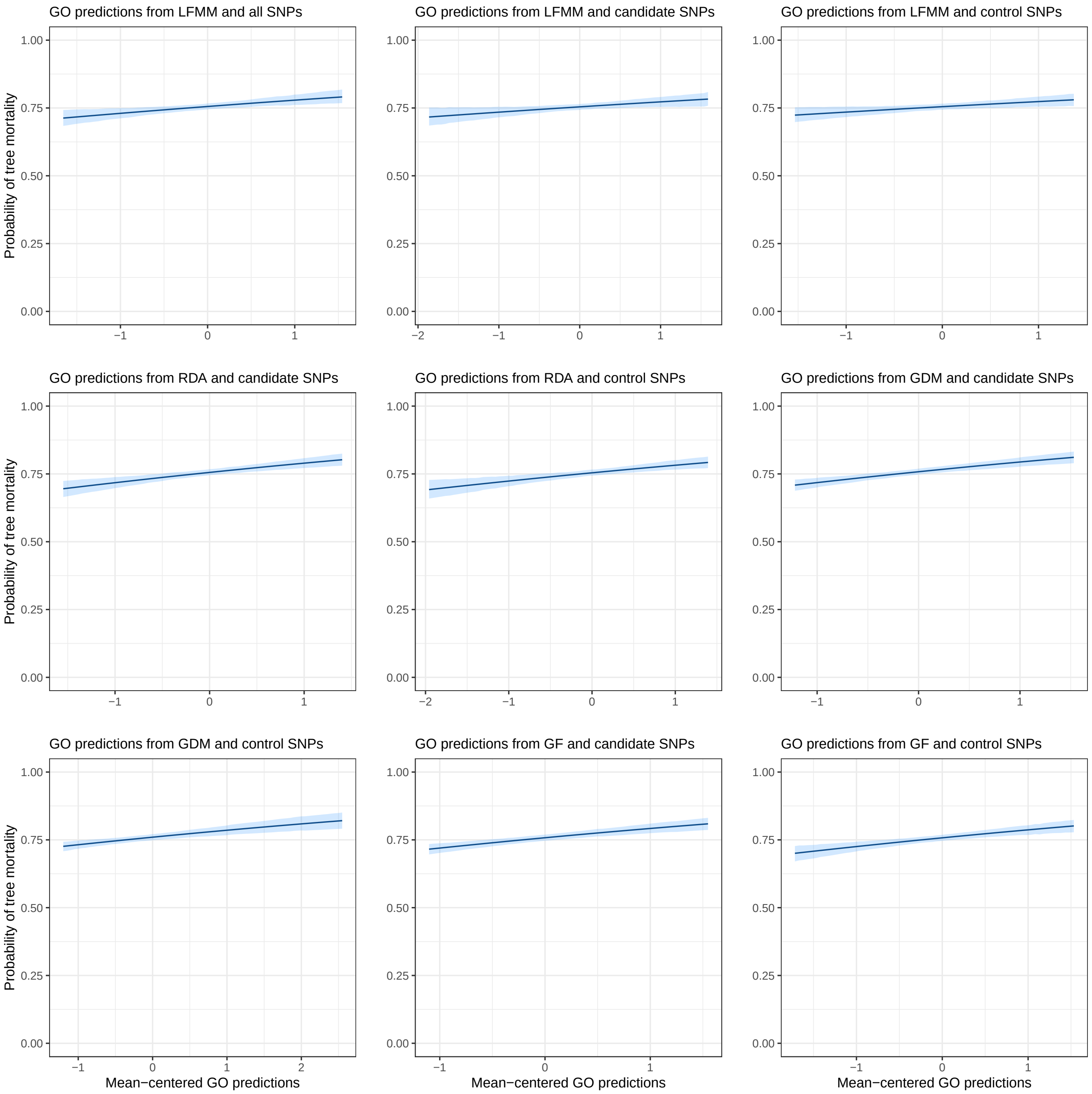

Figure 106: Predicted probability of tree mortality as a function of GO predictions in Fundão (Portugal).

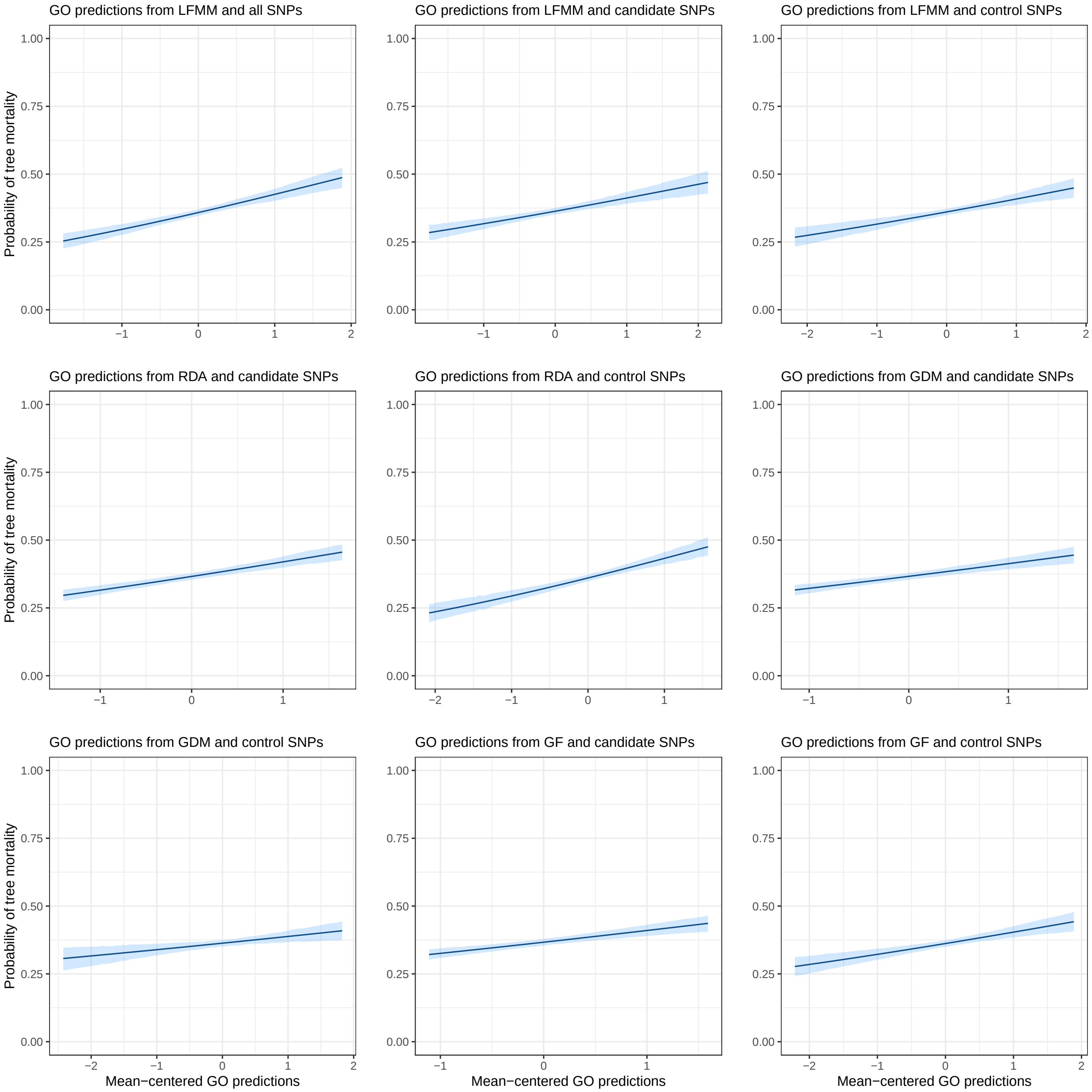

##### 9.4.3.2 Plots with CTDs as covariates

Figure 107: Predicted probability of tree mortality as a function of CTDs in Asturias (Spain).

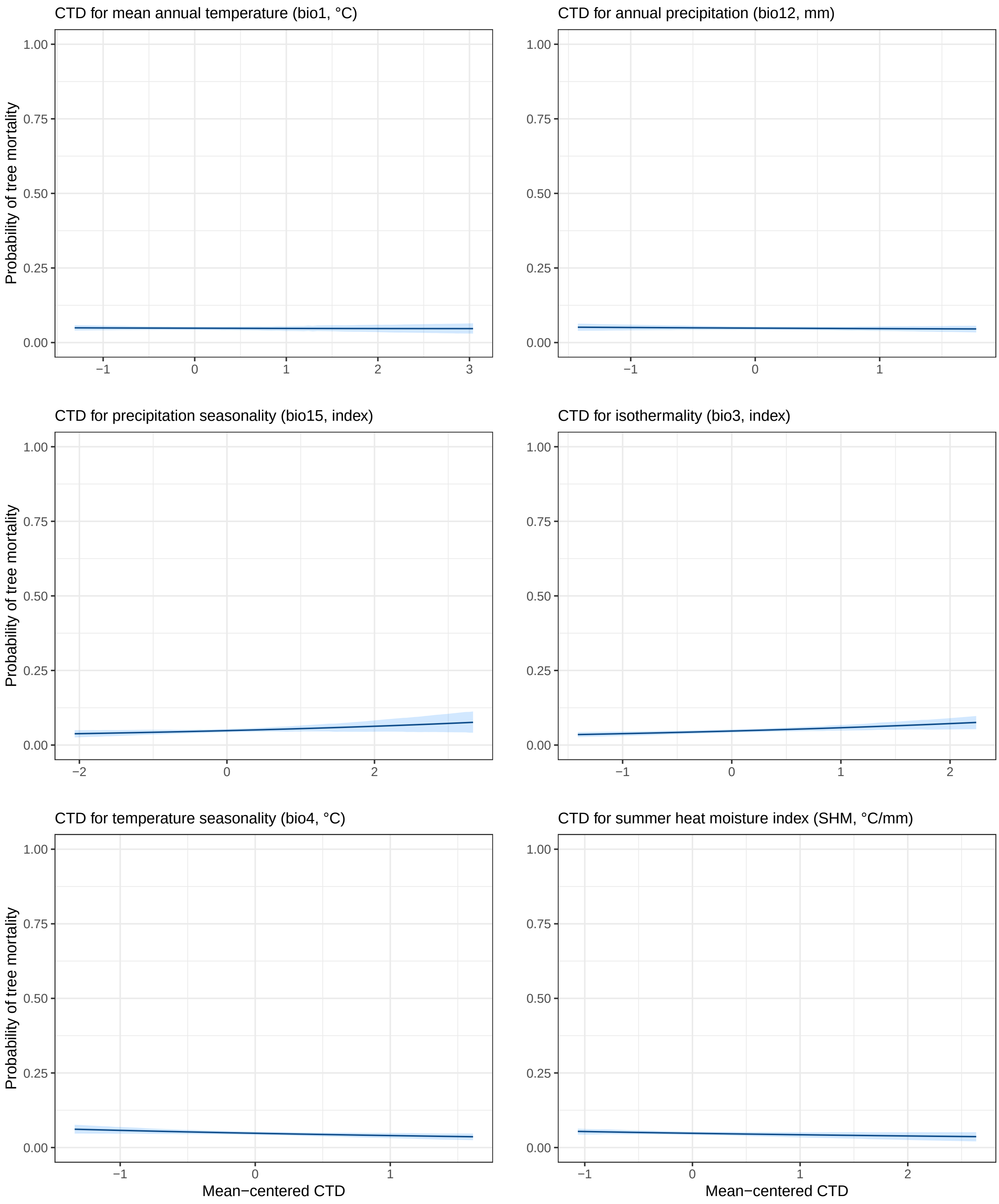

Figure 108: Predicted probability of tree mortality as a function of CTDs in Bordeaux (France).

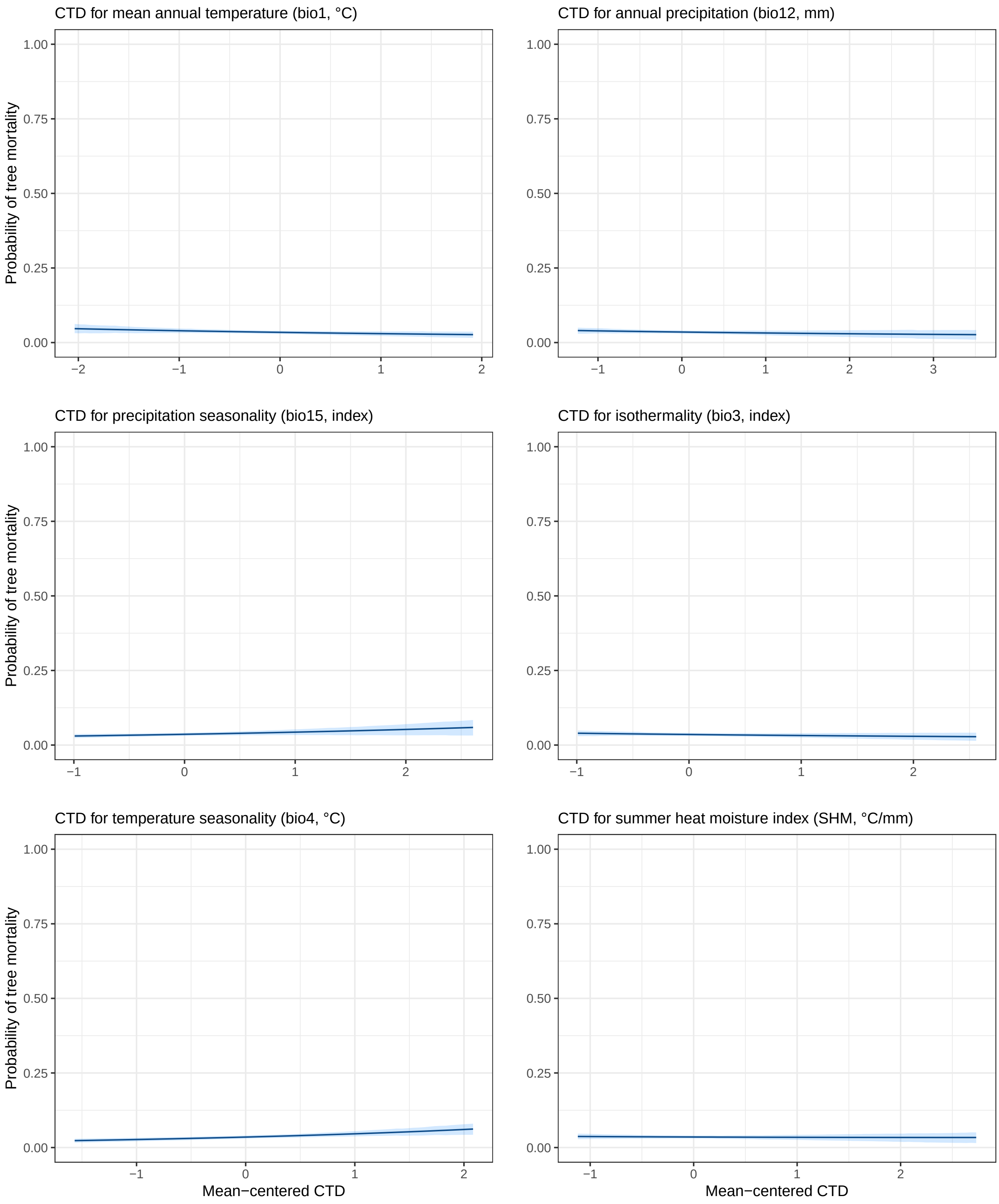

Figure 109: Predicted probability of tree mortality as a function of CTDs in Cáceres (Spain).

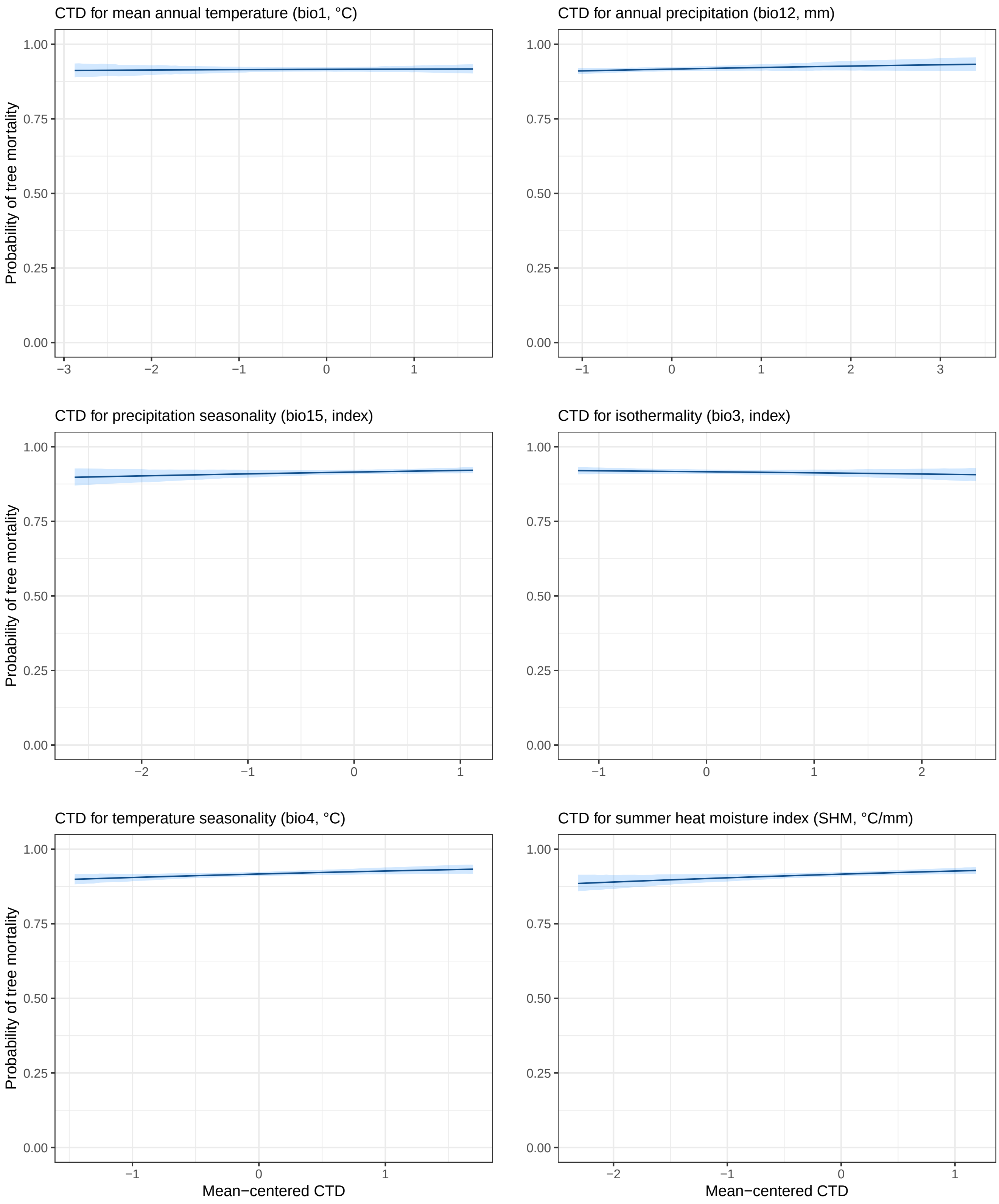

Figure 110: Predicted probability of tree mortality as a function of CTDs in Madrid (Spain).

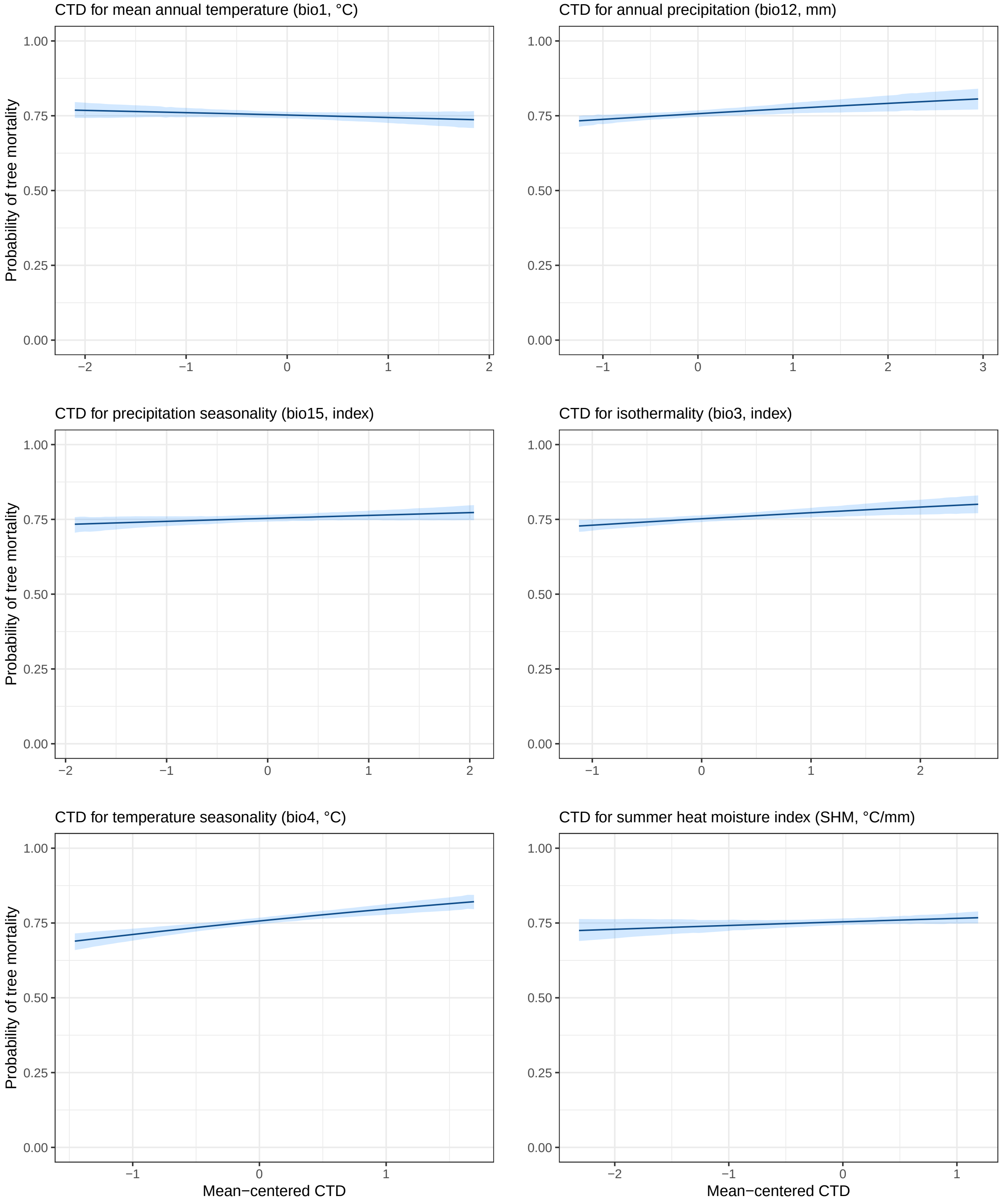

Figure 111: Predicted probability of tree mortality as a function of CTDs in Fundão (Portugal).

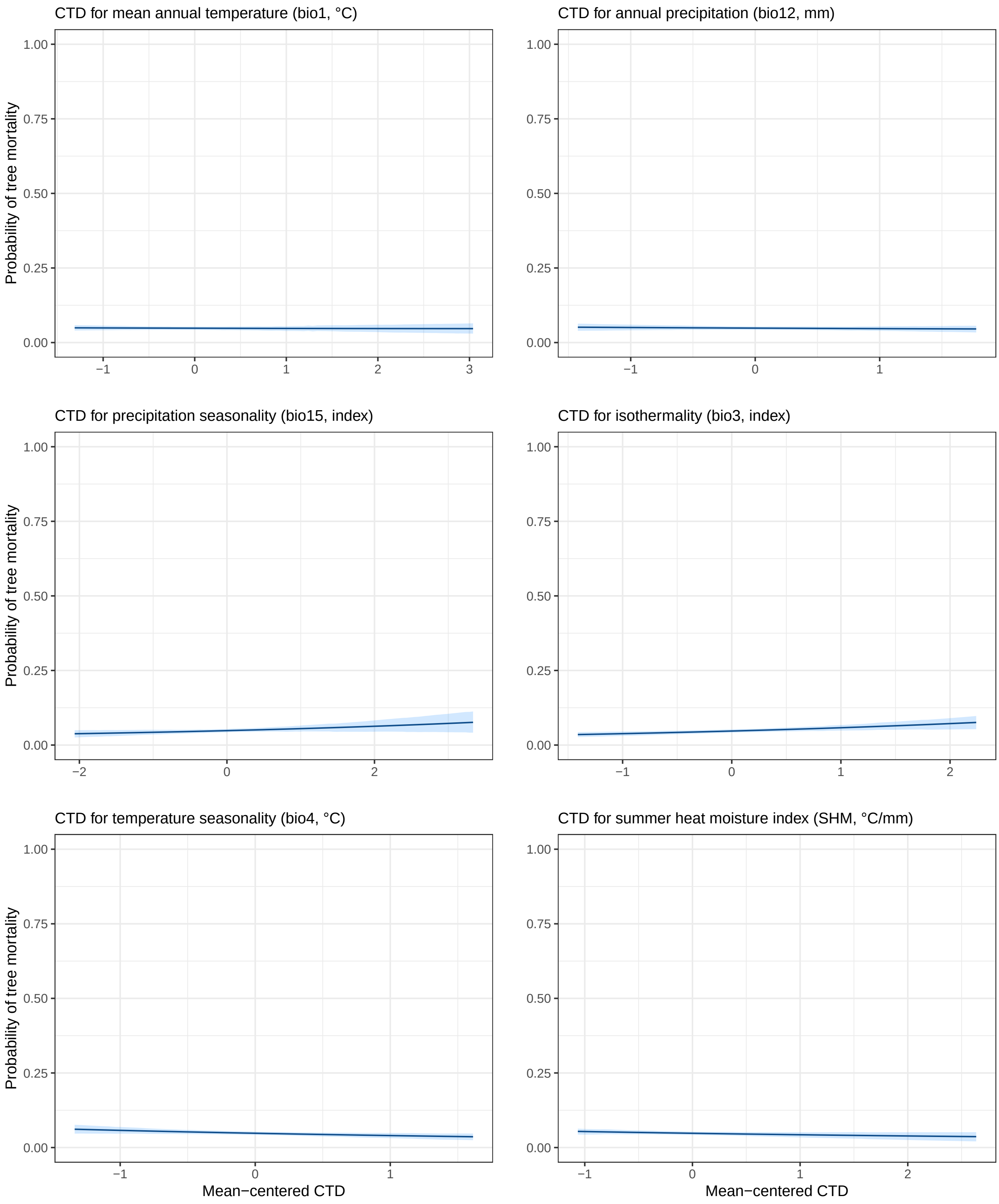

###### 9.4.3.3 Tables

The tables below provide:

- the predictions of tree mortality probability $p$ for different values of the predictor $X$ ($X$={-1, 0, 1}), with $H$ fixed to its mean value.
- the percent change of $p$ for a one-unit increase ($+\Delta_p$) or decrease ($-\Delta_p$) in $X$ (which corresponds to a one-standard deviation increase/decrease as $X$ is mean-centered).

Table 11: Predictions in Asturias (Spain).

| Method | p(x=-1) | p(x=0) | p(x=1) | +Δp | -Δp |
|---|---|---|---|---|---|
| GO predictions from LFMM and all SNPs | 0.048 [0.04;0.056] | 0.048 [0.043;0.053] | 0.049 [0.04;0.057] | 2.083% | 0% |
| GO predictions from LFMM and candidate SNPs | 0.052 [0.042;0.061] | 0.048 [0.043;0.054] | 0.044 [0.035;0.053] | -8.333% | 8.333% |
| GO predictions from LFMM and control SNPs | 0.049 [0.04;0.057] | 0.048 [0.043;0.053] | 0.048 [0.039;0.056] | 0% | 2.083% |
| GO predictions from RDA and candidate SNPs | 0.056 [0.045;0.067] | 0.048 [0.043;0.054] | 0.042 [0.032;0.051] | -12.5% | 16.667% |
| GO predictions from RDA and control SNPs | 0.049 [0.04;0.058] | 0.048 [0.043;0.054] | 0.047 [0.038;0.056] | -2.083% | 2.083% |
| GO predictions from GDM and candidate SNPs | 0.059 [0.047;0.068] | 0.048 [0.042;0.053] | 0.039 [0.03;0.046] | -18.75% | 22.917% |
| GO predictions from GDM and control SNPs | 0.047 [0.039;0.056] | 0.048 [0.043;0.054] | 0.05 [0.04;0.061] | 4.167% | -2.083% |
| GO predictions from GF and candidate SNPs | 0.058 [0.047;0.068] | 0.048 [0.043;0.054] | 0.041 [0.033;0.049] | -14.583% | 20.833% |
| GO predictions from GF and control SNPs | 0.054 [0.044;0.064] | 0.048 [0.043;0.053] | 0.043 [0.034;0.051] | -10.417% | 12.5% |
| CTD for mean annual temperature (bio1, °C) | 0.049 [0.041;0.057] | 0.048 [0.043;0.054] | 0.047 [0.039;0.055] | -2.083% | 2.083% |
| CTD for annual precipitation (bio12, mm) | 0.05 [0.042;0.06] | 0.048 [0.043;0.054] | 0.047 [0.039;0.054] | -2.083% | 4.167% |
| CTD for precipitation seasonality (bio15, index) | 0.043 [0.035;0.051] | 0.048 [0.042;0.053] | 0.055 [0.045;0.065] | 14.583% | -10.417% |
| CTD for isothermality (bio3, index) | 0.038 [0.031;0.045] | 0.047 [0.042;0.053] | 0.058 [0.048;0.067] | 23.404% | -19.149% |
| CTD for temperature seasonality (bio4, °C) | 0.058 [0.047;0.069] | 0.048 [0.042;0.053] | 0.04 [0.031;0.049] | -16.667% | 20.833% |
| CTD for summer heat moisture index (SHM, °C/mm) | 0.054 [0.043;0.062] | 0.048 [0.042;0.053] | 0.043 [0.034;0.051] | -10.417% | 12.5% |

Table 12: Predictions in Bordeaux (France).

| Method | p(x=-1) | p(x=0) | p(x=1) | +Δp | -Δp |
|---|---|---|---|---|---|
| GO predictions from LFMM and all SNPs | 0.031 [0.024;0.038] | 0.036 [0.031;0.042] | 0.044 [0.033;0.055] | 22.222% | -13.889% |
| GO predictions from LFMM and candidate SNPs | 0.029 [0.023;0.035] | 0.036 [0.031;0.042] | 0.045 [0.036;0.054] | 25% | -19.444% |
| GO predictions from LFMM and control SNPs | 0.032 [0.024;0.038] | 0.036 [0.031;0.041] | 0.042 [0.031;0.052] | 16.667% | -11.111% |
| GO predictions from RDA and candidate SNPs | 0.043 [0.031;0.052] | 0.034 [0.029;0.039] | 0.028 [0.019;0.036] | -17.647% | 26.471% |
| GO predictions from RDA and control SNPs | 0.035 [0.028;0.044] | 0.035 [0.03;0.041] | 0.036 [0.026;0.046] | 2.857% | 0% |
| GO predictions from GDM and candidate SNPs | 0.028 [0.022;0.034] | 0.035 [0.03;0.04] | 0.045 [0.037;0.054] | 28.571% | -20% |
| GO predictions from GDM and control SNPs | 0.027 [0.022;0.033] | 0.036 [0.031;0.042] | 0.048 [0.039;0.058] | 33.333% | -25% |
| GO predictions from GF and candidate SNPs | 0.028 [0.021;0.034] | 0.035 [0.03;0.04] | 0.045 [0.036;0.054] | 28.571% | -20% |
| GO predictions from GF and control SNPs | 0.028 [0.022;0.034] | 0.036 [0.031;0.041] | 0.047 [0.037;0.057] | 30.556% | -22.222% |
| CTD for mean annual temperature (bio1, °C) | 0.04 [0.032;0.047] | 0.034 [0.029;0.04] | 0.03 [0.022;0.038] | -11.765% | 17.647% |
| CTD for annual precipitation (bio12, mm) | 0.039 [0.031;0.048] | 0.035 [0.03;0.04] | 0.032 [0.025;0.04] | -8.571% | 11.429% |
| CTD for precipitation seasonality (bio15, index) | 0.03 [0.023;0.037] | 0.036 [0.031;0.041] | 0.043 [0.034;0.053] | 19.444% | -16.667% |
| CTD for isothermality (bio3, index) | 0.04 [0.03;0.049] | 0.035 [0.03;0.041] | 0.032 [0.025;0.04] | -8.571% | 14.286% |
| CTD for temperature seasonality (bio4, °C) | 0.027 [0.021;0.033] | 0.035 [0.03;0.04] | 0.046 [0.038;0.055] | 31.429% | -22.857% |
| CTD for summer heat moisture index (SHM, °C/mm) | 0.037 [0.028;0.046] | 0.035 [0.03;0.04] | 0.034 [0.025;0.043] | -2.857% | 5.714% |

Table 13: Predictions in Cáceres (Spain).

| Method | p(x=-1) | p(x=0) | p(x=1) | +Δp | -Δp |
|---|---|---|---|---|---|
| GO predictions from LFMM and all SNPs | 0.902 [0.887;0.915] | 0.916 [0.909;0.924] | 0.928 [0.917;0.939] | 1.31% | -1.528% |
| GO predictions from LFMM and candidate SNPs | 0.908 [0.895;0.921] | 0.916 [0.908;0.923] | 0.922 [0.912;0.933] | 0.655% | -0.873% |
| GO predictions from LFMM and control SNPs | 0.903 [0.89;0.916] | 0.916 [0.909;0.923] | 0.927 [0.917;0.938] | 1.201% | -1.419% |
| GO predictions from RDA and candidate SNPs | 0.9 [0.887;0.914] | 0.917 [0.91;0.924] | 0.93 [0.92;0.942] | 1.418% | -1.854% |
| GO predictions from RDA and control SNPs | 0.899 [0.885;0.913] | 0.916 [0.909;0.923] | 0.929 [0.918;0.939] | 1.419% | -1.856% |
| GO predictions from GDM and candidate SNPs | 0.906 [0.894;0.918] | 0.917 [0.91;0.924] | 0.926 [0.916;0.937] | 0.981% | -1.2% |
| GO predictions from GDM and control SNPs | 0.907 [0.892;0.922] | 0.916 [0.909;0.923] | 0.923 [0.912;0.936] | 0.764% | -0.983% |

| | | | | | |
|---|---|---|---|---|---|
| GO predictions from GF and candidate SNPs | 0.904 [0.893;0.916] | 0.917 [0.91;0.924] | 0.929 [0.918;0.94] | 1.309% | -1.418% |
| GO predictions from GF and control SNPs | 0.902 [0.888;0.916] | 0.916 [0.909;0.923] | 0.928 [0.916;0.939] | 1.31% | -1.528% |
| CTD for mean annual temperature (bio1, °C) | 0.915 [0.905;0.925] | 0.916 [0.909;0.923] | 0.917 [0.906;0.928] | 0.109% | -0.109% |
| CTD for annual precipitation (bio12, mm) | 0.911 [0.9;0.921] | 0.917 [0.91;0.924] | 0.922 [0.911;0.933] | 0.545% | -0.654% |
| CTD for precipitation seasonality (bio15, index) | 0.909 [0.897;0.922] | 0.915 [0.907;0.922] | 0.921 [0.911;0.93] | 0.656% | -0.656% |
| CTD for isothermality (bio3, index) | 0.919 [0.908;0.93] | 0.916 [0.909;0.924] | 0.913 [0.903;0.923] | -0.328% | 0.328% |
| CTD for temperature seasonality (bio4, °C) | 0.905 [0.892;0.917] | 0.917 [0.91;0.924] | 0.927 [0.916;0.939] | 1.091% | -1.309% |
| CTD for summer heat moisture index (SHM, °C/mm) | 0.904 [0.892;0.916] | 0.916 [0.909;0.923] | 0.927 [0.916;0.937] | 1.201% | -1.31% |

Table 14: Predictions in Madrid (Spain).

| Method | p(x=-1) | p(x=0) | p(x=1) | +Δp | -Δp |
|---|---|---|---|---|---|
| GO predictions from LFMM and all SNPs | 0.73 [0.712;0.749] | 0.755 [0.744;0.766] | 0.779 [0.761;0.799] | 3.179% | -3.311% |
| GO predictions from LFMM and candidate SNPs | 0.735 [0.716;0.755] | 0.754 [0.743;0.765] | 0.773 [0.754;0.79] | 2.52% | -2.52% |
| GO predictions from LFMM and control SNPs | 0.735 [0.717;0.755] | 0.755 [0.744;0.766] | 0.774 [0.755;0.792] | 2.517% | -2.649% |
| GO predictions from RDA and candidate SNPs | 0.718 [0.698;0.738] | 0.756 [0.745;0.767] | 0.79 [0.772;0.808] | 4.497% | -5.026% |
| GO predictions from RDA and control SNPs | 0.724 [0.704;0.743] | 0.754 [0.744;0.766] | 0.782 [0.766;0.801] | 3.714% | -3.979% |
| GO predictions from GDM and candidate SNPs | 0.718 [0.7;0.735] | 0.758 [0.748;0.77] | 0.794 [0.777;0.811] | 4.749% | -5.277% |
| GO predictions from GDM and control SNPs | 0.732 [0.717;0.747] | 0.76 [0.749;0.771] | 0.785 [0.767;0.803] | 3.289% | -3.684% |
| GO predictions from GF and candidate SNPs | 0.72 [0.702;0.738] | 0.758 [0.747;0.769] | 0.792 [0.775;0.81] | 4.485% | -5.013% |
| GO predictions from GF and control SNPs | 0.725 [0.707;0.743] | 0.758 [0.746;0.768] | 0.787 [0.768;0.804] | 3.826% | -4.354% |
| CTD for mean annual temperature (bio1, °C) | 0.76 [0.745;0.776] | 0.752 [0.741;0.763] | 0.744 [0.726;0.763] | -1.064% | 1.064% |
| CTD for annual precipitation (bio12, mm) | 0.738 [0.722;0.753] | 0.757 [0.746;0.768] | 0.775 [0.758;0.793] | 2.378% | -2.51% |
| CTD for precipitation seasonality (bio15, index) | 0.743 [0.727;0.76] | 0.754 [0.743;0.765] | 0.763 [0.747;0.779] | 1.194% | -1.459% |
| CTD for isothermality (bio3, index) | 0.731 [0.713;0.75] | 0.752 [0.741;0.763] | 0.772 [0.757;0.789] | 2.66% | -2.793% |
| CTD for temperature seasonality (bio4, °C) | 0.712 [0.691;0.731] | 0.757 [0.746;0.768] | 0.797 [0.778;0.813] | 5.284% | -5.945% |
| CTD for summer heat moisture index (SHM, °C/mm) | 0.742 [0.724;0.76] | 0.754 [0.743;0.765] | 0.766 [0.748;0.784] | 1.592% | -1.592% |

Table 15: Predictions in Fundão (Portugal).

| Method | p(x=-1) | p(x=0) | p(x=1) | +Δp | -Δp |
|---|---|---|---|---|---|
| GO predictions from LFMM and all SNPs | 0.296 [0.276;0.316] | 0.358 [0.346;0.371] | 0.426 [0.403;0.447] | 18.994% | -17.318% |
| GO predictions from LFMM and candidate SNPs | 0.317 [0.297;0.337] | 0.363 [0.352;0.376] | 0.412 [0.391;0.434] | 13.499% | -12.672% |
| GO predictions from LFMM and control SNPs | 0.316 [0.295;0.337] | 0.361 [0.348;0.372] | 0.408 [0.386;0.429] | 13.019% | -12.465% |
| GO predictions from RDA and candidate SNPs | 0.316 [0.298;0.333] | 0.366 [0.354;0.378] | 0.42 [0.399;0.441] | 14.754% | -13.661% |
| GO predictions from RDA and control SNPs | 0.294 [0.274;0.316] | 0.36 [0.348;0.373] | 0.433 [0.411;0.456] | 20.278% | -18.333% |
| GO predictions from GDM and candidate SNPs | 0.322 [0.305;0.34] | 0.366 [0.354;0.38] | 0.413 [0.393;0.435] | 12.842% | -12.022% |
| GO predictions from GDM and control SNPs | 0.339 [0.318;0.361] | 0.363 [0.351;0.375] | 0.388 [0.367;0.41] | 6.887% | -6.612% |
| GO predictions from GF and candidate SNPs | 0.326 [0.309;0.344] | 0.367 [0.354;0.378] | 0.41 [0.39;0.431] | 11.717% | -11.172% |
| GO predictions from GF and control SNPs | 0.322 [0.302;0.343] | 0.362 [0.349;0.374] | 0.404 [0.384;0.425] | 11.602% | -11.05% |
| CTD for mean annual temperature (bio1, °C) | 0.39 [0.37;0.408] | 0.361 [0.35;0.374] | 0.334 [0.314;0.352] | -7.479% | 8.033% |
| CTD for annual precipitation (bio12, mm) | 0.4 [0.378;0.421] | 0.369 [0.357;0.382] | 0.339 [0.322;0.355] | -8.13% | 8.401% |
| CTD for precipitation seasonality (bio15, index) | 0.347 [0.329;0.369] | 0.361 [0.349;0.374] | 0.375 [0.358;0.392] | 3.878% | -3.878% |
| CTD for isothermality (bio3, index) | 0.346 [0.327;0.366] | 0.363 [0.351;0.375] | 0.38 [0.361;0.399] | 4.683% | -4.683% |
| CTD for temperature seasonality (bio4, °C) | 0.324 [0.305;0.342] | 0.366 [0.354;0.378] | 0.409 [0.39;0.432] | 11.749% | -11.475% |
| CTD for summer heat moisture index (SHM, °C/mm) | 0.325 [0.307;0.343] | 0.365 [0.353;0.377] | 0.408 [0.389;0.429] | 11.781% | -10.959% |

#### 9.5 Height models

##### 9.5.1 Mathematical model

Here is the mathematical model used to estimate the association between tree height and GO predictions or CTD, independently in the five common gardens:

$$
\begin{aligned}
Y_{ipb} &\sim \mathcal{N}(\mu_{pb}, \sigma^2) \\
\mu_{pb} &= B_b + \beta_{X1} X_p + \beta_{X2} X_p^2 + \beta_H H_p
\end{aligned}
$$

- $Y_{ipb}$ is the height of the individual $i$ in the population $p$ and in the block $b$
- $\sigma^2$ is the residual variance of the model.
- $B_b$ are the block intercepts
- $X_p$ is the GO or CTD of the population $p$, with $\beta_{X1}$ and $\beta_{X2}$ being its linear and quadratic coefficients, respectively. The quadratic term was included to allow for potential nonlinearity in the response, following Fitzpatrick et al. (2021).
- $H_p$ is the proxy of the initial tree height of the population $p$, see mortality models above.

We used the following weakly informative priors:

$$
\begin{aligned}
\sigma &\sim \text{Exp}(1) \\
\begin{bmatrix} B_b \\ \beta_{X1} \\ \beta_{X2} \\ \beta_H \end{bmatrix} &\sim \mathcal{N}(0, 1)
\end{aligned}
$$

The proportion of variance explained by the height models was estimated with the Bayesian version of $\mathcal{R}^2$ introduced in Gelman et al. (2019):

$$
\mathcal{R}^2 = \frac{Var_\mu}{Var_\mu + Var_{res}}
$$

where $Var_\mu$ is the variance of the modelled predictive means and $Var_{res}$ is the modelled residual variance, such as $Var_{res} = \sigma^2$.

##### 9.5.2 Estimated regression coefficients

###### 9.5.2.1 Initial tree height

Figure 112: Estimated mean and 95% credible intervals of the regression coefficients $\beta_H$ standing for the association between tree height and the population-level BLUPs for height used as a proxy of the initial tree height of the populations. Graph titles include the time in months corresponding to the age at which height and survival were recorded.

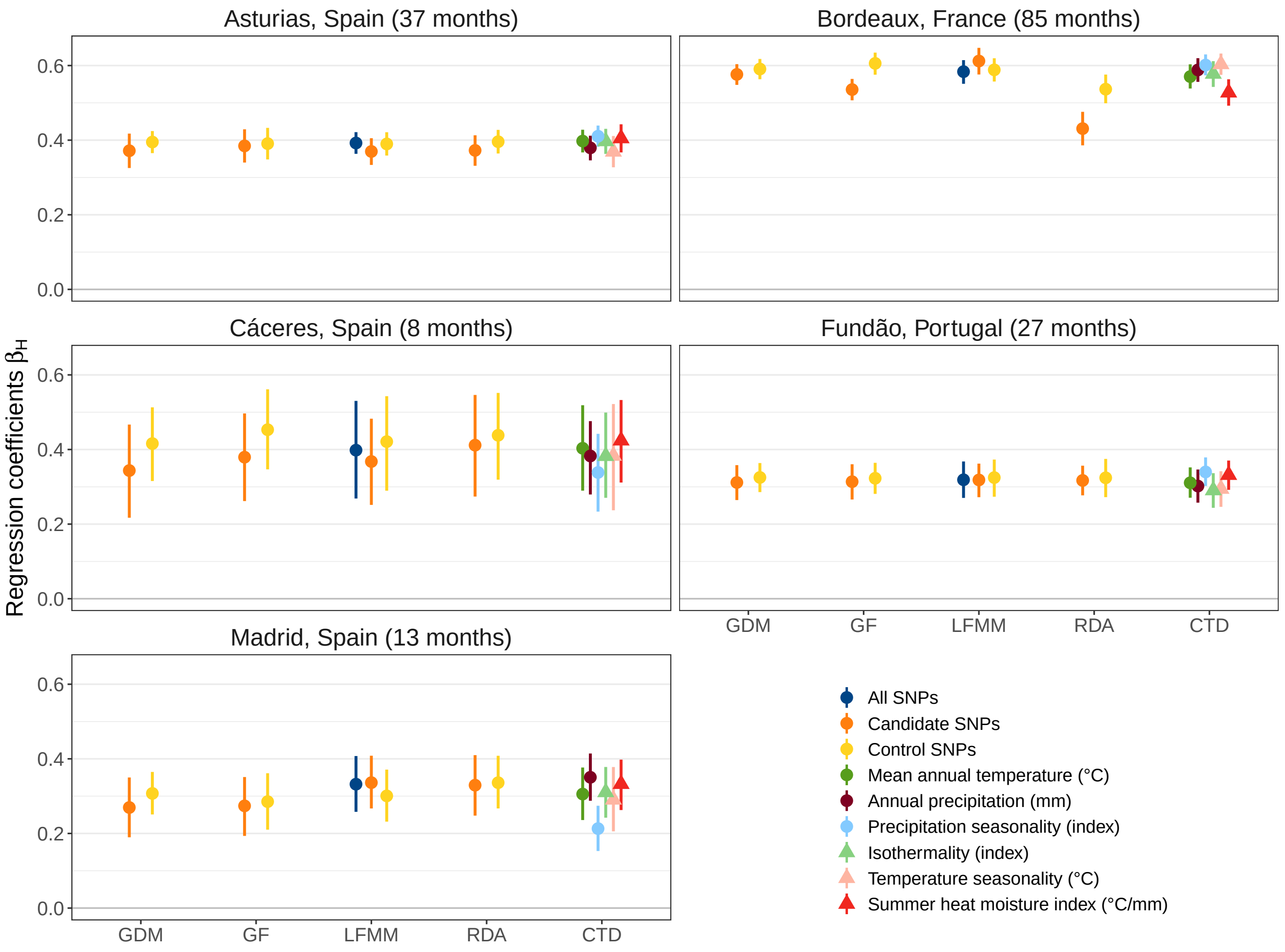

###### 9.5.2.2 GO predictions and CTDs

Figure 113: Estimated mean and 95% credible intervals of the regression coefficients $\beta_{X_1}$ standing for the association between tree height and GO predictions or CTD in the five common gardens. Graph titles include the time in months corresponding to the age at which height and survival were recorded.

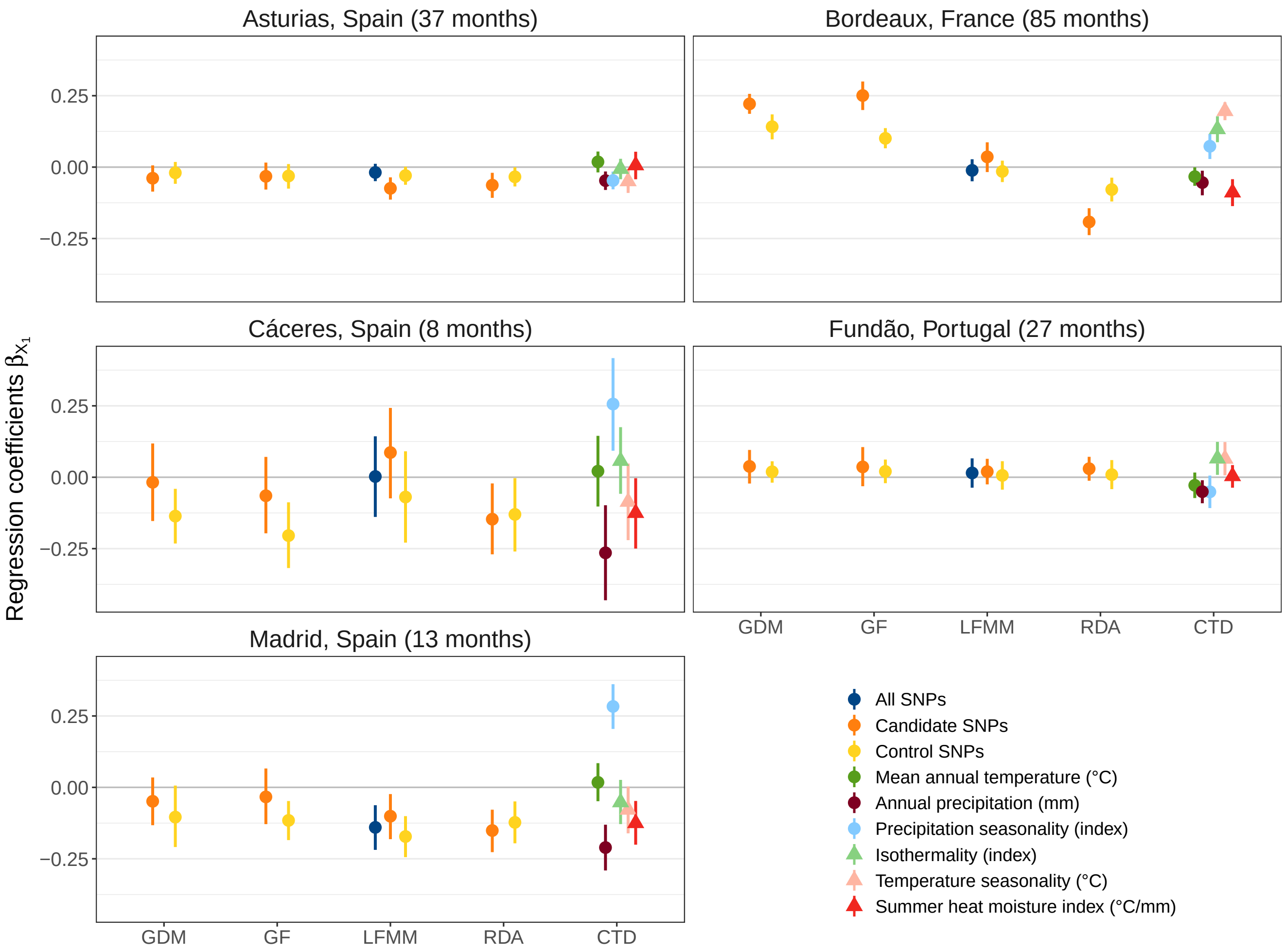

Figure 114: Estimated mean and 95% credible intervals of the regression coefficients $\beta_{X_2}$ standing for the association between tree height and GO predictions or CTD in the five common gardens. Graph titles include the time in months corresponding to the age at which height and survival were recorded.

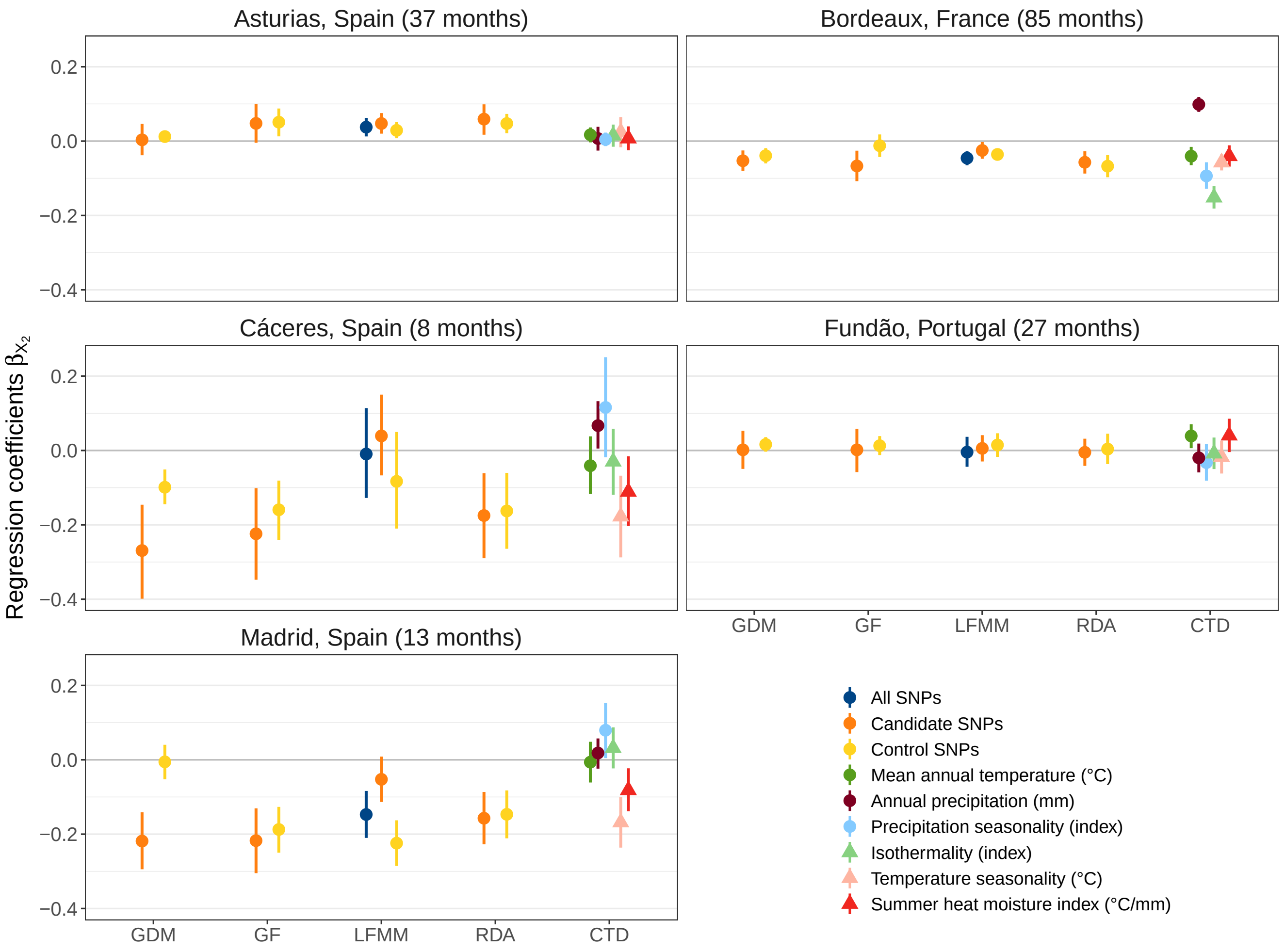

##### 9.5.3 Height predictions

In the graphs below, the blue line corresponds to the mean estimate of the linear predictor $\mu$, i.e. the predicted mean tree height. The narrower uncertainty interval corresponds to the estimated 90% credible intervals of the linear predictor $\mu$. The wider uncertainty interval corresponds to the 90% uncertainty interval of height predictions after accounting for the model uncertainty (i.e. the $\sigma^2$ parameter).

###### 9.5.3.1 Plots with GO predictions as covariates

Figure 115: Tree height predictions as a function of GO predictions in Asturias (Spain).

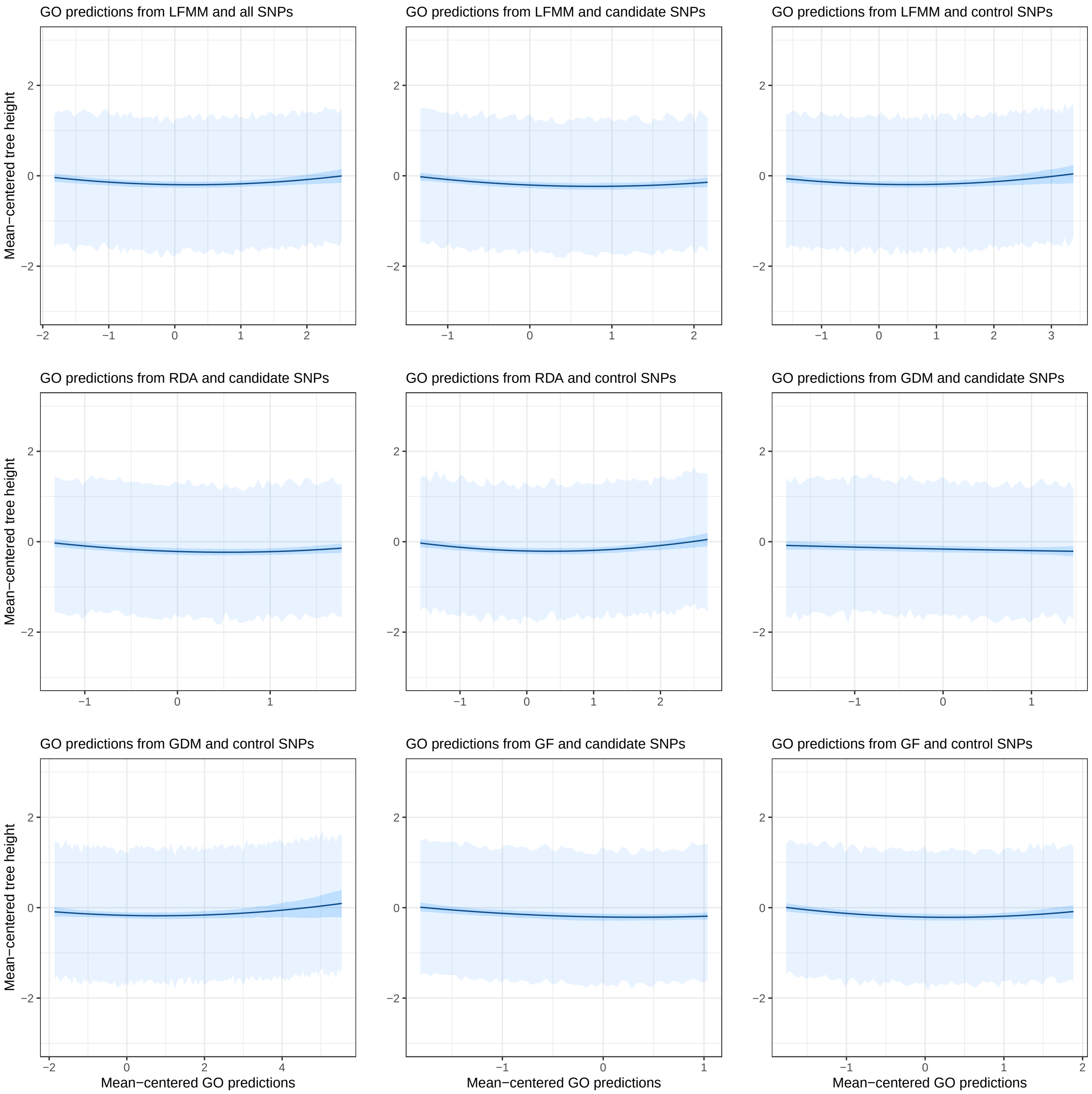

Figure 116: Tree height predictions as a function of GO predictions in Cáceres (Spain).

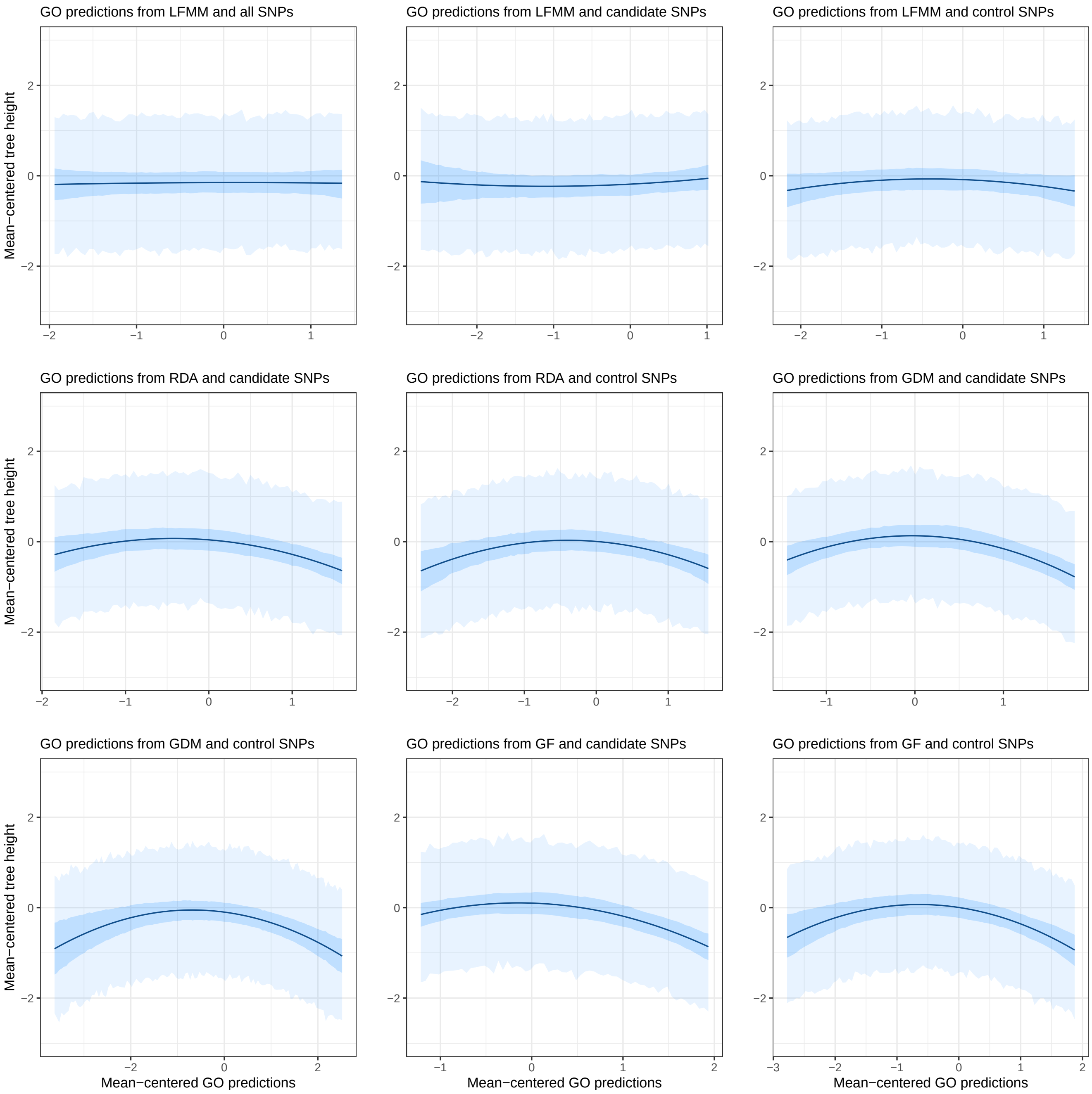

Figure 117: Tree height predictions as a function of GO predictions in Bordeaux (France).

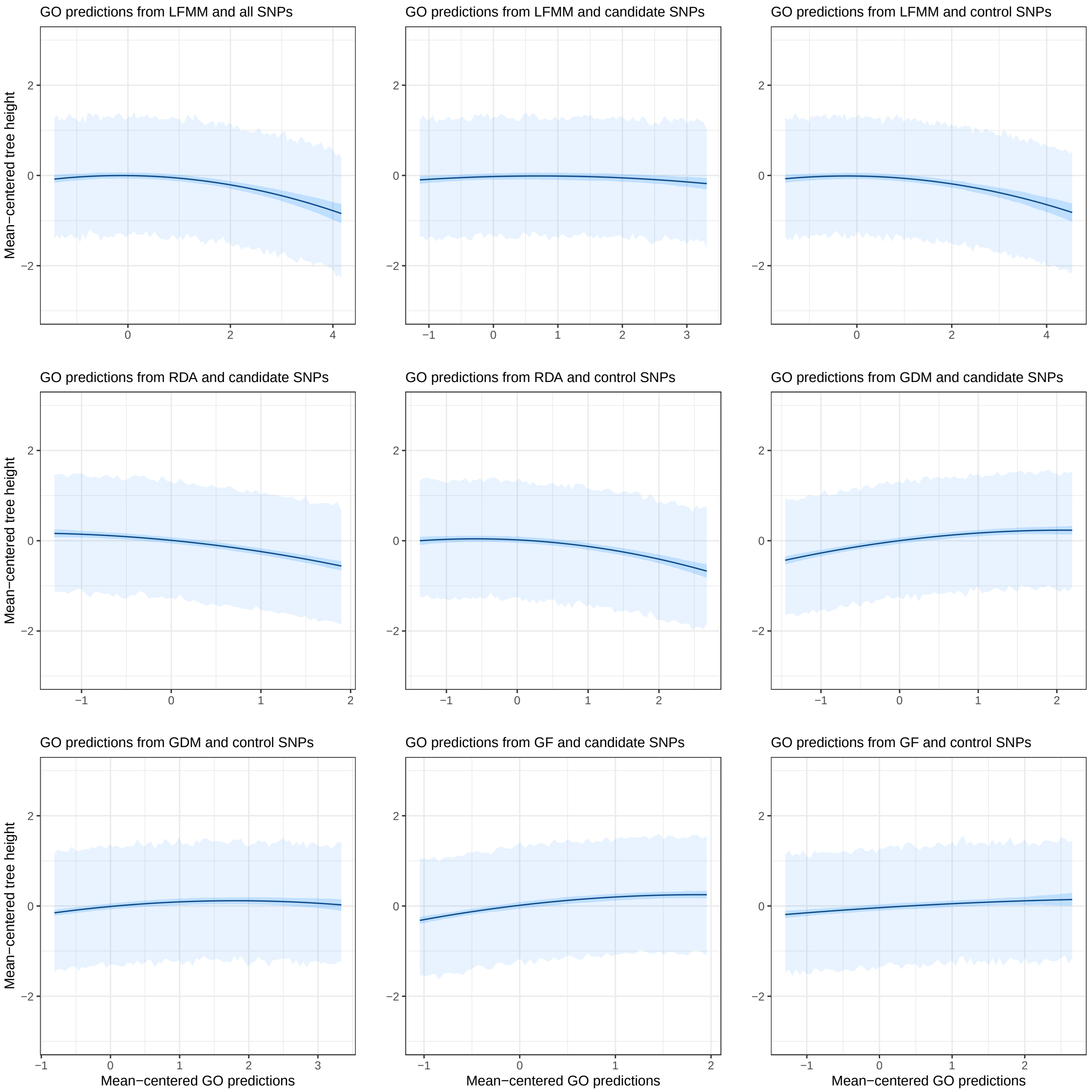

Figure 118: Tree height predictions as a function of GO predictions in Madrid (Spain).

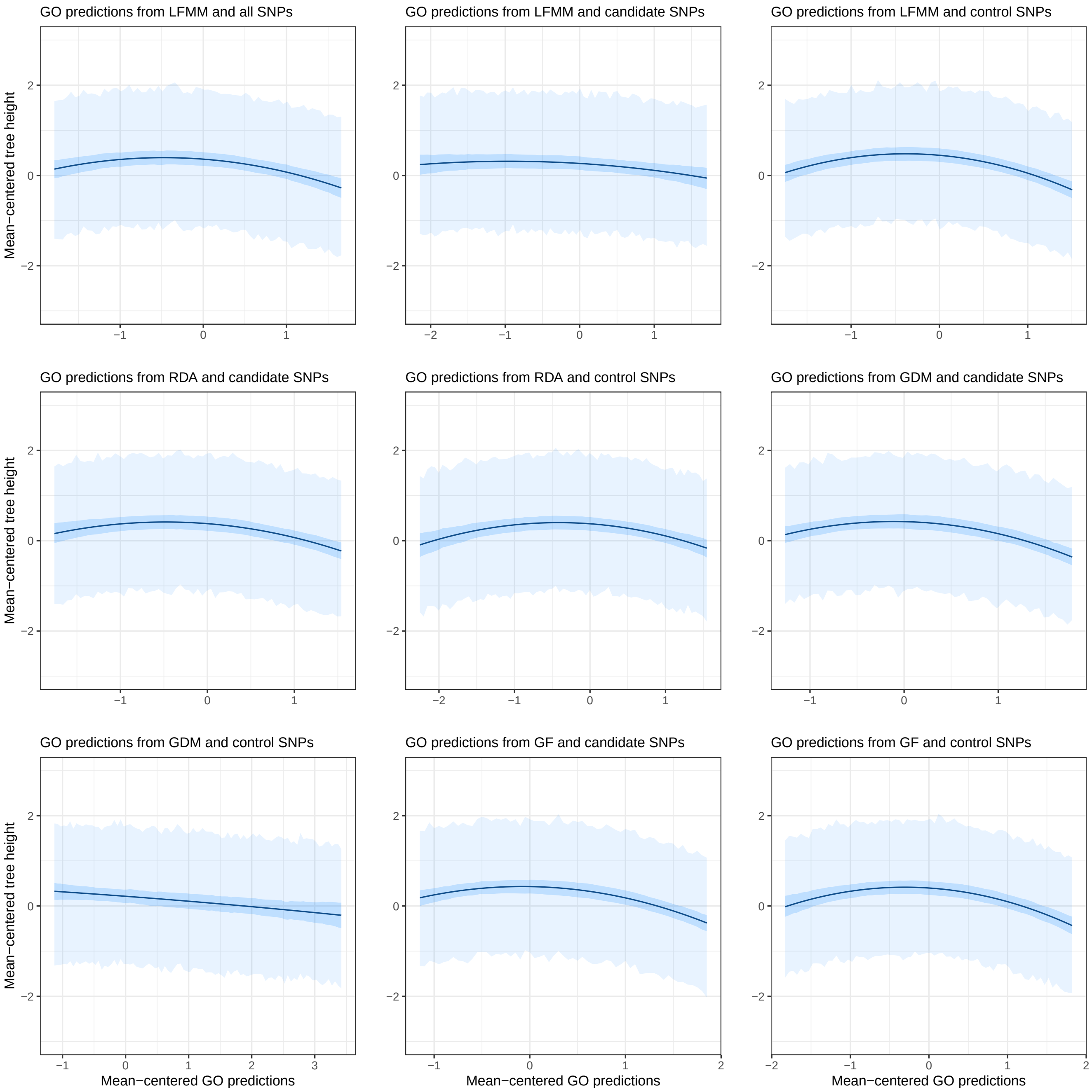

Figure 119: Tree height predictions as a function of GO predictions in Fundão (Portugal).

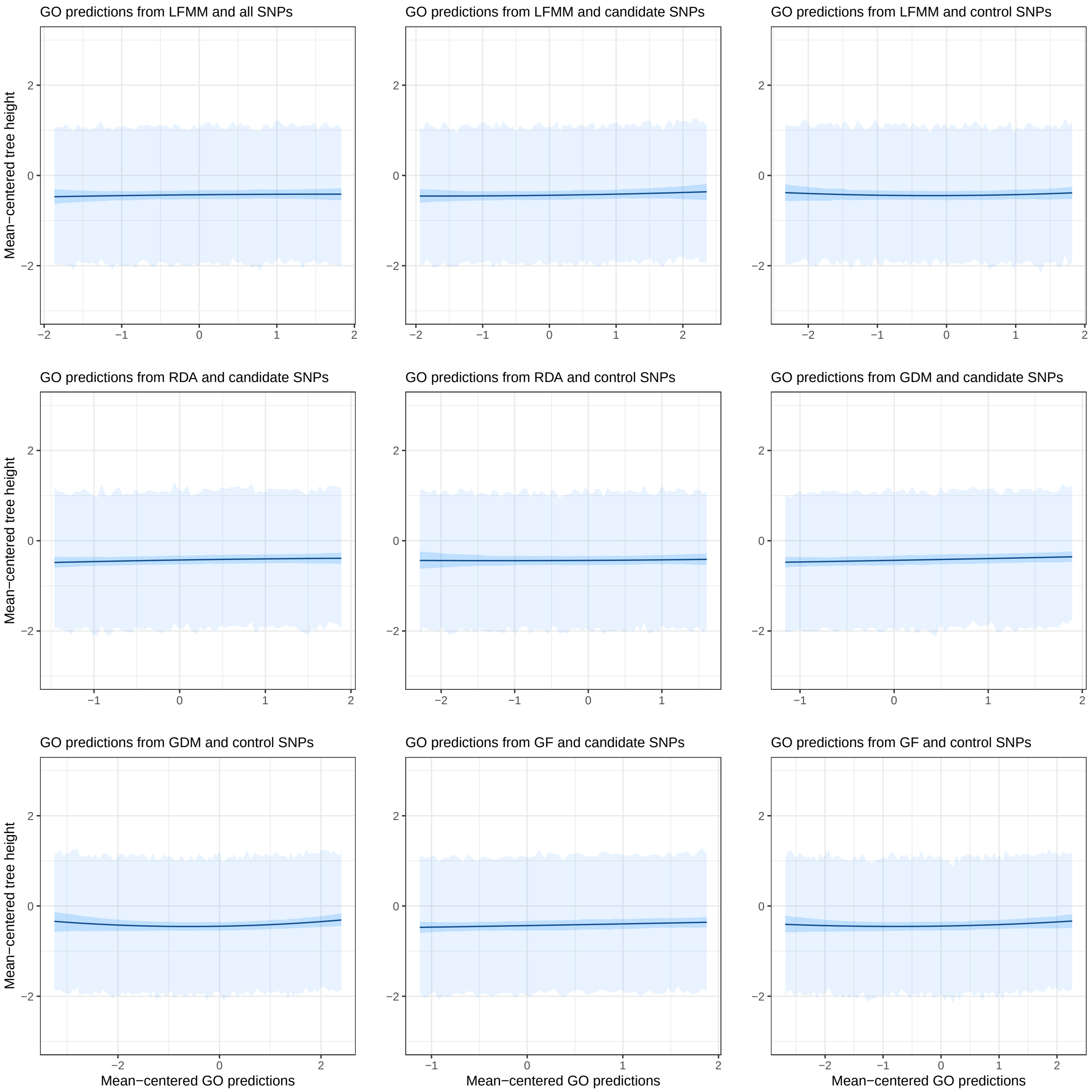

##### 9.5.3.2 Plots with CTDs as covariates

Figure 120: Tree height predictions as a function of CTDs in Asturias (Spain).

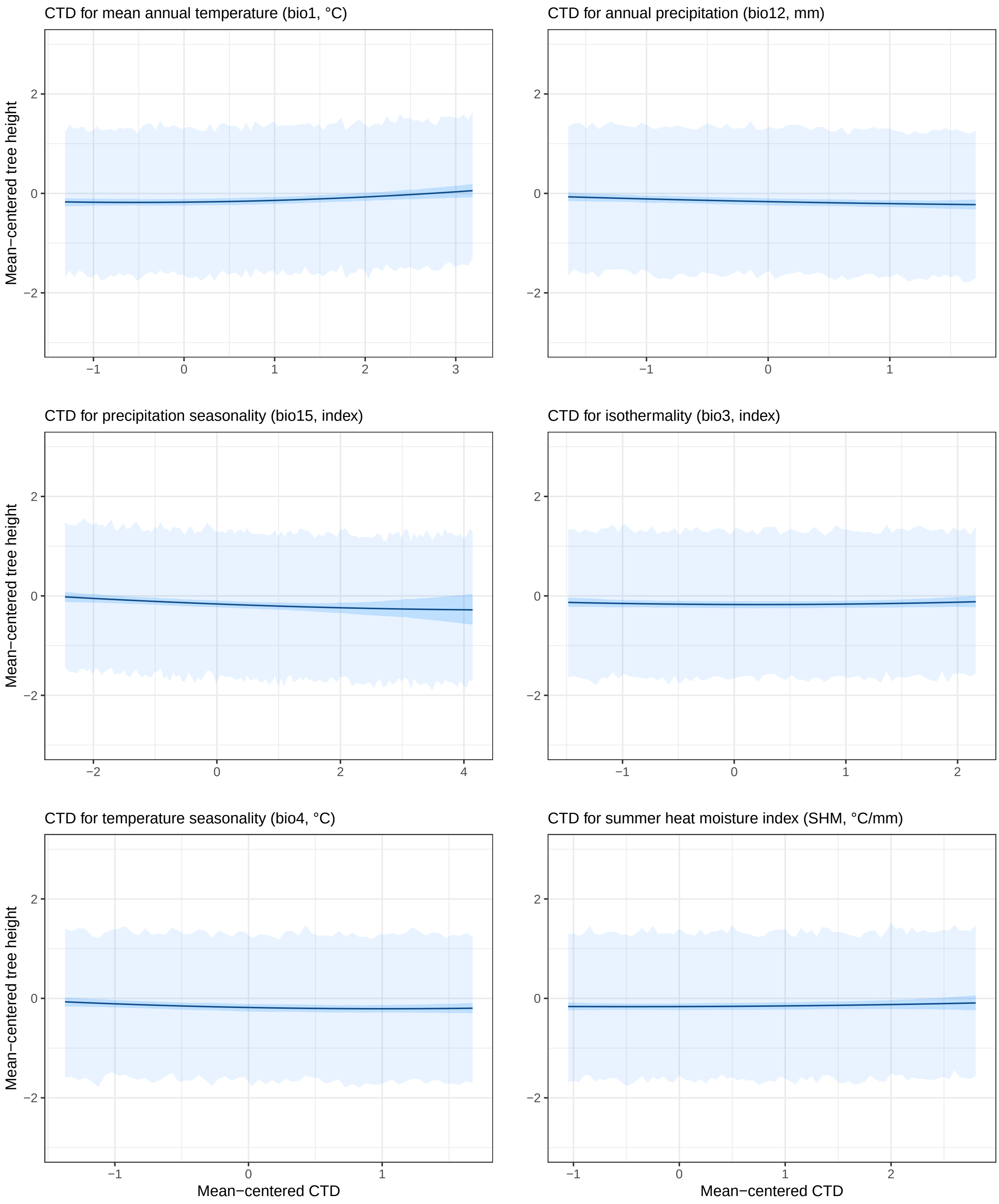

Figure 121: Tree height predictions as a function of CTDs in Bordeaux (France).

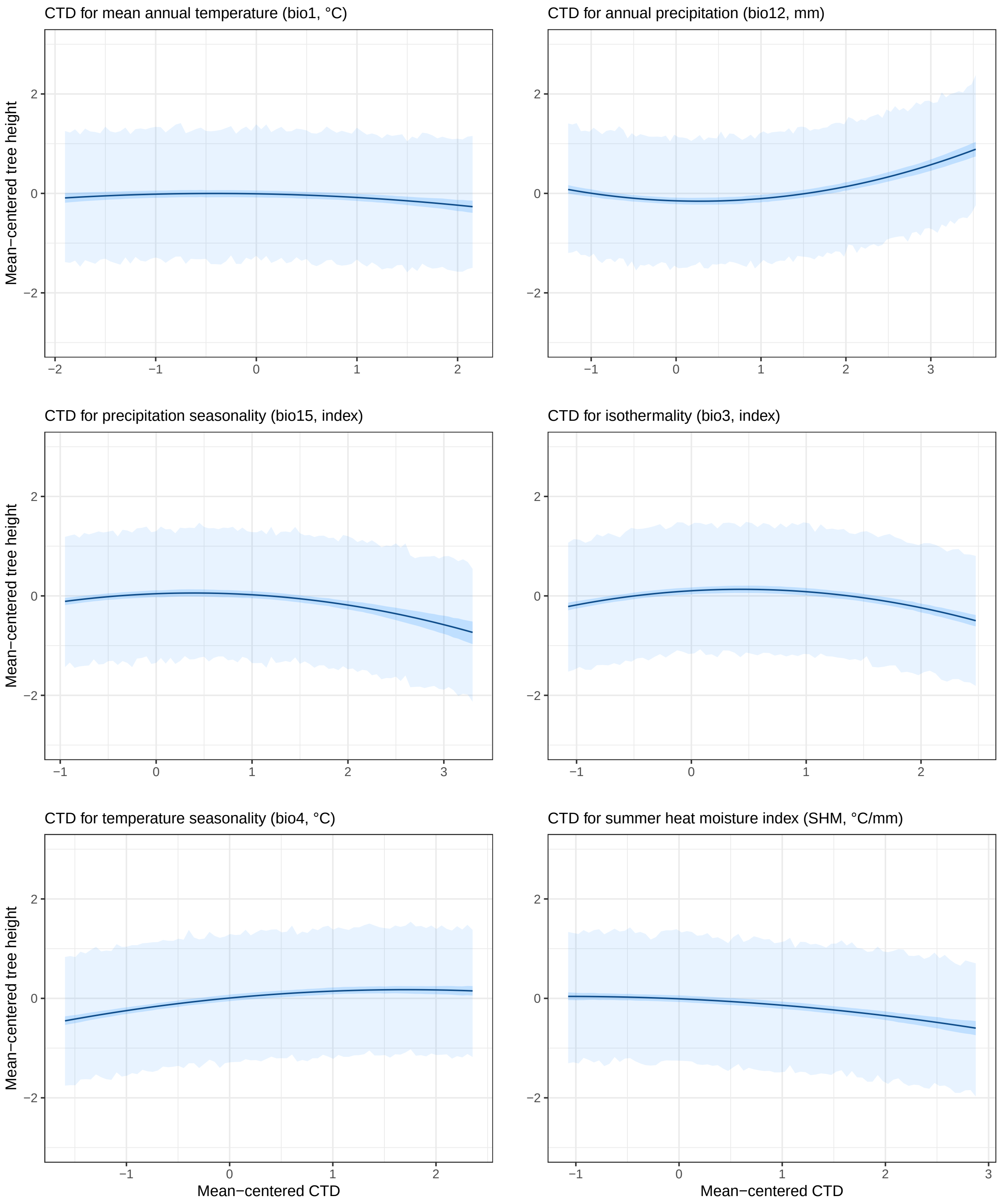

Figure 122: Tree height predictions as a function of CTDs in Cáceres (Spain).

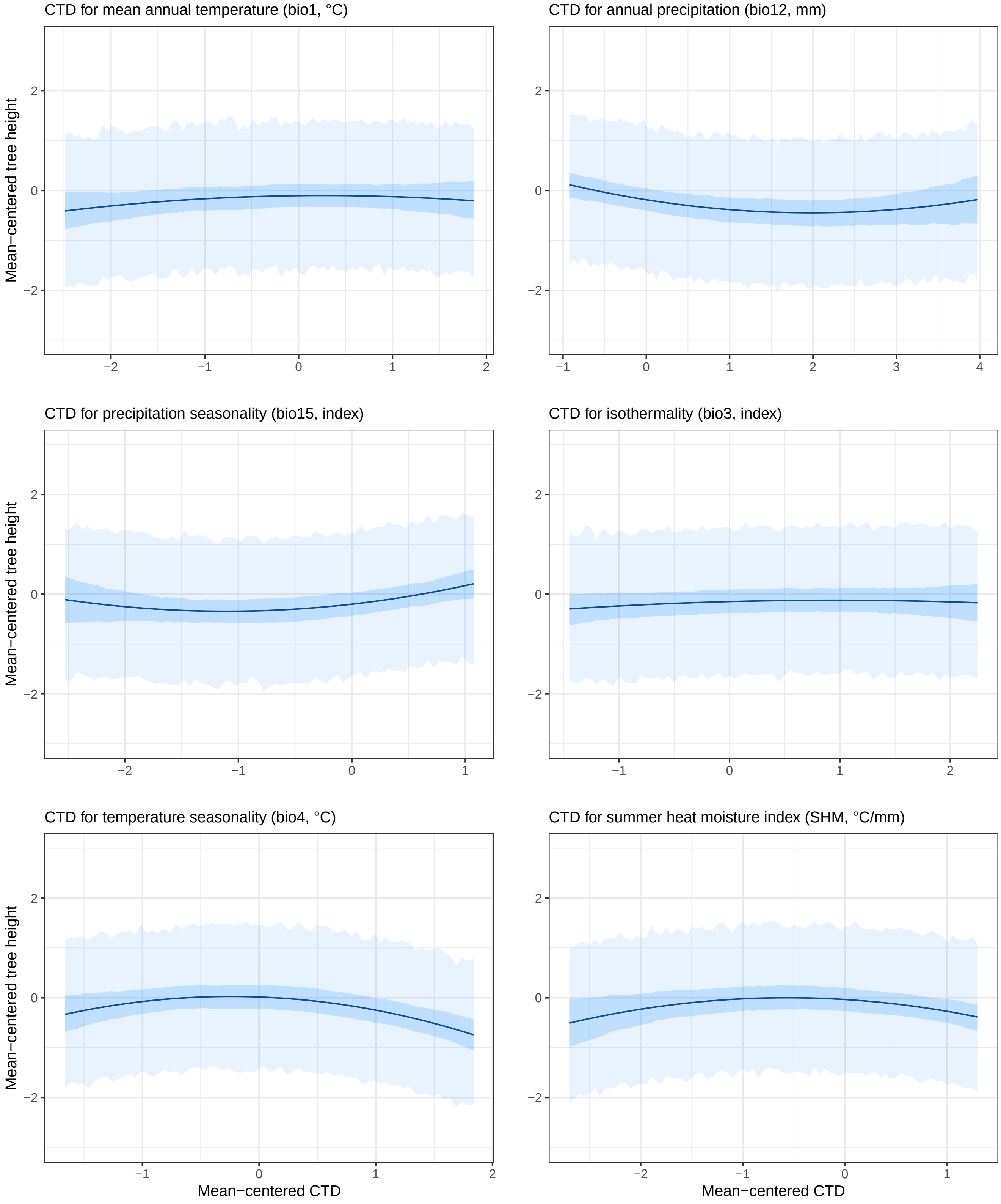

Figure 123: Tree height predictions as a function of CTDs in Madrid (Spain).

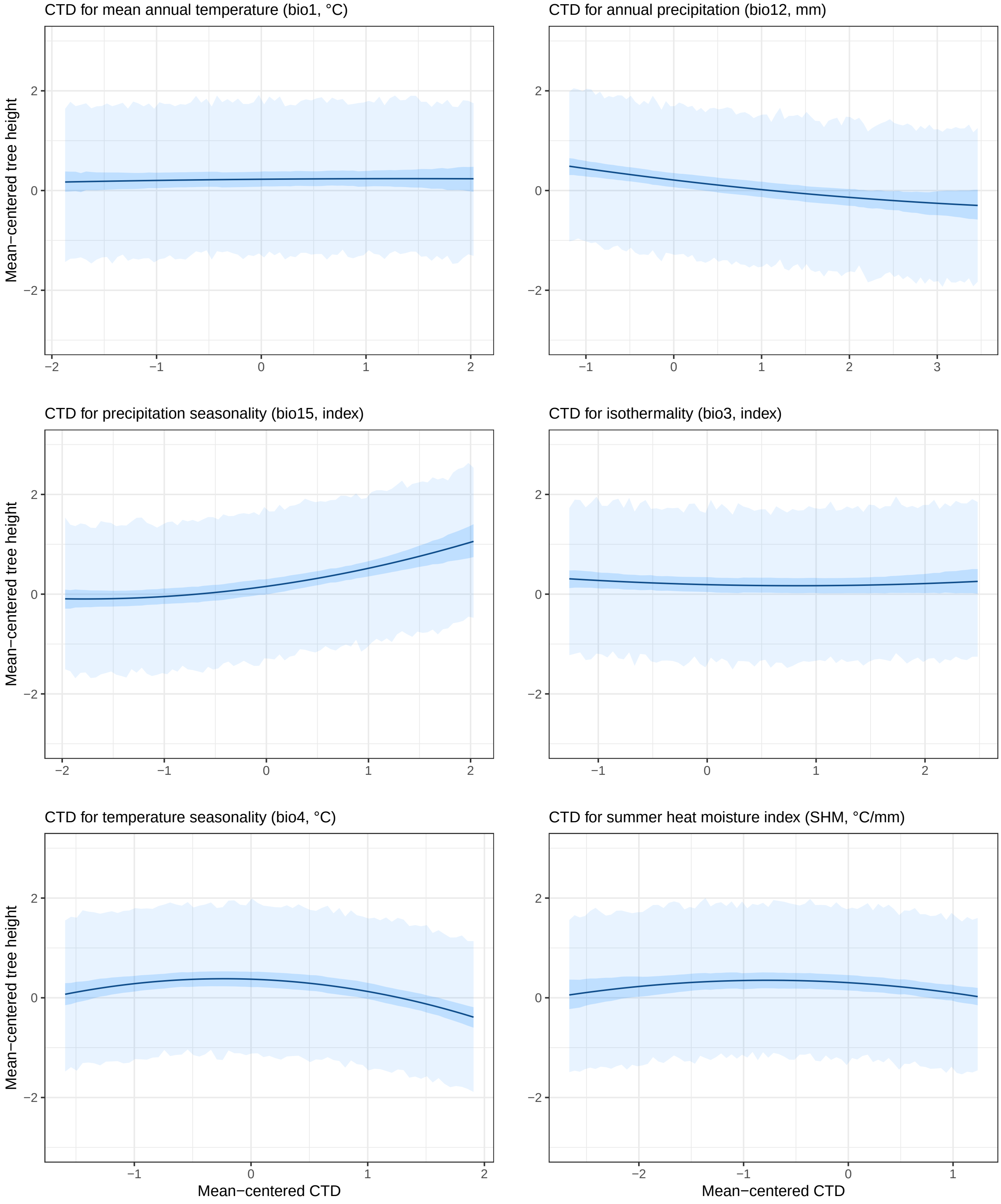

Figure 124: Tree height predictions as a function of CTDs in Fundão (Portugal).

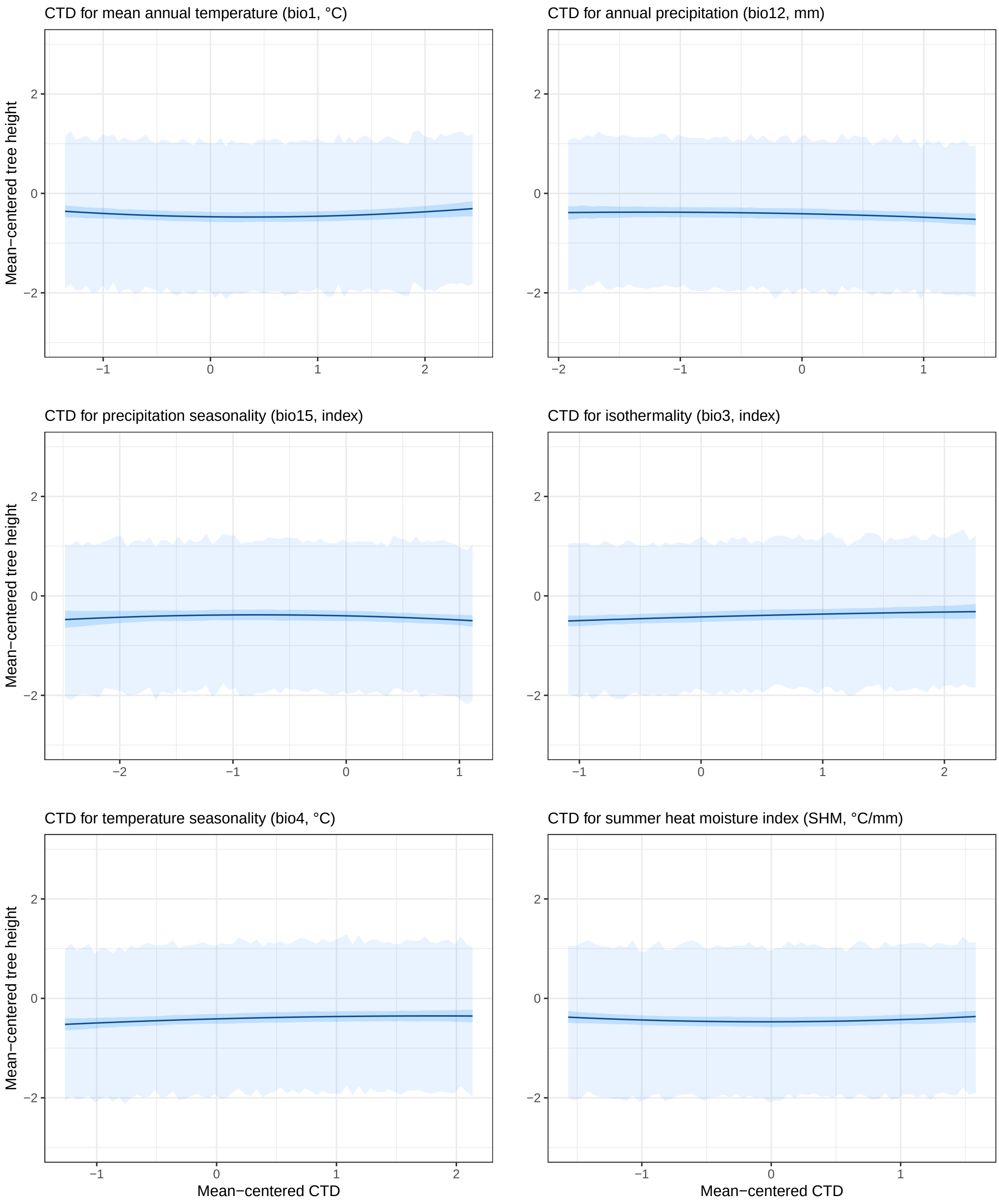

##### 9.5.4 Proportion of variance explained

Figure 125: Estimated mean and 95% credible intervals of the proportion of variance explained $R^2$. Graph titles include the time in months corresponding to the age at which height and survival were recorded. For each common garden, the brown line and uncertainty interval correspond to the estimated mean and 95% credible interval of $R^2$ in a model that does not contain GO predictions or CTDs as predictors (i.e., only including the block intercepts and the initial tree height $H_p$).

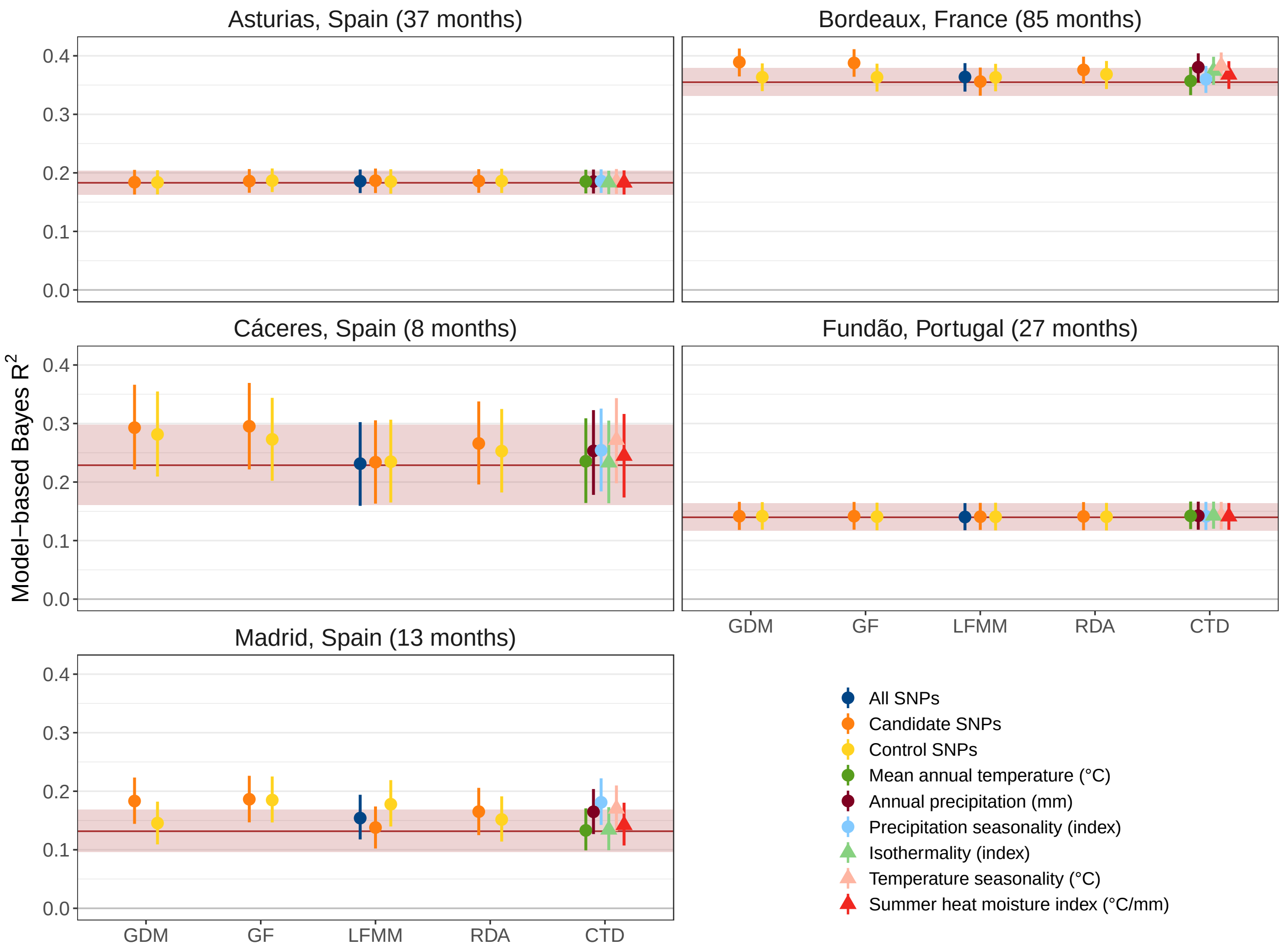

### 10 ALT and ARM populations

#### 10.1 Climate change exposure

The graph below is the same as S4, except that ALT and ARM populations are colored in pink and the other populations are colored in light blue.

Figure 126: Relative climatic distances for the six climatic variables used to make GO predictions. The relative climatic distances were calculated for five different GCMs, each corresponding to a different panel. ALT and ARM populations are colored in pink and the other populations are colored in light blue.

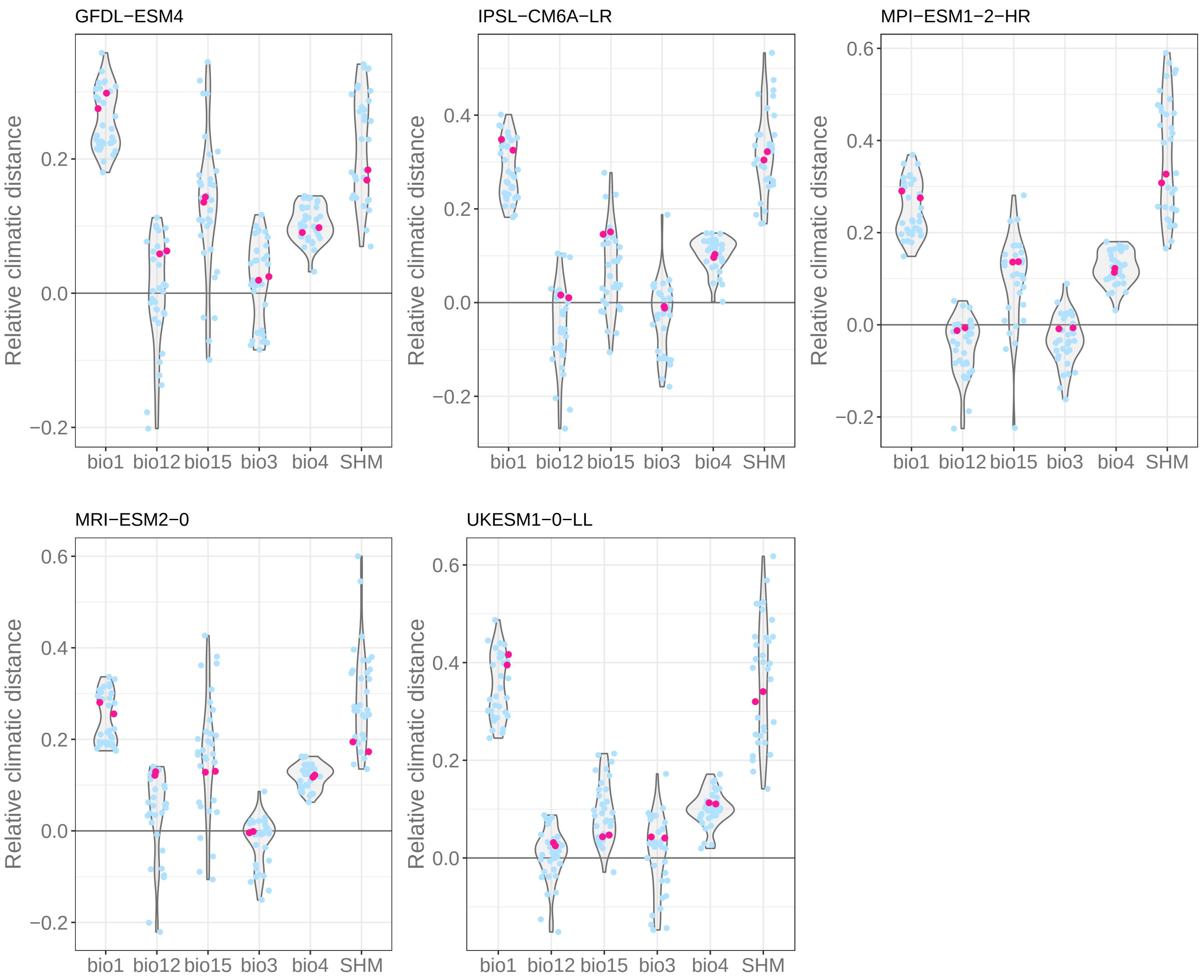

#### 10.2 Genomic composition of control and candidate SNPs

We performed a PCA on the genomic of the two sets of SNPs (i.e. control and candidate SNPs) with the function `PCA` of the `FactoMineR` R package v2.11 (Lê et al. 2008).

Below we show the scree plots of the two PCA and the PCA plots in which the ALT and ARM populations are colored in pink and the other populations in light blue.

Figure 127: Scree plots.

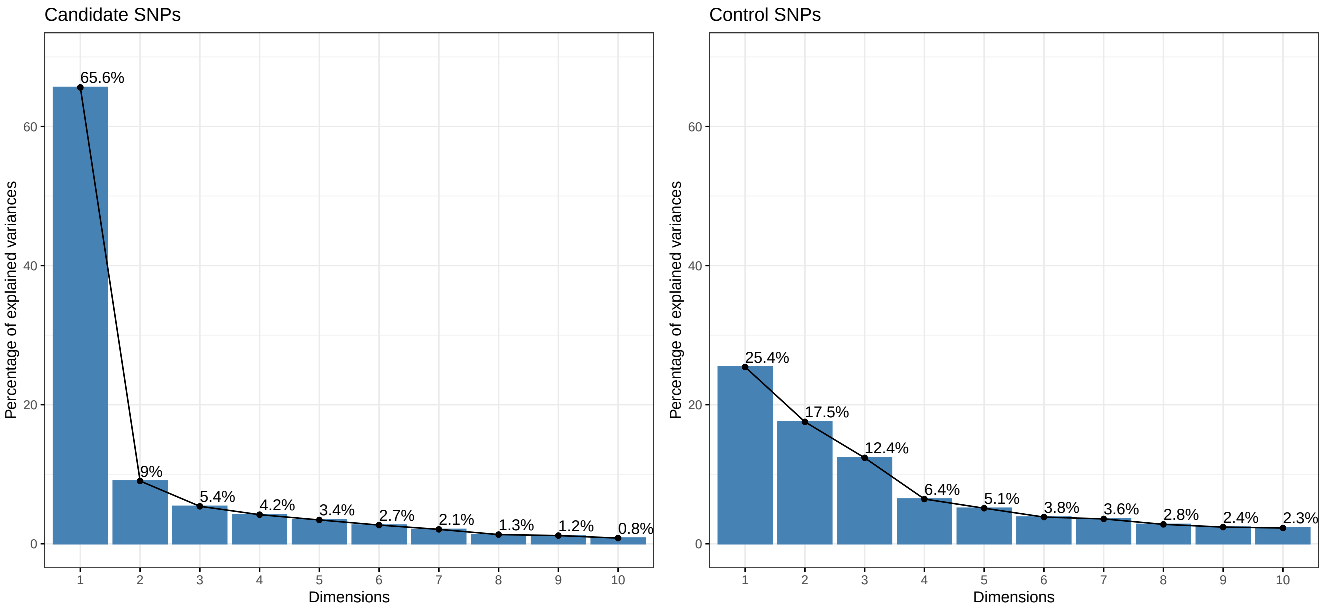

Figure 128: PCA plots.

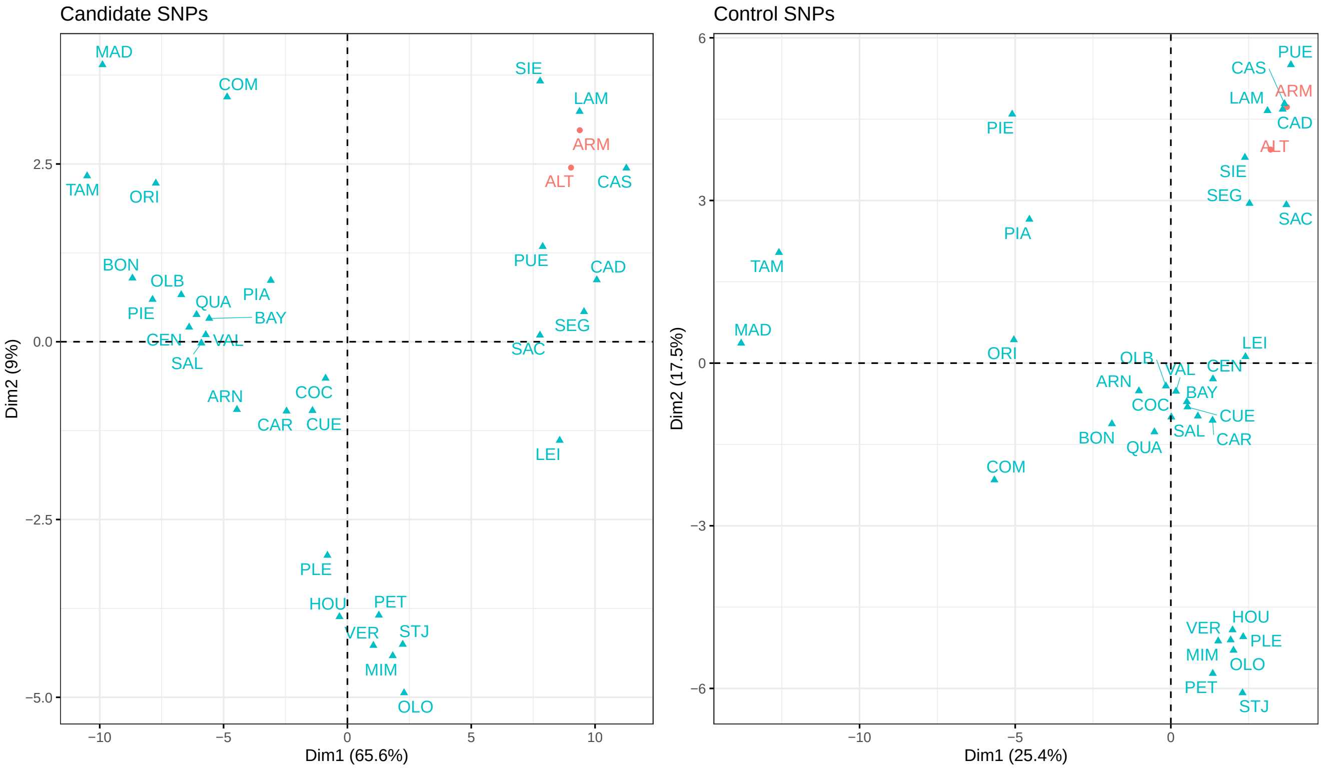
